## Supplementary tables for "Nuclear receptor corepressor 1 controls regulatory T cell subset differentiation and effector function"

### equal first-authorship

Supplementary Tables:

**Supplementary Table 1. DEG between NCOR1-cKO and WT naive Treg cells**

List of all genes that are differentially expressed between NCOR1-cKO and WT naive Treg cells. FDR <0.05.

**Supplementary Table 2. DEG between NCOR1-cKO and WT effector Treg cells**

List of all genes that are differentially expressed between NCOR1-cKO and WT effector Treg cells. FDR <0.05.

**Supplementary Table 3. GSEA hallmark gene sets enriched between NCOR1-cKO and WT Treg cells**

List of all hallmark gene sets, including Padj, NES and leading-edge genes

**Supplementary Table 4. GSEA hallmark gene sets enriched between NCOR1-cKO and WT effector Treg cells**

List of all hallmark gene sets, including Padj, NES and leading-edge genes

**Supplementary Table 5. Top 50 Canonical Pathways between NCOR1-cKO and WT naive Treg cells**

**Supplementary Table 6. Top 50 Canonical Pathways between NCOR1-cKO and WT effector Treg cells**

**Supplementary Table 7. Naive and effector Treg cell gene sets**

List of the 100 most DEG between naive and effector Treg cells.

**Supplementary Table 1. DEG between NCOR1-cKO and WT naive Treg cells**

| | | | | | | | |
|---|---|---|---|---|---|---|---|
| Mpeg1 | protein_coding | 30,1154959 | -8,166010159 | 1,43637112 | -5,685167324 | 1,31E-08 | 2,03E-06 |
| Plbd1 | protein_coding | 18,16401969 | -7,592756435 | 1,440789141 | -5,269859563 | 1,37E-07 | 1,60E-05 |
| Aldh1a2 | protein_coding | 8,516592316 | -7,259665541 | 1,951953383 | -3,719179773 | 2,00E-04 | 0,005709217 |
| Igkv19-93 | IG_V_gene | 5,292411122 | -7,041592648 | 2,134803546 | -3,298473371 | 9,72E-04 | 0,017236453 |
| Ttc8 | protein_coding | 39,62493535 | -7,03567853 | 1,268151291 | -5,547980416 | 2,89E-08 | 4,03E-06 |
| Rapgef3 | protein_coding | 11,73090824 | -6,964159059 | 1,517054704 | -4,590578732 | 4,42E-06 | 3,04E-04 |
| Siglecg | protein_coding | 9,707581966 | -6,929654193 | 1,769302552 | -3,916602159 | 8,98E-05 | 0,003124563 |
| Gata2 | protein_coding | 11,21507887 | -6,901087402 | 1,383948904 | -4,986518927 | 6,15E-07 | 5,73E-05 |
| Wnt10a | protein_coding | 30,86283192 | -6,883778069 | 1,415972475 | -4,861519689 | 1,16E-06 | 1,00E-04 |
| I830077J02Rik | protein_coding | 7,209777673 | -6,767879354 | 1,902301617 | -3,55773201 | 3,74E-04 | 0,009102752 |
| Nr3c2 | protein_coding | 39,72668668 | -6,741579565 | 1,486582384 | -4,53495187 | 5,76E-06 | 3,75E-04 |
| Gm37519 | TEC | 13,46871449 | -6,729456366 | 1,632076693 | -4,123247635 | 3,74E-05 | 0,001653261 |
| Spag16 | protein_coding | 9,236370652 | -6,581616804 | 1,585765741 | -4,150434478 | 3,32E-05 | 0,001521352 |
| Cd5l | protein_coding | 12,68861781 | -6,576006189 | 1,638780043 | -4,012744857 | 6,00E-05 | 0,002338118 |
| Nectin2 | protein_coding | 22,20654955 | -6,43869464 | 1,294795033 | -4,972752041 | 6,60E-07 | 6,09E-05 |
| Gm43915 | antisense | 5,331077157 | -6,430539112 | 1,751737033 | -3,670950028 | 2,42E-04 | 0,006574481 |
| Fn1 | protein_coding | 7,282306952 | -6,314488304 | 2,022695123 | -3,121819118 | 0,001797373 | 0,026165972 |
| Gm18994 | processed_pseudoge | 9,516414407 | -6,310479525 | 1,456978586 | -4,331209522 | 1,48E-05 | 7,91E-04 |
| Fndc7 | protein_coding | 10,95994715 | -6,210975848 | 1,440686052 | -4,311123746 | 1,62E-05 | 8,45E-04 |
| Gfpt2 | protein_coding | 22,59778902 | -6,114950889 | 1,241941669 | -4,923702168 | 8,49E-07 | 7,62E-05 |
| Fzd5 | protein_coding | 8,095743036 | -6,106780077 | 1,806850957 | -3,379791816 | 7,25E-04 | 0,014246269 |
| Gm33195 | lincRNA | 27,87025767 | -6,079742564 | 1,239713507 | -4,904151267 | 9,38E-07 | 8,33E-05 |
| Apc2 | protein_coding | 6,039956444 | -5,972961526 | 1,781245046 | -3,353250884 | 7,99E-04 | 0,015235562 |
| Sytl4 | protein_coding | 6,892145409 | -5,845122048 | 1,957136876 | -2,986567838 | 0,002821282 | 0,03590924 |
| Pla2g7 | protein_coding | 9,436926487 | -5,822820348 | 1,615730461 | -3,603831511 | 3,14E-04 | 0,007932695 |
| Smim10l2a | protein_coding | 104,4945146 | -5,821892039 | 1,021315991 | -5,700382732 | 1,20E-08 | 1,89E-06 |
| Gm44901 | TEC | 8,731143151 | -5,798075997 | 1,588173483 | -3,650782524 | 2,61E-04 | 0,006920945 |
| Amz1 | protein_coding | 10,20587424 | -5,797166593 | 1,679400028 | -3,451927173 | 5,57E-04 | 0,011797317 |
| Tgfbi | protein_coding | 9,581542867 | -5,779522795 | 2,037583246 | -2,836459715 | 0,004561673 | 0,049063856 |
| Cpne7 | protein_coding | 43,8234748 | -5,709818741 | 1,31381135 | -4,345995902 | 1,39E-05 | 7,54E-04 |
| Actg2 | protein_coding | 121,2679274 | -5,708570841 | 1,01755707 | -5,610074375 | 2,02E-08 | 2,97E-06 |
| Tmem132a | protein_coding | 6,23438137 | -5,705965707 | 1,949370697 | -2,927080887 | 0,003421599 | 0,040422508 |
| Gxylt2 | protein_coding | 7,104706816 | -5,638060167 | 1,5891493 | -3,547848001 | 3,88E-04 | 0,009322055 |
| Gas7 | protein_coding | 8,895483205 | -5,632531438 | 1,493541944 | -3,771257622 | 1,62E-04 | 0,005010477 |
| Glt28d2 | protein_coding | 20,17311245 | -5,562137427 | 1,422787422 | -3,909324289 | 9,26E-05 | 0,003200077 |
| Gm45453 | TEC | 5,290665882 | -5,541017962 | 1,60455038 | -3,453315042 | 5,54E-04 | 0,011751791 |
| Gm34680 | lincRNA | 19,50029962 | -5,517726402 | 1,343079175 | -4,10826592 | 3,99E-05 | 0,001743911 |
| Clec4a1 | protein_coding | 8,64457246 | -5,516367132 | 1,734325026 | -3,180699724 | 0,001469198 | 0,0227664 |
| 6030445D17Rik | protein_coding | 5,727935508 | -5,492628452 | 1,75731398 | -3,125581721 | 0,001774538 | 0,025947335 |
| Epb41l5 | protein_coding | 9,850775844 | -5,492518018 | 1,862321768 | -2,949285195 | 0,003185099 | 0,038696855 |
| Col1a1 | protein_coding | 14,08500057 | -5,486221969 | 1,445977389 | -3,79412708 | 1,48E-04 | 0,004665894 |
| Large2 | protein_coding | 8,333148795 | -5,452063571 | 1,560868519 | -3,49296786 | 4,78E-04 | 0,010643164 |
| Arhgap24 | protein_coding | 27,73284723 | -5,44789145 | 1,255858461 | -4,337982042 | 1,44E-05 | 7,72E-04 |
| Fcgr4 | protein_coding | 6,886332543 | -5,431078871 | 1,854601216 | -2,928434869 | 0,003406732 | 0,040403577 |
| Igkv10-96 | IG_V_gene | 6,846373613 | -5,424496428 | 1,636175598 | -3,315351014 | 9,15E-04 | 0,016692314 |
| Tppp | protein_coding | 13,93315424 | -5,401534603 | 1,814441742 | -2,976967779 | 0,002911146 | 0,036435303 |
| Itgad | protein_coding | 12,85706946 | -5,401513836 | 1,521052834 | -3,551167792 | 3,84E-04 | 0,009238018 |
| Slc12a5 | protein_coding | 11,02845249 | -5,393932737 | 1,458630846 | -3,697942321 | 2,17E-04 | 0,006113917 |
| Mmp14 | protein_coding | 28,82127691 | -5,366173075 | 1,439136172 | -3,728745883 | 1,92E-04 | 0,005583364 |
| Ccr10 | protein_coding | 31,66964859 | -5,359394259 | 1,415457491 | -3,786333601 | 1,53E-04 | 0,004763124 |
| Spic | protein_coding | 8,571303629 | -5,346236971 | 1,392000154 | -3,840687055 | 1,23E-04 | 0,003992491 |
| Clec4a3 | protein_coding | 7,661256142 | -5,341192261 | 1,61725707 | -3,302624153 | 9,58E-04 | 0,017076344 |
| Stkld1 | protein_coding | 7,845221249 | -5,315249129 | 1,418670989 | -3,746639756 | 1,79E-04 | 0,005344731 |
| Adgrv1 | protein_coding | 5,367032062 | -5,310276598 | 1,721039225 | -3,085505851 | 0,002032062 | 0,028507274 |
| Tspear | protein_coding | 10,66021438 | -5,224078812 | 1,664169078 | -3,139151473 | 0,001694378 | 0,025264963 |
| Olfr157 | protein_coding | 9,176442654 | -5,196926281 | 1,52555091 | -3,406589874 | 6,58E-04 | 0,013280816 |
| Gm38346 | TEC | 8,614154907 | -5,195996702 | 1,655575402 | -3,138483873 | 0,001698243 | 0,025277164 |
| Zfp449 | protein_coding | 34,40941116 | -5,111623281 | 1,054273886 | -4,84847756 | 1,24E-06 | 1,04E-04 |
| Slc4a5 | protein_coding | 7,389143401 | -5,108015044 | 1,579159563 | -3,234641491 | 0,001217956 | 0,020060149 |
| Gm37490 | TEC | 5,763922538 | -5,095251112 | 1,533680105 | -3,322238513 | 8,93E-04 | 0,016466611 |
| 4931415C17Rik | processed_transcript | 6,692903688 | -5,067442927 | 1,453966343 | -3,485254629 | 4,92E-04 | 0,010807599 |
| Zfp518b | protein_coding | 14,77550409 | -5,046858247 | 1,502006267 | -3,360078023 | 7,79E-04 | 0,015036837 |
| Phf21b | protein_coding | 8,896827231 | -5,043834246 | 1,599124001 | -3,154123284 | 0,001609811 | 0,024398555 |
| Tfr2 | protein_coding | 5,76367155 | -5,042361351 | 1,704472964 | -2,958311136 | 0,003093297 | 0,037859647 |
| Nox1 | protein_coding | 15,43993615 | -4,994708151 | 1,40439936 | -3,556472819 | 3,76E-04 | 0,009127172 |
| G530011O06Rik | processed_transcript | 11,43362074 | -4,967525484 | 1,402348783 | -3,542289581 | 3,97E-04 | 0,0094101 |
| Gm44389 | processed_pseudoge | 17,66491575 | -4,952710545 | 1,105485636 | -4,480122022 | 7,46E-06 | 4,73E-04 |

| | | | | | | | |
|---|---|---|---|---|---|---|---|
| Tex14 | protein_coding | 5,761315926 | -4,921796005 | 1,724142697 | -2,854633791 | 0,004308648 | 0,047205577 |
| Palld | protein_coding | 25,8351764 | -4,898575399 | 1,450681235 | -3,376741409 | 7,33E-04 | 0,014321367 |
| Dab2ip | protein_coding | 26,44873343 | -4,875466875 | 1,293922392 | -3,767974728 | 1,65E-04 | 0,005048661 |
| Igsf9b | protein_coding | 49,1724697 | -4,861486972 | 1,319579492 | -3,684118312 | 2,29E-04 | 0,006337287 |
| Map9 | protein_coding | 13,62597918 | -4,860037683 | 1,527358895 | -3,181988004 | 0,001462679 | 0,02270778 |
| Gm43961 | TEC | 8,532709455 | -4,841372205 | 1,323949596 | -3,656764743 | 2,55E-04 | 0,006814995 |
| Gm42429 | TEC | 6,436890819 | -4,834929667 | 1,62448973 | -2,976275921 | 0,002917722 | 0,036435303 |
| Rin2 | protein_coding | 137,2683561 | -4,826551034 | 1,094480065 | -4,409903103 | 1,03E-05 | 6,02E-04 |
| Pls3 | protein_coding | 23,48138439 | -4,76839209 | 1,468183517 | -3,247817481 | 0,001162938 | 0,01944875 |
| Svip | protein_coding | 8,484899597 | -4,749615474 | 1,529039263 | -3,106274371 | 0,001894608 | 0,027069952 |
| Hgfac | protein_coding | 45,68022394 | -4,730175232 | 1,347311128 | -3,510826217 | 4,47E-04 | 0,010128011 |
| Rnase6 | protein_coding | 14,96386981 | -4,720770253 | 1,230861599 | -3,835337992 | 1,25E-04 | 0,004048641 |
| Ripk4 | protein_coding | 9,515228206 | -4,714679009 | 1,495439076 | -3,152705506 | 0,001617649 | 0,024486298 |
| Nid2 | protein_coding | 73,34422073 | -4,702104194 | 1,328080935 | -3,540525333 | 3,99E-04 | 0,009440601 |
| Il13ra1 | protein_coding | 31,27125793 | -4,621812129 | 1,469940099 | -3,144218008 | 0,001665313 | 0,02499903 |
| Lima1 | protein_coding | 33,89896704 | -4,591152054 | 1,058277759 | -4,338324239 | 1,44E-05 | 7,72E-04 |
| Hif3a | protein_coding | 17,44690705 | -4,564291505 | 1,463860313 | -3,117982955 | 0,001820933 | 0,026393196 |
| Lonrf1 | protein_coding | 7,036325355 | -4,539287012 | 1,579302517 | -2,87423528 | 0,004050071 | 0,04535424 |
| Chdh | protein_coding | 33,0889709 | -4,534750699 | 1,335853593 | -3,394646481 | 6,87E-04 | 0,013740142 |
| Ighg3 | IG_C_gene | 27,59208925 | -4,532669454 | 1,451405451 | -3,122951929 | 0,00179047 | 0,026088357 |
| Gm16702 | lincRNA | 38,53594655 | -4,508623003 | 1,351970843 | -3,334852246 | 8,53E-04 | 0,015950239 |
| Icam5 | protein_coding | 23,30145338 | -4,456893608 | 1,359136195 | -3,279210446 | 0,00104098 | 0,018033507 |
| D830030K20Rik | protein_coding | 24,66427748 | -4,448355848 | 1,352116874 | -3,28991963 | 0,00100216 | 0,017544143 |
| Susd4 | protein_coding | 42,65453796 | -4,415558029 | 1,070336338 | -4,125392992 | 3,70E-05 | 0,001642304 |
| Itpripl2 | protein_coding | 14,06685392 | -4,395700044 | 1,335637331 | -3,291088038 | 9,98E-04 | 0,017526899 |
| Gsta3 | protein_coding | 6,191414263 | -4,368261759 | 1,363975537 | -3,202595383 | 0,001361952 | 0,021567704 |
| Pianp | protein_coding | 18,0937485 | -4,362671001 | 1,183632996 | -3,68583084 | 2,28E-04 | 0,006312851 |
| Rgs9 | protein_coding | 109,6593547 | -4,353568165 | 0,786468954 | -5,53558808 | 3,10E-08 | 4,29E-06 |
| Gna14 | protein_coding | 34,34601028 | -4,327863477 | 1,429012908 | -3,0285685 | 0,002457154 | 0,032571024 |
| 2010300C02Rik | protein_coding | 7,948815669 | -4,277280287 | 1,293200754 | -3,307514533 | 9,41E-04 | 0,016897686 |
| Kdf1 | protein_coding | 12,53952373 | -4,139799383 | 1,457865447 | -2,839630635 | 0,00451658 | 0,048905778 |
| Armc2 | protein_coding | 30,44670458 | -4,104060352 | 1,297746275 | -3,162452039 | 0,001564465 | 0,023863844 |
| Clec4n | protein_coding | 11,26657464 | -4,067134451 | 1,336700382 | -3,042667232 | 0,002344915 | 0,031537911 |
| Itgb8 | protein_coding | 270,0887792 | -4,054223791 | 0,495161511 | -8,187679584 | 2,66E-16 | 2,01E-13 |
| Ticam2 | protein_coding | 15,61254363 | -4,023377855 | 1,174605354 | -3,425301819 | 6,14E-04 | 0,01267647 |
| Zfp112 | protein_coding | 18,09394217 | -3,971429554 | 1,208698384 | -3,285707672 | 0,001017265 | 0,017771088 |
| Ccdc87 | protein_coding | 9,631285351 | -3,8833501 | 1,114641296 | -3,483946013 | 4,94E-04 | 0,010846213 |
| Pirb | protein_coding | 18,15349064 | -3,642575107 | 1,00522503 | -3,62364147 | 2,90E-04 | 0,007485832 |
| Gm48627 | TEC | 8,66153511 | -3,638014111 | 1,256841905 | -2,894567803 | 0,003796809 | 0,043201107 |
| Gm17764 | processed_pseudoge | 105,0493983 | -3,636802038 | 0,851087945 | -4,273121316 | 1,93E-05 | 9,66E-04 |
| Atp6v0d2 | protein_coding | 231,9557521 | -3,578184395 | 0,913218325 | -3,918213528 | 8,92E-05 | 0,003113877 |
| Serpina3f | protein_coding | 203,942006 | -3,53122948 | 0,57488825 | -6,14246243 | 8,13E-10 | 1,90E-07 |
| H2-Q1 | protein_coding | 501,8803359 | -3,521425161 | 0,861158535 | -4,08917176 | 4,33E-05 | 0,001833232 |
| St3gal5 | protein_coding | 34,37937237 | -3,519048345 | 0,874013942 | -4,026306878 | 5,67E-05 | 0,002246054 |
| Gprasp2 | protein_coding | 125,7088543 | -3,505042742 | 0,534906753 | -6,552623837 | 5,65E-11 | 1,59E-08 |
| Mctp1 | protein_coding | 121,0029531 | -3,489155141 | 0,453636727 | -7,691518199 | 1,45E-14 | 7,54E-12 |
| St14 | protein_coding | 177,4843302 | -3,459792028 | 0,580376544 | -5,961288517 | 2,50E-09 | 4,58E-07 |
| Gm43149 | sense_intronic | 7,5856085 | -3,342093616 | 1,175806404 | -2,84238426 | 0,004477749 | 0,048633976 |
| Gm17228 | processed_pseudoge | 133,685409 | -3,283473383 | 0,899991568 | -3,648337939 | 2,64E-04 | 0,006960132 |
| Chadl | protein_coding | 32,09212744 | -3,209021983 | 1,001775853 | -3,203333335 | 0,001358466 | 0,021533055 |
| Camk2a | protein_coding | 34,7629664 | -3,188948923 | 0,720488243 | -4,426094323 | 9,60E-06 | 5,67E-04 |
| Plcd1 | protein_coding | 53,92470817 | -3,166303896 | 1,028548782 | -3,078418789 | 0,002081022 | 0,028973698 |
| Gm5388 | processed_pseudoge | 12,70439493 | -3,086278099 | 0,960098923 | -3,214541778 | 0,001306529 | 0,02101081 |
| Dpt | protein_coding | 12,13376516 | -3,055085503 | 0,940776606 | -3,247408026 | 0,001164613 | 0,01944875 |
| Ppic | protein_coding | 30,04646451 | -3,020593241 | 0,738662423 | -4,08927427 | 4,33E-05 | 0,001833232 |
| Akap2 | protein_coding | 9,425581737 | -3,010144858 | 0,984741891 | -3,056785627 | 0,002237242 | 0,030458795 |
| Gm42666 | processed_pseudoge | 27,95257429 | -2,981675174 | 0,812663404 | -3,669016173 | 2,43E-04 | 0,006606801 |
| Itga7 | protein_coding | 64,199124 | -2,967226169 | 0,636486947 | -4,661880632 | 3,13E-06 | 2,27E-04 |
| Ankrd6 | protein_coding | 174,7750014 | -2,911604402 | 0,515813141 | -5,6446883 | 1,65E-08 | 2,51E-06 |
| Nhsl2 | protein_coding | 42,82487994 | -2,810407912 | 0,803277054 | -3,498678194 | 4,68E-04 | 0,010472066 |
| Myo3b | protein_coding | 108,1803145 | -2,771219077 | 0,530247909 | -5,226270641 | 1,73E-07 | 1,95E-05 |
| Col9a3 | protein_coding | 40,08662975 | -2,766278822 | 0,92505457 | -2,990395282 | 0,002786166 | 0,035825615 |
| A730091E23Rik | TEC | 14,63844193 | -2,756574329 | 0,930957753 | -2,961009048 | 0,003066329 | 0,037667506 |
| Gm10499 | transcribed_unproce | 75,63349638 | -2,72798179 | 0,710230704 | -3,840979806 | 1,23E-04 | 0,003992491 |
| Xk | protein_coding | 56,36077822 | -2,713772772 | 0,612008995 | -4,434204065 | 9,24E-06 | 5,52E-04 |
| H2-Q4 | protein_coding | 4343,468833 | -2,708958031 | 0,680165174 | -3,982794378 | 6,81E-05 | 0,002557349 |
| B4galt4 | protein_coding | 88,45974291 | -2,708233415 | 0,582334419 | -4,650649739 | 3,31E-06 | 2,38E-04 |
| Samd11 | protein_coding | 55,80620711 | -2,682088998 | 0,897979633 | -2,98680382 | 0,002819106 | 0,03590924 |
| Gm4841 | protein_coding | 33,90850804 | -2,671386573 | 0,904483087 | -2,953495329 | 0,003141974 | 0,038341319 |

| | | | | | | | |
|---|---|---|---|---|---|---|---|
| Fam43a | protein_coding | 76,90546311 | -2,635110246 | 0,73057409 | -3,606903504 | 3,10E-04 | 0,007851373 |
| H2-Q2 | protein_coding | 1153,199084 | -2,612467301 | 0,641902427 | -4,069882262 | 4,70E-05 | 0,001946694 |
| Pmaip1 | protein_coding | 208,0085699 | -2,602757049 | 0,711573593 | -3,657748229 | 2,54E-04 | 0,006811776 |
| Gm47692 | processed_pseudoge | 20,65156192 | -2,598378027 | 0,828925144 | -3,13463531 | 0,001720679 | 0,025474028 |
| Tspan9 | protein_coding | 53,47979283 | -2,558111951 | 0,569400793 | -4,492638548 | 7,03E-06 | 4,49E-04 |
| Pstpip2 | protein_coding | 77,3299837 | -2,526169589 | 0,322216077 | -7,839986184 | 4,51E-15 | 2,67E-12 |
| Gm7030 | protein_coding | 709,8098286 | -2,495515051 | 0,64847702 | -3,848270602 | 1,19E-04 | 0,003901523 |
| Gbp2b | protein_coding | 249,7258087 | -2,482103136 | 0,454937109 | -5,455925855 | 4,87E-08 | 6,32E-06 |
| Gm28112 | sense_intronic | 161,0539964 | -2,477730191 | 0,425560504 | -5,822274782 | 5,81E-09 | 9,83E-07 |
| Sh3tc1 | protein_coding | 89,42634282 | -2,473007216 | 0,701977664 | -3,522914391 | 4,27E-04 | 0,009852097 |
| Gas2l1 | protein_coding | 52,07878128 | -2,445997191 | 0,750202021 | -3,26045135 | 0,00111235 | 0,018972432 |
| Gm3194 | protein_coding | 53,49244999 | -2,439501781 | 0,551927587 | -4,419967108 | 9,87E-06 | 5,77E-04 |
| Gm3500 | protein_coding | 48,75876491 | -2,363805122 | 0,699524586 | -3,37915946 | 7,27E-04 | 0,014246269 |
| Bicd1 | protein_coding | 130,5612617 | -2,340060145 | 0,546608557 | -4,281052895 | 1,86E-05 | 9,38E-04 |
| Rhod | protein_coding | 65,73781196 | -2,326523675 | 0,529963692 | -4,389968049 | 1,13E-05 | 6,45E-04 |
| Pnck | protein_coding | 28,98968127 | -2,316599066 | 0,787624621 | -2,941247649 | 0,003268931 | 0,039398093 |
| Vcam1 | protein_coding | 65,34775111 | -2,307481719 | 0,553217723 | -4,171019152 | 3,03E-05 | 0,001421715 |
| Src | protein_coding | 154,889981 | -2,28514376 | 0,340334253 | -6,714410145 | 1,89E-11 | 6,14E-09 |
| Cd80 | protein_coding | 70,98868117 | -2,271515448 | 0,475381903 | -4,778296006 | 1,77E-06 | 1,39E-04 |
| Dclk2 | protein_coding | 234,4150548 | -2,269089147 | 0,487755279 | -4,652105767 | 3,29E-06 | 2,37E-04 |
| Bcas1 | protein_coding | 51,07619762 | -2,262924585 | 0,749238207 | -3,02030057 | 0,002525239 | 0,033234635 |
| Serpinc1 | protein_coding | 93,64706239 | -2,252653969 | 0,49712253 | -4,531385792 | 5,86E-06 | 3,80E-04 |
| Adam4 | protein_coding | 16,8964445 | -2,222341973 | 0,74189478 | -2,995494822 | 0,00274 | 0,035415134 |
| Plcb4 | protein_coding | 195,3872577 | -2,219630599 | 0,372385309 | -5,960575097 | 2,51E-09 | 4,58E-07 |
| Gm46519 | lincRNA | 109,3491263 | -2,218697596 | 0,727219817 | -3,050931155 | 0,002281329 | 0,030831378 |
| Capg | protein_coding | 5963,728336 | -2,146516884 | 0,166332938 | -12,90494182 | 4,22E-38 | 1,44E-34 |
| Plcl1 | protein_coding | 53,65547201 | -2,092871238 | 0,615578898 | -3,399842401 | 6,74E-04 | 0,013530597 |
| Rnf227 | protein_coding | 67,54378493 | -2,061682747 | 0,466214308 | -4,422178198 | 9,77E-06 | 5,73E-04 |
| Sccpdh | protein_coding | 201,4835929 | -2,055355517 | 0,41925228 | -4,902431336 | 9,47E-07 | 8,36E-05 |
| Gpr83 | protein_coding | 3106,526949 | -2,051039581 | 0,232477987 | -8,822510946 | 1,12E-18 | 1,03E-15 |
| Gm28942 | lincRNA | 147,1187839 | -2,045196932 | 0,513885326 | -3,979870272 | 6,90E-05 | 0,002571555 |
| BC064078 | transcribed_unproce | 179,8746493 | -2,041241294 | 0,390620498 | -5,225637933 | 1,74E-07 | 1,95E-05 |
| Gm4017 | processed_pseudoge | 20,9983662 | -2,032784765 | 0,619248393 | -3,28266458 | 0,001028309 | 0,017891393 |
| Gm3752 | protein_coding | 43,73961408 | -1,996852414 | 0,622219284 | -3,209242246 | 0,001330853 | 0,021196582 |
| Itgae | protein_coding | 1485,255132 | -1,955999859 | 0,275119514 | -7,109636936 | 1,16E-12 | 4,95E-10 |
| Rasip1 | protein_coding | 57,59165617 | -1,887307232 | 0,374066238 | -5,045382456 | 4,53E-07 | 4,50E-05 |
| Slc9a7 | protein_coding | 199,3742418 | -1,876825176 | 0,486750313 | -3,855827366 | 1,15E-04 | 0,003806451 |
| Otud1 | protein_coding | 50,00158261 | -1,87535881 | 0,53690336 | -3,49291688 | 4,78E-04 | 0,010643164 |
| Naip2 | protein_coding | 128,8712643 | -1,847374057 | 0,416548249 | -4,434958165 | 9,21E-06 | 5,52E-04 |
| Pi4k2b | protein_coding | 46,42514703 | -1,839024425 | 0,524030399 | -3,509385008 | 4,49E-04 | 0,010169165 |
| Mtmr7 | protein_coding | 81,24728201 | -1,830744794 | 0,491229047 | -3,726865922 | 1,94E-04 | 0,005615285 |
| Gm11127 | protein_coding | 71,2384828 | -1,818371417 | 0,549727758 | -3,307767145 | 9,40E-04 | 0,016897686 |
| Gm14336 | processed_pseudoge | 28,26160202 | -1,803904034 | 0,63161248 | -2,85602975 | 0,004289749 | 0,047085106 |
| H2-Bl | polymorphic_pseudo | 961,037384 | -1,781823878 | 0,542945185 | -3,281774898 | 0,001031559 | 0,017918649 |
| Gm8543 | processed_pseudoge | 646,2274852 | -1,769306906 | 0,562883312 | -3,143292524 | 0,001670588 | 0,025045238 |
| Bcl2l14 | protein_coding | 46,46321781 | -1,741278082 | 0,595854094 | -2,922322932 | 0,003474311 | 0,040864399 |
| Irf6 | protein_coding | 359,915028 | -1,700200326 | 0,221760546 | -7,666829636 | 1,76E-14 | 8,81E-12 |
| Gm3383 | protein_coding | 42,92584297 | -1,683920623 | 0,508852453 | -3,309251261 | 9,35E-04 | 0,016856535 |
| Ubtd2 | protein_coding | 31,31417987 | -1,675096057 | 0,494160896 | -3,389778658 | 6,99E-04 | 0,013902693 |
| Gbp3 | protein_coding | 1312,986171 | -1,668593591 | 0,12106396 | -13,78274416 | 3,24E-43 | 2,69E-39 |
| Rps15a-ps7 | processed_pseudoge | 66,51303053 | -1,657243454 | 0,520024318 | -3,186857609 | 0,001438276 | 0,022403793 |
| Zfp839 | protein_coding | 53,8863669 | -1,626336973 | 0,443657014 | -3,665752873 | 2,47E-04 | 0,006676627 |
| Actg-ps1 | processed_pseudoge | 2174,581278 | -1,619461538 | 0,477955938 | -3,388307185 | 7,03E-04 | 0,013956406 |
| Gprasp1 | protein_coding | 583,211322 | -1,585979522 | 0,157600226 | -10,06330738 | 8,03E-24 | 1,66E-20 |
| H2-Q5 | polymorphic_pseudo | 11954,50182 | -1,578514876 | 0,4385181 | -3,599657295 | 3,19E-04 | 0,008036622 |
| B3gnt5 | protein_coding | 188,4720264 | -1,573199196 | 0,381304149 | -4,125838126 | 3,69E-05 | 0,001642304 |
| Endod1 | protein_coding | 874,1624837 | -1,571229317 | 0,238982588 | -6,574660228 | 4,88E-11 | 1,40E-08 |
| Myo1e | protein_coding | 722,6376001 | -1,563515493 | 0,360339906 | -4,339001777 | 1,43E-05 | 7,72E-04 |
| Entpd1 | protein_coding | 674,7113001 | -1,557282866 | 0,271605696 | -5,733616364 | 9,83E-09 | 1,58E-06 |
| Sep.08 | protein_coding | 95,96583299 | -1,553304166 | 0,300152499 | -5,175049922 | 2,28E-07 | 2,46E-05 |
| Ppm1l | protein_coding | 275,3729939 | -1,547428085 | 0,250961123 | -6,166007177 | 7,00E-10 | 1,68E-07 |
| Igsf3 | protein_coding | 85,55886022 | -1,544725152 | 0,396278432 | -3,89808031 | 9,70E-05 | 0,003283917 |
| 2510009E07Rik | protein_coding | 96,0663001 | -1,534718708 | 0,389422544 | -3,941011457 | 8,11E-05 | 0,002908074 |
| Cul7 | protein_coding | 110,0950292 | -1,521583016 | 0,283495298 | -5,367224886 | 8,00E-08 | 9,76E-06 |
| Scg5 | protein_coding | 41,71799561 | -1,5211575 | 0,493555758 | -3,082037794 | 0,002055887 | 0,028720123 |
| Patl2 | protein_coding | 46,9138595 | -1,50479588 | 0,424330811 | -3,546280025 | 3,91E-04 | 0,00934328 |
| H2-T-ps | unprocessed_pseudo | 2784,208512 | -1,499336753 | 0,452061412 | -3,31666608 | 9,11E-04 | 0,016668903 |
| Apol9b | protein_coding | 514,0177649 | -1,485993187 | 0,243534124 | -6,101786328 | 1,05E-09 | 2,32E-07 |
| Tmem121b | protein_coding | 59,60314713 | -1,448083286 | 0,377404059 | -3,836957373 | 1,25E-04 | 0,004034501 |

| | | | | | | | |
|---|---|---|---|---|---|---|---|
| Gm17241 | processed_pseudoge | 2273,234644 | -1,439172109 | 0,470138414 | -3,06116681 | 0,002204762 | 0,030264875 |
| Gsta4 | protein_coding | 208,4569194 | -1,432295297 | 0,242225297 | -5,913070652 | 3,36E-09 | 6,06E-07 |
| H2-Q7 | protein_coding | 38088,38018 | -1,430660703 | 0,429088789 | -3,334183364 | 8,56E-04 | 0,015970661 |
| Hbegf | protein_coding | 40,31301211 | -1,430487332 | 0,478628648 | -2,98872067 | 0,002801481 | 0,03590924 |
| Epcam | protein_coding | 425,591781 | -1,427050867 | 0,363094415 | -3,930247366 | 8,49E-05 | 0,003009213 |
| Gm8399 | processed_pseudoge | 8218,087193 | -1,421548175 | 0,393793529 | -3,609882011 | 3,06E-04 | 0,00778554 |
| Plxna3 | protein_coding | 90,19303086 | -1,398395546 | 0,4335494 | -3,225458382 | 0,001257711 | 0,020483783 |
| Gm1821 | transcribed_process | 29698,3329 | -1,397745496 | 0,399017105 | -3,502971375 | 4,60E-04 | 0,01037473 |
| H2-K1 | protein_coding | 169234,2267 | -1,394143015 | 0,388505023 | -3,588481312 | 3,33E-04 | 0,008350972 |
| Ubc | protein_coding | 38750,99354 | -1,385640353 | 0,383483254 | -3,613300811 | 3,02E-04 | 0,007707163 |
| H2-D1 | protein_coding | 144607,3112 | -1,385477816 | 0,398206983 | -3,479290607 | 5,03E-04 | 0,010935156 |
| Gm8797 | protein_coding | 4679,278775 | -1,383685129 | 0,449943269 | -3,075243535 | 0,002103307 | 0,029186027 |
| Irf5 | protein_coding | 624,7906278 | -1,376691743 | 0,254090653 | -5,418112505 | 6,02E-08 | 7,63E-06 |
| Colq | protein_coding | 418,7795586 | -1,376274507 | 0,274347457 | -5,016538234 | 5,26E-07 | 5,05E-05 |
| Prelid2 | protein_coding | 66,7329056 | -1,37543993 | 0,416160349 | -3,305072031 | 9,50E-04 | 0,016999172 |
| Gm8464 | transcribed_process | 391,1535137 | -1,356902449 | 0,360107907 | -3,768044026 | 1,65E-04 | 0,005048661 |
| Cd38 | protein_coding | 601,6783473 | -1,356881179 | 0,254348246 | -5,3347377 | 9,57E-08 | 1,14E-05 |
| Gm26880 | lincRNA | 55,28142303 | -1,340975699 | 0,468131341 | -2,864528781 | 0,004176302 | 0,046246379 |
| Ceacam15 | protein_coding | 47,74909126 | -1,336955757 | 0,401148449 | -3,332820456 | 8,60E-04 | 0,016013073 |
| Trav7-1 | TR_V_gene | 64,19336062 | -1,332881267 | 0,326491801 | -4,082434114 | 4,46E-05 | 0,00187722 |
| 1810006J02Rik | lincRNA | 93,39181582 | -1,328876387 | 0,452842411 | -2,934522818 | 0,00334061 | 0,03997171 |
| Gm3252 | protein_coding | 110,2227434 | -1,322229819 | 0,419185172 | -3,154285761 | 0,001608915 | 0,024398555 |
| Ubb | protein_coding | 33354,08808 | -1,319753198 | 0,386491196 | -3,414704424 | 6,39E-04 | 0,012991983 |
| Ptpn5 | protein_coding | 264,0579313 | -1,31719774 | 0,437055038 | -3,013802895 | 0,002579954 | 0,033820333 |
| Shisa5 | protein_coding | 101472,2466 | -1,311178031 | 0,263220048 | -4,981300021 | 6,32E-07 | 5,86E-05 |
| Ecm1 | protein_coding | 5888,521711 | -1,310266122 | 0,215768599 | -6,072552403 | 1,26E-09 | 2,58E-07 |
| Id3 | protein_coding | 3951,235214 | -1,302086152 | 0,149668541 | -8,699798516 | 3,32E-18 | 2,76E-15 |
| H2-Q6 | protein_coding | 18921,18733 | -1,297013952 | 0,299986775 | -4,323570432 | 1,54E-05 | 8,09E-04 |
| Il2rb | protein_coding | 23130,86309 | -1,293804097 | 0,229242886 | -5,643813508 | 1,66E-08 | 2,51E-06 |
| Ccrl2 | protein_coding | 276,9329163 | -1,292739276 | 0,291444714 | -4,435624373 | 9,18E-06 | 5,52E-04 |
| Gm16385 | processed_pseudoge | 604,9677193 | -1,264109978 | 0,367614267 | -3,438685849 | 5,85E-04 | 0,012279888 |
| Foxp3 | protein_coding | 20294,56205 | -1,258690504 | 0,190792168 | -6,597181188 | 4,19E-11 | 1,22E-08 |
| Ltb | protein_coding | 29414,8596 | -1,256836084 | 0,250746736 | -5,012372665 | 5,38E-07 | 5,13E-05 |
| Gbp2 | protein_coding | 1933,244133 | -1,253569417 | 0,153306367 | -8,176890747 | 2,91E-16 | 2,10E-13 |
| H2-K2 | transcribed_unproce | 3473,035449 | -1,251498371 | 0,403317065 | -3,103013683 | 0,001915608 | 0,027312223 |
| Gtf3c4 | protein_coding | 165,0384706 | -1,231259299 | 0,348045191 | -3,537642039 | 4,04E-04 | 0,009490965 |
| Gm3468 | protein_coding | 123,6474935 | -1,225308092 | 0,329686484 | -3,716585765 | 2,02E-04 | 0,00574834 |
| Trps1 | protein_coding | 226,6500035 | -1,223384067 | 0,295049193 | -4,146373208 | 3,38E-05 | 0,001535849 |
| Gm19243 | processed_pseudoge | 220,7009765 | -1,201151838 | 0,35494776 | -3,38402428 | 7,14E-04 | 0,014096079 |
| Tbc1d4 | protein_coding | 2293,457403 | -1,198607287 | 0,13761447 | -8,709892828 | 3,04E-18 | 2,66E-15 |
| B2m | protein_coding | 128311,4647 | -1,198420425 | 0,303088565 | -3,954027181 | 7,68E-05 | 0,002800656 |
| Mdfic | protein_coding | 342,7278467 | -1,187002418 | 0,357690548 | -3,318517713 | 9,05E-04 | 0,016595373 |
| Gm17782 | unprocessed_pseudo | 1642,139625 | -1,184504742 | 0,305491533 | -3,877373388 | 1,06E-04 | 0,003540153 |
| Apol9a | protein_coding | 346,8415457 | -1,181584142 | 0,244219078 | -4,838213913 | 1,31E-06 | 1,08E-04 |
| Txnip | protein_coding | 13537,41394 | -1,180662046 | 0,264960192 | -4,455997854 | 8,35E-06 | 5,15E-04 |
| Gm12715 | processed_pseudoge | 11574,90285 | -1,180366706 | 0,301907495 | -3,909696598 | 9,24E-05 | 0,003200077 |
| Gm8909 | protein_coding | 4547,581212 | -1,179263236 | 0,298268344 | -3,953698943 | 7,70E-05 | 0,002800656 |
| Gm43584 | processed_transcript | 148,6989008 | -1,17882577 | 0,309454009 | -3,809373071 | 1,39E-04 | 0,004455006 |
| Acsbg1 | protein_coding | 1016,282445 | -1,178738373 | 0,264295198 | -4,459931093 | 8,20E-06 | 5,10E-04 |
| Ikzf2 | protein_coding | 2664,877475 | -1,170550017 | 0,090718494 | -12,90310244 | 4,32E-38 | 1,44E-34 |
| Acta1 | protein_coding | 623,7469389 | -1,169716609 | 0,328953301 | -3,555874359 | 3,77E-04 | 0,009127172 |
| Gm10406 | protein_coding | 62,13014552 | -1,157657965 | 0,386919817 | -2,991984166 | 0,002771706 | 0,035737251 |
| Zfp28 | protein_coding | 207,4642268 | -1,155106743 | 0,259493191 | -4,451395196 | 8,53E-06 | 5,22E-04 |
| Osbpl3 | protein_coding | 976,1501542 | -1,145879958 | 0,205592321 | -5,573554272 | 2,50E-08 | 3,51E-06 |
| Gm13604 | processed_pseudoge | 306,5445261 | -1,144746425 | 0,34420687 | -3,325751245 | 8,82E-04 | 0,016278588 |
| Gm49594 | processed_pseudoge | 149,4003857 | -1,135096673 | 0,371523235 | -3,055250835 | 0,002248724 | 0,030545704 |
| Gm30054 | antisense | 250,5123688 | -1,134210459 | 0,21971065 | -5,162291666 | 2,44E-07 | 2,60E-05 |
| Tnfrsf9 | protein_coding | 4705,482046 | -1,129959425 | 0,165757571 | -6,816940045 | 9,30E-12 | 3,15E-09 |
| Ndrg1 | protein_coding | 382,4756557 | -1,129235841 | 0,288222244 | -3,917934391 | 8,93E-05 | 0,003113877 |
| Tns1 | protein_coding | 332,4058337 | -1,123882257 | 0,360505435 | -3,117518204 | 0,001823807 | 0,026411778 |
| Sorbs1 | protein_coding | 272,3558964 | -1,123437854 | 0,206849848 | -5,431175626 | 5,60E-08 | 7,15E-06 |
| Gm5586 | processed_pseudoge | 1553,411573 | -1,122601477 | 0,335213946 | -3,34891042 | 8,11E-04 | 0,015370249 |
| Cd1d2 | polymorphic_pseudo | 486,1141031 | -1,121684752 | 0,172670155 | -6,496112483 | 8,24E-11 | 2,28E-08 |
| Maged2 | protein_coding | 299,0689319 | -1,11757799 | 0,383155206 | -2,916776214 | 0,003536694 | 0,041392791 |
| Rps19-ps3 | processed_pseudoge | 632,4354991 | -1,114923538 | 0,391048917 | -2,851110151 | 0,004356687 | 0,047599463 |
| Gm19246 | processed_pseudoge | 1251,853346 | -1,106102722 | 0,342788808 | -3,226776065 | 0,001251934 | 0,02042979 |
| Gm10925 | unprocessed_pseudo | 86899,88308 | -1,104236513 | 0,303254717 | -3,641283885 | 2,71E-04 | 0,00710125 |
| Lrrc32 | protein_coding | 6013,570429 | -1,102678765 | 0,143069691 | -7,707284148 | 1,29E-14 | 6,88E-12 |
| Fitm2 | protein_coding | 105,7644887 | -1,102045922 | 0,259481996 | -4,247099752 | 2,17E-05 | 0,001066456 |

| | | | | | | | |
|---|---|---|---|---|---|---|---|
| Actg1 | protein_coding | 13827,63328 | -1,100149319 | 0,293299703 | -3,750939084 | 1,76E-04 | 0,005287117 |
| Gm9574 | unprocessed_pseudo | 557,0686228 | -1,098933125 | 0,292927977 | -3,751547173 | 1,76E-04 | 0,005283862 |
| Gm7331 | processed_pseudoge | 245,365734 | -1,095314901 | 0,368612451 | -2,971453886 | 0,002963934 | 0,036873647 |
| Nrp1 | protein_coding | 3704,052472 | -1,095191136 | 0,120134546 | -9,116371406 | 7,77E-20 | 9,21E-17 |
| Ly6e | protein_coding | 70536,02285 | -1,094554548 | 0,285726177 | -3,830781479 | 1,28E-04 | 0,004116353 |
| Coro1a | protein_coding | 68426,59625 | -1,080140452 | 0,26486233 | -4,078120335 | 4,54E-05 | 0,001897933 |
| mt-Cytb | protein_coding | 72067,03761 | -1,078707197 | 0,270193109 | -3,992356433 | 6,54E-05 | 0,002470461 |
| Cst7 | protein_coding | 1302,520224 | -1,077223983 | 0,259993587 | -4,14327137 | 3,42E-05 | 0,001548596 |
| Cyth4 | protein_coding | 8183,43898 | -1,073017721 | 0,184869814 | -5,804180224 | 6,47E-09 | 1,08E-06 |
| Rps27rt | protein_coding | 9029,991876 | -1,071087992 | 0,3311229 | -3,234714335 | 0,001217645 | 0,020060149 |
| Rcn1 | protein_coding | 873,0383784 | -1,066466449 | 0,175357135 | -6,081682684 | 1,19E-09 | 2,50E-07 |
| Actb | protein_coding | 353600,6855 | -1,063535848 | 0,308743584 | -3,444722099 | 5,72E-04 | 0,01205916 |
| Arf2 | protein_coding | 275,0227655 | -1,060383418 | 0,28671705 | -3,698361913 | 2,17E-04 | 0,006113917 |
| Slc35d1 | protein_coding | 611,9587901 | -1,059753481 | 0,19315733 | -5,4864782 | 4,10E-08 | 5,44E-06 |
| Cerk | protein_coding | 481,0784372 | -1,05803236 | 0,317714523 | -3,330135334 | 8,68E-04 | 0,01609604 |
| Ctsw | protein_coding | 2136,717505 | -1,05604225 | 0,19301228 | -5,471373366 | 4,47E-08 | 5,88E-06 |
| Cd96 | protein_coding | 5249,007804 | -1,051986158 | 0,135216612 | -7,780006772 | 7,25E-15 | 4,15E-12 |
| Pfn1 | protein_coding | 54704,82199 | -1,048397585 | 0,306640609 | -3,418978286 | 6,29E-04 | 0,012846927 |
| Gm17541 | processed_pseudoge | 888,5701792 | -1,044174901 | 0,340271032 | -3,068656461 | 0,002150237 | 0,029688297 |
| Nmb | protein_coding | 225,8596006 | -1,043790209 | 0,26602411 | -3,923667704 | 8,72E-05 | 0,003059945 |
| Ighm | IG_C_gene | 23033,37781 | -1,043495025 | 0,226468001 | -4,607693017 | 4,07E-06 | 2,85E-04 |
| Rps27 | protein_coding | 25429,20265 | -1,043056766 | 0,320777576 | -3,251651126 | 0,001147368 | 0,01937102 |
| Arhgap18 | protein_coding | 516,1223165 | -1,032229388 | 0,16935994 | -6,094885183 | 1,10E-09 | 2,39E-07 |
| H2-Q10 | protein_coding | 1518,408715 | -1,030896257 | 0,290789576 | -3,545162354 | 3,92E-04 | 0,009369489 |
| Gm13341 | unprocessed_pseudo | 5252,499945 | -1,030298199 | 0,286337651 | -3,598193234 | 3,20E-04 | 0,008069723 |
| Ggt7 | protein_coding | 226,3870199 | -1,026569506 | 0,328653785 | -3,123559058 | 0,00178678 | 0,026080388 |
| mt-Co1 | protein_coding | 72596,88551 | -1,026370824 | 0,280631767 | -3,657357949 | 2,55E-04 | 0,006811776 |
| Gm5312 | processed_pseudoge | 629,8185512 | -1,026259512 | 0,307602892 | -3,336312952 | 8,49E-04 | 0,015920447 |
| Gm3739 | protein_coding | 137,134298 | -1,025476785 | 0,283100841 | -3,622302147 | 2,92E-04 | 0,007513042 |
| Gm28661 | unprocessed_pseudo | 26852,74075 | -1,023812249 | 0,290952346 | -3,518831389 | 4,33E-04 | 0,009977214 |
| Tgtp1 | protein_coding | 5500,594596 | -1,022705594 | 0,202532495 | -5,049587691 | 4,43E-07 | 4,43E-05 |
| Syde1 | protein_coding | 95,08930048 | -1,02088152 | 0,320960797 | -3,180704719 | 0,001469173 | 0,0227664 |
| Gm15536 | processed_pseudoge | 685,6640471 | -1,017037711 | 0,350614995 | -2,900725088 | 0,003723003 | 0,042658519 |
| Gm28437 | unprocessed_pseudo | 57118,1488 | -1,016299032 | 0,296105901 | -3,43221472 | 5,99E-04 | 0,012457956 |
| H2-T23 | protein_coding | 14236,63877 | -1,01596825 | 0,291430263 | -3,486145333 | 4,90E-04 | 0,010800287 |
| Cnn3 | protein_coding | 405,9624853 | -1,015259505 | 0,165763906 | -6,124732016 | 9,08E-10 | 2,07E-07 |
| Gm7776 | processed_pseudoge | 182,3306812 | -1,014848492 | 0,291416713 | -3,482464964 | 4,97E-04 | 0,010848985 |
| Gm42031 | lincRNA | 200,3500102 | -1,013170076 | 0,354915317 | -2,854681181 | 0,004308005 | 0,047205577 |
| Arhgdib | protein_coding | 33474,80846 | -1,011964386 | 0,269909988 | -3,74926617 | 1,77E-04 | 0,005310445 |
| Gm6394 | processed_pseudoge | 745,0785467 | -1,007751312 | 0,301577689 | -3,341597705 | 8,33E-04 | 0,015691357 |
| Cd81 | protein_coding | 2347,468925 | -1,007293733 | 0,165469257 | -6,087497751 | 1,15E-09 | 2,46E-07 |
| Gm37486 | processed_pseudoge | 256,3304737 | -1,006415528 | 0,338088987 | -2,976777022 | 0,002912958 | 0,036435303 |
| Tgtp2 | protein_coding | 5406,95322 | -1,004761504 | 0,214438029 | -4,68555652 | 2,79E-06 | 2,08E-04 |
| Ctla4 | protein_coding | 6330,290335 | -1,003673784 | 0,149658244 | -6,706438319 | 1,99E-11 | 6,30E-09 |
| AC158975.2 | lincRNA | 97,18226791 | -1,002548856 | 0,306645306 | -3,269408782 | 0,001077725 | 0,018534633 |
| Socs1 | protein_coding | 3689,939195 | -1,002539945 | 0,208517778 | -4,807935109 | 1,52E-06 | 1,23E-04 |
| Arhgap20 | protein_coding | 186,5166668 | -1,001630973 | 0,2256043 | -4,439768978 | 9,01E-06 | 5,45E-04 |
| Gm13493 | processed_pseudoge | 1848,231262 | -0,998139352 | 0,328288954 | -3,040429293 | 0,002362412 | 0,031680938 |
| Gm12094 | processed_pseudoge | 302,8158937 | -0,99393569 | 0,340625455 | -2,917972443 | 0,003523155 | 0,041263428 |
| Gm29216 | unprocessed_pseudo | 47233,61708 | -0,99197902 | 0,286613063 | -3,461039107 | 5,38E-04 | 0,011552676 |
| Gm13340 | unprocessed_pseudo | 18829,5249 | -0,987845566 | 0,278323131 | -3,54927584 | 3,86E-04 | 0,009291167 |
| Gtf2f2 | protein_coding | 611,4074101 | -0,986593689 | 0,270446904 | -3,648012502 | 2,64E-04 | 0,006960132 |
| Ucp2 | protein_coding | 34867,28323 | -0,986175404 | 0,304889257 | -3,234536416 | 0,001218404 | 0,020060149 |
| Gm12183 | processed_pseudoge | 9684,013596 | -0,981411056 | 0,215168729 | -4,561123071 | 5,09E-06 | 3,36E-04 |
| Trbv12-1 | TR_V_gene | 550,3151193 | -0,980412792 | 0,138696648 | -7,068756218 | 1,56E-12 | 6,18E-10 |
| Gm6375 | processed_pseudoge | 318,5806521 | -0,979144888 | 0,334399975 | -2,928065079 | 0,003410786 | 0,040403577 |
| Gm4617 | processed_pseudoge | 853,0476993 | -0,978506926 | 0,296479817 | -3,300416651 | 9,65E-04 | 0,017190996 |
| Fah | protein_coding | 146,880321 | -0,976883738 | 0,314524793 | -3,105903759 | 0,001896984 | 0,027069952 |
| Gm6485 | transcribed_processe | 639,4253761 | -0,976757651 | 0,287176853 | -3,401240876 | 6,71E-04 | 0,013477859 |
| Sdc4 | protein_coding | 1525,889976 | -0,972964169 | 0,176870274 | -5,501004472 | 3,78E-08 | 5,05E-06 |
| Cd79b | protein_coding | 306,3129462 | -0,972700117 | 0,293685197 | -3,312050205 | 9,26E-04 | 0,01677776 |
| E2f2 | protein_coding | 909,5729138 | -0,971685787 | 0,160566351 | -6,051615304 | 1,43E-09 | 2,90E-07 |
| Gm8815 | unprocessed_pseudo | 292,0868865 | -0,967316505 | 0,309223401 | -3,128212494 | 0,00175873 | 0,025807144 |
| Slc14a1 | protein_coding | 1502,22016 | -0,96686835 | 0,165431682 | -5,844517438 | 5,08E-09 | 8,78E-07 |
| Rps11-ps1 | processed_pseudoge | 1638,15755 | -0,966279685 | 0,309626666 | -3,120789618 | 0,001803668 | 0,026234602 |
| Gm1848 | processed_pseudoge | 1103,938526 | -0,964559277 | 0,263735453 | -3,657298498 | 2,55E-04 | 0,006811776 |
| Malat1 | lincRNA | 18945,63695 | -0,96302344 | 0,29451415 | -3,269871551 | 0,001075963 | 0,018523534 |
| Usp31 | protein_coding | 84,87360117 | -0,956053734 | 0,324594099 | -2,945382363 | 0,003225558 | 0,038960231 |
| Spry1 | protein_coding | 168,3374723 | -0,953672278 | 0,300537197 | -3,173225432 | 0,001507554 | 0,023295502 |

| | | | | | | | |
|---|---|---|---|---|---|---|---|
| Ptma | protein_coding | 11811,0363 | -0,952859383 | 0,269877091 | -3,530716078 | 4,14E-04 | 0,009646544 |
| Cdc25b | protein_coding | 1338,625811 | -0,951491992 | 0,237926248 | -3,999104764 | 6,36E-05 | 0,002431373 |
| Swap70 | protein_coding | 1110,111385 | -0,948221711 | 0,174519774 | -5,433319602 | 5,53E-08 | 7,12E-06 |
| Gm26623 | lincRNA | 450,9332208 | -0,948187655 | 0,278183762 | -3,408493892 | 6,53E-04 | 0,013220646 |
| Fau | protein_coding | 18839,97596 | -0,947271561 | 0,323423339 | -2,928890546 | 0,003401742 | 0,040403577 |
| Gm10132 | processed_pseudoge | 515,7560202 | -0,942569433 | 0,319928222 | -2,946190327 | 0,003217144 | 0,038915246 |
| Tnfrsf18 | protein_coding | 7327,41981 | -0,938446893 | 0,21085754 | -4,45062051 | 8,56E-06 | 5,22E-04 |
| Arrdc3 | protein_coding | 639,2496772 | -0,938088153 | 0,256671957 | -3,65481357 | 2,57E-04 | 0,006845024 |
| Prickle1 | protein_coding | 177,8560452 | -0,935889049 | 0,31369718 | -2,983415568 | 0,002850507 | 0,036150946 |
| Samhd1 | protein_coding | 11224,76782 | -0,935728927 | 0,187206116 | -4,99838866 | 5,78E-07 | 5,48E-05 |
| Gm15427 | processed_pseudoge | 15664,38887 | -0,93353609 | 0,312838406 | -2,984084027 | 0,002844287 | 0,036150946 |
| Ddx5 | protein_coding | 68570,94489 | -0,933302775 | 0,208751533 | -4,470878658 | 7,79E-06 | 4,86E-04 |
| Gm13408 | processed_pseudoge | 1648,550027 | -0,933256234 | 0,313295916 | -2,978833069 | 0,002893484 | 0,03641388 |
| Gm10288 | processed_pseudoge | 534,612298 | -0,933169114 | 0,329546282 | -2,831678475 | 0,004630438 | 0,049419136 |
| Arhgap5 | protein_coding | 312,5010764 | -0,931651027 | 0,298321476 | -3,12297673 | 0,001790319 | 0,026088357 |
| Gm8326 | processed_pseudoge | 135,5697255 | -0,930513868 | 0,307008707 | -3,030903839 | 0,002438229 | 0,032430568 |
| Limd2 | protein_coding | 31041,47966 | -0,928431536 | 0,262027234 | -3,543263511 | 3,95E-04 | 0,0094101 |
| Cd99l2 | protein_coding | 328,2604071 | -0,925476386 | 0,263544014 | -3,511657777 | 4,45E-04 | 0,010124038 |
| Klf13 | protein_coding | 11298,94799 | -0,919546632 | 0,25483218 | -3,608440006 | 3,08E-04 | 0,007816963 |
| Gm9800 | processed_pseudoge | 1816,311875 | -0,918449503 | 0,278660385 | -3,295945718 | 9,81E-04 | 0,017355199 |
| Arhgap31 | protein_coding | 879,6799308 | -0,913926934 | 0,219556021 | -4,162613862 | 3,15E-05 | 0,001466719 |
| Rpl18a | protein_coding | 20249,30615 | -0,910477972 | 0,313500071 | -2,904235295 | 0,003681514 | 0,042486058 |
| Irf8 | protein_coding | 697,4093419 | -0,91006088 | 0,187453439 | -4,85486361 | 1,20E-06 | 1,03E-04 |
| Crip1 | protein_coding | 6077,057636 | -0,908073603 | 0,299706355 | -3,029877702 | 0,002446528 | 0,032508067 |
| Cd2 | protein_coding | 16451,95385 | -0,905488548 | 0,23391158 | -3,871071918 | 1,08E-04 | 0,003611057 |
| Gm4977 | processed_pseudoge | 2537,490648 | -0,905138284 | 0,319568386 | -2,832377436 | 0,004620328 | 0,04935133 |
| Cd52 | protein_coding | 17351,61199 | -0,899356076 | 0,268008623 | -3,355698288 | 7,92E-04 | 0,015171136 |
| Tapbp | protein_coding | 12746,70313 | -0,896182447 | 0,172581849 | -5,19279666 | 2,07E-07 | 2,29E-05 |
| Gm6170 | processed_pseudoge | 10843,97576 | -0,895616811 | 0,305078751 | -2,935690563 | 0,003328061 | 0,039850292 |
| Ldhb | protein_coding | 1171,83398 | -0,894370194 | 0,170717623 | -5,23888617 | 1,62E-07 | 1,85E-05 |
| Fth1 | protein_coding | 18044,98968 | -0,894000097 | 0,241753704 | -3,697978899 | 2,17E-04 | 0,006113917 |
| Zscan29 | protein_coding | 970,640484 | -0,890408387 | 0,269934641 | -3,298607332 | 9,72E-04 | 0,017236453 |
| Cnn2 | protein_coding | 23338,24296 | -0,889539009 | 0,220942339 | -4,026113846 | 5,67E-05 | 0,002246054 |
| Slc25a24 | protein_coding | 429,4648259 | -0,888213507 | 0,23816184 | -3,729453498 | 1,92E-04 | 0,005583364 |
| Peli1 | protein_coding | 9667,279165 | -0,887678068 | 0,171216956 | -5,184521958 | 2,17E-07 | 2,36E-05 |
| Ndfip1 | protein_coding | 12860,77086 | -0,885692321 | 0,215731177 | -4,105536966 | 4,03E-05 | 0,001752471 |
| Cnbp | protein_coding | 20812,0188 | -0,88179674 | 0,226131442 | -3,899487532 | 9,64E-05 | 0,003271567 |
| Gm13624 | processed_pseudoge | 68,05556298 | -0,880816532 | 0,283592355 | -3,105924813 | 0,001896849 | 0,027069952 |
| Gm47469 | lincRNA | 121,4628771 | -0,877217908 | 0,21574568 | -4,06598134 | 4,78E-05 | 0,001974625 |
| Rps11 | protein_coding | 11253,55373 | -0,875790291 | 0,289711585 | -3,022972976 | 0,002503046 | 0,033021977 |
| Zfyve16 | protein_coding | 285,522204 | -0,874179525 | 0,267937379 | -3,262626248 | 0,00110385 | 0,018870646 |
| Fam49a | protein_coding | 692,5114571 | -0,873793083 | 0,126107729 | -6,928941566 | 4,24E-12 | 1,64E-09 |
| Igtp | protein_coding | 7789,033067 | -0,871713953 | 0,174502438 | -4,995425642 | 5,87E-07 | 5,54E-05 |
| Gm5869 | processed_pseudoge | 12297,27774 | -0,871286283 | 0,306905353 | -2,838941303 | 0,004526348 | 0,048905778 |
| Eef1a1 | protein_coding | 90266,2872 | -0,869078219 | 0,306986731 | -2,830996037 | 0,00464033 | 0,049435168 |
| Gm43305 | processed_transcript | 404,8591421 | -0,867698136 | 0,223395824 | -3,884128721 | 1,03E-04 | 0,003457135 |
| Gm13456 | processed_pseudoge | 5991,991758 | -0,867414134 | 0,299639422 | -2,894859853 | 0,003793278 | 0,043201107 |
| Rpl36-ps3 | processed_pseudoge | 922,6912909 | -0,867081056 | 0,29856215 | -2,904189488 | 0,003682052 | 0,042486058 |
| Atp1b3 | protein_coding | 19759,15499 | -0,866124461 | 0,254564142 | -3,402382023 | 6,68E-04 | 0,013454278 |
| Pear1 | protein_coding | 1938,832856 | -0,864042513 | 0,156280876 | -5,528779569 | 3,22E-08 | 4,39E-06 |
| Psme1 | protein_coding | 9270,226551 | -0,86330321 | 0,226748193 | -3,807321242 | 1,40E-04 | 0,004466302 |
| AC133509.1 | processed_pseudoge | 5295,868838 | -0,862430099 | 0,301179355 | -2,863510014 | 0,004189756 | 0,046355457 |
| Castor2 | protein_coding | 375,5076653 | -0,857343767 | 0,293110211 | -2,924987714 | 0,003444698 | 0,040631281 |
| Arhgap45 | protein_coding | 26356,08466 | -0,856801434 | 0,184674666 | -4,639517991 | 3,49E-06 | 2,49E-04 |
| Sell | protein_coding | 21628,92501 | -0,856103292 | 0,241717237 | -3,541755245 | 3,97E-04 | 0,0094101 |
| Fcrl1 | protein_coding | 390,6915467 | -0,856093247 | 0,255625551 | -3,349012818 | 8,11E-04 | 0,015370249 |
| Cox7a2l | protein_coding | 4678,367093 | -0,853635006 | 0,241407555 | -3,536074109 | 4,06E-04 | 0,009521571 |
| Rnf216 | protein_coding | 758,50826 | -0,852394892 | 0,168068148 | -5,071721806 | 3,94E-07 | 3,97E-05 |
| Gm7803 | processed_pseudoge | 1775,569976 | -0,851929458 | 0,216197271 | -3,94051902 | 8,13E-05 | 0,002908074 |
| Gnb1 | protein_coding | 5726,740088 | -0,850938303 | 0,255320182 | -3,332828199 | 8,60E-04 | 0,016013073 |
| Eno1b | protein_coding | 1127,021807 | -0,850613403 | 0,251062724 | -3,388051359 | 7,04E-04 | 0,013956406 |
| Erbb3 | protein_coding | 226,7660586 | -0,850396518 | 0,271549037 | -3,131649912 | 0,00173827 | 0,025678852 |
| Ftl1-ps1 | protein_coding | 2302,753592 | -0,849810258 | 0,291023649 | -2,920072863 | 0,003499496 | 0,04104426 |
| Cfl1 | protein_coding | 47954,64902 | -0,84881792 | 0,282455301 | -3,005140694 | 0,00265458 | 0,034578313 |
| Hmgn2 | protein_coding | 2081,987909 | -0,848072314 | 0,250335881 | -3,387737751 | 7,05E-04 | 0,013956406 |
| mt-Nd1 | protein_coding | 40255,89311 | -0,84572353 | 0,27261591 | -3,102253021 | 0,001920537 | 0,027314881 |
| Gm4202 | processed_pseudoge | 2876,168167 | -0,844968782 | 0,271668234 | -3,110296596 | 0,001868996 | 0,026816919 |
| Anxa5 | protein_coding | 1935,66986 | -0,844644037 | 0,13878124 | -6,086154259 | 1,16E-09 | 2,46E-07 |
| Sgpp1 | protein_coding | 451,8470327 | -0,844294508 | 0,179750229 | -4,697042743 | 2,64E-06 | 1,97E-04 |

| | | | | | | | |
|---|---|---|---|---|---|---|---|
| Hspa4l | protein_coding | 357,8051635 | -0,843870895 | 0,242121401 | -3,485321378 | 4,92E-04 | 0,010807599 |
| mt-Nd4 | protein_coding | 32456,89169 | -0,843569003 | 0,246422724 | -3,423259795 | 6,19E-04 | 0,012711322 |
| Ftl1 | protein_coding | 4627,579234 | -0,842314573 | 0,263528713 | -3,196291453 | 0,001392064 | 0,021898291 |
| Gm6430 | processed_pseudoge | 1213,427435 | -0,842120907 | 0,233838335 | -3,601295342 | 3,17E-04 | 0,007998303 |
| Tubb2b | protein_coding | 187,3777719 | -0,841198935 | 0,270357382 | -3,111433202 | 0,001861816 | 0,026763653 |
| Rac2 | protein_coding | 38232,67536 | -0,840807445 | 0,240930561 | -3,489833095 | 4,83E-04 | 0,010709235 |
| Slfn4 | protein_coding | 177,30655 | -0,840594978 | 0,267034333 | -3,147891015 | 0,00164453 | 0,024788936 |
| Coro2a | protein_coding | 2609,587232 | -0,840582324 | 0,155518985 | -5,405014192 | 6,48E-08 | 8,15E-06 |
| Ms4a6b | protein_coding | 24699,14667 | -0,839998962 | 0,243017148 | -3,456541931 | 5,47E-04 | 0,011686704 |
| Tiam1 | protein_coding | 438,3896885 | -0,839844399 | 0,183069302 | -4,587576344 | 4,48E-06 | 3,05E-04 |
| Gm6969 | processed_pseudoge | 519,0685725 | -0,839411125 | 0,292997036 | -2,864913366 | 0,004171233 | 0,046246379 |
| Ephb6 | protein_coding | 1133,714337 | -0,838650666 | 0,1523922 | -5,503238793 | 3,73E-08 | 5,03E-06 |
| Gm12435 | processed_pseudoge | 1422,892082 | -0,835305243 | 0,217715854 | -3,836676236 | 1,25E-04 | 0,004034501 |
| Gm11793 | processed_pseudoge | 1314,478927 | -0,834892731 | 0,207641381 | -4,020839804 | 5,80E-05 | 0,002286031 |
| Gm6180 | processed_pseudoge | 3750,405577 | -0,832517828 | 0,282196411 | -2,950136137 | 0,003176339 | 0,038666038 |
| Gpsm1 | protein_coding | 199,3231016 | -0,831799766 | 0,291038316 | -2,858042119 | 0,004262638 | 0,046911632 |
| Sh3bgrl3 | protein_coding | 8415,412807 | -0,830676074 | 0,250310372 | -3,318584312 | 9,05E-04 | 0,016595373 |
| Trbc1 | TR_C_gene | 10530,71118 | -0,830151341 | 0,243396489 | -3,410695634 | 6,48E-04 | 0,013146422 |
| Gm4735 | processed_pseudoge | 3235,629818 | -0,828989603 | 0,245707371 | -3,373889849 | 7,41E-04 | 0,014443084 |
| Slfn8 | protein_coding | 2716,115013 | -0,828366865 | 0,217569898 | -3,807359724 | 1,40E-04 | 0,004466302 |
| Irf1 | protein_coding | 7277,492603 | -0,827503081 | 0,161956514 | -5,109415234 | 3,23E-07 | 3,31E-05 |
| Usp11 | protein_coding | 286,3282084 | -0,825091438 | 0,232321796 | -3,551502488 | 3,83E-04 | 0,009238018 |
| Ms4a6c | protein_coding | 898,1245236 | -0,825029592 | 0,288467699 | -2,860041501 | 0,004235856 | 0,046709808 |
| Map7 | protein_coding | 232,7798593 | -0,82469184 | 0,192492886 | -4,284271776 | 1,83E-05 | 9,33E-04 |
| Smpdl3a | protein_coding | 4093,186198 | -0,824468485 | 0,153830628 | -5,359586021 | 8,34E-08 | 1,01E-05 |
| Ctss | protein_coding | 8750,66174 | -0,82439588 | 0,190408349 | -4,329620453 | 1,49E-05 | 7,95E-04 |
| H3f3b | protein_coding | 18917,08175 | -0,823763546 | 0,238227203 | -3,457890349 | 5,44E-04 | 0,011643354 |
| Sqor | protein_coding | 284,7300493 | -0,821196446 | 0,210268154 | -3,905472269 | 9,40E-05 | 0,003231291 |
| Gpx4-ps2 | transcribed_processe | 1614,770012 | -0,820468459 | 0,207510117 | -3,95387208 | 7,69E-05 | 0,002800656 |
| Slc41a3 | protein_coding | 318,0154746 | -0,818260309 | 0,199622266 | -4,099043282 | 4,15E-05 | 0,001797661 |
| Hnrnpf | protein_coding | 8045,413776 | -0,816998085 | 0,215580813 | -3,789753242 | 1,51E-04 | 0,004730864 |
| Gbp10 | protein_coding | 1287,614312 | -0,815638799 | 0,215287679 | -3,788599527 | 1,51E-04 | 0,004743916 |
| Gm10054 | processed_pseudoge | 442,8046636 | -0,815055114 | 0,285228609 | -2,857550357 | 0,004269249 | 0,046922155 |
| Mcub | protein_coding | 454,1218101 | -0,813255464 | 0,203501903 | -3,99630398 | 6,43E-05 | 0,002454648 |
| Eno1 | protein_coding | 11652,61485 | -0,812210582 | 0,236460037 | -3,434874627 | 5,93E-04 | 0,012406777 |
| mt-Nd4l | protein_coding | 5219,7214 | -0,806684831 | 0,244502087 | -3,2992963 | 9,69E-04 | 0,017236453 |
| Cnp | protein_coding | 7696,119846 | -0,802453168 | 0,19322915 | -4,152857723 | 3,28E-05 | 0,001509495 |
| Fmo5 | protein_coding | 317,2843852 | -0,802283631 | 0,262387435 | -3,057629763 | 0,00223095 | 0,030448068 |
| Rgs1 | protein_coding | 6525,711224 | -0,802059815 | 0,220228458 | -3,641944469 | 2,71E-04 | 0,007094238 |
| Gm3226 | processed_pseudoge | 1158,39307 | -0,799939945 | 0,245477818 | -3,258705619 | 0,001119217 | 0,019031282 |
| Osgepl1 | protein_coding | 155,7929133 | -0,79890919 | 0,268245809 | -2,978272779 | 0,002898779 | 0,036417965 |
| Gbp7 | protein_coding | 3188,99622 | -0,797758801 | 0,130577219 | -6,10947918 | 1,00E-09 | 2,24E-07 |
| H2-Ob | protein_coding | 1224,539968 | -0,797241193 | 0,131235403 | -6,074894224 | 1,24E-09 | 2,57E-07 |
| Psmb8 | protein_coding | 9030,411657 | -0,793049573 | 0,228634009 | -3,468642197 | 5,23E-04 | 0,011274409 |
| Frmd6 | protein_coding | 689,7809738 | -0,792152897 | 0,276676182 | -2,863104767 | 0,004195118 | 0,046383866 |
| Serp1 | protein_coding | 5516,285117 | -0,785946359 | 0,140523843 | -5,592975115 | 2,23E-08 | 3,22E-06 |
| Prmt2 | protein_coding | 371,0668756 | -0,78448816 | 0,17793364 | -4,408880536 | 1,04E-05 | 6,03E-04 |
| Lsp1 | protein_coding | 16796,14703 | -0,78427907 | 0,23683785 | -3,311460011 | 9,28E-04 | 0,016778683 |
| Tspan32 | protein_coding | 10379,5005 | -0,783839425 | 0,174906839 | -4,481468125 | 7,41E-06 | 4,71E-04 |
| Gm14970 | processed_pseudoge | 928,7858825 | -0,783765034 | 0,231814948 | -3,380994365 | 7,22E-04 | 0,01423551 |
| Il2ra | protein_coding | 7442,926156 | -0,783523293 | 0,102504411 | -7,643800768 | 2,11E-14 | 1,00E-11 |
| Slfn2 | protein_coding | 13287,64374 | -0,782606981 | 0,200610295 | -3,901130701 | 9,57E-05 | 0,003265359 |
| Stx5a | protein_coding | 2618,950759 | -0,782307874 | 0,195113975 | -4,009491763 | 6,08E-05 | 0,002353985 |
| Ephx1 | protein_coding | 2948,161331 | -0,78134314 | 0,110058983 | -7,099312749 | 1,25E-12 | 5,08E-10 |
| Tuba1a | protein_coding | 1560,644386 | -0,781177198 | 0,216279349 | -3,611889914 | 3,04E-04 | 0,007737341 |
| Fxyd5 | protein_coding | 14500,8746 | -0,780556175 | 0,236960288 | -3,294037923 | 9,88E-04 | 0,017399228 |
| Dnmbp | protein_coding | 194,8494852 | -0,77656816 | 0,268082898 | -2,896746362 | 0,003770545 | 0,04303711 |
| Gm12844 | processed_pseudoge | 557,1371188 | -0,775657902 | 0,248052684 | -3,126988555 | 0,001766068 | 0,025869078 |
| Cd7 | protein_coding | 612,7592412 | -0,774031991 | 0,176375452 | -4,38854718 | 1,14E-05 | 6,46E-04 |
| H2afz | protein_coding | 5279,075518 | -0,773469973 | 0,269988858 | -2,864821829 | 0,004172439 | 0,046246379 |
| Map3k1 | protein_coding | 2690,362657 | -0,77206844 | 0,199314762 | -3,873613938 | 1,07E-04 | 0,00358077 |
| Gm14085 | protein_coding | 1438,661474 | -0,771350026 | 0,175361704 | -4,398623012 | 1,09E-05 | 6,26E-04 |
| Gpx4 | protein_coding | 3472,963757 | -0,771100122 | 0,185965614 | -4,146466139 | 3,38E-05 | 0,001535849 |
| Litaf | protein_coding | 378,5337117 | -0,767289904 | 0,181724552 | -4,222268796 | 2,42E-05 | 0,001173635 |
| Gm3756 | processed_pseudoge | 949,4359384 | -0,763059515 | 0,242448805 | -3,147301619 | 0,001647849 | 0,024793925 |
| Gm12185 | protein_coding | 308,9937708 | -0,761852853 | 0,20321876 | -3,748929749 | 1,78E-04 | 0,005310445 |
| Gm7446 | processed_pseudoge | 676,1082794 | -0,761731695 | 0,240383454 | -3,168819157 | 0,001530596 | 0,023464865 |
| Tmbim6 | protein_coding | 14060,1402 | -0,761062533 | 0,186041323 | -4,090825192 | 4,30E-05 | 0,001833232 |
| Slc2a3 | protein_coding | 1753,403271 | -0,759355556 | 0,111253019 | -6,825482674 | 8,76E-12 | 3,03E-09 |

| | | | | | | | |
|---|---|---|---|---|---|---|---|
| Zc3h12d | protein_coding | 833,1618826 | -0,759330994 | 0,135459652 | -5,605587975 | 2,08E-08 | 3,02E-06 |
| Laptm5 | protein_coding | 47957,54724 | -0,759007147 | 0,245350287 | -3,093565352 | 0,00197767 | 0,027933115 |
| Dgka | protein_coding | 18022,50005 | -0,756350726 | 0,165089339 | -4,581463179 | 4,62E-06 | 3,12E-04 |
| Pqlc3 | protein_coding | 821,4752421 | -0,756312223 | 0,204249342 | -3,702886952 | 2,13E-04 | 0,006047182 |
| Trim59 | protein_coding | 3089,749962 | -0,755882705 | 0,164663443 | -4,590470674 | 4,42E-06 | 3,04E-04 |
| Cacna2d4 | protein_coding | 424,0770722 | -0,755805202 | 0,216570335 | -3,489883335 | 4,83E-04 | 0,010709235 |
| Selplg | protein_coding | 16251,5479 | -0,750816299 | 0,220204632 | -3,40962991 | 6,51E-04 | 0,013181781 |
| AC122253.3 | processed_pseudoge | 122,1806391 | -0,750528198 | 0,252903872 | -2,967642174 | 0,003000935 | 0,037194556 |
| Dok2 | protein_coding | 6143,798978 | -0,750004141 | 0,22700243 | -3,303947638 | 9,53E-04 | 0,017030754 |
| BC043934 | lincRNA | 320,6287065 | -0,749902639 | 0,236659624 | -3,168696996 | 0,001531239 | 0,023464865 |
| Trbv5 | TR_V_gene | 129,1350026 | -0,749537829 | 0,232810184 | -3,219523371 | 0,001284039 | 0,020709339 |
| Stat1 | protein_coding | 4652,2502 | -0,749118066 | 0,135488367 | -5,529021305 | 3,22E-08 | 4,39E-06 |
| Chchd10 | protein_coding | 1033,077164 | -0,74886366 | 0,133118048 | -5,6255607 | 1,85E-08 | 2,76E-06 |
| Zfp442 | protein_coding | 161,317077 | -0,74762851 | 0,257470691 | -2,903742196 | 0,003687316 | 0,042496322 |
| Cast | protein_coding | 5183,989303 | -0,745692398 | 0,147860559 | -5,043213721 | 4,58E-07 | 4,50E-05 |
| Mov10 | protein_coding | 1204,895056 | -0,745391169 | 0,167610835 | -4,447153836 | 8,70E-06 | 5,29E-04 |
| Itm2b | protein_coding | 26292,60194 | -0,744158265 | 0,236094784 | -3,15194708 | 0,001621857 | 0,024513964 |
| Itgb7 | protein_coding | 25629,11151 | -0,743335919 | 0,213701165 | -3,478389638 | 5,04E-04 | 0,01095762 |
| AW112010 | lincRNA | 8734,818791 | -0,74262729 | 0,212768924 | -3,490299592 | 4,82E-04 | 0,010709235 |
| Zbtb5 | protein_coding | 225,6336376 | -0,741599701 | 0,248061598 | -2,989578824 | 0,002793624 | 0,035884657 |
| Gm11407 | processed_pseudoge | 328,7746837 | -0,741198987 | 0,246661779 | -3,004920297 | 0,002656505 | 0,034578313 |
| Bin2 | protein_coding | 13452,0248 | -0,737353516 | 0,164895105 | -4,471651952 | 7,76E-06 | 4,86E-04 |
| Lat | protein_coding | 18201,16971 | -0,736483216 | 0,22593006 | -3,259784091 | 0,001114971 | 0,018978514 |
| Tmem123 | protein_coding | 8330,478862 | -0,735116242 | 0,171169122 | -4,294677875 | 1,75E-05 | 9,02E-04 |
| Izumo1r | protein_coding | 12244,55035 | -0,732288913 | 0,178915272 | -4,092936864 | 4,26E-05 | 0,001826605 |
| Pibf1 | protein_coding | 271,4631331 | -0,731536509 | 0,257530479 | -2,840582259 | 0,004503126 | 0,04884567 |
| Myl12a | protein_coding | 3742,606996 | -0,730035741 | 0,242249798 | -3,013565938 | 0,002581969 | 0,033820333 |
| Hspa8 | protein_coding | 18985,04303 | -0,727145728 | 0,246663853 | -2,947921707 | 0,003199181 | 0,038782764 |
| Gm6428 | processed_pseudoge | 332,7129016 | -0,72691513 | 0,243646811 | -2,98347894 | 0,002849916 | 0,036150946 |
| Cd83 | protein_coding | 4948,603699 | -0,723504134 | 0,1492221 | -4,84850524 | 1,24E-06 | 1,04E-04 |
| Natd1 | protein_coding | 452,8054425 | -0,722382853 | 0,249046253 | -2,90059715 | 0,003724524 | 0,042658519 |
| Oaz2-ps | processed_pseudoge | 212,0131424 | -0,722289084 | 0,234141403 | -3,084841361 | 0,002036608 | 0,028546907 |
| Gm12250 | processed_pseudoge | 1291,792729 | -0,721430337 | 0,156158045 | -4,619872995 | 3,84E-06 | 2,70E-04 |
| Cbfa2t2 | protein_coding | 485,8101237 | -0,719182517 | 0,170165822 | -4,226362896 | 2,37E-05 | 0,001155874 |
| AI467606 | protein_coding | 8085,217088 | -0,718641619 | 0,167629233 | -4,287090073 | 1,81E-05 | 9,27E-04 |
| Slc9a3r1 | protein_coding | 6508,799248 | -0,718342026 | 0,183715219 | -3,910084475 | 9,23E-05 | 0,003200077 |
| Rhoh | protein_coding | 10836,98957 | -0,71645755 | 0,165541142 | -4,32797273 | 1,50E-05 | 7,98E-04 |
| Baiap2 | protein_coding | 386,3697549 | -0,715148367 | 0,212049636 | -3,372551733 | 7,45E-04 | 0,014472935 |
| Srgn | protein_coding | 12200,25726 | -0,712703519 | 0,22081895 | -3,22754691 | 0,001248566 | 0,02042979 |
| Capn3 | protein_coding | 767,1728729 | -0,712185754 | 0,193715785 | -3,676446673 | 2,37E-04 | 0,00644506 |
| Itm2c | protein_coding | 7867,404708 | -0,710895063 | 0,151784496 | -4,683581529 | 2,82E-06 | 2,09E-04 |
| Atp5c1 | protein_coding | 5755,124909 | -0,710239424 | 0,223962322 | -3,171245138 | 0,00151787 | 0,023355003 |
| Gabarap | protein_coding | 4806,8338 | -0,708590412 | 0,206996064 | -3,42320717 | 6,19E-04 | 0,012711322 |
| Lad1 | protein_coding | 737,1524381 | -0,708484281 | 0,241702137 | -2,931228869 | 0,003376239 | 0,040224027 |
| Kdm1b | protein_coding | 743,9082669 | -0,705988732 | 0,225460049 | -3,131325193 | 0,001740193 | 0,025678852 |
| Ypel5 | protein_coding | 2024,623249 | -0,705367079 | 0,135861699 | -5,1918023 | 2,08E-07 | 2,29E-05 |
| Cmpk2 | protein_coding | 460,5674323 | -0,704960443 | 0,244516877 | -2,883074786 | 0,003938139 | 0,044370238 |
| Lck | protein_coding | 25805,24191 | -0,703583551 | 0,200247333 | -3,513572643 | 4,42E-04 | 0,010083562 |
| Rbm3-ps | processed_pseudoge | 399,8419329 | -0,703549093 | 0,210643223 | -3,340003459 | 8,38E-04 | 0,015734619 |
| Selenop | protein_coding | 2388,348064 | -0,702635845 | 0,095850244 | -7,330558786 | 2,29E-13 | 1,03E-10 |
| Pglyrp1 | protein_coding | 836,6843123 | -0,701999715 | 0,224463685 | -3,127453399 | 0,001763278 | 0,025851022 |
| Tmem173 | protein_coding | 7110,706203 | -0,699460301 | 0,139224835 | -5,023962135 | 5,06E-07 | 4,88E-05 |
| Psmb10 | protein_coding | 2935,198361 | -0,699385751 | 0,173021391 | -4,042192393 | 5,30E-05 | 0,002138249 |
| Cdk2ap2 | protein_coding | 10186,79645 | -0,697681743 | 0,22167264 | -3,147351625 | 0,001647567 | 0,024793925 |
| Capn2 | protein_coding | 2090,816602 | -0,696395649 | 0,116651495 | -5,969881916 | 2,37E-09 | 4,48E-07 |
| Gimap3 | protein_coding | 40279,68246 | -0,694927868 | 0,223999077 | -3,102369346 | 0,001919783 | 0,027314881 |
| Stat5a | protein_coding | 2538,926687 | -0,692485754 | 0,123210465 | -5,620348496 | 1,91E-08 | 2,82E-06 |
| Tpm3 | protein_coding | 13748,33945 | -0,692066552 | 0,213902762 | -3,235425967 | 0,001214614 | 0,020060149 |
| Sco1 | protein_coding | 250,9515714 | -0,691883284 | 0,172126874 | -4,019612215 | 5,83E-05 | 0,002292529 |
| Ssbp2 | protein_coding | 1218,559847 | -0,691546465 | 0,147028457 | -4,703487189 | 2,56E-06 | 1,93E-04 |
| Nfkbia | protein_coding | 11507,52426 | -0,690981495 | 0,184725391 | -3,740587533 | 1,84E-04 | 0,005402251 |
| Trav12n-2 | TR_V_gene | 128,5844173 | -0,690743699 | 0,240716811 | -2,86952829 | 0,004110846 | 0,045849189 |
| Rgs2 | protein_coding | 657,2642359 | -0,690634186 | 0,243218703 | -2,83956036 | 0,004517575 | 0,048905778 |
| Gch1 | protein_coding | 335,6328037 | -0,690562964 | 0,213781156 | -3,230233081 | 0,001236893 | 0,020304137 |
| Atp5b | protein_coding | 21109,94818 | -0,688714552 | 0,23109604 | -2,980209232 | 0,002880516 | 0,036326016 |
| Apobec1 | protein_coding | 308,2397063 | -0,688659336 | 0,214514262 | -3,210319578 | 0,001325875 | 0,021196582 |
| Il16 | protein_coding | 6765,518501 | -0,686267657 | 0,14901237 | -4,605440865 | 4,12E-06 | 2,87E-04 |
| Arhgdia | protein_coding | 13135,7589 | -0,686023249 | 0,19980178 | -3,433519208 | 5,96E-04 | 0,012421981 |
| Gm47586 | TEC | 210,3173123 | -0,685566647 | 0,201246325 | -3,406604543 | 6,58E-04 | 0,013280816 |

| | | | | | | | |
|---|---|---|---|---|---|---|---|
| Dusp4 | protein_coding | 278,0058372 | -0,685192492 | 0,212811211 | -3,219719901 | 0,001283159 | 0,020709339 |
| Cd200 | protein_coding | 1141,992834 | -0,684051547 | 0,239112802 | -2,86079014 | 0,004225867 | 0,046630642 |
| Dusp2 | protein_coding | 7069,072761 | -0,683861201 | 0,142709434 | -4,791983123 | 1,65E-06 | 1,31E-04 |
| Hmgn1 | protein_coding | 1205,679511 | -0,683826942 | 0,179641205 | -3,806626334 | 1,41E-04 | 0,004470301 |
| Ppp1r16a | protein_coding | 558,4245796 | -0,681584466 | 0,208517533 | -3,268715368 | 0,001080369 | 0,018556026 |
| Btg2 | protein_coding | 5309,175816 | -0,681236238 | 0,149658985 | -4,551923406 | 5,32E-06 | 3,49E-04 |
| Gbp11 | polymorphic_pseudo | 227,4921509 | -0,680408349 | 0,218273637 | -3,117226421 | 0,001825613 | 0,026414885 |
| Gm4076 | processed_pseudoge | 251,5982914 | -0,679235354 | 0,222321489 | -3,05519433 | 0,002249147 | 0,030545704 |
| Cd27 | protein_coding | 10052,23357 | -0,676924116 | 0,13639697 | -4,962897032 | 6,94E-07 | 6,30E-05 |
| Tspan3 | protein_coding | 1117,38186 | -0,672319618 | 0,111426046 | -6,033774368 | 1,60E-09 | 3,13E-07 |
| Tnfrsf4 | protein_coding | 8806,523429 | -0,672303658 | 0,178024869 | -3,77645922 | 1,59E-04 | 0,004943793 |
| Ywhaz | protein_coding | 19010,75182 | -0,671820574 | 0,232900017 | -2,884587913 | 0,003919263 | 0,044217601 |
| Il4ra | protein_coding | 5378,585751 | -0,671805524 | 0,161424038 | -4,161744012 | 3,16E-05 | 0,001468194 |
| Nop53 | protein_coding | 7173,851275 | -0,670835421 | 0,218608293 | -3,068664097 | 0,002150182 | 0,029688297 |
| Gm7592 | transcribed_unproce | 227,4474492 | -0,668899731 | 0,205819656 | -3,249931237 | 0,001154329 | 0,019429256 |
| Cdc42se2 | protein_coding | 9976,459737 | -0,666925238 | 0,14393956 | -4,633369983 | 3,60E-06 | 2,55E-04 |
| Msn | protein_coding | 26010,72134 | -0,665836414 | 0,194475907 | -3,423747578 | 6,18E-04 | 0,012711322 |
| Sep.09 | protein_coding | 11909,48719 | -0,664562214 | 0,162519603 | -4,089120336 | 4,33E-05 | 0,001833232 |
| Gvin1 | protein_coding | 1630,806639 | -0,663442362 | 0,164897978 | -4,023350491 | 5,74E-05 | 0,002267172 |
| Cdc42 | protein_coding | 16108,51498 | -0,659854876 | 0,220916203 | -2,98690122 | 0,002818208 | 0,03590924 |
| Prrc1 | protein_coding | 430,2655456 | -0,659008499 | 0,190828132 | -3,45341377 | 5,54E-04 | 0,011751791 |
| Smc4 | protein_coding | 15231,19843 | -0,658262127 | 0,164795484 | -3,994418488 | 6,49E-05 | 0,002462939 |
| Gm5620 | processed_pseudoge | 1336,52905 | -0,657881729 | 0,213487804 | -3,081589291 | 0,002058987 | 0,028739235 |
| Ift80 | protein_coding | 4179,56884 | -0,656829153 | 0,143809469 | -4,567356767 | 4,94E-06 | 3,29E-04 |
| Bmyc | protein_coding | 422,3290751 | -0,656617612 | 0,175386888 | -3,743823844 | 1,81E-04 | 0,005352084 |
| Itgb2 | protein_coding | 13148,52141 | -0,65621103 | 0,184033522 | -3,565714674 | 3,63E-04 | 0,008898836 |
| Spcs2 | protein_coding | 6591,090428 | -0,654784101 | 0,178969688 | -3,658631303 | 2,54E-04 | 0,00680935 |
| mt-Nd5 | protein_coding | 21050,26115 | -0,654664966 | 0,201972851 | -3,241351317 | 0,001189645 | 0,019802752 |
| Snx3 | protein_coding | 4446,500661 | -0,652376076 | 0,184677376 | -3,532517566 | 4,12E-04 | 0,009594513 |
| Prpsap2 | protein_coding | 340,255964 | -0,651334673 | 0,203624614 | -3,198703044 | 0,001380473 | 0,021819359 |
| Tap1 | protein_coding | 8893,833482 | -0,650622792 | 0,163401317 | -3,981747544 | 6,84E-05 | 0,002562843 |
| Gm10156 | processed_pseudoge | 293,7695165 | -0,650493157 | 0,224542581 | -2,896970156 | 0,003767856 | 0,04303602 |
| Clec2i | protein_coding | 1103,279192 | -0,650391644 | 0,215093465 | -3,023762917 | 0,00249652 | 0,032987457 |
| Vcp | protein_coding | 6327,261416 | -0,650214003 | 0,150791403 | -4,312009779 | 1,62E-05 | 8,44E-04 |
| Trbv12-2 | TR_V_gene | 953,254192 | -0,649992166 | 0,154221711 | -4,214660591 | 2,50E-05 | 0,00120335 |
| Camk2d | protein_coding | 2135,569694 | -0,649852214 | 0,097135378 | -6,690170255 | 2,23E-11 | 6,73E-09 |
| Gata1 | protein_coding | 668,2628918 | -0,649344346 | 0,187052607 | -3,471453063 | 5,18E-04 | 0,011186089 |
| Phf13 | protein_coding | 395,1873574 | -0,647667798 | 0,182298527 | -3,552786796 | 3,81E-04 | 0,009221504 |
| Gm10157 | processed_pseudoge | 718,5840259 | -0,6462858 | 0,219347229 | -2,946405123 | 0,00321491 | 0,038915246 |
| Arhgef1 | protein_coding | 30094,22796 | -0,646035181 | 0,187070576 | -3,453430229 | 5,54E-04 | 0,011751791 |
| Rbm3 | protein_coding | 5205,473182 | -0,645641556 | 0,203575571 | -3,171508025 | 0,001516497 | 0,023355003 |
| Eif1 | protein_coding | 7149,27359 | -0,644005591 | 0,193376901 | -3,330312924 | 8,67E-04 | 0,01609604 |
| L1cam | protein_coding | 618,215568 | -0,642287906 | 0,17980289 | -3,57217787 | 3,54E-04 | 0,008782356 |
| Tubb4b-ps2 | processed_pseudoge | 178,5708408 | -0,642221318 | 0,222716558 | -2,883581372 | 0,003931811 | 0,044329027 |
| Trav4d-4 | TR_V_gene | 130,3392909 | -0,642111998 | 0,218390192 | -2,940205284 | 0,003279949 | 0,039502197 |
| Cd1d1 | protein_coding | 1208,077183 | -0,642095006 | 0,152712364 | -4,204603921 | 2,62E-05 | 0,001254484 |
| Ccnd2 | protein_coding | 17255,72527 | -0,641484696 | 0,212136994 | -3,023917153 | 0,002495248 | 0,032987457 |
| Pld3 | protein_coding | 3492,338634 | -0,640830534 | 0,166731544 | -3,843487072 | 1,21E-04 | 0,003962733 |
| Zfp758 | protein_coding | 460,0411873 | -0,639207089 | 0,1544779 | -4,137854597 | 3,51E-05 | 0,001568203 |
| Cyth1 | protein_coding | 6752,472784 | -0,638745409 | 0,131597998 | -4,853762341 | 1,21E-06 | 1,03E-04 |
| Pitpnm1 | protein_coding | 4434,389148 | -0,638739932 | 0,118962328 | -5,369262224 | 7,91E-08 | 9,72E-06 |
| B4galnt1 | protein_coding | 8487,783413 | -0,636180431 | 0,172267322 | -3,692983804 | 2,22E-04 | 0,006202878 |
| Tpm3-rs7 | protein_coding | 3496,72478 | -0,636130477 | 0,216492192 | -2,938352975 | 0,003299611 | 0,039611538 |
| Rrp1 | protein_coding | 4746,87561 | -0,635926862 | 0,147375402 | -4,315013587 | 1,60E-05 | 8,36E-04 |
| Slc7a11 | protein_coding | 378,785786 | -0,635610197 | 0,207677186 | -3,06056822 | 0,002209174 | 0,030300377 |
| Nr1d2 | protein_coding | 694,6032925 | -0,635564922 | 0,197791597 | -3,213305986 | 0,001312164 | 0,021081003 |
| Gbp6 | protein_coding | 914,8932755 | -0,633260699 | 0,205320621 | -3,084252797 | 0,002040641 | 0,028560753 |
| S1pr4 | protein_coding | 7689,700037 | -0,629048436 | 0,167066306 | -3,765262139 | 1,66E-04 | 0,005094373 |
| Gm7964 | processed_pseudoge | 2046,275311 | -0,628004137 | 0,216252237 | -2,904035334 | 0,003683866 | 0,042486058 |
| Disp1 | protein_coding | 416,1717436 | -0,627888656 | 0,155598143 | -4,035322305 | 5,45E-05 | 0,00218909 |
| Pisd | protein_coding | 567,6746132 | -0,625116193 | 0,128934759 | -4,848313971 | 1,25E-06 | 1,04E-04 |
| Oaz2 | protein_coding | 1519,723464 | -0,624589804 | 0,168583212 | -3,704934782 | 2,11E-04 | 0,006008807 |
| Gbp9 | protein_coding | 3188,865133 | -0,622606641 | 0,129651247 | -4,802164683 | 1,57E-06 | 1,26E-04 |
| Zfp282 | protein_coding | 321,9102485 | -0,621983478 | 0,175612387 | -3,5417973 | 3,97E-04 | 0,0094101 |
| Gimap5 | protein_coding | 4076,599369 | -0,621946383 | 0,190202147 | -3,269923038 | 0,001075767 | 0,018523534 |
| Eif4a-ps4 | processed_pseudoge | 2906,153421 | -0,62098392 | 0,204397106 | -3,038124818 | 0,002380553 | 0,031835345 |
| Traf1 | protein_coding | 4444,880195 | -0,620528453 | 0,117285152 | -5,290767327 | 1,22E-07 | 1,44E-05 |
| Os9 | protein_coding | 3076,428213 | -0,620439256 | 0,142470543 | -4,354859899 | 1,33E-05 | 7,32E-04 |
| Dazap2 | protein_coding | 10253,12834 | -0,619135114 | 0,169547825 | -3,651684212 | 2,61E-04 | 0,006917906 |

| | | | | | | | |
|---|---|---|---|---|---|---|---|
| Cish | protein_coding | 4081,575106 | -0,61871635 | 0,154590501 | -4,002292163 | 6,27E-05 | 0,002415537 |
| Cndp2 | protein_coding | 1568,064927 | -0,617408777 | 0,155870995 | -3,961024165 | 7,46E-05 | 0,002746214 |
| Fam107b | protein_coding | 8174,677526 | -0,616816221 | 0,183036629 | -3,369905935 | 7,52E-04 | 0,014578471 |
| Emp3 | protein_coding | 5364,282282 | -0,616556277 | 0,198503761 | -3,106018112 | 0,001896251 | 0,027069952 |
| Tmem18 | protein_coding | 439,4971852 | -0,615523965 | 0,205232198 | -2,999158861 | 0,002707261 | 0,035128776 |
| Clic1 | protein_coding | 9511,893412 | -0,614664145 | 0,202046756 | -3,042187641 | 0,002348654 | 0,03156135 |
| Rnh1 | protein_coding | 2204,341536 | -0,614436771 | 0,149597207 | -4,10727435 | 4,00E-05 | 0,001743911 |
| Ets1 | protein_coding | 15525,46535 | -0,614091678 | 0,185003496 | -3,319351749 | 9,02E-04 | 0,016582528 |
| Vcp-rs | processed_pseudoge | 2364,33707 | -0,61405278 | 0,152475585 | -4,027220366 | 5,64E-05 | 0,002246054 |
| Sp100 | protein_coding | 10006,76661 | -0,613283879 | 0,144886711 | -4,232851132 | 2,31E-05 | 0,001132983 |
| Trbv15 | TR_V_gene | 699,755628 | -0,612800289 | 0,106137707 | -5,773634155 | 7,76E-09 | 1,29E-06 |
| Dpp4 | protein_coding | 1717,556745 | -0,61233557 | 0,175406444 | -3,490952527 | 4,81E-04 | 0,010707351 |
| Spopl | protein_coding | 1616,582204 | -0,611968996 | 0,179098879 | -3,416933711 | 6,33E-04 | 0,012927875 |
| Tax1bp1 | protein_coding | 4701,454171 | -0,611471004 | 0,132109393 | -4,628520262 | 3,68E-06 | 2,60E-04 |
| Ctsd | protein_coding | 3852,93351 | -0,610151205 | 0,165903884 | -3,677739127 | 2,35E-04 | 0,006423841 |
| Fam129a | protein_coding | 1824,733082 | -0,608825244 | 0,148702729 | -4,094243925 | 4,24E-05 | 0,001822414 |
| Ndrg3 | protein_coding | 3292,693734 | -0,607337185 | 0,116034305 | -5,234117469 | 1,66E-07 | 1,88E-05 |
| Rabac1 | protein_coding | 4066,960828 | -0,606737711 | 0,172755256 | -3,512123012 | 4,45E-04 | 0,010120195 |
| Bnip3l-ps | transcribed_processe | 567,4093353 | -0,606575793 | 0,175573205 | -3,454831233 | 5,51E-04 | 0,011730909 |
| Hnrnpk | protein_coding | 21297,71276 | -0,606448689 | 0,208720021 | -2,905560698 | 0,003665957 | 0,042397372 |
| Trbv13-3 | TR_V_gene | 1116,750442 | -0,603193058 | 0,170736417 | -3,532890442 | 4,11E-04 | 0,009594467 |
| P4ha1 | protein_coding | 1869,307816 | -0,603152349 | 0,137400899 | -4,389726365 | 1,13E-05 | 6,45E-04 |
| Ppp1r18 | protein_coding | 7427,796605 | -0,602676791 | 0,171445229 | -3,515273041 | 4,39E-04 | 0,010042228 |
| Neu3 | protein_coding | 651,7004977 | -0,602599898 | 0,168313 | -3,580233834 | 3,43E-04 | 0,008567201 |
| Slfn1 | protein_coding | 6566,843945 | -0,601316805 | 0,191193984 | -3,145061323 | 0,00166052 | 0,02496195 |
| Eif4a1 | protein_coding | 8849,265341 | -0,600887981 | 0,196520569 | -3,057634034 | 0,002230918 | 0,030448068 |
| Ccdc125 | protein_coding | 352,0638452 | -0,599302164 | 0,163515035 | -3,665119636 | 2,47E-04 | 0,006682268 |
| Tmem50a | protein_coding | 5733,790214 | -0,598462764 | 0,171839844 | -3,482677532 | 4,96E-04 | 0,010848985 |
| Sf3b1 | protein_coding | 15459,7349 | -0,595151813 | 0,146024167 | -4,07570764 | 4,59E-05 | 0,001912903 |
| Tmem63a | protein_coding | 1303,785953 | -0,594289683 | 0,150807257 | -3,940723368 | 8,12E-05 | 0,002908074 |
| Rsl1 | protein_coding | 343,6979861 | -0,594069966 | 0,191006517 | -3,110207841 | 0,001869557 | 0,026816919 |
| Eva1b | protein_coding | 705,9174075 | -0,593568377 | 0,177567367 | -3,342778498 | 8,29E-04 | 0,01565973 |
| Gm6335 | processed_pseudoge | 470,7538614 | -0,591571508 | 0,1903408 | -3,107959548 | 0,001883839 | 0,026975137 |
| Rnf187 | protein_coding | 5900,839364 | -0,591083256 | 0,178719666 | -3,30732072 | 9,42E-04 | 0,016897686 |
| Oat | protein_coding | 1417,903225 | -0,590679694 | 0,157090156 | -3,760131818 | 1,70E-04 | 0,005161898 |
| Tubb4b | protein_coding | 2619,476315 | -0,589934915 | 0,17457689 | -3,379226852 | 7,27E-04 | 0,014246269 |
| Mical1 | protein_coding | 2125,554049 | -0,589349273 | 0,122389846 | -4,815344527 | 1,47E-06 | 1,20E-04 |
| Grhpr | protein_coding | 396,7031214 | -0,588162476 | 0,160517061 | -3,664174227 | 2,48E-04 | 0,00669609 |
| Calm2 | protein_coding | 6123,783498 | -0,587670549 | 0,196733935 | -2,987133602 | 0,002816066 | 0,03590924 |
| Wls | protein_coding | 2051,3997 | -0,587539133 | 0,156923015 | -3,744123404 | 1,81E-04 | 0,005352084 |
| Slc22a5 | protein_coding | 654,5348982 | -0,586885176 | 0,193034106 | -3,04031856 | 0,00236328 | 0,031680938 |
| Fbxo32 | protein_coding | 588,6875177 | -0,586015212 | 0,176090284 | -3,327924743 | 8,75E-04 | 0,016206198 |
| Vwa5a | protein_coding | 614,1509464 | -0,585995339 | 0,175682237 | -3,335541199 | 8,51E-04 | 0,015946684 |
| Serpinb9 | protein_coding | 564,1472991 | -0,585159338 | 0,204758469 | -2,857802858 | 0,004265853 | 0,046915906 |
| Cd6 | protein_coding | 14964,14968 | -0,583986595 | 0,158643333 | -3,68112914 | 2,32E-04 | 0,006396815 |
| Btbd6 | protein_coding | 602,165601 | -0,583703224 | 0,184067554 | -3,171135866 | 0,001518441 | 0,023355003 |
| Xlr4a | protein_coding | 802,718276 | -0,583591518 | 0,190372666 | -3,065521599 | 0,002172907 | 0,029901792 |
| Atp2a3 | protein_coding | 4955,57799 | -0,583587268 | 0,096667682 | -6,03704624 | 1,57E-09 | 3,10E-07 |
| Pfdn5 | protein_coding | 2663,129792 | -0,582884952 | 0,205797472 | -2,832323187 | 0,004621112 | 0,04935133 |
| Mtmr3 | protein_coding | 1721,895169 | -0,582050748 | 0,166107696 | -3,504056477 | 4,58E-04 | 0,010346605 |
| Tbc1d10c | protein_coding | 2897,149241 | -0,581300532 | 0,19709205 | -2,949385995 | 0,00318406 | 0,038696855 |
| Arhgap9 | protein_coding | 8948,956616 | -0,580560097 | 0,147176057 | -3,944664029 | 7,99E-05 | 0,00287683 |
| Tubgcp4 | protein_coding | 787,1963943 | -0,578507945 | 0,171291743 | -3,377325341 | 7,32E-04 | 0,014307828 |
| Myo18a | protein_coding | 1059,878372 | -0,578189676 | 0,16731421 | -3,455711709 | 5,49E-04 | 0,011707689 |
| Cdc42ep3 | protein_coding | 824,4227372 | -0,578189516 | 0,159110567 | -3,633885094 | 2,79E-04 | 0,007273718 |
| Card6 | protein_coding | 1501,591666 | -0,578156508 | 0,143286334 | -4,034973133 | 5,46E-05 | 0,00218909 |
| Nt5e | protein_coding | 2204,386058 | -0,577995991 | 0,181330214 | -3,187532723 | 0,001434922 | 0,022402605 |
| Kctd10 | protein_coding | 1904,045677 | -0,577518456 | 0,111476762 | -5,180617405 | 2,21E-07 | 2,40E-05 |
| Rab3ip | protein_coding | 1178,429625 | -0,577019708 | 0,138196754 | -4,17534922 | 2,98E-05 | 0,001402783 |
| Ass1 | protein_coding | 4095,418411 | -0,576594318 | 0,139165516 | -4,143226949 | 3,42E-05 | 0,001548596 |
| Ncor1 | protein_coding | 1545,045843 | -0,576365555 | 0,120693277 | -4,775456987 | 1,79E-06 | 1,40E-04 |
| Fmnl3 | protein_coding | 759,551797 | -0,575726135 | 0,159246851 | -3,615306235 | 3,00E-04 | 0,007659508 |
| Gm49396 | protein_coding | 588,2212386 | -0,574945848 | 0,164219571 | -3,501079957 | 4,63E-04 | 0,010420327 |
| Flna | protein_coding | 6709,231231 | -0,573897737 | 0,095514996 | -6,008456932 | 1,87E-09 | 3,61E-07 |
| Sf3b2 | protein_coding | 6197,550081 | -0,57378394 | 0,141394597 | -4,058032988 | 4,95E-05 | 0,002022904 |
| Ndufa5 | protein_coding | 696,9845485 | -0,573269188 | 0,192573085 | -2,976891538 | 0,00291187 | 0,036435303 |
| Ifi47 | protein_coding | 8256,125578 | -0,573077815 | 0,167477524 | -3,421819241 | 6,22E-04 | 0,012760591 |
| Gm5424 | processed_pseudoge | 660,4246315 | -0,5727704 | 0,153412359 | -3,733534918 | 1,89E-04 | 0,005530164 |
| Ccng1 | protein_coding | 4005,693734 | -0,572724473 | 0,136463337 | -4,196910954 | 2,71E-05 | 0,001294107 |

| | | | | | | | |
|---|---|---|---|---|---|---|---|
| Mat2b | protein_coding | 4478,78456 | -0,572212543 | 0,160832664 | -3,557813009 | 3,74E-04 | 0,009102752 |
| Arl6ip1 | protein_coding | 10450,03864 | -0,569113692 | 0,190512347 | -2,987279825 | 0,00281472 | 0,03590924 |
| Kdm4c | protein_coding | 829,3940232 | -0,569111392 | 0,190825212 | -2,982370019 | 0,002860261 | 0,036150946 |
| Lta | protein_coding | 1965,811767 | -0,568322476 | 0,128304706 | -4,429474905 | 9,45E-06 | 5,62E-04 |
| Pja2 | protein_coding | 838,2026443 | -0,567461232 | 0,188815803 | -3,005369371 | 0,002652585 | 0,034578313 |
| Rassf5 | protein_coding | 10435,39255 | -0,566896544 | 0,162595899 | -3,486536553 | 4,89E-04 | 0,010800287 |
| Sipa1 | protein_coding | 5075,323611 | -0,565186058 | 0,132005376 | -4,281538167 | 1,86E-05 | 9,38E-04 |
| Pnrc1 | protein_coding | 6017,370007 | -0,564153141 | 0,141255556 | -3,993847451 | 6,50E-05 | 0,002463244 |
| Trbj1-2 | TR_J_gene | 241,60083 | -0,564003719 | 0,182099137 | -3,097234447 | 0,001953353 | 0,027649911 |
| Slc35b4 | protein_coding | 660,7557191 | -0,56369862 | 0,174428704 | -3,231684972 | 0,001230626 | 0,020221264 |
| Socs2 | protein_coding | 1398,890578 | -0,561625821 | 0,114813385 | -4,891640669 | 1,00E-06 | 8,73E-05 |
| Il17ra | protein_coding | 4277,440101 | -0,561403581 | 0,123912217 | -4,530655599 | 5,88E-06 | 3,80E-04 |
| Gm15542 | processed_pseudoge | 716,3935544 | -0,560908459 | 0,190359407 | -2,946575985 | 0,003213135 | 0,038915246 |
| 2310011J03Rik | protein_coding | 1652,911736 | -0,560175847 | 0,185346044 | -3,022324269 | 0,002508417 | 0,033065675 |
| Lcp1 | protein_coding | 24557,34379 | -0,559862352 | 0,191069635 | -2,930148232 | 0,003388003 | 0,040335226 |
| Gmppb | protein_coding | 443,0054971 | -0,55964827 | 0,151803657 | -3,686658672 | 2,27E-04 | 0,006312851 |
| Tacc1 | protein_coding | 1651,154368 | -0,558174115 | 0,197058624 | -2,832528229 | 0,004618149 | 0,04935133 |
| Snx20 | protein_coding | 2424,771037 | -0,556324689 | 0,162646729 | -3,420448073 | 6,25E-04 | 0,012793463 |
| Psme2b | protein_coding | 1068,181214 | -0,555529098 | 0,190639892 | -2,914023356 | 0,003568032 | 0,041671401 |
| Etfbkmt | protein_coding | 1095,293745 | -0,553925517 | 0,182885891 | -3,028803991 | 0,002455239 | 0,032571024 |
| Ube2l6 | protein_coding | 606,4362547 | -0,55391128 | 0,177604865 | -3,118784385 | 0,001815988 | 0,026344525 |
| Wdr6 | protein_coding | 2281,553603 | -0,553730556 | 0,148810945 | -3,721033804 | 1,98E-04 | 0,005694644 |
| Cd5 | protein_coding | 15973,44914 | -0,553523735 | 0,172449404 | -3,209774712 | 0,00132839 | 0,021196582 |
| Arid5a | protein_coding | 2468,131602 | -0,551567327 | 0,131594956 | -4,19140173 | 2,77E-05 | 0,001314575 |
| Trp53i11 | protein_coding | 3808,628683 | -0,551304067 | 0,092403002 | -5,966300423 | 2,43E-09 | 4,53E-07 |
| Sh2d2a | protein_coding | 7711,336963 | -0,550898003 | 0,182479545 | -3,018957563 | 0,002536461 | 0,033336628 |
| Zfp677 | protein_coding | 495,5784201 | -0,550880718 | 0,181565436 | -3,0340616 | 0,002412852 | 0,032163602 |
| Tuba4a | protein_coding | 3964,891952 | -0,549609494 | 0,173881166 | -3,160833966 | 0,001573181 | 0,02395277 |
| Il6ra | protein_coding | 4730,64895 | -0,549070183 | 0,098286844 | -5,586405684 | 2,32E-08 | 3,29E-06 |
| Sdf4 | protein_coding | 9307,71447 | -0,548515214 | 0,163685818 | -3,351024665 | 8,05E-04 | 0,015323468 |
| Wdr1 | protein_coding | 13401,95875 | -0,54810644 | 0,168635457 | -3,250244348 | 0,001153059 | 0,019429256 |
| Dis3l | protein_coding | 484,47385 | -0,547265986 | 0,149995457 | -3,648550402 | 2,64E-04 | 0,006960132 |
| Lcp2 | protein_coding | 8991,260757 | -0,546648426 | 0,157880162 | -3,462426294 | 5,35E-04 | 0,011523103 |
| Tubb4b-ps1 | processed_pseudoge | 772,790897 | -0,546458143 | 0,187831457 | -2,909300461 | 0,003622385 | 0,04208083 |
| Dnajc9 | protein_coding | 1895,031805 | -0,544245957 | 0,118268114 | -4,60179788 | 4,19E-06 | 2,91E-04 |
| Golph3l | protein_coding | 997,6183496 | -0,543394113 | 0,17041058 | -3,188734604 | 0,00142897 | 0,022353882 |
| Gm12216 | protein_coding | 370,7947628 | -0,543350956 | 0,188939786 | -2,875788998 | 0,00403019 | 0,045192592 |
| Rdx | protein_coding | 1769,87384 | -0,542778953 | 0,155128668 | -3,498895207 | 4,67E-04 | 0,010472066 |
| Rptor | protein_coding | 499,4817541 | -0,541877981 | 0,160751654 | -3,370901428 | 7,49E-04 | 0,014542883 |
| Zfp329 | protein_coding | 378,4208028 | -0,539895466 | 0,168200111 | -3,209840121 | 0,001328088 | 0,021196582 |
| Stx7 | protein_coding | 785,1903917 | -0,538896254 | 0,139699072 | -3,857550718 | 1,15E-04 | 0,003793856 |
| Cipc | protein_coding | 581,7666506 | -0,538234998 | 0,160281508 | -3,358060489 | 7,85E-04 | 0,015121975 |
| Tmem154 | protein_coding | 2852,094938 | -0,537669753 | 0,150673431 | -3,568444347 | 3,59E-04 | 0,008872388 |
| Syf2 | protein_coding | 3322,143979 | -0,537400206 | 0,165850955 | -3,240259948 | 0,001194208 | 0,019838911 |
| Iigp1 | protein_coding | 1359,645377 | -0,537358923 | 0,158321925 | -3,39409038 | 6,89E-04 | 0,013751506 |
| Stat5b | protein_coding | 3492,796862 | -0,53706229 | 0,105448207 | -5,093138195 | 3,52E-07 | 3,59E-05 |
| Sep.01 | protein_coding | 8423,382703 | -0,536499152 | 0,189114387 | -2,836902902 | 0,004555346 | 0,049059395 |
| Ezh1 | protein_coding | 1659,575315 | -0,535720288 | 0,16529991 | -3,240898859 | 0,001191535 | 0,019814335 |
| Lta4h | protein_coding | 4606,518362 | -0,532875222 | 0,150630953 | -3,53762099 | 4,04E-04 | 0,009490965 |
| Exoc2 | protein_coding | 1803,561601 | -0,532327768 | 0,121022866 | -4,39857181 | 1,09E-05 | 6,26E-04 |
| Lrp10 | protein_coding | 7469,552039 | -0,532035045 | 0,105533422 | -5,041389109 | 4,62E-07 | 4,50E-05 |
| Sqstm1 | protein_coding | 7682,799678 | -0,529182738 | 0,14383385 | -3,679125166 | 2,34E-04 | 0,00641992 |
| Traf3ip3 | protein_coding | 7200,821888 | -0,528222378 | 0,159704089 | -3,307506916 | 9,41E-04 | 0,016897686 |
| Ptprc | protein_coding | 31748,52865 | -0,527761104 | 0,162224647 | -3,253273241 | 0,001140837 | 0,019300038 |
| Relb | protein_coding | 1647,946336 | -0,526549381 | 0,103805359 | -5,0724682 | 3,93E-07 | 3,97E-05 |
| Mfap1a | protein_coding | 1754,236012 | -0,526432787 | 0,131787594 | -3,994554963 | 6,48E-05 | 0,002462939 |
| Sh3bp5 | protein_coding | 1616,766136 | -0,526428334 | 0,132505856 | -3,972868454 | 7,10E-05 | 0,002639318 |
| Iws1 | protein_coding | 900,5781768 | -0,525508554 | 0,133456842 | -3,93766664 | 8,23E-05 | 0,002936521 |
| Irgm1 | protein_coding | 3093,414868 | -0,525169847 | 0,160937279 | -3,263195756 | 0,001101634 | 0,018867613 |
| Serpinb6a | protein_coding | 678,8402334 | -0,522996272 | 0,111184431 | -4,70386245 | 2,55E-06 | 1,93E-04 |
| Ppp1ca | protein_coding | 10543,47058 | -0,522758393 | 0,18191092 | -2,873705396 | 0,004056872 | 0,045369169 |
| Tnfrsf13b | protein_coding | 1534,379501 | -0,522162312 | 0,110450175 | -4,727582498 | 2,27E-06 | 1,75E-04 |
| Nod1 | protein_coding | 1291,269318 | -0,522067749 | 0,156312888 | -3,339889334 | 8,38E-04 | 0,015734619 |
| Ifi35 | protein_coding | 1887,390311 | -0,520895865 | 0,15309603 | -3,402412624 | 6,68E-04 | 0,013454278 |
| Sorl1 | protein_coding | 544,9674859 | -0,52074019 | 0,172886682 | -3,012031835 | 0,002595054 | 0,033938151 |
| Cd82 | protein_coding | 8373,056254 | -0,520703754 | 0,153365961 | -3,39517158 | 6,86E-04 | 0,013738705 |
| Zfp110 | protein_coding | 1408,876847 | -0,520625898 | 0,126478277 | -4,116326622 | 3,85E-05 | 0,001694635 |
| Plekhb2 | protein_coding | 2197,33529 | -0,51982101 | 0,136932808 | -3,796175776 | 1,47E-04 | 0,004636321 |
| Cd9 | protein_coding | 1338,447738 | -0,519028563 | 0,104462849 | -4,968546907 | 6,75E-07 | 6,15E-05 |

| | | | | | | | |
|---|---|---|---|---|---|---|---|
| Ube2d3 | protein_coding | 9472,77584 | -0,518620456 | 0,176506465 | -2,938251893 | 0,003300687 | 0,039611538 |
| Eif4g2 | protein_coding | 14717,62777 | -0,518498628 | 0,162363068 | -3,193451782 | 0,001405828 | 0,022081672 |
| Triobp | protein_coding | 1174,378186 | -0,518421316 | 0,127103807 | -4,078723734 | 4,53E-05 | 0,001897795 |
| Bnip3l | protein_coding | 3296,533184 | -0,517935095 | 0,148276063 | -3,493045918 | 4,78E-04 | 0,010643164 |
| Def6 | protein_coding | 3477,464489 | -0,51720878 | 0,139008602 | -3,720696214 | 1,99E-04 | 0,005694644 |
| Ikzf4 | protein_coding | 3221,90821 | -0,516948886 | 0,100194122 | -5,15947319 | 2,48E-07 | 2,62E-05 |
| Gm10093 | processed_pseudoge | 952,1124891 | -0,51535345 | 0,118506181 | -4,348747438 | 1,37E-05 | 7,47E-04 |
| Irgm2 | protein_coding | 1734,181262 | -0,515116853 | 0,126158337 | -4,083097982 | 4,44E-05 | 0,001876626 |
| Xbp1 | protein_coding | 2632,154471 | -0,512189821 | 0,129525981 | -3,954340407 | 7,67E-05 | 0,002800656 |
| Tspo | protein_coding | 3889,586848 | -0,509015977 | 0,178349379 | -2,854038401 | 0,004316731 | 0,047253663 |
| Gnai3 | protein_coding | 4676,475661 | -0,507058364 | 0,136298882 | -3,720194606 | 1,99E-04 | 0,005696129 |
| Slamf1 | protein_coding | 1961,123647 | -0,506951579 | 0,117754254 | -4,305165736 | 1,67E-05 | 8,65E-04 |
| Akna | protein_coding | 9227,999459 | -0,505673778 | 0,11347211 | -4,456370642 | 8,34E-06 | 5,15E-04 |
| Pbxip1 | protein_coding | 3125,456042 | -0,505223399 | 0,151236983 | -3,3406075 | 8,36E-04 | 0,015729567 |
| Ptges3 | protein_coding | 2445,064993 | -0,505190996 | 0,143787674 | -3,513451328 | 4,42E-04 | 0,010083562 |
| Slc26a2 | protein_coding | 801,5892245 | -0,504833566 | 0,171821919 | -2,938120862 | 0,003302083 | 0,039611538 |
| Ablim1 | protein_coding | 12420,8018 | -0,503821271 | 0,15540546 | -3,241979221 | 0,001187027 | 0,019779012 |
| Btg1 | protein_coding | 10842,91426 | -0,502530972 | 0,164296487 | -3,058683603 | 0,002223118 | 0,030416214 |
| Tnfaip8l2 | protein_coding | 2842,63894 | -0,50098999 | 0,15393229 | -3,254612727 | 0,001135471 | 0,019228852 |
| Akr1b3 | protein_coding | 2167,091702 | -0,499996561 | 0,148712036 | -3,362179509 | 7,73E-04 | 0,014957645 |
| Commd4 | protein_coding | 1815,133405 | -0,499983448 | 0,173043675 | -2,889348297 | 0,003860412 | 0,043732015 |
| Tpm4 | protein_coding | 5100,06805 | -0,499786374 | 0,132002374 | -3,786192308 | 1,53E-04 | 0,004763124 |
| Zfand6 | protein_coding | 3346,714542 | -0,497970318 | 0,158650023 | -3,138797645 | 0,001696426 | 0,025272781 |
| Fam118a | protein_coding | 877,3598654 | -0,497650099 | 0,172005235 | -2,893226469 | 0,003813063 | 0,043284258 |
| Sat1 | protein_coding | 4222,813647 | -0,496764702 | 0,144742655 | -3,432054662 | 5,99E-04 | 0,012457956 |
| Cirbp | protein_coding | 2548,016634 | -0,495484428 | 0,167445214 | -2,959083846 | 0,003085551 | 0,037819653 |
| Hip1r | protein_coding | 1033,018659 | -0,495151219 | 0,13847148 | -3,575835386 | 3,49E-04 | 0,008699476 |
| Psen1 | protein_coding | 2165,798965 | -0,495090952 | 0,148606921 | -3,331547067 | 8,64E-04 | 0,016068481 |
| Rac1 | protein_coding | 9058,441351 | -0,494269886 | 0,161691358 | -3,05687263 | 0,002236593 | 0,030458795 |
| Akr1a1 | protein_coding | 5376,298978 | -0,493365492 | 0,162250523 | -3,040763642 | 0,00235979 | 0,031680938 |
| Ezr | protein_coding | 8470,460952 | -0,492816852 | 0,15173132 | -3,247957329 | 0,001162367 | 0,01944875 |
| Bsdc1 | protein_coding | 650,942346 | -0,491147413 | 0,154121139 | -3,186762158 | 0,00143875 | 0,022403793 |
| Zbp1 | protein_coding | 1779,077702 | -0,491122222 | 0,166195188 | -2,955092906 | 0,003125749 | 0,038171399 |
| Slc44a2 | protein_coding | 11306,47577 | -0,490402323 | 0,162949271 | -3,009539836 | 0,002616438 | 0,034190866 |
| Rnf114 | protein_coding | 5571,11695 | -0,490393914 | 0,132739037 | -3,694421207 | 2,20E-04 | 0,00618877 |
| Acss1 | protein_coding | 1047,037069 | -0,489567221 | 0,155871968 | -3,140829148 | 0,001684703 | 0,025188584 |
| Cdc42se1 | protein_coding | 6675,097927 | -0,489518348 | 0,149511325 | -3,274122201 | 0,001059908 | 0,018285059 |
| Bcap31 | protein_coding | 3536,310944 | -0,489137448 | 0,147646805 | -3,312888816 | 9,23E-04 | 0,01676626 |
| Map1lc3b | protein_coding | 3824,695461 | -0,489103658 | 0,127089968 | -3,848483615 | 1,19E-04 | 0,003901523 |
| Ttc7 | protein_coding | 1182,36241 | -0,488189766 | 0,169995907 | -2,87177365 | 0,004081752 | 0,045616672 |
| Sep.07 | protein_coding | 8151,216622 | -0,487395228 | 0,168871663 | -2,886187175 | 0,003899402 | 0,044083428 |
| Rela | protein_coding | 3773,016246 | -0,487153492 | 0,102766326 | -4,740400016 | 2,13E-06 | 1,65E-04 |
| Bid | protein_coding | 890,2264507 | -0,48550613 | 0,136087229 | -3,567609793 | 3,60E-04 | 0,008883729 |
| Prkch | protein_coding | 6570,942762 | -0,485104265 | 0,148994753 | -3,255847978 | 0,001130543 | 0,019184546 |
| Eps15 | protein_coding | 1853,110392 | -0,484742926 | 0,166981216 | -2,902978778 | 0,003696317 | 0,042511486 |
| Reep5 | protein_coding | 2680,160589 | -0,484725499 | 0,12446053 | -3,894612207 | 9,84E-05 | 0,003317714 |
| Rgs14 | protein_coding | 2545,779481 | -0,484419159 | 0,105585552 | -4,58793034 | 4,48E-06 | 3,05E-04 |
| Sun2 | protein_coding | 3932,709442 | -0,484266246 | 0,141230772 | -3,428900359 | 6,06E-04 | 0,012555334 |
| Dhx40 | protein_coding | 910,3261397 | -0,48382969 | 0,169536002 | -2,853846291 | 0,004319342 | 0,047253663 |
| Aldh4a1 | protein_coding | 361,7954409 | -0,483634311 | 0,159919431 | -3,024237318 | 0,002492608 | 0,032987457 |
| Tnfrsf1b | protein_coding | 5898,514631 | -0,483582084 | 0,115472143 | -4,187867934 | 2,82E-05 | 0,001331402 |
| Cant1 | protein_coding | 2342,685831 | -0,482990367 | 0,123652439 | -3,90603187 | 9,38E-05 | 0,003230509 |
| Hk1 | protein_coding | 2953,706054 | -0,482266213 | 0,151337078 | -3,186702287 | 0,001439048 | 0,022403793 |
| Vapa | protein_coding | 3041,186268 | -0,481817303 | 0,154612452 | -3,116290421 | 0,001831418 | 0,026452754 |
| Cd164 | protein_coding | 6319,244115 | -0,48018581 | 0,164984617 | -2,910488387 | 0,003608644 | 0,042026447 |
| Hsdl2 | protein_coding | 768,5394308 | -0,479187369 | 0,1142153 | -4,195474409 | 2,72E-05 | 0,001298594 |
| Eef1akmt1 | protein_coding | 837,2065494 | -0,478766913 | 0,164649942 | -2,907786708 | 0,003639965 | 0,042218396 |
| Cep250 | protein_coding | 898,8926846 | -0,477965579 | 0,144941829 | -3,297637287 | 9,75E-04 | 0,017269404 |
| Zdhhc20 | protein_coding | 3565,963457 | -0,477525231 | 0,098742276 | -4,836076791 | 1,32E-06 | 1,09E-04 |
| Sdha | protein_coding | 6702,717852 | -0,477520834 | 0,152569683 | -3,129854007 | 0,001748932 | 0,025728143 |
| Aaed1 | protein_coding | 527,8124585 | -0,477356996 | 0,141502976 | -3,373476728 | 7,42E-04 | 0,014443084 |
| Cers4 | protein_coding | 990,4217598 | -0,477354262 | 0,148929149 | -3,205244003 | 0,00134948 | 0,021452083 |
| Ubac2 | protein_coding | 2813,126613 | -0,477274044 | 0,124664075 | -3,828481015 | 1,29E-04 | 0,00414696 |
| Irf9 | protein_coding | 4809,250247 | -0,477254434 | 0,104086285 | -4,585180789 | 4,54E-06 | 3,07E-04 |
| Ncbp3 | protein_coding | 903,8761456 | -0,476228174 | 0,162558242 | -2,929584917 | 0,003394151 | 0,040379443 |
| Mob3a | protein_coding | 1875,147683 | -0,475457019 | 0,167527059 | -2,838090888 | 0,004538426 | 0,048940685 |
| Stk10 | protein_coding | 6862,094247 | -0,475166132 | 0,129090048 | -3,680888943 | 2,32E-04 | 0,006396815 |
| Timm29 | protein_coding | 894,9715434 | -0,474942205 | 0,120698001 | -3,934963296 | 8,32E-05 | 0,002963398 |
| Ramp1 | protein_coding | 981,2918663 | -0,474735343 | 0,147257742 | -3,223839621 | 0,001264842 | 0,020499336 |

| | | | | | | | |
|---|---|---|---|---|---|---|---|
| Tap2 | protein_coding | 4768,791956 | -0,474679857 | 0,13112242 | -3,620127333 | 2,94E-04 | 0,007553054 |
| Prkca | protein_coding | 2030,527662 | -0,472858034 | 0,137527012 | -3,438292075 | 5,85E-04 | 0,012282213 |
| Nek7 | protein_coding | 2189,867163 | -0,47281214 | 0,146973172 | -3,216996226 | 0,001295403 | 0,020852094 |
| 2610021A01Rik | protein_coding | 443,4530092 | -0,471689237 | 0,161315383 | -2,92401896 | 0,003455437 | 0,040729 |
| Pcmtd2 | protein_coding | 1410,218566 | -0,470286363 | 0,14989334 | -3,137473379 | 0,001704107 | 0,025316597 |
| Ap3m1 | protein_coding | 1003,907779 | -0,469676543 | 0,119602229 | -3,926988213 | 8,60E-05 | 0,003030838 |
| Rmnd5b | protein_coding | 1337,433709 | -0,469306342 | 0,121587465 | -3,859825039 | 1,13E-04 | 0,003772149 |
| Zfp148 | protein_coding | 1463,872379 | -0,468864092 | 0,148642982 | -3,154296855 | 0,001608853 | 0,024398555 |
| Mfap1b | protein_coding | 2165,144849 | -0,468471986 | 0,102359979 | -4,576710458 | 4,72E-06 | 3,17E-04 |
| Cybc1 | protein_coding | 2604,921443 | -0,467828797 | 0,126468428 | -3,6991746 | 2,16E-04 | 0,006113917 |
| H2-T22 | protein_coding | 4312,687347 | -0,466400316 | 0,164032935 | -2,843333366 | 0,004464435 | 0,048521126 |
| Parvg | protein_coding | 2962,596953 | -0,466199856 | 0,10045839 | -4,640725954 | 3,47E-06 | 2,48E-04 |
| Cd3e | protein_coding | 8224,319058 | -0,465811937 | 0,160437175 | -2,90339154 | 0,003691448 | 0,042511486 |
| Cpne3 | protein_coding | 631,33271 | -0,465678019 | 0,146301586 | -3,183000489 | 0,001457574 | 0,02267094 |
| Maf1 | protein_coding | 5340,937651 | -0,465674495 | 0,14345623 | -3,246108545 | 0,001169942 | 0,019513931 |
| Coro1b | protein_coding | 6224,581819 | -0,465031347 | 0,124160358 | -3,74540921 | 1,80E-04 | 0,005352084 |
| Leprot | protein_coding | 1170,140303 | -0,463982669 | 0,126144918 | -3,678171719 | 2,35E-04 | 0,006423841 |
| Micu2 | protein_coding | 1599,495092 | -0,463654393 | 0,146415078 | -3,166712056 | 0,001541729 | 0,023582058 |
| Parp10 | protein_coding | 2017,342474 | -0,46003349 | 0,121948584 | -3,772356131 | 1,62E-04 | 0,004997751 |
| Ttc3 | protein_coding | 1050,33883 | -0,459814047 | 0,142611494 | -3,224242541 | 0,001263064 | 0,020497763 |
| Sfxn3 | protein_coding | 2309,273316 | -0,457534593 | 0,151352925 | -3,022964991 | 0,002503112 | 0,033021977 |
| Rpa1 | protein_coding | 3553,047412 | -0,457518049 | 0,1291552 | -3,542389693 | 3,97E-04 | 0,0094101 |
| Gm4070 | protein_coding | 1474,386029 | -0,457510123 | 0,134512244 | -3,40125263 | 6,71E-04 | 0,013477859 |
| Gipc1 | protein_coding | 819,8592851 | -0,457173911 | 0,154178445 | -2,96522585 | 0,003024608 | 0,03736633 |
| Acap2 | protein_coding | 866,0017396 | -0,457085795 | 0,137857112 | -3,315648992 | 9,14E-04 | 0,016692314 |
| Itpk1 | protein_coding | 1023,608289 | -0,456838525 | 0,121369432 | -3,764032811 | 1,67E-04 | 0,005100675 |
| Ptpre | protein_coding | 700,1243794 | -0,454826182 | 0,134617426 | -3,378657544 | 7,28E-04 | 0,014255471 |
| Slc28a2 | protein_coding | 2698,216174 | -0,454796929 | 0,110537859 | -4,114399639 | 3,88E-05 | 0,001704332 |
| Rabgap1l | protein_coding | 2357,767127 | -0,452944297 | 0,12516302 | -3,618834847 | 2,96E-04 | 0,007579158 |
| Phf1 | protein_coding | 2110,637216 | -0,45245677 | 0,12755109 | -3,547259134 | 3,89E-04 | 0,009322055 |
| Hdac1 | protein_coding | 3903,471131 | -0,451761287 | 0,109175253 | -4,137945875 | 3,50E-05 | 0,001568203 |
| Sep.06 | protein_coding | 4745,80098 | -0,448944186 | 0,135557257 | -3,311841768 | 9,27E-04 | 0,01677776 |
| Spata13 | protein_coding | 3302,808783 | -0,447528457 | 0,111545482 | -4,012071568 | 6,02E-05 | 0,002339307 |
| Jak1 | protein_coding | 14024,21457 | -0,446402146 | 0,150451156 | -2,967090162 | 0,003006328 | 0,037233595 |
| Trbv19 | TR_V_gene | 1032,281028 | -0,446100311 | 0,1250164 | -3,568334317 | 3,59E-04 | 0,008872388 |
| Il21r | protein_coding | 5452,30331 | -0,446054325 | 0,140613428 | -3,172202904 | 0,001512873 | 0,023334233 |
| Slc2a1 | protein_coding | 1029,597236 | -0,445324386 | 0,142959045 | -3,115048695 | 0,001839145 | 0,026531886 |
| Tmem59 | protein_coding | 4374,261295 | -0,445075335 | 0,152126457 | -2,925693171 | 0,003436897 | 0,040568099 |
| Hadha | protein_coding | 2543,098333 | -0,443539974 | 0,156708198 | -2,830355913 | 0,004649625 | 0,049464856 |
| Puf60 | protein_coding | 5185,314763 | -0,442719143 | 0,145301105 | -3,046908296 | 0,002312082 | 0,031170852 |
| Sigirr | protein_coding | 3258,574827 | -0,442680026 | 0,120113104 | -3,685526485 | 2,28E-04 | 0,006312851 |
| Arl5c | protein_coding | 1864,381714 | -0,442538694 | 0,133517256 | -3,314468163 | 9,18E-04 | 0,016726734 |
| Sit1 | protein_coding | 2307,293813 | -0,442159857 | 0,151041398 | -2,927408391 | 0,003417997 | 0,040422508 |
| Syngr2 | protein_coding | 2009,155166 | -0,441873261 | 0,137101661 | -3,222960666 | 0,00126873 | 0,02052226 |
| Ncln | protein_coding | 2644,025791 | -0,440774937 | 0,117186665 | -3,761306281 | 1,69E-04 | 0,005147144 |
| Hbp1 | protein_coding | 3051,282517 | -0,438458718 | 0,113189823 | -3,873658494 | 1,07E-04 | 0,00358077 |
| Gba | protein_coding | 1484,284298 | -0,4366142 | 0,136279159 | -3,203822223 | 0,001356162 | 0,021520401 |
| Hexb | protein_coding | 1263,362899 | -0,433170645 | 0,131999732 | -3,281602454 | 0,00103219 | 0,017918649 |
| Casp8 | protein_coding | 4706,487289 | -0,432399392 | 0,100707117 | -4,293632895 | 1,76E-05 | 9,03E-04 |
| Rpn2 | protein_coding | 4255,916789 | -0,431661787 | 0,130847437 | -3,298970143 | 9,70E-04 | 0,017236453 |
| Scamp2 | protein_coding | 1924,884464 | -0,430850314 | 0,12570607 | -3,427442402 | 6,09E-04 | 0,012592614 |
| Mgrn1 | protein_coding | 1296,790595 | -0,430422076 | 0,114889129 | -3,746412569 | 1,79E-04 | 0,005344731 |
| Ccm2 | protein_coding | 4368,931973 | -0,429768795 | 0,112872042 | -3,807575264 | 1,40E-04 | 0,004466302 |
| Tbc1d14 | protein_coding | 1749,87439 | -0,429326338 | 0,137223347 | -3,128668327 | 0,001756004 | 0,025789949 |
| Rhof | protein_coding | 5208,565956 | -0,429247755 | 0,109172806 | -3,931819376 | 8,43E-05 | 0,002996001 |
| Arl6ip5 | protein_coding | 2985,183536 | -0,426907462 | 0,141446913 | -3,018146194 | 0,002543262 | 0,033392384 |
| Tnfrsf25 | protein_coding | 3473,921956 | -0,426407721 | 0,132768987 | -3,21165154 | 0,001319743 | 0,021120986 |
| Anp32e | protein_coding | 1965,26434 | -0,426177809 | 0,143674869 | -2,966265512 | 0,003014401 | 0,037305745 |
| Crbn | protein_coding | 1622,659351 | -0,425939117 | 0,146483719 | -2,907757383 | 0,003640306 | 0,042218396 |
| Derl1 | protein_coding | 2706,21243 | -0,422572278 | 0,098839718 | -4,275328665 | 1,91E-05 | 9,60E-04 |
| Zfp869 | protein_coding | 941,5335482 | -0,422532947 | 0,111169768 | -3,800790047 | 1,44E-04 | 0,004559489 |
| Dtx1 | protein_coding | 4250,235398 | -0,42238807 | 0,126265477 | -3,345237989 | 8,22E-04 | 0,015557472 |
| Vps35 | protein_coding | 2544,223877 | -0,422368623 | 0,13769189 | -3,067490924 | 0,00215864 | 0,029745752 |
| Ciz1 | protein_coding | 1472,098113 | -0,421898512 | 0,115830523 | -3,642377689 | 2,70E-04 | 0,007093512 |
| Bin1 | protein_coding | 2487,290294 | -0,421214234 | 0,134833198 | -3,123965307 | 0,001784315 | 0,026067333 |
| Ilvbl | protein_coding | 1503,257346 | -0,421176458 | 0,138580588 | -3,039216854 | 0,002371941 | 0,031745746 |
| Plekhg2 | protein_coding | 2650,001836 | -0,417441213 | 0,128514075 | -3,248213963 | 0,001161319 | 0,01944875 |
| Nfyc | protein_coding | 1176,661319 | -0,417157095 | 0,114019103 | -3,658659694 | 2,54E-04 | 0,00680935 |
| Ccndbp1 | protein_coding | 1827,550969 | -0,416724241 | 0,141839329 | -2,93800206 | 0,003303348 | 0,039611538 |

| | | | | | | | |
|---|---|---|---|---|---|---|---|
| Actr10 | protein_coding | 1979,103706 | -0,416451931 | 0,114169404 | -3,647666679 | 2,65E-04 | 0,006960132 |
| Xlr4b | protein_coding | 1614,763977 | -0,415646445 | 0,139072544 | -2,988702393 | 0,002801649 | 0,03590924 |
| Tfip11 | protein_coding | 685,3808817 | -0,415566669 | 0,142183806 | -2,92274261 | 0,003469632 | 0,040839818 |
| Lipa | protein_coding | 2541,079182 | -0,415067149 | 0,119049934 | -3,486496268 | 4,89E-04 | 0,010800287 |
| Ano6 | protein_coding | 1234,616765 | -0,412807163 | 0,142618997 | -2,894475284 | 0,003797928 | 0,043201107 |
| Dhx8 | protein_coding | 1290,321674 | -0,412347741 | 0,142034843 | -2,903144987 | 0,003694356 | 0,042511486 |
| Fam53b | protein_coding | 3627,261443 | -0,411587478 | 0,118496434 | -3,473416574 | 5,14E-04 | 0,011119041 |
| Sufu | protein_coding | 864,7829828 | -0,411155955 | 0,142044082 | -2,894565898 | 0,003796832 | 0,043201107 |
| Lamp1 | protein_coding | 4960,107676 | -0,411063877 | 0,137841248 | -2,982154339 | 0,002862277 | 0,036150946 |
| Il10rb | protein_coding | 1468,135872 | -0,410947328 | 0,098524753 | -4,171005913 | 3,03E-05 | 0,001421715 |
| Vps11 | protein_coding | 1382,824902 | -0,409576342 | 0,129403588 | -3,165108078 | 0,001550254 | 0,023668821 |
| Acot9 | protein_coding | 1500,556473 | -0,408452094 | 0,122200568 | -3,34247297 | 8,30E-04 | 0,01565973 |
| Iqgap1 | protein_coding | 3599,23205 | -0,407334519 | 0,096533208 | -4,219631028 | 2,45E-05 | 0,001183989 |
| Wwp2 | protein_coding | 2237,564753 | -0,40690738 | 0,088984603 | -4,572784104 | 4,81E-06 | 3,22E-04 |
| Lrrc61 | protein_coding | 757,6662455 | -0,406459246 | 0,138628315 | -2,932007392 | 0,003367787 | 0,040181019 |
| Mbp | protein_coding | 2404,527521 | -0,406405527 | 0,133065174 | -3,054184009 | 0,002256736 | 0,030598687 |
| Frmd8 | protein_coding | 6710,632735 | -0,406273547 | 0,137720589 | -2,949984097 | 0,003177903 | 0,038666038 |
| Nmi | protein_coding | 1717,176376 | -0,406168088 | 0,108145585 | -3,755752828 | 1,73E-04 | 0,005220917 |
| Gsto1 | protein_coding | 799,2558928 | -0,40465963 | 0,142171923 | -2,846269644 | 0,004423472 | 0,048138979 |
| Tex261 | protein_coding | 2311,372054 | -0,403203934 | 0,124164103 | -3,247347065 | 0,001164862 | 0,01944875 |
| Arhgap27 | protein_coding | 2674,016262 | -0,402995661 | 0,092411841 | -4,360866067 | 1,30E-05 | 7,19E-04 |
| Clcf1 | protein_coding | 1016,569987 | -0,402763281 | 0,10925535 | -3,686439885 | 2,27E-04 | 0,006312851 |
| Efr3a | protein_coding | 3754,400052 | -0,402048398 | 0,132281341 | -3,03934323 | 0,002370946 | 0,031745746 |
| Gata3 | protein_coding | 4303,836616 | -0,401368944 | 0,127737553 | -3,142137404 | 0,001677193 | 0,025121567 |
| Arid4b | protein_coding | 1824,591159 | -0,400603363 | 0,125460453 | -3,193064863 | 0,001407713 | 0,022081672 |
| Ctps2 | protein_coding | 2487,412362 | -0,400558588 | 0,127641906 | -3,138143273 | 0,001700217 | 0,025283879 |
| Man2b2 | protein_coding | 1410,147752 | -0,400379987 | 0,12323238 | -3,248983654 | 0,001158181 | 0,01944875 |
| Bcl2l1 | protein_coding | 1642,83626 | -0,399520182 | 0,137645904 | -2,902521406 | 0,003701719 | 0,042536227 |
| Casp6 | protein_coding | 714,3984379 | -0,399374972 | 0,111349113 | -3,586691989 | 3,35E-04 | 0,008383096 |
| Kcnab2 | protein_coding | 2948,990118 | -0,398731651 | 0,113568594 | -3,510932359 | 4,47E-04 | 0,010128011 |
| Vps28 | protein_coding | 4539,055904 | -0,39869363 | 0,139856354 | -2,850736624 | 0,004361808 | 0,047624059 |
| Abtb1 | protein_coding | 1560,302391 | -0,396202195 | 0,107422264 | -3,688268895 | 2,26E-04 | 0,006297693 |
| Trex1 | protein_coding | 1449,105666 | -0,396104452 | 0,095370453 | -4,153324654 | 3,28E-05 | 0,001509495 |
| Fam111a | protein_coding | 1006,690476 | -0,395115615 | 0,115710434 | -3,414693056 | 6,39E-04 | 0,012991983 |
| Inpp5f | protein_coding | 1391,506087 | -0,39420266 | 0,132948447 | -2,96507908 | 0,003026051 | 0,03736633 |
| Mtmr6 | protein_coding | 1483,629696 | -0,393407072 | 0,128866517 | -3,052826136 | 0,002266972 | 0,030712387 |
| Cbx7 | protein_coding | 5212,366002 | -0,391936207 | 0,10943171 | -3,581559744 | 3,42E-04 | 0,008536668 |
| Ip6k1 | protein_coding | 1772,560913 | -0,39150899 | 0,122820347 | -3,187655776 | 0,001434312 | 0,022402605 |
| Akap8 | protein_coding | 4462,46343 | -0,39074398 | 0,116016175 | -3,36801295 | 7,57E-04 | 0,014661809 |
| Tagap | protein_coding | 2062,659073 | -0,390489954 | 0,116431341 | -3,353821682 | 7,97E-04 | 0,015221666 |
| Pou2f2 | protein_coding | 1535,121603 | -0,390046682 | 0,118975901 | -3,278367126 | 0,001044095 | 0,018068615 |
| Supt20 | protein_coding | 3392,849784 | -0,38948555 | 0,091527762 | -4,255381575 | 2,09E-05 | 0,001036966 |
| Ubl7 | protein_coding | 1787,521882 | -0,38939061 | 0,108541398 | -3,587484759 | 3,34E-04 | 0,008370281 |
| Prkd2 | protein_coding | 4354,735584 | -0,389044549 | 0,087963163 | -4,422812203 | 9,74E-06 | 5,73E-04 |
| Rbmx | protein_coding | 969,1058109 | -0,387615521 | 0,104145668 | -3,721859266 | 1,98E-04 | 0,005688121 |
| Lasp1 | protein_coding | 5800,927879 | -0,385982933 | 0,106281557 | -3,631701892 | 2,82E-04 | 0,007324045 |
| Fam3c | protein_coding | 2193,434742 | -0,385513984 | 0,109282369 | -3,527686936 | 4,19E-04 | 0,009716718 |
| Ap1g2 | protein_coding | 2416,133079 | -0,384870241 | 0,134983776 | -2,851233337 | 0,004355 | 0,047599463 |
| Igf2r | protein_coding | 2628,505035 | -0,383818819 | 0,117429776 | -3,268496548 | 0,001081205 | 0,018556026 |
| Uba1 | protein_coding | 6091,603311 | -0,382889758 | 0,120901269 | -3,166962287 | 0,001540403 | 0,023582058 |
| Uba7 | protein_coding | 4450,219266 | -0,382499353 | 0,128593312 | -2,974488701 | 0,002934773 | 0,036620667 |
| Psmc3 | protein_coding | 3503,886304 | -0,379451096 | 0,127048324 | -2,986667472 | 0,002820363 | 0,03590924 |
| Fam189b | protein_coding | 4481,3678 | -0,379025396 | 0,130606654 | -2,902037418 | 0,003707443 | 0,042550979 |
| Tapbpl | protein_coding | 2008,855007 | -0,375069764 | 0,121633063 | -3,083616858 | 0,002045008 | 0,028592206 |
| Rftn1 | protein_coding | 1964,406038 | -0,374622198 | 0,119509774 | -3,134657403 | 0,001720549 | 0,025474028 |
| Pdlim2 | protein_coding | 1921,67166 | -0,374061891 | 0,11947125 | -3,130978295 | 0,00174225 | 0,025678852 |
| Ssh2 | protein_coding | 2967,032176 | -0,373581234 | 0,113566993 | -3,289522982 | 0,001003574 | 0,017550377 |
| Snx2 | protein_coding | 2654,13254 | -0,372633344 | 0,1102414 | -3,380157934 | 7,24E-04 | 0,014245068 |
| Wsb1 | protein_coding | 1951,513781 | -0,370084531 | 0,121356603 | -3,049562372 | 0,00229175 | 0,030947022 |
| Selenok | protein_coding | 2005,541596 | -0,369981697 | 0,112452407 | -3,290118066 | 0,001001454 | 0,017544143 |
| Stat3 | protein_coding | 5153,839568 | -0,369463884 | 0,121085818 | -3,051256467 | 0,002278858 | 0,030831378 |
| Rnf31 | protein_coding | 1443,235668 | -0,367456725 | 0,12689426 | -2,895771048 | 0,003782282 | 0,043141415 |
| Skap1 | protein_coding | 7781,855336 | -0,36742408 | 0,12359565 | -2,972791366 | 0,00295105 | 0,036740901 |
| Fam32a | protein_coding | 3651,174359 | -0,367200005 | 0,11431875 | -3,212071545 | 0,001317816 | 0,021111915 |
| Fmnl1 | protein_coding | 6230,826406 | -0,366344824 | 0,118091066 | -3,10222303 | 0,001920732 | 0,027314881 |
| Cnot2 | protein_coding | 2387,482969 | -0,365814127 | 0,102617348 | -3,564837075 | 3,64E-04 | 0,008898836 |
| Tgoln1 | protein_coding | 2453,801528 | -0,3648953 | 0,121676911 | -2,998886936 | 0,002709678 | 0,035132674 |
| Mar.02 | protein_coding | 1666,120856 | -0,36468021 | 0,122847385 | -2,968563077 | 0,002991957 | 0,037110999 |
| Inpp1 | protein_coding | 1280,041869 | -0,3641751 | 0,119365221 | -3,050931397 | 0,002281327 | 0,030831378 |

| | | | | | | | |
|---|---|---|---|---|---|---|---|
| Ptpn6 | protein_coding | 5027,237088 | -0,364054376 | 0,11051156 | -3,294265117 | 9,87E-04 | 0,017399228 |
| Slfn10-ps | transcribed_unproce | 981,1386457 | -0,36404432 | 0,11589476 | -3,141162896 | 0,001682784 | 0,025182583 |
| Vav1 | protein_coding | 4182,744558 | -0,362144841 | 0,096419821 | -3,755916977 | 1,73E-04 | 0,005220917 |
| Tecpr1 | protein_coding | 4799,2094 | -0,359156305 | 0,125460941 | -2,862694183 | 0,004200558 | 0,046413087 |
| Pdha1 | protein_coding | 2610,919514 | -0,357860859 | 0,119112865 | -3,004384631 | 0,002661187 | 0,034612111 |
| Cdipt | protein_coding | 5247,86932 | -0,357594731 | 0,109956187 | -3,252156516 | 0,001145329 | 0,019356299 |
| Abcf3 | protein_coding | 1428,449741 | -0,355123103 | 0,107029403 | -3,317995747 | 9,07E-04 | 0,016608065 |
| Tnip1 | protein_coding | 4489,597281 | -0,354729014 | 0,094456944 | -3,755457239 | 1,73E-04 | 0,005220917 |
| Dapp1 | protein_coding | 2118,299755 | -0,354245499 | 0,118754581 | -2,983004915 | 0,002854334 | 0,036150946 |
| Sec24c | protein_coding | 4750,957237 | -0,353765411 | 0,119335061 | -2,964471694 | 0,003032031 | 0,037412336 |
| Atp6v1d | protein_coding | 2288,508269 | -0,353737692 | 0,116496555 | -3,036464838 | 0,0023937 | 0,031985386 |
| Glud1 | protein_coding | 3295,072981 | -0,353721172 | 0,121667947 | -2,907266713 | 0,003646022 | 0,042254061 |
| Taz | protein_coding | 2133,713981 | -0,353012768 | 0,120841264 | -2,921293235 | 0,003485816 | 0,040941682 |
| Arhgef2 | protein_coding | 2262,553115 | -0,352468468 | 0,109733109 | -3,212052133 | 0,001317905 | 0,021111915 |
| Tars2 | protein_coding | 1249,707285 | -0,352460076 | 0,124502036 | -2,83095832 | 0,004640877 | 0,049435168 |
| Trim27 | protein_coding | 2429,334541 | -0,35243544 | 0,121797192 | -2,893625322 | 0,003808223 | 0,043258909 |
| Irf3 | protein_coding | 3178,334372 | -0,351359753 | 0,113338351 | -3,100095871 | 0,00193458 | 0,027464749 |
| Hsd17b4 | protein_coding | 1253,993095 | -0,35034686 | 0,117512247 | -2,981364655 | 0,002869669 | 0,036216746 |
| Dcaf11 | protein_coding | 2659,020377 | -0,349718989 | 0,110009036 | -3,179002393 | 0,001477829 | 0,022857451 |
| Psmd3 | protein_coding | 2060,12974 | -0,345347915 | 0,116311447 | -2,969165315 | 0,002986099 | 0,037066043 |
| Sars | protein_coding | 2689,240807 | -0,344811882 | 0,108784229 | -3,169686307 | 0,001526036 | 0,023450085 |
| Sin3b | protein_coding | 2394,362908 | -0,342247524 | 0,116716906 | -2,932287497 | 0,003364751 | 0,040173675 |
| Ambra1 | protein_coding | 947,0731985 | -0,341134703 | 0,116999295 | -2,915698801 | 0,003548929 | 0,041506716 |
| Nrbp1 | protein_coding | 3182,813874 | -0,338215518 | 0,106066672 | -3,188706786 | 0,001429108 | 0,022353882 |
| Ss18 | protein_coding | 2188,53135 | -0,338104558 | 0,085387861 | -3,95963262 | 7,51E-05 | 0,002756153 |
| Supt5 | protein_coding | 4230,220276 | -0,337210197 | 0,09810567 | -3,437214139 | 5,88E-04 | 0,012315631 |
| Uba2 | protein_coding | 2727,338294 | -0,334686554 | 0,113194166 | -2,956747369 | 0,003109027 | 0,03799933 |
| 9930111J21Rik1 | protein_coding | 1559,997511 | -0,333878824 | 0,110695819 | -3,016182781 | 0,002559789 | 0,033582813 |
| Ptpn7 | protein_coding | 6284,670365 | -0,331179963 | 0,088779942 | -3,730346683 | 1,91E-04 | 0,005583364 |
| Atf7ip | protein_coding | 6503,65491 | -0,329877293 | 0,116739929 | -2,825745195 | 0,004717077 | 0,049958271 |
| Mpp1 | protein_coding | 1636,57542 | -0,329512715 | 0,112772209 | -2,921931915 | 0,003478676 | 0,040886758 |
| Adcy7 | protein_coding | 2689,450592 | -0,327844848 | 0,106090018 | -3,090251605 | 0,00199987 | 0,02817474 |
| Rab5b | protein_coding | 1618,128578 | -0,326900421 | 0,113869186 | -2,870841823 | 0,004093803 | 0,045720564 |
| Prkag2 | protein_coding | 894,5332154 | -0,326189194 | 0,115217246 | -2,83107958 | 0,004639118 | 0,049435168 |
| Cers2 | protein_coding | 4037,803462 | -0,324824327 | 0,112013421 | -2,899869722 | 0,003733178 | 0,042728152 |
| Hp1bp3 | protein_coding | 4103,104189 | -0,324479982 | 0,108479634 | -2,99116037 | 0,002779195 | 0,035782405 |
| Rin3 | protein_coding | 2996,864759 | -0,324291846 | 0,080761586 | -4,015421961 | 5,93E-05 | 0,002322639 |
| Lonp2 | protein_coding | 2323,490685 | -0,318731137 | 0,109052502 | -2,922731085 | 0,00346976 | 0,040839818 |
| Rasgrp1 | protein_coding | 6268,010735 | -0,317173795 | 0,111718171 | -2,839052878 | 0,004524766 | 0,048905778 |
| Cog5 | protein_coding | 770,3624111 | -0,317074757 | 0,111265855 | -2,849704038 | 0,004375993 | 0,047716147 |
| Phc3 | protein_coding | 2807,765566 | -0,316640465 | 0,108744361 | -2,911787453 | 0,003593671 | 0,041882416 |
| Skiv2l | protein_coding | 3017,407817 | -0,316494118 | 0,078263896 | -4,043935134 | 5,26E-05 | 0,002127586 |
| Mapre1 | protein_coding | 4238,953113 | -0,316144338 | 0,102322211 | -3,089694176 | 0,002003627 | 0,028203725 |
| Smu1 | protein_coding | 2872,035186 | -0,313993726 | 0,100676584 | -3,11883572 | 0,001815672 | 0,026344525 |
| Arhgef3 | protein_coding | 6674,149245 | -0,313532234 | 0,102823109 | -3,049238985 | 0,002294219 | 0,030955169 |
| Camta2 | protein_coding | 898,2328953 | -0,311199446 | 0,108641291 | -2,864467495 | 0,00417711 | 0,046246379 |
| Actr1a | protein_coding | 2529,340541 | -0,307863559 | 0,103023847 | -2,988274746 | 0,002805572 | 0,03590924 |
| Nbr1 | protein_coding | 1772,651005 | -0,305359366 | 0,09712147 | -3,144097455 | 0,001665999 | 0,02499903 |
| Anapc2 | protein_coding | 2396,016158 | -0,305236854 | 0,076460718 | -3,99207414 | 6,55E-05 | 0,002470461 |
| Copa | protein_coding | 5199,003444 | -0,30374725 | 0,101312876 | -2,998111011 | 0,002716587 | 0,035170538 |
| Gsdmd | protein_coding | 2051,724617 | -0,303623046 | 0,098537978 | -3,081279459 | 0,002061131 | 0,028744983 |
| Akap8l | protein_coding | 3015,516527 | -0,303333923 | 0,099192055 | -3,058046561 | 0,00222785 | 0,030448068 |
| Eftud2 | protein_coding | 2037,348534 | -0,303205811 | 0,105733473 | -2,867642605 | 0,004135424 | 0,04609234 |
| Smc1a | protein_coding | 2067,681205 | -0,300699244 | 0,105717212 | -2,844373579 | 0,004449884 | 0,048394676 |
| Gltp | protein_coding | 4958,379572 | -0,299499774 | 0,104491562 | -2,86625798 | 0,004153556 | 0,046246379 |
| St6gal1 | protein_coding | 4145,209994 | -0,298431214 | 0,09182553 | -3,249980831 | 0,001154128 | 0,019429256 |
| Abhd8 | protein_coding | 3265,410068 | -0,298235858 | 0,100189156 | -2,976727921 | 0,002913424 | 0,036435303 |
| Clcn7 | protein_coding | 962,1858166 | -0,297069974 | 0,101165886 | -2,936463903 | 0,003319775 | 0,039779768 |
| Tes | protein_coding | 4028,699559 | -0,295728577 | 0,100293009 | -2,948645956 | 0,003191694 | 0,038748611 |
| Ppm1m | protein_coding | 1863,097091 | -0,292361208 | 0,097885226 | -2,986775644 | 0,002819365 | 0,03590924 |
| Myo1g | protein_coding | 5030,082851 | -0,27511204 | 0,095142832 | -2,891568742 | 0,003833237 | 0,043453829 |
| Psd4 | protein_coding | 4093,471617 | -0,263561475 | 0,078496321 | -3,357628389 | 7,86E-04 | 0,015121975 |
| Vps4b | protein_coding | 2696,766482 | -0,25392323 | 0,088622375 | -2,865227105 | 0,004167103 | 0,046246379 |
| Paics | protein_coding | 2386,385533 | 0,293197677 | 0,082070755 | 3,572498839 | 3,54E-04 | 0,008782356 |
| Gimap1os | processed_transcript | 1465,520952 | 0,296248818 | 0,104377821 | 2,838235318 | 0,004536373 | 0,048940685 |
| Srrm2 | protein_coding | 4231,825853 | 0,298112322 | 0,095812017 | 3,111429352 | 0,00186184 | 0,026763653 |
| Sod2 | protein_coding | 1635,841596 | 0,316540007 | 0,111703993 | 2,833739402 | 0,004600683 | 0,04935133 |
| Lysmd2 | protein_coding | 1044,971645 | 0,334312748 | 0,107783823 | 3,101696882 | 0,001924149 | 0,027340045 |
| Celf1 | protein_coding | 2325,964277 | 0,341645119 | 0,116008637 | 2,944997263 | 0,003229575 | 0,038980385 |

| | | | | | | | |
|---|---|---|---|---|---|---|---|
| Dgkz | protein_coding | 1585,566328 | 0,343554614 | 0,106549396 | 3,224369433 | 0,001262504 | 0,020497763 |
| Btla | protein_coding | 1988,185534 | 0,346132261 | 0,111249832 | 3,111305921 | 0,001862619 | 0,026763653 |
| Zbtb7b | protein_coding | 1157,775361 | 0,347778398 | 0,122788501 | 2,832336861 | 0,004620914 | 0,04935133 |
| Mrps17 | protein_coding | 1166,700687 | 0,352492714 | 0,120188056 | 2,932843133 | 0,003358735 | 0,040130723 |
| Pa2g4 | protein_coding | 2685,272437 | 0,357428475 | 0,101994091 | 3,504403736 | 4,58E-04 | 0,010346605 |
| Tor1aip2 | protein_coding | 994,5129593 | 0,364645303 | 0,114068629 | 3,196718558 | 0,001390005 | 0,021898291 |
| Macf1 | protein_coding | 3672,065808 | 0,369012805 | 0,076857681 | 4,801248213 | 1,58E-06 | 1,26E-04 |
| Tomm5 | protein_coding | 729,6727549 | 0,369809829 | 0,121997891 | 3,031280516 | 0,002435189 | 0,032430568 |
| Man1a | protein_coding | 1206,233092 | 0,371084586 | 0,124823649 | 2,972870839 | 0,002950286 | 0,036740901 |
| Wdr24 | protein_coding | 940,2742908 | 0,374592095 | 0,094941008 | 3,945524716 | 7,96E-05 | 0,002872747 |
| Art2b | protein_coding | 3436,406885 | 0,389726933 | 0,128816075 | 3,025452634 | 0,002482613 | 0,032882237 |
| Ago2 | protein_coding | 1252,623471 | 0,389747823 | 0,130897467 | 2,977504698 | 0,002906052 | 0,036435303 |
| Pop5 | protein_coding | 949,0347591 | 0,389821189 | 0,111975386 | 3,481311429 | 4,99E-04 | 0,010881502 |
| Mri1 | protein_coding | 1060,341012 | 0,394476064 | 0,099869311 | 3,949922764 | 7,82E-05 | 0,002832786 |
| Nup153 | protein_coding | 1360,519575 | 0,396004538 | 0,128184458 | 3,089333487 | 0,002006061 | 0,028214061 |
| Stat4 | protein_coding | 1476,470226 | 0,396786107 | 0,129711257 | 3,058995151 | 0,002220807 | 0,03040967 |
| Ndufab1 | protein_coding | 484,8801207 | 0,397472406 | 0,130633411 | 3,042655045 | 0,00234501 | 0,031537911 |
| Ndufa1 | protein_coding | 1141,9924 | 0,403486632 | 0,117862373 | 3,423371032 | 6,18E-04 | 0,012711322 |
| Dync1h1 | protein_coding | 1267,780181 | 0,406980984 | 0,123980452 | 3,282622187 | 0,001028464 | 0,017891393 |
| Prmt5 | protein_coding | 1099,558205 | 0,411455634 | 0,118248114 | 3,479595731 | 5,02E-04 | 0,010935156 |
| Rere | protein_coding | 1159,002085 | 0,413496708 | 0,127245596 | 3,249595442 | 0,001155693 | 0,019432502 |
| Huwe1 | protein_coding | 1909,776012 | 0,418528096 | 0,147696242 | 2,833708493 | 0,004601128 | 0,04935133 |
| Zfp444 | protein_coding | 698,4002341 | 0,419478743 | 0,134952769 | 3,10833743 | 0,001881431 | 0,026963932 |
| Ddx21 | protein_coding | 1884,994411 | 0,423383083 | 0,107961354 | 3,921617017 | 8,80E-05 | 0,003079598 |
| Nsmce3 | protein_coding | 614,4541613 | 0,424860615 | 0,138332168 | 3,071307431 | 0,002131236 | 0,029499575 |
| Aatf | protein_coding | 1075,591996 | 0,427652237 | 0,127228961 | 3,361280578 | 7,76E-04 | 0,014988945 |
| Ndufb1-ps | bidirectional_promo | 688,22125 | 0,433691155 | 0,131646387 | 3,294364289 | 9,86E-04 | 0,017399228 |
| Tmem126a | protein_coding | 688,9808606 | 0,434943998 | 0,14455255 | 3,008898817 | 0,002621964 | 0,034236128 |
| Trat1 | protein_coding | 2373,06644 | 0,43495166 | 0,130421311 | 3,334973848 | 8,53E-04 | 0,015950239 |
| Scaf1 | protein_coding | 582,4167094 | 0,437029464 | 0,147360029 | 2,965725967 | 0,003019694 | 0,037343402 |
| Pced1b | protein_coding | 984,598694 | 0,438688732 | 0,153133766 | 2,864742003 | 0,004173491 | 0,046246379 |
| Pycr2 | protein_coding | 1306,116404 | 0,44134284 | 0,152266701 | 2,898485596 | 0,003749695 | 0,042858088 |
| Zfp692 | protein_coding | 695,3051564 | 0,441961568 | 0,111310638 | 3,970524086 | 7,17E-05 | 0,002655239 |
| Eif4ebp2 | protein_coding | 1074,060267 | 0,442761804 | 0,138466071 | 3,197619474 | 0,00138567 | 0,021880665 |
| Ldb1 | protein_coding | 1035,914669 | 0,44395228 | 0,122145291 | 3,634624615 | 2,78E-04 | 0,00726429 |
| Phykpl | protein_coding | 280,2555521 | 0,444143842 | 0,152781293 | 2,907056438 | 0,003648474 | 0,042254061 |
| Usmg5 | protein_coding | 883,2849443 | 0,447930799 | 0,135546694 | 3,304623564 | 9,51E-04 | 0,017008046 |
| Apbb1ip | protein_coding | 582,0405713 | 0,451782559 | 0,140764879 | 3,20948352 | 0,001329737 | 0,021196582 |
| Usp36 | protein_coding | 776,1183711 | 0,45384223 | 0,151770541 | 2,99031833 | 0,002786868 | 0,035825615 |
| Uhrf1bp1l | protein_coding | 993,6198659 | 0,459270917 | 0,142265783 | 3,228259868 | 0,001245458 | 0,020424523 |
| Aimp2 | protein_coding | 399,563547 | 0,460579937 | 0,151736282 | 3,035397538 | 0,002402188 | 0,032072982 |
| Ap3s1 | protein_coding | 411,9247691 | 0,464507684 | 0,16272315 | 2,854588804 | 0,004309258 | 0,047205577 |
| Kri1 | protein_coding | 288,3240584 | 0,464778011 | 0,164060091 | 2,832974245 | 0,00461171 | 0,04935133 |
| Sdhd | protein_coding | 1379,970923 | 0,466005929 | 0,112619551 | 4,137877692 | 3,51E-05 | 0,001568203 |
| Mrpl57 | protein_coding | 655,2091248 | 0,47053996 | 0,118445902 | 3,972614956 | 7,11E-05 | 0,002639318 |
| Pycard | protein_coding | 1449,919024 | 0,472816705 | 0,119157329 | 3,968003548 | 7,25E-05 | 0,002672961 |
| Tceanc2 | protein_coding | 452,9056523 | 0,476716028 | 0,159800991 | 2,983185686 | 0,002852649 | 0,036150946 |
| Foxred1 | protein_coding | 633,3000486 | 0,477602304 | 0,168940147 | 2,827050364 | 0,004697894 | 0,049819147 |
| Fam162a | protein_coding | 597,9621436 | 0,477861664 | 0,124197013 | 3,84760996 | 1,19E-04 | 0,00390434 |
| Mrpl19 | protein_coding | 552,0713271 | 0,478304136 | 0,119331884 | 4,008183885 | 6,12E-05 | 0,002361552 |
| Lage3 | protein_coding | 377,4481816 | 0,478339958 | 0,169213494 | 2,826842859 | 0,004700939 | 0,049819147 |
| Pkn1 | protein_coding | 681,1572902 | 0,482445811 | 0,150586542 | 3,20377775 | 0,001356371 | 0,021520401 |
| Nob1 | protein_coding | 968,1639095 | 0,486293706 | 0,13960521 | 3,483349266 | 4,95E-04 | 0,010848985 |
| Rbm15 | protein_coding | 743,2439081 | 0,48694914 | 0,155523729 | 3,131027928 | 0,001741956 | 0,025678852 |
| Abhd11 | protein_coding | 630,7292244 | 0,48707436 | 0,163829476 | 2,973056936 | 0,002948498 | 0,036740901 |
| Fam114a2 | protein_coding | 1181,732777 | 0,487467487 | 0,164494871 | 2,96342059 | 0,003042406 | 0,037506102 |
| Nup93 | protein_coding | 691,7126166 | 0,488244599 | 0,156906571 | 3,111689938 | 0,001860198 | 0,026763653 |
| Kmt2c | protein_coding | 1230,146194 | 0,490175565 | 0,155065667 | 3,161083772 | 0,001571833 | 0,02395277 |
| Ctdsp2 | protein_coding | 512,6696976 | 0,492646117 | 0,138189311 | 3,56500885 | 3,64E-04 | 0,008898836 |
| Mknk2 | protein_coding | 1192,402303 | 0,49265186 | 0,123121442 | 4,001349019 | 6,30E-05 | 0,002416777 |
| Herc2 | protein_coding | 764,1078994 | 0,497099345 | 0,14124466 | 3,519420461 | 4,32E-04 | 0,009968911 |
| Ankrd11 | protein_coding | 895,7921988 | 0,497583169 | 0,162003871 | 3,071427642 | 0,002130378 | 0,029499575 |
| Noa1 | protein_coding | 607,293355 | 0,498067247 | 0,135135083 | 3,685699055 | 2,28E-04 | 0,006312851 |
| Ubr4 | protein_coding | 1732,224648 | 0,498485921 | 0,154610441 | 3,224141386 | 0,00126351 | 0,020497763 |
| Npepl1 | protein_coding | 747,0799902 | 0,499837075 | 0,12723853 | 3,92834683 | 8,55E-05 | 0,003020182 |
| Rnaseh1 | protein_coding | 398,2086885 | 0,500985892 | 0,169066695 | 2,963244124 | 0,00304415 | 0,037506102 |
| 3830406C13Rik | protein_coding | 477,9726477 | 0,501208387 | 0,155317701 | 3,226988187 | 0,001251006 | 0,02042979 |
| Mycbp2 | protein_coding | 2051,309363 | 0,502976627 | 0,135098997 | 3,723022667 | 1,97E-04 | 0,005671794 |
| Nfatc3 | protein_coding | 1072,018566 | 0,503964222 | 0,153493233 | 3,283299286 | 0,001025997 | 0,017885969 |

| | | | | | | | |
|---|---|---|---|---|---|---|---|
| Mprip | protein_coding | 483,7189198 | 0,508210403 | 0,178821962 | 2,841990977 | 0,004483276 | 0,048662164 |
| Nktr | protein_coding | 1225,958793 | 0,508668573 | 0,125314236 | 4,059144346 | 4,93E-05 | 0,002022904 |
| Aacs | protein_coding | 1335,609196 | 0,508851027 | 0,166121116 | 3,063132738 | 0,002190329 | 0,030091636 |
| Clcn3 | protein_coding | 700,1064282 | 0,512560035 | 0,128750426 | 3,981035628 | 6,86E-05 | 0,002564742 |
| Mtrr | protein_coding | 475,3717025 | 0,514092112 | 0,179952186 | 2,856826166 | 0,004279001 | 0,046998215 |
| Wdr75 | protein_coding | 809,0125499 | 0,515947058 | 0,165638583 | 3,114896591 | 0,001840094 | 0,026531886 |
| Chchd4 | protein_coding | 376,8882942 | 0,51686221 | 0,161666568 | 3,197087781 | 0,001388227 | 0,021898291 |
| Gnl3 | protein_coding | 1322,566262 | 0,516898845 | 0,146913715 | 3,518383868 | 4,34E-04 | 0,009980214 |
| Gstp3 | protein_coding | 1093,085115 | 0,520040656 | 0,127630685 | 4,074573901 | 4,61E-05 | 0,001917428 |
| Slc19a1 | protein_coding | 638,9119529 | 0,523721864 | 0,167287269 | 3,130673765 | 0,001744058 | 0,025682683 |
| Zdhhc18 | protein_coding | 782,305633 | 0,527482077 | 0,153132058 | 3,444622128 | 5,72E-04 | 0,01205916 |
| Atad3a | protein_coding | 1103,27466 | 0,528551219 | 0,125033545 | 4,227275315 | 2,37E-05 | 0,001154585 |
| Srebf1 | protein_coding | 2569,172647 | 0,529204488 | 0,155100236 | 3,412016013 | 6,45E-04 | 0,013098921 |
| Mettl14 | protein_coding | 454,2167832 | 0,529229233 | 0,153443929 | 3,449007304 | 5,63E-04 | 0,011910413 |
| Alkbh5 | protein_coding | 1864,633749 | 0,532952215 | 0,180497496 | 2,952684813 | 0,003150234 | 0,038402849 |
| Emb | protein_coding | 3579,828897 | 0,533992654 | 0,153310239 | 3,4830854 | 4,96E-04 | 0,010848985 |
| Rgp1 | protein_coding | 507,2601652 | 0,535357581 | 0,149837221 | 3,572927852 | 3,53E-04 | 0,008782356 |
| Bach2 | protein_coding | 684,4772934 | 0,537078904 | 0,179413197 | 2,993530657 | 0,002757698 | 0,035616153 |
| Lnpep | protein_coding | 1545,483822 | 0,537641104 | 0,147081636 | 3,65539247 | 2,57E-04 | 0,006840558 |
| Fastkd2 | protein_coding | 352,865323 | 0,538160775 | 0,160601131 | 3,35091522 | 8,05E-04 | 0,015323468 |
| Epn1 | protein_coding | 334,7765711 | 0,539510225 | 0,146605191 | 3,680021309 | 2,33E-04 | 0,006407992 |
| Etf1 | protein_coding | 1750,133478 | 0,543963212 | 0,138449276 | 3,928971158 | 8,53E-05 | 0,003018776 |
| Pola2 | protein_coding | 687,2575563 | 0,545990463 | 0,193110809 | 2,827342839 | 0,004693605 | 0,049819147 |
| Pno1 | protein_coding | 475,3116023 | 0,547671546 | 0,159207719 | 3,439981101 | 5,82E-04 | 0,012252288 |
| Srm | protein_coding | 1204,678694 | 0,549289595 | 0,134118904 | 4,095541925 | 4,21E-05 | 0,001820296 |
| Spen | protein_coding | 346,1972614 | 0,555061929 | 0,179562237 | 3,091195216 | 0,001993525 | 0,028109214 |
| Hnrnpd | protein_coding | 338,5973407 | 0,555291936 | 0,180641992 | 3,07399144 | 0,002112155 | 0,029284313 |
| Usp18 | protein_coding | 2317,745702 | 0,555960059 | 0,166238382 | 3,344354377 | 8,25E-04 | 0,015589336 |
| Ybx1 | protein_coding | 3328,357542 | 0,558122242 | 0,193265282 | 2,887855686 | 0,003878778 | 0,043880162 |
| Cyp4f17 | protein_coding | 259,1033604 | 0,562540013 | 0,169860303 | 3,311780341 | 9,27E-04 | 0,01677776 |
| Irak1 | protein_coding | 356,4811751 | 0,562631467 | 0,175476872 | 3,206299832 | 0,001344538 | 0,021394012 |
| Sigmar1 | protein_coding | 665,7144634 | 0,563060726 | 0,174552558 | 3,225737463 | 0,001256486 | 0,020483783 |
| Desi2 | protein_coding | 492,4728251 | 0,565431097 | 0,194109146 | 2,912954437 | 0,003580268 | 0,041784904 |
| Tnfsf11 | protein_coding | 759,3902246 | 0,566258898 | 0,191786601 | 2,952546714 | 0,003151644 | 0,038402849 |
| Psmg3 | protein_coding | 206,6479092 | 0,567428354 | 0,18881016 | 3,005285062 | 0,002653321 | 0,034578313 |
| Cul3 | protein_coding | 449,257956 | 0,567908104 | 0,160623368 | 3,535650573 | 4,07E-04 | 0,009521571 |
| Rnf213 | protein_coding | 1346,95577 | 0,569617441 | 0,192252065 | 2,962867745 | 0,003047875 | 0,037524134 |
| CAAA01118383.1 | protein_coding | 372,9738543 | 0,569787051 | 0,180286057 | 3,160460995 | 0,001575197 | 0,023961474 |
| Gm17018 | protein_coding | 362,7432504 | 0,573246927 | 0,162993809 | 3,516985903 | 4,36E-04 | 0,010005209 |
| Ric1 | protein_coding | 370,6015196 | 0,573407184 | 0,169976849 | 3,373442847 | 7,42E-04 | 0,014443084 |
| Trim28 | protein_coding | 837,1148338 | 0,574440024 | 0,167291648 | 3,433763921 | 5,95E-04 | 0,012421981 |
| Ankib1 | protein_coding | 422,9674086 | 0,575533477 | 0,167778594 | 3,430315294 | 6,03E-04 | 0,012522404 |
| Atp2b1 | protein_coding | 2540,129765 | 0,58114619 | 0,149674893 | 3,882723265 | 1,03E-04 | 0,003470146 |
| Bicdl1 | protein_coding | 306,04596 | 0,581199127 | 0,168051876 | 3,458450699 | 5,43E-04 | 0,011634154 |
| Acot7 | protein_coding | 1259,801686 | 0,584241466 | 0,161503578 | 3,617514073 | 2,97E-04 | 0,007606189 |
| Rictor | protein_coding | 645,8507434 | 0,585353451 | 0,206270939 | 2,837789232 | 0,004542717 | 0,048955149 |
| Snord78 | snoRNA | 180,0382606 | 0,585402241 | 0,174383454 | 3,356982722 | 7,88E-04 | 0,01512453 |
| Kpnb1 | protein_coding | 785,5238264 | 0,588143178 | 0,132880737 | 4,426098101 | 9,60E-06 | 5,67E-04 |
| Arf6 | protein_coding | 400,5192539 | 0,589626564 | 0,191624355 | 3,076991776 | 0,002091011 | 0,029039677 |
| Gpr137 | protein_coding | 244,7680676 | 0,592474449 | 0,208057783 | 2,847643774 | 0,004404419 | 0,047963087 |
| Hmgcr | protein_coding | 2029,432219 | 0,594152373 | 0,126373136 | 4,701571816 | 2,58E-06 | 1,94E-04 |
| Cep83 | protein_coding | 240,1719209 | 0,594264657 | 0,2054394 | 2,892651837 | 0,003820045 | 0,043333879 |
| Prrc2a | protein_coding | 1608,395987 | 0,595727592 | 0,114721742 | 5,192804609 | 2,07E-07 | 2,29E-05 |
| Nudt3 | protein_coding | 306,1946874 | 0,596474116 | 0,169618758 | 3,516557503 | 4,37E-04 | 0,010007547 |
| Hdac7 | protein_coding | 1905,970669 | 0,598584575 | 0,15346579 | 3,900443058 | 9,60E-05 | 0,003265359 |
| Crebl2 | protein_coding | 202,1997093 | 0,602511827 | 0,202284334 | 2,978539237 | 0,002896259 | 0,03641388 |
| Arl4c | protein_coding | 859,3663858 | 0,603844808 | 0,163541915 | 3,692293842 | 2,22E-04 | 0,006209264 |
| Birc6 | protein_coding | 2361,096939 | 0,606976289 | 0,108642507 | 5,586913503 | 2,31E-08 | 3,29E-06 |
| Rnf126 | protein_coding | 295,435958 | 0,607615861 | 0,203747803 | 2,982195886 | 0,002861888 | 0,036150946 |
| Stk26 | protein_coding | 791,0792195 | 0,610460102 | 0,18711043 | 3,262565861 | 0,001104085 | 0,018870646 |
| Fam120a | protein_coding | 812,2984406 | 0,611043329 | 0,158777651 | 3,848421525 | 1,19E-04 | 0,003901523 |
| Ccdc6 | protein_coding | 494,9436644 | 0,611340925 | 0,209443283 | 2,918885329 | 0,003512854 | 0,041171842 |
| Myc | protein_coding | 4741,605711 | 0,611715067 | 0,160580177 | 3,809405854 | 1,39E-04 | 0,004455006 |
| Alas1 | protein_coding | 420,3855029 | 0,612617056 | 0,194324737 | 3,152542822 | 0,001618551 | 0,024486298 |
| Shmt1 | protein_coding | 338,2567923 | 0,614663876 | 0,199698916 | 3,077952988 | 0,002084278 | 0,028980096 |
| Lap3 | protein_coding | 530,1232343 | 0,615323274 | 0,163045029 | 3,773946858 | 1,61E-04 | 0,004975239 |
| Als2cl | protein_coding | 1158,852322 | 0,617277979 | 0,168632249 | 3,660497814 | 2,52E-04 | 0,006781883 |
| Wdr26 | protein_coding | 897,8991362 | 0,617445887 | 0,163810204 | 3,769276092 | 1,64E-04 | 0,005041049 |
| Gm11131 | antisense | 288,2970436 | 0,619722159 | 0,206706142 | 2,998082947 | 0,002716837 | 0,035170538 |

| | | | | | | | |
|---|---|---|---|---|---|---|---|
| 2810004N23Rik | protein_coding | 480,4801639 | 0,621692755 | 0,144468488 | 4,303310453 | 1,68E-05 | 8,70E-04 |
| Igfbp4 | protein_coding | 3706,253023 | 0,623551674 | 0,208399385 | 2,992099403 | 0,00277066 | 0,035737251 |
| Dusp6 | protein_coding | 769,0067864 | 0,624857682 | 0,199571085 | 3,131003081 | 0,001742103 | 0,025678852 |
| Psme3 | protein_coding | 1067,434339 | 0,626590907 | 0,220904384 | 2,836480182 | 0,004561381 | 0,049063856 |
| Kcnq1ot1 | antisense | 883,2028117 | 0,627680895 | 0,167609992 | 3,74488947 | 1,80E-04 | 0,005352084 |
| Tjap1 | protein_coding | 565,1994808 | 0,631006688 | 0,184429832 | 3,421391655 | 6,23E-04 | 0,012764894 |
| Rc3h1 | protein_coding | 712,4856819 | 0,631145772 | 0,135142028 | 4,670240482 | 3,01E-06 | 2,20E-04 |
| Cdkn1b | protein_coding | 1172,105578 | 0,631299743 | 0,154345902 | 4,090161999 | 4,31E-05 | 0,001833232 |
| Taf1d | protein_coding | 516,0885177 | 0,631588339 | 0,190456938 | 3,316173967 | 9,13E-04 | 0,016679894 |
| Usp34 | protein_coding | 732,0378456 | 0,63168139 | 0,177864238 | 3,551480603 | 3,83E-04 | 0,009238018 |
| Ppp3cc | protein_coding | 240,0463513 | 0,632372523 | 0,219439593 | 2,881761283 | 0,003954592 | 0,044495194 |
| Smn1 | protein_coding | 541,0432564 | 0,636181497 | 0,170814062 | 3,724409396 | 1,96E-04 | 0,00565052 |
| Rab3a | protein_coding | 204,9705488 | 0,638279232 | 0,219022114 | 2,914222769 | 0,003565754 | 0,041671401 |
| Gm34220 | TEC | 175,6677182 | 0,640936378 | 0,206859465 | 3,098414562 | 0,001945591 | 0,027573887 |
| Msmo1 | protein_coding | 1471,480027 | 0,640957014 | 0,108521085 | 5,906290123 | 3,50E-09 | 6,24E-07 |
| Dph5 | protein_coding | 1007,806571 | 0,641910384 | 0,136484768 | 4,703165004 | 2,56E-06 | 1,93E-04 |
| Ubxn11 | protein_coding | 484,0986408 | 0,644682777 | 0,172601468 | 3,7350944 | 1,88E-04 | 0,005511775 |
| D6Wsu163e | protein_coding | 175,3344629 | 0,644921009 | 0,211144757 | 3,054402193 | 0,002255095 | 0,030598687 |
| I730030J21Rik | processed_transcript | 676,5179739 | 0,645973194 | 0,199833293 | 3,232560416 | 0,001226862 | 0,020179386 |
| Themis | protein_coding | 1088,439752 | 0,647503724 | 0,148561116 | 4,358500666 | 1,31E-05 | 7,24E-04 |
| Hnrnpl | protein_coding | 505,4248179 | 0,648888209 | 0,201408584 | 3,221750516 | 0,0012741 | 0,020589063 |
| Fmc1 | protein_coding | 148,3536238 | 0,650556698 | 0,216555047 | 3,004116998 | 0,002663529 | 0,034615445 |
| Tnfrsf14 | protein_coding | 181,9547399 | 0,652557669 | 0,221543588 | 2,945504646 | 0,003224283 | 0,038960231 |
| A430093F15Rik | processed_transcript | 707,2496443 | 0,65441542 | 0,175657874 | 3,725511442 | 1,95E-04 | 0,00563569 |
| Ece1 | protein_coding | 803,9130819 | 0,655410032 | 0,173020004 | 3,788059272 | 1,52E-04 | 0,004745288 |
| Ring1 | protein_coding | 561,1344619 | 0,655662703 | 0,19527979 | 3,357555348 | 7,86E-04 | 0,015121975 |
| Qtrt1 | protein_coding | 326,6395679 | 0,656507875 | 0,213300565 | 3,077853432 | 0,002084974 | 0,028980096 |
| Snhg5 | processed_transcript | 638,3894934 | 0,65784934 | 0,189313506 | 3,474920279 | 5,11E-04 | 0,011071334 |
| Hnrnpa0 | protein_coding | 169,3294675 | 0,658880851 | 0,219466565 | 3,002192383 | 0,002680427 | 0,034807801 |
| Cisd1 | protein_coding | 328,3933758 | 0,66158569 | 0,210692304 | 3,140056263 | 0,001689154 | 0,025232402 |
| Rflnb | protein_coding | 650,3784476 | 0,66413435 | 0,196453256 | 3,380622774 | 7,23E-04 | 0,014237863 |
| Atp5s | protein_coding | 179,6795443 | 0,664451349 | 0,223174061 | 2,977278572 | 0,002908196 | 0,036435303 |
| Dus2 | protein_coding | 383,8006615 | 0,667218987 | 0,232091333 | 2,87481216 | 0,004042679 | 0,045302029 |
| Daglb | protein_coding | 362,8378745 | 0,668208345 | 0,189977319 | 3,517305895 | 4,36E-04 | 0,010005209 |
| Mid1ip1 | protein_coding | 354,684545 | 0,672682646 | 0,159537038 | 4,216466932 | 2,48E-05 | 0,001197226 |
| Phf12 | protein_coding | 346,877969 | 0,674233971 | 0,179556109 | 3,755004355 | 1,73E-04 | 0,005220917 |
| Ptpn9 | protein_coding | 232,9408889 | 0,676873626 | 0,228478148 | 2,962531126 | 0,00305121 | 0,037537342 |
| Sympk | protein_coding | 336,2370628 | 0,678133862 | 0,203569913 | 3,331208679 | 8,65E-04 | 0,016070012 |
| Adgrl1 | protein_coding | 207,2417767 | 0,678525647 | 0,183721487 | 3,693229665 | 2,21E-04 | 0,006202878 |
| 2700038G22Rik | lincRNA | 118,9104165 | 0,680037321 | 0,211317065 | 3,218089934 | 0,001290473 | 0,020792911 |
| 5430405H02Rik | processed_transcript | 126,1846099 | 0,680044339 | 0,240348454 | 2,829410077 | 0,00466339 | 0,049579517 |
| Usp7 | protein_coding | 627,3523766 | 0,680225424 | 0,207080678 | 3,284832903 | 0,001020429 | 0,017807607 |
| Trim56 | protein_coding | 545,6493849 | 0,680417839 | 0,187473366 | 3,629410692 | 2,84E-04 | 0,007377792 |
| Cox10 | protein_coding | 517,9235685 | 0,68106335 | 0,184074328 | 3,699936625 | 2,16E-04 | 0,006107479 |
| Capn10 | protein_coding | 234,1906363 | 0,684951958 | 0,209033965 | 3,276749581 | 0,001050095 | 0,01813462 |
| Eif4g3 | protein_coding | 1155,300304 | 0,687361416 | 0,211181948 | 3,254830361 | 0,001134601 | 0,019228852 |
| Nsdhl | protein_coding | 510,2560577 | 0,690336664 | 0,158890786 | 4,344724323 | 1,39E-05 | 7,56E-04 |
| Prim1 | protein_coding | 251,4546627 | 0,691433842 | 0,228851507 | 3,021320907 | 0,002516745 | 0,033149123 |
| Cpm | protein_coding | 984,7860252 | 0,691933111 | 0,16745476 | 4,132059964 | 3,60E-05 | 0,001603953 |
| Usf3 | protein_coding | 268,2296603 | 0,695850974 | 0,23763719 | 2,92820739 | 0,003409226 | 0,040403577 |
| Mettl1 | protein_coding | 216,2497706 | 0,700985832 | 0,187985738 | 3,728930936 | 1,92E-04 | 0,005583364 |
| Cox16 | protein_coding | 292,7583915 | 0,701280759 | 0,206854663 | 3,390210049 | 6,98E-04 | 0,013897476 |
| Dusp5 | protein_coding | 1261,249805 | 0,703893706 | 0,228653867 | 3,078424672 | 0,002080981 | 0,028973698 |
| Sfxn1 | protein_coding | 1064,51473 | 0,704560844 | 0,148337631 | 4,74971078 | 2,04E-06 | 1,58E-04 |
| Tmem104 | protein_coding | 404,1274288 | 0,704617112 | 0,235517463 | 2,991782869 | 0,002773535 | 0,035737251 |
| Tnfrsf22 | protein_coding | 471,5247951 | 0,705137718 | 0,173595321 | 4,061962705 | 4,87E-05 | 0,002003951 |
| Gcsh | protein_coding | 322,8591629 | 0,705155317 | 0,160169703 | 4,402551206 | 1,07E-05 | 6,19E-04 |
| Acsf3 | protein_coding | 307,6347791 | 0,706009067 | 0,21733945 | 3,24841656 | 0,001160492 | 0,01944875 |
| Tmem256 | protein_coding | 321,771398 | 0,706131863 | 0,218283931 | 3,234923699 | 0,001216752 | 0,020060149 |
| Tpi1 | protein_coding | 1528,053558 | 0,712497194 | 0,186477991 | 3,820811185 | 1,33E-04 | 0,004269809 |
| B230354K17Rik | bidirectional_promoter | 190,7757011 | 0,712558381 | 0,252058992 | 2,826950852 | 0,004699354 | 0,049819147 |
| Gm23187 | snoRNA | 107,5583503 | 0,712734594 | 0,250084169 | 2,849978858 | 0,004372214 | 0,047706283 |
| D17H6S53E | protein_coding | 343,5633826 | 0,712902572 | 0,20431611 | 3,489213715 | 4,84E-04 | 0,010719769 |
| Rock2 | protein_coding | 281,8501632 | 0,713963117 | 0,180898157 | 3,946768328 | 7,92E-05 | 0,002864099 |
| Cmss1 | protein_coding | 149,7207823 | 0,714039031 | 0,244458083 | 2,920905791 | 0,003490153 | 0,04096364 |
| Sart1 | protein_coding | 479,4548966 | 0,718255219 | 0,177505534 | 4,046382109 | 5,20E-05 | 0,002110626 |
| G6pc3 | protein_coding | 293,2767528 | 0,719123257 | 0,217744911 | 3,302595017 | 9,58E-04 | 0,017076344 |
| Setd5 | protein_coding | 407,9530165 | 0,722169915 | 0,17976955 | 4,017198218 | 5,89E-05 | 0,002310657 |
| Stard4 | protein_coding | 370,4209515 | 0,724401084 | 0,150987219 | 4,797764279 | 1,60E-06 | 1,27E-04 |

| | | | | | | | |
|---|---|---|---|---|---|---|---|
| Rcc2 | protein_coding | 518,4595126 | 0,726956126 | 0,244815442 | 2,969404701 | 0,002983773 | 0,037064898 |
| Tfdp1 | protein_coding | 576,1930147 | 0,728239043 | 0,205964519 | 3,53574997 | 4,07E-04 | 0,009521571 |
| Fdft1 | protein_coding | 1198,615738 | 0,728380817 | 0,118492491 | 6,147063061 | 7,89E-10 | 1,87E-07 |
| Fads1 | protein_coding | 379,7678912 | 0,729549613 | 0,181114941 | 4,028102863 | 5,62E-05 | 0,002243267 |
| Timm9 | protein_coding | 315,1612864 | 0,732052925 | 0,16042301 | 4,563266351 | 5,04E-06 | 3,34E-04 |
| 5830444F18Rik | TEC | 142,0221935 | 0,733486207 | 0,254560103 | 2,881387141 | 0,00395929 | 0,04451787 |
| Tfap4 | protein_coding | 225,7397557 | 0,73359275 | 0,20629547 | 3,556029362 | 3,77E-04 | 0,009127172 |
| Arhgef10 | protein_coding | 375,5937758 | 0,737104468 | 0,238706565 | 3,087910331 | 0,002015693 | 0,028301555 |
| Ppp4r1 | protein_coding | 418,979909 | 0,739990462 | 0,213884767 | 3,459762341 | 5,41E-04 | 0,011592592 |
| Thy1 | protein_coding | 1294,404133 | 0,740038818 | 0,115101616 | 6,429438962 | 1,28E-10 | 3,43E-08 |
| Jak2 | protein_coding | 669,9927636 | 0,741709424 | 0,161585348 | 4,590202238 | 4,43E-06 | 3,04E-04 |
| Rpp38 | protein_coding | 179,1895273 | 0,742569437 | 0,225369396 | 3,29489918 | 9,85E-04 | 0,017399228 |
| Igsf23 | protein_coding | 476,8888686 | 0,744539899 | 0,181256716 | 4,107654141 | 4,00E-05 | 0,001743911 |
| 1110038B12Rik | processed_transcript | 157,8101552 | 0,746920508 | 0,245362291 | 3,044153633 | 0,002333359 | 0,03143217 |
| Clock | protein_coding | 313,0438372 | 0,751461726 | 0,239378881 | 3,1392148 | 0,001694012 | 0,025264963 |
| Gar1 | protein_coding | 291,6195665 | 0,75323717 | 0,198722078 | 3,790405072 | 1,50E-04 | 0,004727402 |
| Rbm26 | protein_coding | 406,4467158 | 0,753766413 | 0,236056234 | 3,193164607 | 0,001407227 | 0,022081672 |
| Rab21 | protein_coding | 305,3047797 | 0,754563714 | 0,208221019 | 3,623859477 | 2,90E-04 | 0,007485832 |
| Urb1 | protein_coding | 272,6590102 | 0,755910465 | 0,250393315 | 3,018892353 | 0,002537007 | 0,033336628 |
| Mrpl27 | protein_coding | 361,0182761 | 0,760356236 | 0,175522611 | 4,331956051 | 1,48E-05 | 7,91E-04 |
| Art2a-ps | polymorphic_pseudo | 353,1084003 | 0,760389295 | 0,157007607 | 4,84300926 | 1,28E-06 | 1,06E-04 |
| Ydjc | protein_coding | 180,3215399 | 0,761071012 | 0,235857201 | 3,226829662 | 0,0012517 | 0,02042979 |
| 1190007I07Rik | protein_coding | 211,6096595 | 0,764317963 | 0,208325755 | 3,66885969 | 2,44E-04 | 0,006606801 |
| Acsl3 | protein_coding | 354,7933204 | 0,766298066 | 0,216814776 | 3,534344296 | 4,09E-04 | 0,00955528 |
| Acvrl1 | protein_coding | 217,380544 | 0,767614998 | 0,250005167 | 3,070396528 | 0,002137747 | 0,029565047 |
| Cd40lg | protein_coding | 586,9177013 | 0,767971185 | 0,196861571 | 3,901072112 | 9,58E-05 | 0,003265359 |
| Dpp7 | protein_coding | 349,8199187 | 0,77417282 | 0,208279696 | 3,716986516 | 2,02E-04 | 0,00574834 |
| Abhd2 | protein_coding | 878,1853231 | 0,777844273 | 0,185488895 | 4,193481634 | 2,75E-05 | 0,001306306 |
| Snhg6 | lincRNA | 300,9481503 | 0,779100783 | 0,178390936 | 4,367378749 | 1,26E-05 | 7,03E-04 |
| Tmem38b | protein_coding | 191,1582684 | 0,784896201 | 0,255834775 | 3,067980892 | 0,002155104 | 0,029730761 |
| Fbrs | protein_coding | 304,9921676 | 0,786727334 | 0,216426142 | 3,635084598 | 2,78E-04 | 0,007262761 |
| Fbxw17 | protein_coding | 250,41213 | 0,78827506 | 0,266461888 | 2,958303214 | 0,003093377 | 0,037859647 |
| Slc39a4 | protein_coding | 520,1546183 | 0,796104936 | 0,229084738 | 3,475154835 | 5,11E-04 | 0,011071334 |
| Dusp10 | protein_coding | 941,4745946 | 0,802274593 | 0,172062427 | 4,662694864 | 3,12E-06 | 2,27E-04 |
| Zfas1 | processed_transcript | 391,6007768 | 0,803105059 | 0,14895666 | 5,391535093 | 6,99E-08 | 8,65E-06 |
| Snhg12 | lincRNA | 117,9755196 | 0,805690869 | 0,270124403 | 2,982665982 | 0,002857497 | 0,036150946 |
| Arap2 | protein_coding | 996,4695323 | 0,808846487 | 0,163546797 | 4,945657765 | 7,59E-07 | 6,84E-05 |
| Mvd | protein_coding | 901,8098552 | 0,809613441 | 0,165818616 | 4,882524413 | 1,05E-06 | 9,10E-05 |
| Zcchc2 | protein_coding | 265,4535553 | 0,815591597 | 0,281347565 | 2,898875618 | 0,003745034 | 0,042834314 |
| Gpt | protein_coding | 152,6049322 | 0,817578057 | 0,252460212 | 3,238443206 | 0,001201839 | 0,019925802 |
| Wtap | protein_coding | 2039,036942 | 0,822815156 | 0,150555083 | 5,46521006 | 4,62E-08 | 6,04E-06 |
| Tmem238 | protein_coding | 204,5934641 | 0,827967365 | 0,28659291 | 2,889001566 | 0,003864671 | 0,043750401 |
| Cited2 | protein_coding | 218,9161519 | 0,828792589 | 0,215851896 | 3,839635429 | 1,23E-04 | 0,004001784 |
| Zfc3h1 | protein_coding | 700,5984958 | 0,831665546 | 0,20043836 | 4,149233446 | 3,34E-05 | 0,001525143 |
| Pfas | protein_coding | 513,2718501 | 0,832531835 | 0,169158132 | 4,921618758 | 8,58E-07 | 7,66E-05 |
| Mfhas1 | protein_coding | 250,3130528 | 0,832912924 | 0,200352967 | 4,157227791 | 3,22E-05 | 0,001489169 |
| Cyp51 | protein_coding | 1792,742796 | 0,833352431 | 0,167039895 | 4,988942498 | 6,07E-07 | 5,69E-05 |
| Cd3eap | protein_coding | 233,6671264 | 0,834971413 | 0,204968326 | 4,073660688 | 4,63E-05 | 0,001920153 |
| Chd7 | protein_coding | 630,0645054 | 0,835849016 | 0,240853146 | 3,470367855 | 5,20E-04 | 0,011216781 |
| Ccdc28b | protein_coding | 185,8317552 | 0,841769906 | 0,277734818 | 3,030840398 | 0,002438741 | 0,032430568 |
| Ppp1r14b | protein_coding | 122,9139982 | 0,844526191 | 0,257646477 | 3,277848779 | 0,001046014 | 0,01808297 |
| Rasa1 | protein_coding | 759,1383427 | 0,846716999 | 0,195955459 | 4,320966624 | 1,55E-05 | 8,16E-04 |
| Hmgcs1 | protein_coding | 1281,789717 | 0,850210298 | 0,191517395 | 4,439337218 | 9,02E-06 | 5,45E-04 |
| Ifrd2 | protein_coding | 404,7067847 | 0,850495549 | 0,174947451 | 4,861434357 | 1,17E-06 | 1,00E-04 |
| Bmp2k | protein_coding | 330,7306934 | 0,851947943 | 0,235293555 | 3,620787416 | 2,94E-04 | 0,007545471 |
| Iffo2 | protein_coding | 79,36030431 | 0,852900864 | 0,300924912 | 2,834264732 | 0,004593126 | 0,049350061 |
| Ung | protein_coding | 261,6530537 | 0,858074478 | 0,183490742 | 4,676391118 | 2,92E-06 | 2,15E-04 |
| Ccnk | protein_coding | 171,729397 | 0,860549039 | 0,287057661 | 2,997826419 | 0,002719125 | 0,035172719 |
| Utp20 | protein_coding | 264,2008957 | 0,865382784 | 0,185188303 | 4,672988347 | 2,97E-06 | 2,18E-04 |
| Qtrt2 | protein_coding | 193,6191812 | 0,865410666 | 0,251612012 | 3,439464835 | 5,83E-04 | 0,012260121 |
| Dhcr24 | protein_coding | 87,37391843 | 0,865486589 | 0,297498304 | 2,909215202 | 0,003623373 | 0,04208083 |
| Gm3571 | processed_pseudoge | 142,9810726 | 0,867747892 | 0,203109906 | 4,272307091 | 1,93E-05 | 9,67E-04 |
| Hdhd2 | protein_coding | 76,59297312 | 0,868800107 | 0,304966221 | 2,848840455 | 0,004387888 | 0,047814436 |
| Mboat1 | protein_coding | 222,5444418 | 0,870056101 | 0,284015497 | 3,063410662 | 0,002188295 | 0,030088608 |
| 2900005J15Rik | processed_transcript | 281,7205494 | 0,870260938 | 0,271917014 | 3,200465187 | 0,001372059 | 0,021707053 |
| Suds3 | protein_coding | 215,3203432 | 0,890044601 | 0,307533845 | 2,894135443 | 0,003802041 | 0,043218271 |
| Snhg15 | processed_transcript | 234,3045694 | 0,891625093 | 0,169525376 | 5,259537612 | 1,44E-07 | 1,68E-05 |
| Gsk3a | protein_coding | 64,8180172 | 0,894295464 | 0,312136789 | 2,865075497 | 0,004169098 | 0,046246379 |
| 9330162G02Rik | TEC | 205,1907564 | 0,896080535 | 0,297473603 | 3,01230269 | 0,002592739 | 0,033934622 |

| | | | | | | | |
|---|---|---|---|---|---|---|---|
| Pole2 | protein_coding | 71,06291652 | 0,901367955 | 0,31408718 | 2,869801796 | 0,004107292 | 0,045840358 |
| Rec8 | protein_coding | 335,7493688 | 0,905834912 | 0,304076477 | 2,978970692 | 0,002892184 | 0,03641388 |
| Zfp691 | protein_coding | 110,4138946 | 0,906241248 | 0,289567854 | 3,129633468 | 0,001750245 | 0,025728143 |
| Pan3 | protein_coding | 1007,231118 | 0,910570559 | 0,168705421 | 5,397399529 | 6,76E-08 | 8,44E-06 |
| Extl2 | protein_coding | 132,9011091 | 0,910970188 | 0,320863654 | 2,839119288 | 0,004523824 | 0,048905778 |
| Uros | protein_coding | 588,1986199 | 0,911046493 | 0,200162979 | 4,551523455 | 5,33E-06 | 3,49E-04 |
| Acaca | protein_coding | 240,075092 | 0,913215008 | 0,285708477 | 3,196317506 | 0,001391938 | 0,021898291 |
| BC024978 | protein_coding | 94,35884391 | 0,913482225 | 0,293123024 | 3,116378274 | 0,001830873 | 0,026452754 |
| Egr3 | protein_coding | 114,6742936 | 0,915494442 | 0,278215385 | 3,290596028 | 1,00E-03 | 0,01753902 |
| Pag1 | protein_coding | 528,4261902 | 0,917763568 | 0,228602983 | 4,01466138 | 5,95E-05 | 0,002324658 |
| Plekha5 | protein_coding | 193,2615843 | 0,918333031 | 0,32118544 | 2,85919882 | 0,004247125 | 0,046802979 |
| Fdps | protein_coding | 128,1040338 | 0,927313403 | 0,218337678 | 4,247152444 | 2,17E-05 | 0,001066456 |
| 1810032O08Rik | processed_transcript | 193,0253764 | 0,928437558 | 0,2378645 | 3,903220349 | 9,49E-05 | 0,003254778 |
| Ldlr | protein_coding | 1833,63119 | 0,937175518 | 0,182248212 | 5,142302957 | 2,71E-07 | 2,82E-05 |
| Gm26637 | lincRNA | 113,2730319 | 0,938567114 | 0,322501036 | 2,91027628 | 0,003611094 | 0,042026447 |
| Fam102b | protein_coding | 118,1958558 | 0,940293835 | 0,284002596 | 3,310863523 | 9,30E-04 | 0,016796192 |
| Insig1 | protein_coding | 1153,804771 | 0,941782774 | 0,155936421 | 6,039530512 | 1,55E-09 | 3,09E-07 |
| Car12 | protein_coding | 199,1092849 | 0,944561294 | 0,215656203 | 4,379940292 | 1,19E-05 | 6,68E-04 |
| Ptger2 | protein_coding | 1343,888073 | 0,94644055 | 0,16130533 | 5,867385458 | 4,43E-09 | 7,82E-07 |
| Scpep1 | protein_coding | 308,1774153 | 0,94698347 | 0,270516592 | 3,500648386 | 4,64E-04 | 0,010423091 |
| Snhg1 | processed_transcript | 343,5082523 | 0,947496417 | 0,183701468 | 5,157805369 | 2,50E-07 | 2,62E-05 |
| BC048403 | protein_coding | 138,5490161 | 0,951086536 | 0,320211528 | 2,970182058 | 0,002976233 | 0,036998925 |
| 2310022B05Rik | protein_coding | 167,2322996 | 0,953789639 | 0,280939081 | 3,395005195 | 6,86E-04 | 0,013738705 |
| Cnpy4 | protein_coding | 417,7681051 | 0,964872252 | 0,284348883 | 3,393269002 | 6,91E-04 | 0,013776232 |
| Rasgrp4 | protein_coding | 374,2668629 | 0,966056288 | 0,151820881 | 6,363131901 | 1,98E-10 | 5,21E-08 |
| Tent5a | protein_coding | 556,3001342 | 0,967293878 | 0,165822099 | 5,833323079 | 5,43E-09 | 9,30E-07 |
| Atg9b | protein_coding | 169,9039797 | 0,974482442 | 0,284222874 | 3,428585565 | 6,07E-04 | 0,012555334 |
| Zfp365 | protein_coding | 153,8044146 | 0,9754048 | 0,305671255 | 3,191025601 | 0,001417687 | 0,022217126 |
| Ube2q1 | protein_coding | 209,9989549 | 0,976922492 | 0,277004584 | 3,52673764 | 4,21E-04 | 0,009724457 |
| Pygm | protein_coding | 118,9745029 | 0,977836416 | 0,274262909 | 3,565325041 | 3,63E-04 | 0,008898836 |
| Csnk1e | protein_coding | 107,8033485 | 0,980281725 | 0,294601389 | 3,327485071 | 8,76E-04 | 0,016213697 |
| Oas2 | protein_coding | 255,2870063 | 0,984813059 | 0,302279127 | 3,257959188 | 0,001122165 | 0,019061881 |
| AC121821.1 | processed_transcript | 83,19894988 | 0,987825126 | 0,294850153 | 3,350261535 | 8,07E-04 | 0,015330468 |
| Mvb12b | protein_coding | 79,59002556 | 0,988333618 | 0,297743674 | 3,319410971 | 9,02E-04 | 0,016582528 |
| Mtln | protein_coding | 194,4613342 | 0,989504 | 0,226067979 | 4,377019718 | 1,20E-05 | 6,75E-04 |
| Mccc2 | protein_coding | 198,2920133 | 0,993846314 | 0,254330819 | 3,907691245 | 9,32E-05 | 0,003215079 |
| Ssh3 | protein_coding | 232,9879437 | 0,99453801 | 0,278884349 | 3,566130591 | 3,62E-04 | 0,008898836 |
| Abcb9 | protein_coding | 862,2994213 | 0,995041808 | 0,148398654 | 6,705194277 | 2,01E-11 | 6,30E-09 |
| Cmtr2 | protein_coding | 198,0962731 | 0,996966497 | 0,303962564 | 3,279898959 | 0,001038443 | 0,018008354 |
| Spaca1 | protein_coding | 63,59962429 | 1,007658588 | 0,338285688 | 2,97872072 | 0,002894545 | 0,03641388 |
| Kdm8 | protein_coding | 155,6306654 | 1,009966646 | 0,351148103 | 2,876184256 | 0,004025147 | 0,045166555 |
| B630019A10Rik | antisense | 466,8856192 | 1,014512156 | 0,139863134 | 7,253606625 | 4,06E-13 | 1,77E-10 |
| Pcsk4 | protein_coding | 418,7493132 | 1,017294594 | 0,197267209 | 5,156937134 | 2,51E-07 | 2,62E-05 |
| Xrcc3 | protein_coding | 102,1725821 | 1,017332363 | 0,322485388 | 3,154661894 | 0,001606842 | 0,024398555 |
| Tm7sf2 | protein_coding | 210,8695289 | 1,020416268 | 0,255035491 | 4,001075551 | 6,31E-05 | 0,002416777 |
| Fasl | protein_coding | 153,0025968 | 1,021278284 | 0,28955377 | 3,527076453 | 4,20E-04 | 0,009724457 |
| Tbl1x | protein_coding | 1896,325856 | 1,023856956 | 0,103362286 | 9,905517763 | 3,94E-23 | 7,26E-20 |
| Abca1 | protein_coding | 346,0291353 | 1,024676164 | 0,29839984 | 3,433903199 | 5,95E-04 | 0,012421981 |
| Gm42047 | lincRNA | 81,07184858 | 1,028490435 | 0,363775536 | 2,827266635 | 0,004694722 | 0,049819147 |
| Hsd17b7 | protein_coding | 214,1566953 | 1,030786038 | 0,288759202 | 3,569708021 | 3,57E-04 | 0,008852339 |
| Gm35363 | lincRNA | 190,4870309 | 1,031465362 | 0,242601731 | 4,251681796 | 2,12E-05 | 0,001051102 |
| Slc39a1 | protein_coding | 817,8537018 | 1,037495947 | 0,146102067 | 7,101172267 | 1,24E-12 | 5,08E-10 |
| Gm6781 | processed_pseudogene | 105,6388825 | 1,044246548 | 0,267931606 | 3,89743698 | 9,72E-05 | 0,003285944 |
| Trav9-2 | TR_V_gene | 316,2121023 | 1,047336608 | 0,31199809 | 3,356868658 | 7,88E-04 | 0,01512453 |
| Samd4b | protein_coding | 131,1690924 | 1,048747721 | 0,296350933 | 3,538871001 | 4,02E-04 | 0,009472973 |
| Gm17066 | processed_transcript | 108,6417589 | 1,054395247 | 0,323304271 | 3,261309364 | 0,00110899 | 0,018934973 |
| Rcbtb2 | protein_coding | 1448,628519 | 1,055357714 | 0,171893015 | 6,139619546 | 8,27E-10 | 1,91E-07 |
| Scarb1 | protein_coding | 245,216603 | 1,062533527 | 0,247927742 | 4,285658067 | 1,82E-05 | 9,30E-04 |
| Ccdc17 | protein_coding | 214,8364657 | 1,070362891 | 0,265724268 | 4,028096118 | 5,62E-05 | 0,002243267 |
| Sik1 | protein_coding | 164,2805539 | 1,07042294 | 0,371584296 | 2,880700153 | 0,003967929 | 0,044584803 |
| Hsf2 | protein_coding | 170,8995878 | 1,075801909 | 0,265209471 | 4,05642342 | 4,98E-05 | 0,002031884 |
| Bbs9 | protein_coding | 318,4853291 | 1,083728239 | 0,265593139 | 4,080407517 | 4,50E-05 | 0,001888869 |
| Socs7 | protein_coding | 124,2833643 | 1,085317594 | 0,36317141 | 2,988444478 | 0,002804014 | 0,03590924 |
| Mvk | protein_coding | 311,8212075 | 1,087391614 | 0,203689048 | 5,338488378 | 9,37E-08 | 1,13E-05 |
| Sema4a | protein_coding | 1048,492119 | 1,091463163 | 0,191257422 | 5,706775472 | 1,15E-08 | 1,84E-06 |
| 6720489N17Rik | protein_coding | 56,55970548 | 1,097250157 | 0,387393153 | 2,832394299 | 0,004620084 | 0,04935133 |
| Trak1 | protein_coding | 386,5220434 | 1,098668121 | 0,135058577 | 8,134752661 | 4,13E-16 | 2,74E-13 |
| Gm49616 | TEC | 237,9944987 | 1,101767096 | 0,368910382 | 2,986544021 | 0,002821502 | 0,03590924 |
| Tmem97 | protein_coding | 213,2738006 | 1,102287605 | 0,244868481 | 4,501549569 | 6,75E-06 | 4,32E-04 |

| | | | | | | | |
|---|---|---|---|---|---|---|---|
| Bcl6 | protein_coding | 574,7899265 | 1,115222885 | 0,276370312 | 4,035248491 | 5,45E-05 | 0,00218909 |
| Gm31718 | lincRNA | 275,6357032 | 1,122112433 | 0,381781688 | 2,939146814 | 0,003291171 | 0,039608615 |
| Gngt2 | protein_coding | 188,0776834 | 1,122455799 | 0,229223316 | 4,896778475 | 9,74E-07 | 8,55E-05 |
| Sqle | protein_coding | 1405,869022 | 1,127971666 | 0,124610363 | 9,051989211 | 1,40E-19 | 1,55E-16 |
| Osgin1 | protein_coding | 604,3913437 | 1,133687363 | 0,189643518 | 5,977991638 | 2,26E-09 | 4,31E-07 |
| Gm42646 | TEC | 265,4281468 | 1,134217038 | 0,169416835 | 6,694830751 | 2,16E-11 | 6,64E-09 |
| Dlgap5 | protein_coding | 130,3474998 | 1,13848407 | 0,361389639 | 3,150295271 | 0,001631055 | 0,024608176 |
| Palm3 | protein_coding | 40,87899145 | 1,146383336 | 0,404483878 | 2,834187956 | 0,00459423 | 0,049350061 |
| Nrip1 | protein_coding | 930,0552526 | 1,153556218 | 0,174277842 | 6,619064147 | 3,61E-11 | 1,07E-08 |
| Trio | protein_coding | 154,9402608 | 1,15757056 | 0,319276845 | 3,625601356 | 2,88E-04 | 0,007452426 |
| Gimap7 | protein_coding | 4436,659712 | 1,158146442 | 0,149787771 | 7,731915878 | 1,06E-14 | 5,86E-12 |
| Gm17173 | antisense | 183,6798872 | 1,160528527 | 0,266862863 | 4,348782425 | 1,37E-05 | 7,47E-04 |
| Hist1h2bc | protein_coding | 87,31851552 | 1,16105258 | 0,325625071 | 3,565611756 | 3,63E-04 | 0,008898836 |
| Neb | protein_coding | 620,0738054 | 1,166517298 | 0,275856347 | 4,228712915 | 2,35E-05 | 0,001150617 |
| Chek2 | protein_coding | 81,27990479 | 1,174537372 | 0,378982435 | 3,099186836 | 0,001940526 | 0,027525616 |
| Ncf1 | protein_coding | 450,3116181 | 1,175858068 | 0,260969701 | 4,505726385 | 6,61E-06 | 4,25E-04 |
| Pex26 | protein_coding | 139,5277217 | 1,183550756 | 0,316112435 | 3,74408161 | 1,81E-04 | 0,005352084 |
| Gm13546 | antisense | 149,8600705 | 1,199346699 | 0,241289476 | 4,97057195 | 6,68E-07 | 6,12E-05 |
| Il18r1 | protein_coding | 373,5750795 | 1,203038346 | 0,228279404 | 5,270025797 | 1,36E-07 | 1,60E-05 |
| Gm13502 | processed_pseudoge | 198,5034244 | 1,204695392 | 0,229689887 | 5,244877821 | 1,56E-07 | 1,80E-05 |
| Gm11725 | antisense | 88,00840844 | 1,223424121 | 0,368516657 | 3,319861118 | 9,01E-04 | 0,016582528 |
| Akap1 | protein_coding | 63,16080001 | 1,224949602 | 0,358740556 | 3,414583555 | 6,39E-04 | 0,012991983 |
| Lss | protein_coding | 347,6224073 | 1,225876188 | 0,215527517 | 5,687794325 | 1,29E-08 | 2,01E-06 |
| Arhgef39 | protein_coding | 59,99413692 | 1,245294516 | 0,387617305 | 3,212690713 | 0,001314978 | 0,02110578 |
| Clybl | protein_coding | 116,1455931 | 1,245743095 | 0,37844167 | 3,291770422 | 9,96E-04 | 0,017521513 |
| Itgb3 | protein_coding | 942,6872423 | 1,250643348 | 0,180735658 | 6,919737724 | 4,52E-12 | 1,66E-09 |
| Tk1 | protein_coding | 178,4988851 | 1,257687069 | 0,415136704 | 3,029573286 | 0,002448995 | 0,032514814 |
| Idi1 | protein_coding | 624,2997424 | 1,262161804 | 0,183006168 | 6,896826565 | 5,32E-12 | 1,88E-09 |
| Gna15 | protein_coding | 582,5560015 | 1,26283082 | 0,200979179 | 6,283391283 | 3,31E-10 | 8,59E-08 |
| Gm37593 | TEC | 60,21527601 | 1,263287142 | 0,377021286 | 3,350705087 | 8,06E-04 | 0,015323468 |
| Vipr1 | protein_coding | 284,3561567 | 1,285963306 | 0,206227615 | 6,235650374 | 4,50E-10 | 1,13E-07 |
| Hist1h2ac | protein_coding | 67,40395232 | 1,29295722 | 0,445484364 | 2,902362743 | 0,003703594 | 0,042536227 |
| Bhlhe40 | protein_coding | 519,972973 | 1,295980348 | 0,400030378 | 3,239704829 | 0,001196535 | 0,019857694 |
| Pde3b | protein_coding | 73,68132456 | 1,302886705 | 0,451458375 | 2,885950903 | 0,00390233 | 0,044086503 |
| Thra | protein_coding | 102,4770584 | 1,310174917 | 0,316340106 | 4,141665548 | 3,45E-05 | 0,001554941 |
| Stra6 | protein_coding | 204,3169904 | 1,315194841 | 0,289629953 | 4,540948979 | 5,60E-06 | 3,66E-04 |
| Repin1 | protein_coding | 105,8919442 | 1,320880309 | 0,415383888 | 3,179902608 | 0,001473246 | 0,022807821 |
| Snord104 | snoRNA | 74,44933199 | 1,328411289 | 0,336252668 | 3,950634194 | 7,79E-05 | 0,002830558 |
| Gpr183 | protein_coding | 3506,46209 | 1,328688338 | 0,118457149 | 11,21661589 | 3,38E-29 | 9,35E-26 |
| Rdh10 | protein_coding | 103,6455792 | 1,329228373 | 0,406886859 | 3,26682552 | 0,001087607 | 0,018646616 |
| Bcat1 | protein_coding | 104,7441934 | 1,332990215 | 0,463363382 | 2,876770732 | 0,004017674 | 0,045113204 |
| Gm42856 | TEC | 51,98740037 | 1,334863353 | 0,455871565 | 2,928156648 | 0,003409782 | 0,040403577 |
| Zfp608 | protein_coding | 78,79247855 | 1,341079761 | 0,343799263 | 3,900763922 | 9,59E-05 | 0,003265359 |
| AA465934 | processed_transcript | 61,88746149 | 1,34381154 | 0,348188059 | 3,859441769 | 1,14E-04 | 0,003772149 |
| Arglu1 | protein_coding | 753,2586698 | 1,343856503 | 0,175349262 | 7,66388458 | 1,80E-14 | 8,81E-12 |
| Abhd14a | protein_coding | 167,065837 | 1,34494574 | 0,306335884 | 4,390428317 | 1,13E-05 | 6,45E-04 |
| Phtf1os | antisense | 41,10128187 | 1,353310565 | 0,465636179 | 2,906369019 | 0,003656499 | 0,042317479 |
| Gm5113 | lincRNA | 58,47312588 | 1,356635026 | 0,478001652 | 2,838138781 | 0,004537745 | 0,048940685 |
| Id2 | protein_coding | 1512,612146 | 1,358291377 | 0,201751387 | 6,732500802 | 1,67E-11 | 5,54E-09 |
| Gm35570 | processed_pseudoge | 36,58693876 | 1,366410435 | 0,390235209 | 3,501504746 | 4,63E-04 | 0,010417846 |
| E130317F20Rik | bidirectional_promo | 50,01214622 | 1,382122466 | 0,435653276 | 3,172528569 | 0,001511177 | 0,02332976 |
| Ppp2cb | protein_coding | 72,15135401 | 1,388884518 | 0,389884412 | 3,562298147 | 3,68E-04 | 0,008972153 |
| Tmem140 | protein_coding | 34,89699323 | 1,391793538 | 0,446210785 | 3,11913917 | 0,001813803 | 0,026344525 |
| Rapgef4 | protein_coding | 118,8100974 | 1,393917251 | 0,34347728 | 4,058251691 | 4,94E-05 | 0,002022904 |
| 1700113A16Rik | antisense | 27,29812047 | 1,399516615 | 0,475314993 | 2,944398208 | 0,003235834 | 0,039027541 |
| Gm12940 | processed_transcript | 287,0142315 | 1,40684566 | 0,217383199 | 6,471731341 | 9,69E-11 | 2,64E-08 |
| Gm37169 | TEC | 89,89323941 | 1,410852287 | 0,323198261 | 4,365284274 | 1,27E-05 | 7,07E-04 |
| Mirt1 | lincRNA | 145,2286922 | 1,412884681 | 0,294372064 | 4,799656125 | 1,59E-06 | 1,27E-04 |
| Gm19705 | lincRNA | 79,06090174 | 1,417460471 | 0,353434333 | 4,010534172 | 6,06E-05 | 0,002349091 |
| Podnl1 | protein_coding | 505,7054244 | 1,453844099 | 0,331538619 | 4,385142525 | 1,16E-05 | 0,000654299 |
| Zfp523 | protein_coding | 100,0725186 | 1,460440937 | 0,367858225 | 3,970119029 | 7,18E-05 | 0,002655239 |
| Gm39323 | lincRNA | 135,0618874 | 1,472631828 | 0,405980894 | 3,627342693 | 2,86E-04 | 0,007425495 |
| Gm19585 | lincRNA | 770,6443714 | 1,485411153 | 0,155031674 | 9,58133986 | 9,58E-22 | 1,22E-18 |
| Mfsd2a | protein_coding | 75,05577451 | 1,512493575 | 0,4006699 | 3,774911903 | 1,60E-04 | 0,004965282 |
| Fgd6 | protein_coding | 83,13503013 | 1,528179569 | 0,406794619 | 3,756636639 | 1,72E-04 | 0,005220917 |
| Gm42908 | TEC | 45,84411661 | 1,528847805 | 0,472518714 | 3,235528585 | 0,001214177 | 0,020060149 |
| Arid3a | protein_coding | 58,51782559 | 1,532344461 | 0,517709808 | 2,959852097 | 0,003077868 | 0,037753357 |
| Rap1gap2 | protein_coding | 74,7131642 | 1,535675264 | 0,360177407 | 4,263663503 | 2,01E-05 | 0,00100225 |
| Ptgir | protein_coding | 299,3926274 | 1,53694498 | 0,26278037 | 5,848781542 | 4,95E-09 | 8,65E-07 |

| | | | | | | | |
|---|---|---|---|---|---|---|---|
| Gm36931 | TEC | 29,64233192 | 1,545949699 | 0,460885863 | 3,35430054 | 7,96E-04 | 0,015221666 |
| Traj23 | TR_J_gene | 74,59926325 | 1,567845513 | 0,547815075 | 2,861997754 | 0,004209799 | 0,046484247 |
| Pde4a | protein_coding | 55,52013531 | 1,570893311 | 0,50933656 | 3,084195075 | 0,002041037 | 0,028560753 |
| Gm37488 | TEC | 121,2390412 | 1,587645957 | 0,314967285 | 5,04066941 | 4,64E-07 | 4,50E-05 |
| Lmna | protein_coding | 255,8159024 | 1,589773451 | 0,280931883 | 5,658928545 | 1,52E-08 | 2,34E-06 |
| Cd86 | protein_coding | 1123,528334 | 1,601261882 | 0,166673034 | 9,607204276 | 7,45E-22 | 1,03E-18 |
| Gins2 | protein_coding | 64,51402273 | 1,606543213 | 0,459766617 | 3,494258073 | 4,75E-04 | 0,010632651 |
| Erf | protein_coding | 48,58838291 | 1,621051884 | 0,530237704 | 3,057217303 | 0,002234023 | 0,030458795 |
| Ect2 | protein_coding | 91,60400526 | 1,633862363 | 0,501159246 | 3,260166054 | 0,00111347 | 0,018972432 |
| Cnga1 | protein_coding | 50,05959326 | 1,638471284 | 0,495066502 | 3,309598364 | 9,34E-04 | 0,016853947 |
| Top2a | protein_coding | 179,5442048 | 1,644959294 | 0,560792403 | 2,93327671 | 0,003354048 | 0,04010359 |
| Doc2g | protein_coding | 24,33212056 | 1,653054204 | 0,544763394 | 3,034444354 | 0,002409792 | 0,032148641 |
| Rab11fip3 | protein_coding | 74,77999183 | 1,655912703 | 0,320734563 | 5,162875762 | 2,43E-07 | 2,60E-05 |
| Ccdc114 | protein_coding | 42,86585947 | 1,669048886 | 0,563860822 | 2,960036985 | 0,003076021 | 0,037753357 |
| Zfp987 | protein_coding | 18,72994508 | 1,678922293 | 0,585799677 | 2,866034858 | 0,004156485 | 0,046246379 |
| Tbl1xr1 | protein_coding | 1596,922979 | 1,727465713 | 0,167730957 | 10,29902736 | 7,12E-25 | 1,69E-21 |
| Fhl3 | protein_coding | 151,2087501 | 1,744023641 | 0,280396924 | 6,219838697 | 4,98E-10 | 1,23E-07 |
| Atp8b4 | protein_coding | 36,78331232 | 1,768154778 | 0,548600018 | 3,223030841 | 0,001268419 | 0,02052226 |
| Mfsd13a | protein_coding | 173,8831524 | 1,794782089 | 0,372023835 | 4,824373921 | 1,40E-06 | 1,15E-04 |
| Tctn1 | protein_coding | 45,19848506 | 1,8073154 | 0,460407048 | 3,925472919 | 8,66E-05 | 0,00304352 |
| Plac8 | protein_coding | 94,17942058 | 1,813022611 | 0,528741099 | 3,428942094 | 6,06E-04 | 0,012555334 |
| Aurka | protein_coding | 90,15867588 | 1,815401791 | 0,54788082 | 3,313497614 | 9,21E-04 | 0,016748126 |
| Hist2h3c2 | protein_coding | 36,88530326 | 1,822971259 | 0,621861888 | 2,931472878 | 0,003373588 | 0,040221315 |
| Gm43672 | lincRNA | 248,0424019 | 1,827469567 | 0,223992686 | 8,158612667 | 3,39E-16 | 2,34E-13 |
| 4833407H14Rik | lincRNA | 343,0110506 | 1,82777407 | 0,229355737 | 7,969166572 | 1,60E-15 | 1,02E-12 |
| Aldoc | protein_coding | 247,8412473 | 1,849240617 | 0,267096702 | 6,923487286 | 4,41E-12 | 1,66E-09 |
| Gm12592 | processed_transcript | 31,06526724 | 1,849952961 | 0,464128417 | 3,98586446 | 6,72E-05 | 0,002530227 |
| Wfikkn1 | protein_coding | 47,9164 | 1,850269687 | 0,479870858 | 3,855765888 | 1,15E-04 | 0,003806451 |
| Gm12892 | processed_pseudoge | 62,95819042 | 1,866114993 | 0,36315569 | 5,138608711 | 2,77E-07 | 2,85E-05 |
| Gm32633 | lincRNA | 143,4688512 | 1,926972092 | 0,334206143 | 5,765818881 | 8,13E-09 | 1,32E-06 |
| Rad54l | protein_coding | 44,80159863 | 1,948268668 | 0,56278472 | 3,461836468 | 5,37E-04 | 0,011533425 |
| Ccdc85c | protein_coding | 15,72596632 | 1,950108841 | 0,676035851 | 2,88462341 | 0,003918821 | 0,044217601 |
| Gm6598 | antisense | 28,39226339 | 1,978314673 | 0,483214806 | 4,094068826 | 4,24E-05 | 0,001822414 |
| Snx8 | protein_coding | 49,95846863 | 1,979370214 | 0,622000231 | 3,182266045 | 0,001461275 | 0,022707232 |
| Scd2 | protein_coding | 2778,589368 | 2,000100254 | 0,131747034 | 15,18136839 | 4,70E-52 | 7,80E-48 |
| Ncaph | protein_coding | 74,91715635 | 2,023779096 | 0,683454505 | 2,961102868 | 0,003065395 | 0,037667506 |
| Zfp239 | protein_coding | 69,88921897 | 2,034554806 | 0,658875663 | 3,087919196 | 0,002015633 | 0,028301555 |
| Cks1b | protein_coding | 168,776347 | 2,036844772 | 0,470825083 | 4,326117797 | 1,52E-05 | 8,02E-04 |
| Cdca8 | protein_coding | 119,0334183 | 2,0370673 | 0,695514417 | 2,928864233 | 0,00340203 | 0,040403577 |
| Cldn10 | protein_coding | 63,32793709 | 2,042511862 | 0,456159057 | 4,477630843 | 7,55E-06 | 4,76E-04 |
| Trav9n-2 | TR_V_gene | 88,99806444 | 2,044486166 | 0,648850181 | 3,150937192 | 0,001627475 | 0,024576499 |
| Ckb | protein_coding | 943,2200026 | 2,048830751 | 0,155506588 | 13,17520227 | 1,22E-39 | 6,74E-36 |
| Gm7240 | transcribed_processe | 31,08347597 | 2,059701561 | 0,658852145 | 3,126196941 | 0,001770829 | 0,025915945 |
| Cenpe | protein_coding | 40,50930554 | 2,067314253 | 0,623887639 | 3,313600275 | 9,21E-04 | 0,016748126 |
| Nme4 | protein_coding | 31,04390756 | 2,086094895 | 0,717006534 | 2,909450329 | 0,003620649 | 0,04208083 |
| Slco4a1 | protein_coding | 27,66866249 | 2,117851517 | 0,600020155 | 3,529633961 | 4,16E-04 | 0,009658987 |
| Scd4 | protein_coding | 35,99247455 | 2,131921183 | 0,734929112 | 2,900852816 | 0,003721486 | 0,042658519 |
| Bag2 | protein_coding | 36,0153828 | 2,173562343 | 0,58292087 | 3,728743393 | 1,92E-04 | 0,005583364 |
| Alox8 | protein_coding | 133,5005001 | 2,194099762 | 0,708438497 | 3,097092793 | 0,001954287 | 0,027649911 |
| Adam8 | protein_coding | 502,0513716 | 2,200667762 | 0,744295239 | 2,956713473 | 0,003109369 | 0,03799933 |
| Sytl3 | protein_coding | 33,23691572 | 2,247555565 | 0,771857204 | 2,911880012 | 0,003592606 | 0,041882416 |
| Myo6 | protein_coding | 336,4298492 | 2,265526119 | 0,255516398 | 8,866460766 | 7,55E-19 | 7,37E-16 |
| Cdca3 | protein_coding | 78,53570322 | 2,28482606 | 0,606933735 | 3,764539564 | 1,67E-04 | 0,005099717 |
| Pdlim4 | protein_coding | 72,6394737 | 2,285817784 | 0,549825931 | 4,157348093 | 3,22E-05 | 0,001489169 |
| 0610038B21Rik | antisense | 15,12251388 | 2,315024373 | 0,754735066 | 3,067333793 | 0,002159775 | 0,029745752 |
| Cenph | protein_coding | 26,86087243 | 2,317839892 | 0,718053611 | 3,227948241 | 0,001246815 | 0,020426601 |
| Gm48876 | lincRNA | 113,6646203 | 2,324992148 | 0,305017735 | 7,622481845 | 2,49E-14 | 1,15E-11 |
| Cenps | protein_coding | 33,1289234 | 2,371865199 | 0,553899066 | 4,282125291 | 1,85E-05 | 9,38E-04 |
| Chek1 | protein_coding | 34,69381278 | 2,375637349 | 0,70183948 | 3,384872777 | 7,12E-04 | 0,014069309 |
| Dapl1 | protein_coding | 168,8609604 | 2,390600292 | 0,248161467 | 9,633245321 | 5,79E-22 | 8,73E-19 |
| Gm28935 | lincRNA | 37,32395469 | 2,398928557 | 0,502788202 | 4,77125069 | 1,83E-06 | 1,43E-04 |
| Cobll1 | protein_coding | 118,8399097 | 2,414900228 | 0,540031397 | 4,471777462 | 7,76E-06 | 4,86E-04 |
| Gm49312 | bidirectional_promot | 25,52296911 | 2,433870382 | 0,67122887 | 3,625991805 | 2,88E-04 | 0,007452426 |
| Dtl | protein_coding | 64,07064042 | 2,444097924 | 0,593596807 | 4,117437789 | 3,83E-05 | 0,001690973 |
| Mastl | protein_coding | 24,80857273 | 2,450582237 | 0,681596133 | 3,595358187 | 3,24E-04 | 0,008145758 |
| Gm11973 | antisense | 16,49821877 | 2,481153104 | 0,700333068 | 3,542818718 | 3,96E-04 | 0,0094101 |
| Exo1 | protein_coding | 35,7455926 | 2,515313346 | 0,741473382 | 3,392317791 | 6,93E-04 | 0,013807555 |
| Fut7 | protein_coding | 150,3225225 | 2,517931999 | 0,499218815 | 5,043744189 | 4,57E-07 | 4,50E-05 |
| Siah3 | protein_coding | 39,22025087 | 2,52432664 | 0,674552853 | 3,742222166 | 1,82E-04 | 0,005376755 |

| | | | | | | | |
|---|---|---|---|---|---|---|---|
| Asns | protein_coding | 37,09758622 | 2,525622707 | 0,815877297 | 3,095591353 | 0,001964209 | 0,027766615 |
| Ccdc73 | protein_coding | 14,34470722 | 2,609427797 | 0,874514643 | 2,983858323 | 0,002846385 | 0,036150946 |
| Dnph1 | protein_coding | 62,5032881 | 2,610394759 | 0,537049742 | 4,860620075 | 1,17E-06 | 1,00E-04 |
| Gm29336 | lincRNA | 11,60679171 | 2,634445057 | 0,860870589 | 3,060210316 | 0,002211816 | 0,030311561 |
| Plk1 | protein_coding | 64,57704929 | 2,640574354 | 0,802281319 | 3,29133222 | 9,97E-04 | 0,017526899 |
| Acoxl | protein_coding | 41,71849537 | 2,65279873 | 0,749405159 | 3,539872519 | 4,00E-04 | 0,009450521 |
| Il1rl1 | protein_coding | 448,7726476 | 2,718653333 | 0,393021779 | 6,917309614 | 4,60E-12 | 1,66E-09 |
| Mmp9 | protein_coding | 657,588885 | 2,855156431 | 0,347644596 | 8,21286011 | 2,16E-16 | 1,71E-13 |
| Ramp3 | protein_coding | 70,93809379 | 3,024051597 | 0,382235274 | 7,911492735 | 2,54E-15 | 1,56E-12 |
| Gm44651 | processed_pseudoge | 10,8985555 | 3,183011266 | 1,005292552 | 3,166253704 | 0,001544161 | 0,023597503 |
| Nexn | protein_coding | 8,918912635 | 3,18895649 | 1,115526414 | 2,858701012 | 0,004253795 | 0,046845377 |
| Slc6a13 | protein_coding | 7,817960069 | 3,216050204 | 1,080538271 | 2,976340857 | 0,002917104 | 0,036435303 |
| Nuf2 | protein_coding | 28,97853727 | 3,248974457 | 1,146787214 | 2,833110117 | 0,00460975 | 0,04935133 |
| Gm14005 | lincRNA | 65,05217552 | 3,253192778 | 0,871383151 | 3,73336663 | 1,89E-04 | 0,005530164 |
| Cdk1 | protein_coding | 77,65699636 | 3,267188553 | 0,750095747 | 4,355695346 | 1,33E-05 | 7,31E-04 |
| Lrrc75b | protein_coding | 7,453357715 | 3,302952466 | 1,123973154 | 2,938639997 | 0,003296557 | 0,039611538 |
| Gm43560 | sense_intronic | 8,029347718 | 3,382868373 | 1,191224474 | 2,839824439 | 0,004513837 | 0,048905778 |
| Gas2l3 | protein_coding | 18,61539012 | 3,399575253 | 0,986194945 | 3,447163534 | 5,67E-04 | 0,011976721 |
| Nek2 | protein_coding | 25,46729669 | 3,415977399 | 1,176281766 | 2,904046886 | 0,00368373 | 0,042486058 |
| Amy1 | protein_coding | 16,35464399 | 3,485059029 | 1,231313242 | 2,830359417 | 0,004649574 | 0,049464856 |
| Cdkn3 | protein_coding | 24,3481673 | 3,517384933 | 1,150945159 | 3,056083867 | 0,002242485 | 0,030505151 |
| Aurkb | protein_coding | 51,04253562 | 3,529723409 | 0,847478256 | 4,16497224 | 3,11E-05 | 0,001455728 |
| Rasl11a | protein_coding | 44,39260909 | 3,566429568 | 0,976902985 | 3,650751019 | 2,61E-04 | 0,006920945 |
| Tmem273 | protein_coding | 139,9597381 | 3,80210249 | 0,386816898 | 9,829204746 | 8,43E-23 | 1,40E-19 |
| Ccna2 | protein_coding | 79,75130964 | 3,958980697 | 1,076480567 | 3,677707539 | 2,35E-04 | 0,006423841 |
| Selenom | protein_coding | 45,2353349 | 3,980985198 | 1,359943365 | 2,927316903 | 0,003419003 | 0,040422508 |
| Depdc1a | protein_coding | 14,56151532 | 4,001532989 | 1,392443417 | 2,873749081 | 0,004056311 | 0,045369169 |
| 1810010D01Rik | protein_coding | 6,092278854 | 4,195773839 | 1,43345635 | 2,927032859 | 0,003422127 | 0,040422508 |
| Tuba8 | protein_coding | 34,3638834 | 4,200216696 | 0,892172832 | 4,707850928 | 2,50E-06 | 1,91E-04 |
| Ccnb2 | protein_coding | 134,3365472 | 4,344484655 | 1,385399003 | 3,135908605 | 0,001713226 | 0,025409023 |
| 1700001O22Rik | protein_coding | 33,20342363 | 4,376130807 | 1,323272349 | 3,307052255 | 9,43E-04 | 0,016897686 |
| Cacnb4 | protein_coding | 10,98497822 | 4,688212163 | 1,494375043 | 3,137239333 | 0,001705468 | 0,025316597 |
| Gm38326 | lincRNA | 7,315467944 | 4,773052592 | 1,480128311 | 3,22475596 | 0,001260801 | 0,020497763 |
| Neil3 | protein_coding | 13,47382455 | 4,870157138 | 1,699884972 | 2,864992171 | 0,004170196 | 0,046246379 |
| 2210011C24Rik | protein_coding | 10,98403525 | 5,004963547 | 1,545915622 | 3,237539925 | 0,001205651 | 0,019969041 |
| Gm35584 | lincRNA | 7,662846513 | 5,03252625 | 1,418612051 | 3,547499998 | 3,89E-04 | 0,009322055 |
| Pbk | protein_coding | 23,25008035 | 5,143516057 | 1,519481563 | 3,385046704 | 7,12E-04 | 0,014069309 |
| Gm47782 | lincRNA | 8,392474101 | 5,321074159 | 1,586560339 | 3,353842919 | 7,97E-04 | 0,015221666 |
| Ltf | protein_coding | 16,02450342 | 5,369231895 | 1,821240765 | 2,948117568 | 0,003197154 | 0,038782764 |
| Gm15428 | processed_pseudoge | 20,30778823 | 5,497419498 | 1,445447342 | 3,803265147 | 1,43E-04 | 0,00452277 |
| Gatm | protein_coding | 8,076547722 | 5,565673957 | 1,800292736 | 3,091538307 | 0,001991223 | 0,028100625 |
| Cyb561 | protein_coding | 21,81890839 | 5,742614991 | 1,626722381 | 3,530175191 | 4,15E-04 | 0,009652749 |
| Pltp | protein_coding | 7,679476703 | 6,021218476 | 1,900124453 | 3,168854791 | 0,001530408 | 0,023464865 |
| 1700012B07Rik | protein_coding | 7,726810383 | 6,029308232 | 2,091531157 | 2,882724558 | 0,00394252 | 0,04438946 |
| Parpbp | protein_coding | 15,81589371 | 6,312895215 | 1,558390175 | 4,050907994 | 5,10E-05 | 0,002075284 |
| Bpifa2 | protein_coding | 62,31426255 | 6,330385522 | 1,013638807 | 6,24520833 | 4,23E-10 | 1,08E-07 |
| Cyp26b1 | protein_coding | 41,19204133 | 6,53505248 | 1,132281374 | 5,771579955 | 7,85E-09 | 1,29E-06 |
| Ispd | protein_coding | 20,24172691 | 6,539370616 | 1,469264698 | 4,450777743 | 8,56E-06 | 5,22E-04 |
| Smr3a | protein_coding | 6,8434385 | 6,689034982 | 2,010738444 | 3,326655937 | 8,79E-04 | 0,016243903 |
| Xist | lincRNA | 719,0420469 | 8,399180722 | 0,92934982 | 9,037695542 | 1,60E-19 | 1,66E-16 |
| Vax2 | protein_coding | 14,80582812 | 19,34111145 | 3,124924328 | 6,189305537 | 6,04E-10 | 1,47E-07 |

**Supplementary Table 2. DEG between NCOR1-cKO and WT effector Treg cells**

| gene_name | gene_biotype | baseMean | log2FoldChange | lfcSE | stat | pvalue | padj |
|---|---|---|---|---|---|---|---|
| Mpo | protein_coding | 26,38714309 | -22,00333324 | 3,128243022 | -7,033767225 | 2,01E-12 | 7,00E-10 |
| Mpeg1 | protein_coding | 30,1154959 | -5,855577157 | 1,0261529 | -5,706339825 | 1,15E-08 | 1,98E-06 |
| Wnt10a | protein_coding | 30,86283192 | -5,445328705 | 0,88948181 | -6,121911255 | 9,25E-10 | 2,09E-07 |
| Prss12 | protein_coding | 38,10758783 | -3,376407585 | 0,924910991 | -3,650521637 | 2,62E-04 | 0,01081392 |
| Ccr10 | protein_coding | 31,66964859 | -2,806914798 | 0,831365676 | -3,376269769 | 7,35E-04 | 0,023497627 |
| Maged1 | protein_coding | 66,8934247 | -2,709020521 | 0,63367123 | -4,275119954 | 1,91E-05 | 0,001360672 |
| Gk5 | protein_coding | 34,64611307 | -2,596286363 | 0,766069737 | -3,38909924 | 7,01E-04 | 0,022657166 |
| Gbp2b | protein_coding | 249,7258087 | -2,585554948 | 0,43194445 | -5,985850606 | 2,15E-09 | 4,27E-07 |
| Gm10499 | transcribed_unproce | 75,63349638 | -2,50063913 | 0,71555232 | -3,494697816 | 4,75E-04 | 0,017150194 |
| Wscd2 | protein_coding | 49,25980686 | -2,342687112 | 0,49504477 | -4,732273227 | 2,22E-06 | 2,30E-04 |
| Arhgap20 | protein_coding | 186,5166668 | -2,130904256 | 0,282238157 | -7,550021857 | 4,35E-14 | 2,34E-11 |
| Pla2g4f | protein_coding | 91,80603878 | -2,054367267 | 0,455246735 | -4,512645803 | 6,40E-06 | 5,61E-04 |
| Gpm6b | protein_coding | 251,2585449 | -2,035102279 | 0,42211559 | -4,821196672 | 1,43E-06 | 1,64E-04 |
| Matn2 | protein_coding | 298,4778649 | -1,963751235 | 0,218091198 | -9,004266333 | 2,17E-19 | 1,87E-16 |
| Il1rn | protein_coding | 27,97880062 | -1,929276492 | 0,497599506 | -3,877167219 | 1,06E-04 | 0,005487727 |
| Cst7 | protein_coding | 1302,520224 | -1,768154125 | 0,245987989 | -7,187969352 | 6,58E-13 | 2,57E-10 |
| Vcam1 | protein_coding | 65,34775111 | -1,656764103 | 0,517312085 | -3,202639475 | 0,001361743 | 0,03623593 |
| Gm17764 | processed_pseudoge | 105,0493983 | -1,64535502 | 0,34092194 | -4,826192829 | 1,39E-06 | 1,62E-04 |
| Gsta4 | protein_coding | 208,4569194 | -1,630936093 | 0,245700668 | -6,637898488 | 3,18E-11 | 8,92E-09 |
| Gm28942 | lincRNA | 147,1187839 | -1,618486076 | 0,274794971 | -5,889795106 | 3,87E-09 | 7,44E-07 |
| 1810006J02Rik | lincRNA | 93,39181582 | -1,610286324 | 0,450977021 | -3,570661586 | 3,56E-04 | 0,013953169 |
| Plekho1 | protein_coding | 35,77362148 | -1,583871615 | 0,391047407 | -4,050331459 | 5,11E-05 | 0,002983543 |
| Rin2 | protein_coding | 137,2683561 | -1,582289832 | 0,470600339 | -3,362279413 | 7,73E-04 | 0,024275496 |
| Tmem121b | protein_coding | 59,60314713 | -1,579894922 | 0,40907352 | -3,862129533 | 1,12E-04 | 0,005705113 |
| Ptpn5 | protein_coding | 264,0579313 | -1,570856376 | 0,314049335 | -5,001941412 | 5,68E-07 | 7,32E-05 |
| Itgae | protein_coding | 1485,255132 | -1,503259547 | 0,247994129 | -6,061673935 | 1,35E-09 | 2,85E-07 |
| Serpina3f | protein_coding | 203,942006 | -1,489380405 | 0,331889504 | -4,487579112 | 7,20E-06 | 6,19E-04 |
| Rgmb | protein_coding | 101,1851597 | -1,429008072 | 0,279528015 | -5,112217713 | 3,18E-07 | 4,23E-05 |
| Jazf1 | protein_coding | 113,1429182 | -1,3769191 | 0,394785177 | -3,48776798 | 4,87E-04 | 0,017353694 |
| Tspan5 | protein_coding | 173,72699 | -1,351119621 | 0,365358978 | -3,698060541 | 2,17E-04 | 0,009398747 |
| Asap2 | protein_coding | 57,29534138 | -1,317669629 | 0,415268316 | -3,173056021 | 0,001508434 | 0,039124458 |
| Actg2 | protein_coding | 121,2679274 | -1,291254416 | 0,300891139 | -4,291433843 | 1,78E-05 | 0,001285746 |
| Gm6831 | transcribed_unproce | 45,06775914 | -1,286175059 | 0,376820375 | -3,413231196 | 6,42E-04 | 0,021233232 |
| Trib3 | protein_coding | 99,30861129 | -1,214141604 | 0,360297789 | -3,369828073 | 7,52E-04 | 0,02380248 |
| Sccpdh | protein_coding | 201,4835929 | -1,203959013 | 0,20747458 | -5,802923011 | 6,52E-09 | 1,22E-06 |
| Cd7 | protein_coding | 612,7592412 | -1,202915108 | 0,184877184 | -6,506563351 | 7,69E-11 | 1,98E-08 |
| Fam174b | protein_coding | 141,2336248 | -1,189634841 | 0,249929934 | -4,759873389 | 1,94E-06 | 2,06E-04 |
| Gdpd5 | protein_coding | 367,2384371 | -1,18799623 | 0,316098008 | -3,758316087 | 1,71E-04 | 0,007876123 |
| Fgf13 | protein_coding | 279,5940786 | -1,184477257 | 0,272545366 | -4,345982013 | 1,39E-05 | 0,001045337 |
| B3gnt5 | protein_coding | 188,4720264 | -1,181013277 | 0,3735525 | -3,161572412 | 0,001569198 | 0,0400351 |
| Mgmt | protein_coding | 117,6601245 | -1,166461375 | 0,217175413 | -5,371056312 | 7,83E-08 | 1,17E-05 |
| Colq | protein_coding | 418,7795586 | -1,162102346 | 0,274154024 | -4,238866632 | 2,25E-05 | 0,001548771 |
| Laptm4b | protein_coding | 257,3913019 | -1,124449736 | 0,218806222 | -5,139020851 | 2,76E-07 | 3,75E-05 |
| Sh3bgrl2 | protein_coding | 424,5723128 | -1,110802072 | 0,233989991 | -4,747220456 | 2,06E-06 | 2,16E-04 |
| Capg | protein_coding | 5963,728336 | -1,088744085 | 0,165155737 | -6,592226851 | 4,33E-11 | 1,19E-08 |
| Iigp1 | protein_coding | 1359,645377 | -1,079394511 | 0,156638861 | -6,890975208 | 5,54E-12 | 1,72E-09 |
| Fam49a | protein_coding | 692,5114571 | -1,053418694 | 0,128260217 | -8,213136696 | 2,15E-16 | 1,39E-13 |
| Pstpip2 | protein_coding | 77,3299837 | -1,048223636 | 0,301961784 | -3,471378473 | 5,18E-04 | 0,018272553 |
| Mctp1 | protein_coding | 121,0029531 | -1,028381732 | 0,329541596 | -3,120643173 | 0,001804566 | 0,043566402 |
| Irf8 | protein_coding | 697,4093419 | -1,015453649 | 0,180001919 | -5,64134902 | 1,69E-08 | 2,82E-06 |
| Gpr83 | protein_coding | 3106,526949 | -0,996164875 | 0,233603643 | -4,264337928 | 2,00E-05 | 0,001406567 |
| Arhgap5 | protein_coding | 312,5010764 | -0,995731938 | 0,292876613 | -3,399834241 | 6,74E-04 | 0,021951142 |
| Serpinc1 | protein_coding | 93,64706239 | -0,987326815 | 0,279679837 | -3,530203765 | 4,15E-04 | 0,015652838 |
| Trav13n-1 | TR_V_gene | 74,60962783 | -0,984773088 | 0,277566649 | -3,547879723 | 3,88E-04 | 0,014812279 |
| Ddx3y | protein_coding | 1560,867423 | -0,972407754 | 0,245465889 | -3,961478134 | 7,45E-05 | 0,004103799 |
| Uty | protein_coding | 422,3845167 | -0,970653495 | 0,310352209 | -3,127586872 | 0,001762477 | 0,043362322 |
| Kdm5d | protein_coding | 461,9421005 | -0,95734408 | 0,198772318 | -4,816284727 | 1,46E-06 | 1,67E-04 |
| Acsl1 | protein_coding | 250,5648606 | -0,94403854 | 0,187481076 | -5,035380431 | 4,77E-07 | 6,21E-05 |
| Serpini1 | protein_coding | 269,9158978 | -0,930506916 | 0,223085896 | -4,171070128 | 3,03E-05 | 0,00192537 |
| Dtx1 | protein_coding | 4250,235398 | -0,92341842 | 0,127177001 | -7,260891636 | 3,85E-13 | 1,65E-10 |
| Cdyl2 | protein_coding | 341,3698912 | -0,90136214 | 0,224890578 | -4,008003129 | 6,12E-05 | 0,00350859 |
| Rrm2b | protein_coding | 338,7146157 | -0,89899118 | 0,246928709 | -3,640691204 | 2,72E-04 | 0,011128337 |
| Pard6g | protein_coding | 362,1339608 | -0,891782174 | 0,17849672 | -4,996070385 | 5,85E-07 | 7,47E-05 |
| Rflnb | protein_coding | 650,3784476 | -0,89081083 | 0,198869704 | -4,479369223 | 7,49E-06 | 6,39E-04 |
| Cnn3 | protein_coding | 405,9624853 | -0,889137525 | 0,226967557 | -3,917465276 | 8,95E-05 | 0,004826944 |
| Nedd4l | protein_coding | 188,3681977 | -0,88862952 | 0,249387978 | -3,56324121 | 3,66E-04 | 0,014267035 |
| Slc25a24 | protein_coding | 429,4648259 | -0,88085923 | 0,23634434 | -3,727016393 | 1,94E-04 | 0,008613628 |

| | | | | | | | |
|---|---|---|---|---|---|---|---|
| L3mbtl3 | protein_coding | 346,1677051 | -0,869977601 | 0,2248571 | -3,869024379 | 1,09E-04 | 0,005634924 |
| Irf6 | protein_coding | 359,915028 | -0,857806279 | 0,227322012 | -3,773529324 | 1,61E-04 | 0,007518198 |
| Supt7l | protein_coding | 470,3728319 | -0,838614679 | 0,191476054 | -4,379736581 | 1,19E-05 | 9,17E-04 |
| Camsap1 | protein_coding | 144,2208781 | -0,836306168 | 0,258564812 | -3,234416013 | 0,001218918 | 0,033528588 |
| Fam129a | protein_coding | 1824,733082 | -0,82149743 | 0,141818281 | -5,792606039 | 6,93E-09 | 1,28E-06 |
| Hmgn5 | protein_coding | 289,7081204 | -0,819783061 | 0,196034616 | -4,181828082 | 2,89E-05 | 0,001860055 |
| Cd96 | protein_coding | 5249,007804 | -0,81804655 | 0,135361186 | -6,043435138 | 1,51E-09 | 3,14E-07 |
| Ces2c | protein_coding | 264,2728983 | -0,815816437 | 0,233374248 | -3,495743192 | 4,73E-04 | 0,017150194 |
| Atp6v0d2 | protein_coding | 231,9557521 | -0,813474307 | 0,251358532 | -3,236310698 | 0,001210855 | 0,033528588 |
| Usp6nl | protein_coding | 270,9506382 | -0,812033042 | 0,246513568 | -3,294070382 | 9,87E-04 | 0,028858201 |
| Fry | protein_coding | 172,6981449 | -0,811387917 | 0,256719636 | -3,16059936 | 0,001574449 | 0,0400351 |
| Itprid2 | protein_coding | 133,8833278 | -0,799889756 | 0,258204124 | -3,097896902 | 0,001948992 | 0,045601465 |
| Endod1 | protein_coding | 874,1624837 | -0,795943114 | 0,230973066 | -3,446042983 | 5,69E-04 | 0,019714359 |
| Sh3bgrl | protein_coding | 2244,768085 | -0,783473545 | 0,166932825 | -4,693346229 | 2,69E-06 | 2,68E-04 |
| Tbc1d4 | protein_coding | 2293,457403 | -0,782524437 | 0,133640095 | -5,855461555 | 4,76E-09 | 9,02E-07 |
| AW112010 | lincRNA | 8734,818791 | -0,775412513 | 0,211908356 | -3,659188001 | 2,53E-04 | 0,010522188 |
| Ctse | protein_coding | 497,6970325 | -0,763110273 | 0,19758429 | -3,86220116 | 1,12E-04 | 0,005705113 |
| Ephb6 | protein_coding | 1133,714337 | -0,76184633 | 0,153358685 | -4,967741672 | 6,77E-07 | 8,48E-05 |
| Hspa4l | protein_coding | 357,8051635 | -0,755339313 | 0,236950305 | -3,187754128 | 0,001433824 | 0,037570852 |
| Litaf | protein_coding | 378,5337117 | -0,753055167 | 0,169423611 | -4,444806491 | 8,80E-06 | 7,18E-04 |
| Ppp1r21 | protein_coding | 330,8196852 | -0,751499888 | 0,200785987 | -3,742790519 | 1,82E-04 | 0,008290419 |
| Cdc25b | protein_coding | 1338,625811 | -0,750218141 | 0,236587238 | -3,17100004 | 0,001519151 | 0,03919414 |
| Fbxo32 | protein_coding | 588,6875177 | -0,740106059 | 0,175086152 | -4,227096497 | 2,37E-05 | 0,001614748 |
| Tnfrsf9 | protein_coding | 4705,482046 | -0,738872089 | 0,163962678 | -4,506343129 | 6,60E-06 | 5,75E-04 |
| Slc14a1 | protein_coding | 1502,22016 | -0,735941802 | 0,164900686 | -4,462939587 | 8,08E-06 | 6,72E-04 |
| Clip1 | protein_coding | 1168,289592 | -0,735860826 | 0,159923348 | -4,601334554 | 4,20E-06 | 3,89E-04 |
| Armcx2 | protein_coding | 394,7603489 | -0,726822635 | 0,209108704 | -3,475812447 | 5,09E-04 | 0,01803847 |
| Atg14 | protein_coding | 354,0767441 | -0,721254833 | 0,235536551 | -3,062177957 | 0,002197328 | 0,049524387 |
| Arhgap31 | protein_coding | 879,6799308 | -0,710993969 | 0,21587635 | -3,293524136 | 9,89E-04 | 0,028858201 |
| Dcun1d4 | protein_coding | 450,090493 | -0,702313393 | 0,175829525 | -3,994285901 | 6,49E-05 | 0,003685268 |
| Selenop | protein_coding | 2388,348064 | -0,686065505 | 0,094649354 | -7,248496426 | 4,21E-13 | 1,71E-10 |
| Nt5e | protein_coding | 2204,386058 | -0,682661872 | 0,176601295 | -3,865554162 | 1,11E-04 | 0,005670293 |
| Wdr47 | protein_coding | 295,0533713 | -0,67786648 | 0,201712153 | -3,3605634 | 7,78E-04 | 0,024275496 |
| Gbp3 | protein_coding | 1312,986171 | -0,67004807 | 0,115763901 | -5,788057123 | 7,12E-09 | 1,29E-06 |
| Ncor1 | protein_coding | 1545,045843 | -0,669546579 | 0,120644887 | -5,549730236 | 2,86E-08 | 4,55E-06 |
| Id3 | protein_coding | 3951,235214 | -0,668270928 | 0,149990135 | -4,455432534 | 8,37E-06 | 6,92E-04 |
| Flna | protein_coding | 6709,231231 | -0,668077813 | 0,095824329 | -6,971901804 | 3,13E-12 | 1,01E-09 |
| Tiam1 | protein_coding | 438,3896885 | -0,667873707 | 0,165457186 | -4,036534904 | 5,42E-05 | 0,003136078 |
| Igf1r | protein_coding | 507,6371919 | -0,666703797 | 0,178492715 | -3,735187714 | 1,88E-04 | 0,008396596 |
| Ift140 | protein_coding | 492,588342 | -0,665230317 | 0,13386152 | -4,969541025 | 6,71E-07 | 8,48E-05 |
| Slc41a1 | protein_coding | 848,1113317 | -0,664269317 | 0,155151616 | -4,28142054 | 1,86E-05 | 0,001330054 |
| Nrde2 | protein_coding | 529,4301168 | -0,662765918 | 0,173458137 | -3,820898398 | 1,33E-04 | 0,006493194 |
| Cdc7 | protein_coding | 332,9500974 | -0,656546441 | 0,213722937 | -3,07195124 | 0,002126645 | 0,048353972 |
| Vwa5a | protein_coding | 614,1509464 | -0,635115836 | 0,169889338 | -3,738409032 | 1,85E-04 | 0,008377017 |
| Gbp2 | protein_coding | 1933,244133 | -0,633319711 | 0,152850222 | -4,143400667 | 3,42E-05 | 0,002111597 |
| Pip4k2a | protein_coding | 489,2483226 | -0,632975907 | 0,197412061 | -3,206368967 | 0,001344215 | 0,035879132 |
| Crbn | protein_coding | 1622,659351 | -0,632154007 | 0,145196091 | -4,353794959 | 1,34E-05 | 0,001014683 |
| Exoc4 | protein_coding | 736,9488585 | -0,630735966 | 0,19984532 | -3,156120779 | 0,001598826 | 0,040415824 |
| Ldhb | protein_coding | 1171,83398 | -0,630006057 | 0,170700466 | -3,690710828 | 2,24E-04 | 0,009545269 |
| Ssbp2 | protein_coding | 1218,559847 | -0,627311157 | 0,147411747 | -4,255503182 | 2,09E-05 | 0,001453517 |
| Trim59 | protein_coding | 3089,749962 | -0,626311434 | 0,164498284 | -3,80740405 | 1,40E-04 | 0,006780767 |
| Lipa | protein_coding | 2541,079182 | -0,624849498 | 0,119056119 | -5,248361043 | 1,53E-07 | 2,22E-05 |
| Odf2 | protein_coding | 910,371311 | -0,62239747 | 0,14486146 | -4,296501433 | 1,74E-05 | 0,001263815 |
| Eif4enif1 | protein_coding | 603,1416952 | -0,616743925 | 0,138907518 | -4,43996072 | 9,00E-06 | 7,25E-04 |
| Ccdc125 | protein_coding | 352,0638452 | -0,613567824 | 0,159529684 | -3,846104425 | 1,20E-04 | 0,005950681 |
| L1cam | protein_coding | 618,215568 | -0,609554109 | 0,18324601 | -3,326425003 | 8,80E-04 | 0,026435418 |
| Ppm1d | protein_coding | 649,4588734 | -0,600675362 | 0,163173089 | -3,681215863 | 2,32E-04 | 0,009811631 |
| Cd82 | protein_coding | 8373,056254 | -0,595326151 | 0,152829927 | -3,895350628 | 9,81E-05 | 0,005159788 |
| Adamts6 | protein_coding | 553,5277178 | -0,591743908 | 0,186621755 | -3,170819547 | 0,001520095 | 0,03919414 |
| Rad17 | protein_coding | 559,8994174 | -0,589893131 | 0,161953923 | -3,642351604 | 2,70E-04 | 0,011091994 |
| Rgcc | protein_coding | 216,0974687 | -0,586387681 | 0,189943424 | -3,087170218 | 0,002020718 | 0,046603046 |
| Ptpre | protein_coding | 700,1243794 | -0,584840269 | 0,134126786 | -4,360354015 | 1,30E-05 | 9,91E-04 |
| Stat1 | protein_coding | 4652,2502 | -0,58474469 | 0,135178909 | -4,325709504 | 1,52E-05 | 0,001126507 |
| Cldnd1 | protein_coding | 1263,397543 | -0,583303775 | 0,179713317 | -3,245745976 | 0,001171433 | 0,032688563 |
| Lta | protein_coding | 1965,811767 | -0,583267288 | 0,12900181 | -4,521388412 | 6,14E-06 | 5,50E-04 |
| Gzf1 | protein_coding | 355,4352345 | -0,574482055 | 0,172053616 | -3,338971128 | 8,41E-04 | 0,025628352 |
| Spata13 | protein_coding | 3302,808783 | -0,572096159 | 0,110753302 | -5,165499798 | 2,40E-07 | 3,36E-05 |
| Orai2 | protein_coding | 1570,488276 | -0,566669514 | 0,162314949 | -3,491172667 | 4,81E-04 | 0,017221755 |
| Slamf6 | protein_coding | 4816,433926 | -0,563410812 | 0,126718545 | -4,446159112 | 8,74E-06 | 7,18E-04 |

| | | | | | | | |
|---|---|---|---|---|---|---|---|
| Nfatc1 | protein_coding | 4213,419108 | -0,557500425 | 0,118977564 | -4,685760956 | 2,79E-06 | 2,72E-04 |
| Card6 | protein_coding | 1501,591666 | -0,557035774 | 0,143515824 | -3,881354385 | 1,04E-04 | 0,005443796 |
| Eps15 | protein_coding | 1853,110392 | -0,556407662 | 0,167191512 | -3,327965973 | 8,75E-04 | 0,026351053 |
| Pdk1 | protein_coding | 2502,482911 | -0,552416563 | 0,128286139 | -4,306128206 | 1,66E-05 | 0,001223906 |
| Tgfbr1 | protein_coding | 1493,028995 | -0,550689359 | 0,161616909 | -3,407374646 | 6,56E-04 | 0,021516528 |
| Pdlim5 | protein_coding | 634,6443931 | -0,549937896 | 0,160413072 | -3,428261103 | 6,07E-04 | 0,020447483 |
| Anp32e | protein_coding | 1965,26434 | -0,549164173 | 0,143718402 | -3,821112436 | 1,33E-04 | 0,006493194 |
| Hif1a | protein_coding | 6006,524052 | -0,539436852 | 0,136679939 | -3,946715647 | 7,92E-05 | 0,004309875 |
| Susd6 | protein_coding | 2002,098136 | -0,538048105 | 0,114668629 | -4,6921997 | 2,70E-06 | 2,68E-04 |
| Pcmtd2 | protein_coding | 1410,218566 | -0,537057652 | 0,149993458 | -3,580540505 | 3,43E-04 | 0,013518235 |
| Slc2a3 | protein_coding | 1753,403271 | -0,53408611 | 0,110159274 | -4,848308171 | 1,25E-06 | 1,49E-04 |
| Cbfa2t2 | protein_coding | 485,8101237 | -0,533741849 | 0,169446112 | -3,149920892 | 0,001633147 | 0,041202599 |
| Lclat1 | protein_coding | 1269,345974 | -0,532492773 | 0,125433187 | -4,245230368 | 2,18E-05 | 0,001513555 |
| Cd1d2 | polymorphic_pseudo | 486,1141031 | -0,530841353 | 0,170433566 | -3,114652624 | 0,001841617 | 0,044073183 |
| Hsd11b1 | protein_coding | 1587,957334 | -0,529105365 | 0,137875296 | -3,837564661 | 1,24E-04 | 0,006114376 |
| Armc7 | protein_coding | 1443,76693 | -0,528862302 | 0,140655769 | -3,759975915 | 1,70E-04 | 0,007852093 |
| Zc3h12d | protein_coding | 833,1618826 | -0,527375994 | 0,130641925 | -4,036805133 | 5,42E-05 | 0,003136078 |
| Prnp | protein_coding | 527,9032954 | -0,526656548 | 0,143480609 | -3,670576459 | 2,42E-04 | 0,010129599 |
| Pitpnm2 | protein_coding | 694,98939 | -0,526183559 | 0,150658102 | -3,492567292 | 4,78E-04 | 0,017179794 |
| Rfwd3 | protein_coding | 724,8914438 | -0,525926997 | 0,154731007 | -3,398976112 | 6,76E-04 | 0,021964663 |
| Madd | protein_coding | 1862,106903 | -0,525501781 | 0,138936685 | -3,7823112 | 1,55E-04 | 0,007310749 |
| Gprasp1 | protein_coding | 583,211322 | -0,523156344 | 0,152669605 | -3,426722332 | 6,11E-04 | 0,020510145 |
| Camk2d | protein_coding | 2135,569694 | -0,521339713 | 0,095327889 | -5,468910693 | 4,53E-08 | 7,03E-06 |
| Capn2 | protein_coding | 2090,816602 | -0,517346833 | 0,116520423 | -4,439966997 | 9,00E-06 | 7,25E-04 |
| Dok1 | protein_coding | 828,4945339 | -0,516158653 | 0,156863575 | -3,290494 | 0,001000116 | 0,028909193 |
| Cyfip1 | protein_coding | 1123,152107 | -0,513417218 | 0,158840699 | -3,23227751 | 0,001228077 | 0,033614379 |
| Dgka | protein_coding | 18022,50005 | -0,512163763 | 0,165256049 | -3,099213399 | 0,001940352 | 0,045594789 |
| Micu2 | protein_coding | 1599,495092 | -0,510075037 | 0,14596429 | -3,494519358 | 4,75E-04 | 0,017150194 |
| Mboat7 | protein_coding | 1388,903611 | -0,508625849 | 0,124155282 | -4,096691169 | 4,19E-05 | 0,002501391 |
| Triobp | protein_coding | 1174,378186 | -0,507095816 | 0,12681308 | -3,998765877 | 6,37E-05 | 0,003632214 |
| Cux1 | protein_coding | 1112,044627 | -0,504928402 | 0,12305494 | -4,10327615 | 4,07E-05 | 0,002442528 |
| Rab3ip | protein_coding | 1178,429625 | -0,504408107 | 0,143428644 | -3,516787811 | 4,37E-04 | 0,016275323 |
| Lrrc32 | protein_coding | 6013,570429 | -0,502536535 | 0,143251616 | -3,508068879 | 4,51E-04 | 0,01667361 |
| Ift80 | protein_coding | 4179,56884 | -0,502235944 | 0,144338395 | -3,479572732 | 5,02E-04 | 0,017836207 |
| Ikzf1 | protein_coding | 4757,138589 | -0,498569399 | 0,10586637 | -4,709421888 | 2,48E-06 | 2,50E-04 |
| Nrp1 | protein_coding | 3704,052472 | -0,49611904 | 0,118150109 | -4,199056992 | 2,68E-05 | 0,001772014 |
| Lcp2 | protein_coding | 8991,260757 | -0,495488004 | 0,157749344 | -3,140982986 | 0,001683818 | 0,042233042 |
| Trim26 | protein_coding | 1803,978866 | -0,493938262 | 0,134814568 | -3,663834475 | 2,48E-04 | 0,010366478 |
| Tmem131l | protein_coding | 1345,796503 | -0,489954836 | 0,126993571 | -3,858107392 | 1,14E-04 | 0,005736266 |
| Tagap | protein_coding | 2062,659073 | -0,48923277 | 0,116488932 | -4,199821909 | 2,67E-05 | 0,001772014 |
| Ipcef1 | protein_coding | 1829,584974 | -0,489133699 | 0,151635218 | -3,225726219 | 0,001256535 | 0,034165663 |
| Cand1 | protein_coding | 1969,542438 | -0,488834897 | 0,109307644 | -4,472101666 | 7,75E-06 | 6,53E-04 |
| Cd84 | protein_coding | 2400,073847 | -0,487371862 | 0,13173006 | -3,699777119 | 2,16E-04 | 0,009398747 |
| Ephx1 | protein_coding | 2948,161331 | -0,484088817 | 0,107857862 | -4,488210767 | 7,18E-06 | 6,19E-04 |
| Pigk | protein_coding | 983,5667963 | -0,482932939 | 0,155861312 | -3,0984786 | 0,00194517 | 0,045594789 |
| Oxct1 | protein_coding | 1149,298656 | -0,482593979 | 0,140225509 | -3,441556274 | 5,78E-04 | 0,019932181 |
| Ccng1 | protein_coding | 4005,693734 | -0,480710648 | 0,135572259 | -3,545789174 | 3,91E-04 | 0,014853663 |
| Nedd9 | protein_coding | 2541,862832 | -0,480332248 | 0,14717729 | -3,263630201 | 0,001099946 | 0,030961807 |
| Dennd2d | protein_coding | 2179,551962 | -0,479952246 | 0,11680858 | -4,108878356 | 3,98E-05 | 0,002395173 |
| Cltc | protein_coding | 3161,712491 | -0,479773122 | 0,141943261 | -3,380034529 | 7,25E-04 | 0,023243033 |
| Tbc1d14 | protein_coding | 1749,87439 | -0,478633991 | 0,136217784 | -3,513740836 | 4,42E-04 | 0,016415702 |
| Ttc3 | protein_coding | 1050,33883 | -0,478021336 | 0,142793378 | -3,347643591 | 8,15E-04 | 0,025136857 |
| Il6st | protein_coding | 3129,014358 | -0,477075664 | 0,122993115 | -3,878881056 | 1,05E-04 | 0,005477171 |
| Mbnl2 | protein_coding | 1103,693953 | -0,476907179 | 0,150016354 | -3,179034591 | 0,001477665 | 0,038407362 |
| Il16 | protein_coding | 6765,518501 | -0,476769447 | 0,148882015 | -3,202330707 | 0,001363204 | 0,03623593 |
| Gbp7 | protein_coding | 3188,99622 | -0,474938053 | 0,129998902 | -3,653400501 | 2,59E-04 | 0,010727723 |
| Pitpnm1 | protein_coding | 4434,389148 | -0,472459344 | 0,118852874 | -3,975161291 | 7,03E-05 | 0,003925172 |
| Cnst | protein_coding | 888,7183825 | -0,470020318 | 0,139265542 | -3,374993634 | 7,38E-04 | 0,023497627 |
| Iqgap1 | protein_coding | 3599,23205 | -0,469459984 | 0,09637863 | -4,870996652 | 1,11E-06 | 1,35E-04 |
| Pbxip1 | protein_coding | 3125,456042 | -0,468640445 | 0,150471146 | -3,114487102 | 0,00184265 | 0,044073183 |
| Msh2 | protein_coding | 1205,603533 | -0,467307953 | 0,12279373 | -3,805633654 | 1,41E-04 | 0,006803975 |
| Btg2 | protein_coding | 5309,175816 | -0,464071941 | 0,150007578 | -3,093656646 | 0,001977062 | 0,045842225 |
| Fam102a | protein_coding | 2731,648404 | -0,459560143 | 0,130115406 | -3,531942593 | 4,13E-04 | 0,015595876 |
| Mms19 | protein_coding | 1411,613371 | -0,458236608 | 0,145145344 | -3,157087885 | 0,001593533 | 0,040361157 |
| Tmem229b | protein_coding | 1314,846926 | -0,45784959 | 0,133823547 | -3,42129318 | 6,23E-04 | 0,020815606 |
| Cntrl | protein_coding | 1589,841919 | -0,456718822 | 0,123950923 | -3,684674653 | 2,29E-04 | 0,009711194 |
| Atp2a3 | protein_coding | 4955,57799 | -0,456612954 | 0,096635218 | -4,725119515 | 2,30E-06 | 2,35E-04 |
| Fam3c | protein_coding | 2193,434742 | -0,456247576 | 0,10812367 | -4,219682687 | 2,45E-05 | 0,00165999 |
| Hsdl2 | protein_coding | 768,5394308 | -0,455305816 | 0,116452546 | -3,909796994 | 9,24E-05 | 0,004941419 |

| | | | | | | | |
|---|---|---|---|---|---|---|---|
| Ypel5 | protein_coding | 2024,623249 | -0,454626176 | 0,135841274 | -3,346745537 | 8,18E-04 | 0,02515824 |
| Rfk | protein_coding | 1835,226498 | -0,453529548 | 0,147457774 | -3,075657097 | 0,002100392 | 0,048123525 |
| Tnfaip8 | protein_coding | 1883,573204 | -0,453057459 | 0,122452935 | -3,699849743 | 2,16E-04 | 0,009398747 |
| Anxa5 | protein_coding | 1935,66986 | -0,452023994 | 0,138089641 | -3,273409879 | 0,001062583 | 0,030041263 |
| Ivns1abp | protein_coding | 3355,802713 | -0,449559383 | 0,097639628 | -4,604271797 | 4,14E-06 | 3,87E-04 |
| Hip1r | protein_coding | 1033,018659 | -0,446405761 | 0,131843431 | -3,385877917 | 7,10E-04 | 0,022867488 |
| Arhgef6 | protein_coding | 2155,878999 | -0,44371072 | 0,139572983 | -3,179058802 | 0,001477541 | 0,038407362 |
| Dock2 | protein_coding | 7111,407829 | -0,440338987 | 0,119315998 | -3,690527636 | 2,24E-04 | 0,009545269 |
| Arid5a | protein_coding | 2468,131602 | -0,437639168 | 0,132657714 | -3,299010321 | 9,70E-04 | 0,028493477 |
| Tgoln1 | protein_coding | 2453,801528 | -0,436632321 | 0,12175947 | -3,586023501 | 3,36E-04 | 0,013318773 |
| Traf1 | protein_coding | 4444,880195 | -0,434698512 | 0,116888209 | -3,718925259 | 2,00E-04 | 0,008833326 |
| Tspyl1 | protein_coding | 1349,902905 | -0,434456627 | 0,109400802 | -3,971238029 | 7,15E-05 | 0,003973186 |
| Acot9 | protein_coding | 1500,556473 | -0,434398804 | 0,12155631 | -3,573642567 | 3,52E-04 | 0,01383726 |
| Tollip | protein_coding | 1096,068032 | -0,432669667 | 0,132302695 | -3,270301239 | 0,00107433 | 0,030306923 |
| Rnf19a | protein_coding | 1095,132864 | -0,432653557 | 0,12610278 | -3,430959708 | 6,01E-04 | 0,020387452 |
| Ssh2 | protein_coding | 2967,032176 | -0,42993934 | 0,114680779 | -3,749009585 | 1,78E-04 | 0,008145097 |
| Zfp110 | protein_coding | 1408,876847 | -0,428120071 | 0,12651898 | -3,383840689 | 7,15E-04 | 0,022980381 |
| Fli1 | protein_coding | 2816,548657 | -0,426605224 | 0,118310847 | -3,605799748 | 3,11E-04 | 0,012459319 |
| Hsd17b4 | protein_coding | 1253,993095 | -0,426028288 | 0,118417131 | -3,597691362 | 3,21E-04 | 0,01281434 |
| Stag1 | protein_coding | 1341,217624 | -0,424517318 | 0,134363514 | -3,159468714 | 0,001580571 | 0,040111648 |
| Trip12 | protein_coding | 2006,251571 | -0,422317902 | 0,1143618 | -3,692823146 | 2,22E-04 | 0,009530551 |
| Hp1bp3 | protein_coding | 4103,104189 | -0,419434396 | 0,108720075 | -3,857929599 | 1,14E-04 | 0,005736266 |
| Rab3gap1 | protein_coding | 1345,596359 | -0,417987706 | 0,130762196 | -3,196548521 | 0,001390824 | 0,036812387 |
| Mapk9 | protein_coding | 850,4693583 | -0,416392348 | 0,123885282 | -3,361112321 | 7,76E-04 | 0,024275496 |
| Zdhhc17 | protein_coding | 497,5016873 | -0,415986216 | 0,135388275 | -3,072542403 | 0,002122437 | 0,04834356 |
| Atp6v1d | protein_coding | 2288,508269 | -0,412209849 | 0,116259867 | -3,545590247 | 3,92E-04 | 0,014853663 |
| Rabgap1l | protein_coding | 2357,767127 | -0,411127089 | 0,122689805 | -3,350947464 | 8,05E-04 | 0,024898434 |
| Mbp | protein_coding | 2404,527521 | -0,410918916 | 0,13319011 | -3,085205908 | 0,002034113 | 0,046828182 |
| Scamp2 | protein_coding | 1924,884464 | -0,410539754 | 0,124892174 | -3,28715357 | 0,001012056 | 0,029123726 |
| Cog5 | protein_coding | 770,3624111 | -0,408066355 | 0,110339056 | -3,698294777 | 2,17E-04 | 0,009398747 |
| Trappc12 | protein_coding | 1501,54442 | -0,406841535 | 0,128644732 | -3,162519977 | 0,0015641 | 0,0400351 |
| Tax1bp1 | protein_coding | 4701,454171 | -0,406042198 | 0,132207432 | -3,071250929 | 0,002131639 | 0,048382207 |
| Dnajc1 | protein_coding | 975,9205795 | -0,404606431 | 0,121227116 | -3,337590173 | 8,45E-04 | 0,025695305 |
| Glud1 | protein_coding | 3295,072981 | -0,403117313 | 0,121611437 | -3,314797715 | 9,17E-04 | 0,02724236 |
| Plekhg2 | protein_coding | 2650,001836 | -0,399874237 | 0,12910709 | -3,097229103 | 0,001953388 | 0,045621528 |
| Zfand5 | protein_coding | 1426,069988 | -0,399262873 | 0,129580609 | -3,081193054 | 0,002061729 | 0,047379347 |
| Tbc1d15 | protein_coding | 1994,753209 | -0,398837146 | 0,118407644 | -3,368339523 | 7,56E-04 | 0,023836774 |
| Tnfrsf1b | protein_coding | 5898,514631 | -0,394373044 | 0,112319319 | -3,511177306 | 4,46E-04 | 0,016527201 |
| Exoc2 | protein_coding | 1803,561601 | -0,391999302 | 0,119660364 | -3,275932715 | 0,001053136 | 0,02986952 |
| Stat3 | protein_coding | 5153,839568 | -0,391987786 | 0,121244316 | -3,233040514 | 0,001224802 | 0,033596056 |
| Slc9a9 | protein_coding | 902,2572958 | -0,384724965 | 0,113467009 | -3,390632836 | 6,97E-04 | 0,022587384 |
| Polg | protein_coding | 1871,732483 | -0,382520639 | 0,115997644 | -3,297658698 | 9,75E-04 | 0,028565907 |
| Stat5b | protein_coding | 3492,796862 | -0,381581921 | 0,105605507 | -3,613276722 | 3,02E-04 | 0,012258765 |
| Lrp10 | protein_coding | 7469,552039 | -0,378910259 | 0,105051265 | -3,606908101 | 3,10E-04 | 0,01244489 |
| Hsp90b1 | protein_coding | 3836,754221 | -0,378298541 | 0,11750103 | -3,219533814 | 0,001283992 | 0,034702782 |
| Epsti1 | protein_coding | 1977,194391 | -0,375700355 | 0,114694901 | -3,275650021 | 0,001054191 | 0,02986952 |
| Dcaf11 | protein_coding | 2659,020377 | -0,374580077 | 0,10963896 | -3,416486956 | 6,34E-04 | 0,021131788 |
| Il6ra | protein_coding | 4730,64895 | -0,366396677 | 0,099205931 | -3,693294089 | 2,21E-04 | 0,009530551 |
| Vsir | protein_coding | 5489,001939 | -0,365246642 | 0,109892726 | -3,323665302 | 8,88E-04 | 0,026574496 |
| Cd28 | protein_coding | 6081,572398 | -0,363708582 | 0,118818083 | -3,061054124 | 0,002205592 | 0,0496239 |
| Fam53b | protein_coding | 3627,261443 | -0,363516324 | 0,118394223 | -3,07038904 | 0,002137801 | 0,048389859 |
| Zfp869 | protein_coding | 941,5335482 | -0,362636489 | 0,110676265 | -3,276551563 | 0,001050831 | 0,02986952 |
| Srpk2 | protein_coding | 1358,837479 | -0,362019233 | 0,11277909 | -3,209985409 | 0,001327417 | 0,035504276 |
| Kctd10 | protein_coding | 1904,045677 | -0,361245636 | 0,111694719 | -3,234223057 | 0,001219742 | 0,033528588 |
| Adss | protein_coding | 2117,740146 | -0,360923565 | 0,108222198 | -3,33502343 | 8,53E-04 | 0,025811906 |
| Slc28a2 | protein_coding | 2698,216174 | -0,356393897 | 0,110130023 | -3,236119331 | 0,001211667 | 0,033528588 |
| Hbp1 | protein_coding | 3051,282517 | -0,352797458 | 0,113121385 | -3,118751235 | 0,001816192 | 0,043714627 |
| Adcy7 | protein_coding | 2689,450592 | -0,3492171 | 0,104486239 | -3,342230558 | 8,31E-04 | 0,025418749 |
| Abcf3 | protein_coding | 1428,449741 | -0,342664225 | 0,107310071 | -3,193215912 | 0,001406977 | 0,037093554 |
| Wipf1 | protein_coding | 2241,129 | -0,341838489 | 0,111746165 | -3,059062385 | 0,002220309 | 0,049867987 |
| Brd8 | protein_coding | 1998,978615 | -0,341570467 | 0,109854628 | -3,109295204 | 0,001875342 | 0,044772064 |
| Wwp2 | protein_coding | 2237,564753 | -0,335188822 | 0,088396736 | -3,791868776 | 1,50E-04 | 0,007060751 |
| Lrch1 | protein_coding | 1149,055005 | -0,334360293 | 0,098331033 | -3,400353704 | 6,73E-04 | 0,021951142 |
| Smc1a | protein_coding | 2067,681205 | -0,332764499 | 0,106209122 | -3,13310659 | 0,001729666 | 0,042965041 |
| Myh9 | protein_coding | 4683,050146 | -0,307663295 | 0,098505405 | -3,123313827 | 0,00178827 | 0,04343521 |
| Arhgap27 | protein_coding | 2674,016262 | -0,304737672 | 0,092755515 | -3,285386037 | 0,001018427 | 0,02920798 |
| Ptpn7 | protein_coding | 6284,670365 | -0,282808519 | 0,088090686 | -3,210424769 | 0,00132539 | 0,035504276 |
| Prkd2 | protein_coding | 4354,735584 | -0,27591056 | 0,089040779 | -3,098698849 | 0,001943725 | 0,045594789 |
| Rin3 | protein_coding | 2996,864759 | -0,252520878 | 0,080452231 | -3,138767878 | 0,001696598 | 0,042353333 |

| | | | | | | | |
|---|---|---|---|---|---|---|---|
| Psd4 | protein_coding | 4093,471617 | -0,25028028 | 0,078310403 | -3,19600297 | 0,001393457 | 0,036812387 |
| Ncoa3 | protein_coding | 4260,167021 | 0,270222288 | 0,07298921 | 3,702222425 | 2,14E-04 | 0,009398747 |
| Ndufb3 | protein_coding | 970,4975099 | 0,308258807 | 0,098627761 | 3,125477095 | 0,001775169 | 0,04343521 |
| Mapkapk3 | protein_coding | 2901,281024 | 0,312776432 | 0,100295748 | 3,118541278 | 0,001817487 | 0,043714627 |
| Pa2g4 | protein_coding | 2685,272437 | 0,331463388 | 0,102482501 | 3,234341327 | 0,001219236 | 0,033528588 |
| Gm16286 | protein_coding | 2461,230234 | 0,350093949 | 0,101025404 | 3,465405078 | 5,29E-04 | 0,018597971 |
| Bccip | protein_coding | 1119,081851 | 0,357848842 | 0,116395138 | 3,074431184 | 0,002109044 | 0,048123525 |
| Gemin7 | protein_coding | 1254,662948 | 0,359488085 | 0,110174072 | 3,262910023 | 0,001102745 | 0,030972967 |
| Rad23a | protein_coding | 1506,563059 | 0,370441777 | 0,107703465 | 3,439460145 | 5,83E-04 | 0,019932181 |
| Snrnp27 | protein_coding | 767,3711613 | 0,373098287 | 0,119250285 | 3,128699349 | 0,001755819 | 0,043281097 |
| Mrpl39 | protein_coding | 703,3176244 | 0,37330525 | 0,120167033 | 3,106552936 | 0,001892824 | 0,044939759 |
| Mrpl19 | protein_coding | 552,0713271 | 0,37932972 | 0,118846873 | 3,191751784 | 0,001414128 | 0,037205996 |
| Erg28 | protein_coding | 1054,598568 | 0,379577601 | 0,113792959 | 3,335686181 | 8,51E-04 | 0,025811045 |
| Cox7a2 | protein_coding | 1418,568683 | 0,379642014 | 0,117329482 | 3,23569156 | 0,001213484 | 0,033528588 |
| Ep400 | protein_coding | 2050,054882 | 0,382406568 | 0,124374535 | 3,074637165 | 0,002107588 | 0,048123525 |
| Med15 | protein_coding | 2355,75965 | 0,388144072 | 0,103852325 | 3,737461551 | 1,86E-04 | 0,008379237 |
| Sem1 | protein_coding | 1747,005945 | 0,388244707 | 0,099516994 | 3,901290558 | 9,57E-05 | 0,005076224 |
| Nutf2 | protein_coding | 807,0445536 | 0,388797371 | 0,125258036 | 3,103971469 | 0,001909417 | 0,045084782 |
| Pop5 | protein_coding | 949,0347591 | 0,389737576 | 0,113068987 | 3,446900747 | 5,67E-04 | 0,019714359 |
| Fam162a | protein_coding | 597,9621436 | 0,394184596 | 0,122167295 | 3,22659674 | 0,001252719 | 0,03414387 |
| Guk1 | protein_coding | 979,1380007 | 0,396931585 | 0,113633137 | 3,493097138 | 4,77E-04 | 0,017179794 |
| Tomm20 | protein_coding | 1601,844592 | 0,401268159 | 0,126955899 | 3,160689353 | 0,001573963 | 0,0400351 |
| Ppdpf | protein_coding | 1671,608092 | 0,411040521 | 0,133882699 | 3,070154121 | 0,002139483 | 0,048389859 |
| Dnmt3a | protein_coding | 1044,82914 | 0,417209323 | 0,12675253 | 3,29152659 | 9,96E-04 | 0,028867997 |
| Macf1 | protein_coding | 3672,065808 | 0,420013164 | 0,077455994 | 5,42260377 | 5,87E-08 | 8,91E-06 |
| Ifi206 | protein_coding | 4765,138076 | 0,421836334 | 0,1226114 | 3,440433233 | 5,81E-04 | 0,019932181 |
| Uqcrq | protein_coding | 1260,676204 | 0,422478691 | 0,134703806 | 3,136353031 | 0,001710632 | 0,04257425 |
| Atad3a | protein_coding | 1103,27466 | 0,422880947 | 0,127881098 | 3,30682918 | 9,44E-04 | 0,027865159 |
| Tomm7 | protein_coding | 1031,327089 | 0,422980263 | 0,138017298 | 3,064690217 | 0,002178956 | 0,049196315 |
| Nfkbiz | protein_coding | 2358,363085 | 0,426856207 | 0,135554431 | 3,148965351 | 0,001638496 | 0,041256823 |
| Prrc2a | protein_coding | 1608,395987 | 0,436474883 | 0,114890656 | 3,799045972 | 1,45E-04 | 0,006910024 |
| Mrpl30 | protein_coding | 1171,378263 | 0,442924849 | 0,113679883 | 3,89624652 | 9,77E-05 | 0,005159788 |
| Gpr183 | protein_coding | 3506,46209 | 0,454136375 | 0,117897567 | 3,851957129 | 1,17E-04 | 0,005832637 |
| Hmg20b | protein_coding | 1147,519201 | 0,457037724 | 0,147864944 | 3,09091331 | 0,001995419 | 0,046102042 |
| Mrps24 | protein_coding | 1091,933943 | 0,458851101 | 0,118647326 | 3,86735308 | 1,10E-04 | 0,005651066 |
| Tomm5 | protein_coding | 729,6727549 | 0,462122553 | 0,122748106 | 3,764803931 | 1,67E-04 | 0,007757532 |
| Mid1ip1 | protein_coding | 354,684545 | 0,462296119 | 0,149333117 | 3,095737418 | 0,001963241 | 0,045658273 |
| Dmac1 | protein_coding | 691,9423364 | 0,465427447 | 0,143594164 | 3,241269943 | 0,001189984 | 0,033134512 |
| Ndufb1-ps | bidirectional_promo | 688,22125 | 0,466211009 | 0,131303441 | 3,550638162 | 3,84E-04 | 0,014745162 |
| Nt5c | protein_coding | 2402,005866 | 0,469023797 | 0,11322957 | 4,142237743 | 3,44E-05 | 0,002111597 |
| Elovl5 | protein_coding | 5222,65527 | 0,471354947 | 0,141964666 | 3,320227202 | 8,99E-04 | 0,026841686 |
| Stard4 | protein_coding | 370,4209515 | 0,477129237 | 0,14958991 | 3,189581678 | 0,001424789 | 0,037410134 |
| Sdhd | protein_coding | 1379,970923 | 0,481818808 | 0,112536081 | 4,281460719 | 1,86E-05 | 0,001330054 |
| Atxn2l | protein_coding | 3793,407412 | 0,482546344 | 0,128925196 | 3,742839707 | 1,82E-04 | 0,008290419 |
| Tsfm | protein_coding | 424,454969 | 0,488878915 | 0,124721302 | 3,919770772 | 8,86E-05 | 0,004801091 |
| Hspe1 | protein_coding | 1210,858873 | 0,488890351 | 0,126850425 | 3,854069481 | 1,16E-04 | 0,005804928 |
| Atp5mpl | protein_coding | 1123,702954 | 0,489291892 | 0,133079066 | 3,676700683 | 2,36E-04 | 0,009954225 |
| Rpf2 | protein_coding | 584,5454323 | 0,491529203 | 0,13471317 | 3,648709354 | 2,64E-04 | 0,010855685 |
| Cd200r1 | protein_coding | 953,0720765 | 0,492192118 | 0,148882779 | 3,305903622 | 9,47E-04 | 0,027865159 |
| Bcat2 | protein_coding | 1036,207931 | 0,492778065 | 0,140509289 | 3,507085326 | 4,53E-04 | 0,016687547 |
| Ndufa1 | protein_coding | 1141,9924 | 0,49466764 | 0,117693567 | 4,203013391 | 2,63E-05 | 0,001768521 |
| Gas5 | processed_transcript | 3218,554435 | 0,501558286 | 0,12597565 | 3,981390743 | 6,85E-05 | 0,003840315 |
| Eif4ebp2 | protein_coding | 1074,060267 | 0,501875525 | 0,13889856 | 3,613252191 | 3,02E-04 | 0,012258765 |
| B630019A10Rik | antisense | 466,8856192 | 0,504576693 | 0,14433309 | 3,495918322 | 4,72E-04 | 0,017150194 |
| Slc25a22 | protein_coding | 1191,537496 | 0,508546025 | 0,154877276 | 3,283541892 | 0,001025114 | 0,029303261 |
| Lyar | protein_coding | 625,9835916 | 0,510195187 | 0,153984247 | 3,31329469 | 9,22E-04 | 0,027326231 |
| 2410006H16Rik | processed_transcript | 845,4047831 | 0,510428095 | 0,10923843 | 4,672605538 | 2,97E-06 | 2,88E-04 |
| Rpa3 | protein_coding | 280,327083 | 0,511569039 | 0,164793075 | 3,104311514 | 0,001907224 | 0,045084782 |
| Mknk2 | protein_coding | 1192,402303 | 0,513742214 | 0,123505125 | 4,159683359 | 3,19E-05 | 0,00201399 |
| BC005537 | protein_coding | 990,8809637 | 0,51741672 | 0,127415421 | 4,060864195 | 4,89E-05 | 0,002878119 |
| Ptger2 | protein_coding | 1343,888073 | 0,517877882 | 0,155431151 | 3,331879596 | 8,63E-04 | 0,026044125 |
| Usp16 | protein_coding | 1035,08173 | 0,520586459 | 0,151465361 | 3,437000096 | 5,88E-04 | 0,020007955 |
| Romo1 | protein_coding | 795,0535357 | 0,520780166 | 0,10710631 | 4,862273432 | 1,16E-06 | 1,40E-04 |
| Tmem181b-ps | transcribed_unproce | 669,9528705 | 0,521011677 | 0,155091953 | 3,359372719 | 7,81E-04 | 0,024275496 |
| Csrp1 | protein_coding | 724,2242756 | 0,522273758 | 0,150819622 | 3,462903231 | 5,34E-04 | 0,018720745 |
| Ldb1 | protein_coding | 1035,914669 | 0,523464457 | 0,123667567 | 4,232835429 | 2,31E-05 | 0,001582448 |
| Bst2 | protein_coding | 1488,573774 | 0,525860049 | 0,142513349 | 3,689900311 | 2,24E-04 | 0,009545269 |
| Prmt7 | protein_coding | 815,4313012 | 0,526991024 | 0,156355413 | 3,370468696 | 7,50E-04 | 0,02380248 |
| Rcbtb2 | protein_coding | 1448,628519 | 0,528604202 | 0,171010612 | 3,091060812 | 0,001994428 | 0,046102042 |

| | | | | | | | |
|---|---|---|---|---|---|---|---|
| Sesn1 | protein_coding | 7027,893073 | 0,531776712 | 0,170591548 | 3,117251222 | 0,001825459 | 0,043824624 |
| Ubxn7 | protein_coding | 692,9662203 | 0,533681767 | 0,171011395 | 3,120738051 | 0,001803984 | 0,043566402 |
| Tesc | protein_coding | 1178,445553 | 0,538056541 | 0,150188612 | 3,582538874 | 3,40E-04 | 0,01345636 |
| Chchd1 | protein_coding | 670,1803977 | 0,54214181 | 0,134955238 | 4,017197238 | 5,89E-05 | 0,003389585 |
| Nsdhl | protein_coding | 510,2560577 | 0,542922156 | 0,157827001 | 3,439982729 | 5,82E-04 | 0,019932181 |
| St8sia6 | protein_coding | 2436,3748 | 0,544266237 | 0,120040463 | 4,534023128 | 5,79E-06 | 5,22E-04 |
| Rap2b | protein_coding | 519,202318 | 0,54612083 | 0,156080903 | 3,498959946 | 4,67E-04 | 0,017106686 |
| Sirt6 | protein_coding | 596,7552681 | 0,547091533 | 0,16025625 | 3,413854582 | 6,41E-04 | 0,021233232 |
| Ndufab1 | protein_coding | 484,8801207 | 0,548105474 | 0,13136233 | 4,172470715 | 3,01E-05 | 0,001923042 |
| Rasgrp4 | protein_coding | 374,2668629 | 0,548182699 | 0,154426608 | 3,549794331 | 3,86E-04 | 0,01474861 |
| Sigmar1 | protein_coding | 665,7144634 | 0,549618785 | 0,175861368 | 3,12529574 | 0,001776264 | 0,04343521 |
| Usmg5 | protein_coding | 883,2849443 | 0,549632408 | 0,135450976 | 4,057795831 | 4,95E-05 | 0,002902929 |
| Aacs | protein_coding | 1335,609196 | 0,555346524 | 0,167302299 | 3,319419573 | 9,02E-04 | 0,026857277 |
| Bola2 | protein_coding | 542,287775 | 0,557476357 | 0,175721444 | 3,172500436 | 0,001511323 | 0,039124458 |
| Tob1 | protein_coding | 1249,791679 | 0,559506328 | 0,159775341 | 3,50183153 | 4,62E-04 | 0,016971591 |
| Srm | protein_coding | 1204,678694 | 0,559696958 | 0,136786366 | 4,09175984 | 4,28E-05 | 0,002543419 |
| Pet100 | protein_coding | 352,9848724 | 0,559896328 | 0,177839524 | 3,148323366 | 0,001642099 | 0,041266948 |
| Snhg6 | lincRNA | 300,9481503 | 0,562799218 | 0,181773116 | 3,096163114 | 0,001960425 | 0,045658273 |
| Ppm1j | protein_coding | 846,9217215 | 0,56319515 | 0,120714259 | 4,665522982 | 3,08E-06 | 2,96E-04 |
| Mxd1 | protein_coding | 1704,197574 | 0,569972006 | 0,149841284 | 3,803838235 | 1,42E-04 | 0,006816297 |
| Hdac5 | protein_coding | 710,8496173 | 0,572666686 | 0,1775597 | 3,225206437 | 0,001258819 | 0,034165663 |
| Aars2 | protein_coding | 349,2850187 | 0,573897566 | 0,181385095 | 3,163973122 | 0,001556312 | 0,039968068 |
| Wapl | protein_coding | 932,8691011 | 0,578684154 | 0,164049734 | 3,527492191 | 4,20E-04 | 0,015722099 |
| Notch2 | protein_coding | 737,8864796 | 0,580099507 | 0,134959617 | 4,298319143 | 1,72E-05 | 0,00126062 |
| Ung | protein_coding | 261,6530537 | 0,58217995 | 0,18204745 | 3,197957182 | 0,001384048 | 0,036714302 |
| Gemin2 | protein_coding | 365,5451198 | 0,586574088 | 0,171024389 | 3,429768649 | 6,04E-04 | 0,020387452 |
| Dph5 | protein_coding | 1007,806571 | 0,587273041 | 0,140282289 | 4,186366253 | 2,83E-05 | 0,001845612 |
| Rock2 | protein_coding | 281,8501632 | 0,588375837 | 0,182885186 | 3,217186961 | 0,001294542 | 0,034914721 |
| Ost4 | protein_coding | 580,3120409 | 0,588784072 | 0,154349919 | 3,81460564 | 1,36E-04 | 0,006635759 |
| Nme1 | protein_coding | 1123,593307 | 0,589885051 | 0,123904062 | 4,760820916 | 1,93E-06 | 2,06E-04 |
| Skil | protein_coding | 2218,462933 | 0,590344969 | 0,152290106 | 3,876449915 | 1,06E-04 | 0,005487727 |
| Ntpcr | protein_coding | 306,8981202 | 0,596670331 | 0,192761367 | 3,095383377 | 0,001965587 | 0,045658273 |
| Abhd2 | protein_coding | 878,1853231 | 0,597498981 | 0,185123619 | 3,227567522 | 0,001248476 | 0,034100316 |
| Gm4459 | processed_pseudoge | 191,2398556 | 0,602028038 | 0,188985153 | 3,185583785 | 0,001444623 | 0,037777029 |
| Hemk1 | protein_coding | 386,1865387 | 0,606058843 | 0,134284712 | 4,513237829 | 6,38E-06 | 5,61E-04 |
| Dazap1 | protein_coding | 269,3692869 | 0,60869734 | 0,165614838 | 3,675379251 | 2,37E-04 | 0,009973299 |
| Mettl1 | protein_coding | 216,2497706 | 0,611042524 | 0,193268935 | 3,161617905 | 0,001568953 | 0,0400351 |
| Tmf1 | protein_coding | 1153,604043 | 0,612638884 | 0,175659407 | 3,487652021 | 4,87E-04 | 0,017353694 |
| Elovl6 | protein_coding | 293,2521855 | 0,628695669 | 0,20075269 | 3,131692377 | 0,001738019 | 0,043074425 |
| Rln3 | protein_coding | 1159,901044 | 0,629092166 | 0,135700831 | 4,635875571 | 3,55E-06 | 3,37E-04 |
| Ifi213 | protein_coding | 2630,570137 | 0,630315964 | 0,129108756 | 4,88205435 | 1,05E-06 | 1,29E-04 |
| Gar1 | protein_coding | 291,6195665 | 0,635373577 | 0,203437395 | 3,123189708 | 0,001789024 | 0,04343521 |
| Mterf1a | protein_coding | 318,7347053 | 0,640933889 | 0,1861573 | 3,442969406 | 5,75E-04 | 0,019886321 |
| Cdca7 | protein_coding | 527,5311092 | 0,642257468 | 0,192127232 | 3,342875762 | 8,29E-04 | 0,025418749 |
| Pfas | protein_coding | 513,2718501 | 0,642878476 | 0,168842436 | 3,807564563 | 1,40E-04 | 0,006780767 |
| Birc6 | protein_coding | 2361,096939 | 0,645157774 | 0,109196996 | 5,908200754 | 3,46E-09 | 6,76E-07 |
| Nudt3 | protein_coding | 306,1946874 | 0,650395121 | 0,17407949 | 3,736196148 | 1,87E-04 | 0,008392151 |
| Uros | protein_coding | 588,1986199 | 0,657508157 | 0,200149497 | 3,285085228 | 0,001019515 | 0,02920798 |
| Crebl2 | protein_coding | 202,1997093 | 0,664771374 | 0,19546925 | 3,400900003 | 6,72E-04 | 0,021951142 |
| H2afy | protein_coding | 681,683265 | 0,667916605 | 0,197897874 | 3,375057006 | 7,38E-04 | 0,023497627 |
| Snhg15 | processed_transcript | 234,3045694 | 0,669230295 | 0,180919734 | 3,699045313 | 2,16E-04 | 0,009398747 |
| Malt1 | protein_coding | 1565,766431 | 0,671727222 | 0,204036044 | 3,292198815 | 9,94E-04 | 0,028863942 |
| Nop16 | protein_coding | 657,3433329 | 0,678483649 | 0,131013998 | 5,17871111 | 2,23E-07 | 3,17E-05 |
| Flot1 | protein_coding | 655,5401507 | 0,679452621 | 0,161741512 | 4,200854886 | 2,66E-05 | 0,001772014 |
| Arf6 | protein_coding | 400,5192539 | 0,679970973 | 0,186899243 | 3,638168677 | 2,75E-04 | 0,011202315 |
| Smox | protein_coding | 464,9606141 | 0,683127304 | 0,21265344 | 3,212397146 | 0,001316323 | 0,035387803 |
| Rnf213 | protein_coding | 1346,95577 | 0,68973293 | 0,193495138 | 3,564600832 | 3,64E-04 | 0,014236297 |
| Sytl2 | protein_coding | 754,1463584 | 0,691950515 | 0,170193353 | 4,065672981 | 4,79E-05 | 0,00283234 |
| Cox16 | protein_coding | 292,7583915 | 0,692676908 | 0,207269845 | 3,341908756 | 8,32E-04 | 0,025418749 |
| Ifit1 | protein_coding | 1612,60729 | 0,695804288 | 0,222774627 | 3,123355198 | 0,001788018 | 0,04343521 |
| Ptpmt1 | processed_transcript | 262,1169326 | 0,701120772 | 0,176810734 | 3,965374466 | 7,33E-05 | 0,004054662 |
| Wdfy1 | protein_coding | 200,408403 | 0,702305206 | 0,199029802 | 3,528643447 | 4,18E-04 | 0,015699505 |
| Kdm6b | protein_coding | 2045,275709 | 0,704366636 | 0,109227183 | 6,448638667 | 1,13E-10 | 2,80E-08 |
| Pcyt2 | protein_coding | 2205,380149 | 0,713616416 | 0,134188149 | 5,318028613 | 1,05E-07 | 1,55E-05 |
| Cdk9 | protein_coding | 279,8187507 | 0,7165665 | 0,213309146 | 3,359286345 | 7,81E-04 | 0,024275496 |
| Jak2 | protein_coding | 669,9927636 | 0,72286365 | 0,158719076 | 4,55435899 | 5,25E-06 | 4,77E-04 |
| Wtap | protein_coding | 2039,036942 | 0,724326828 | 0,15076378 | 4,804382248 | 1,55E-06 | 1,74E-04 |
| Nle1 | protein_coding | 691,8323223 | 0,725973083 | 0,17362883 | 4,18117823 | 2,90E-05 | 0,001860055 |
| Ydjc | protein_coding | 180,3215399 | 0,726317756 | 0,23369252 | 3,108006004 | 0,001883542 | 0,044884711 |

| | | | | | | | |
|---|---|---|---|---|---|---|---|
| Lysmd2 | protein_coding | 1044,971645 | 0,727077006 | 0,111068724 | 6,546190335 | 5,90E-11 | 1,59E-08 |
| Uck2 | protein_coding | 1343,922257 | 0,735110014 | 0,152048663 | 4,834702245 | 1,33E-06 | 1,58E-04 |
| D17H6S53E | protein_coding | 343,5633826 | 0,736533007 | 0,204171961 | 3,607415061 | 3,09E-04 | 0,01244489 |
| Npm3 | protein_coding | 225,4809354 | 0,737851877 | 0,215131596 | 3,429769926 | 6,04E-04 | 0,020387452 |
| Tatdn1 | protein_coding | 153,3319394 | 0,743923299 | 0,19951234 | 3,728708204 | 1,92E-04 | 0,008585619 |
| Abhd11 | protein_coding | 630,7292244 | 0,744260166 | 0,170022566 | 4,377419904 | 1,20E-05 | 9,22E-04 |
| Srebf1 | protein_coding | 2569,172647 | 0,747153829 | 0,155674109 | 4,799473922 | 1,59E-06 | 1,77E-04 |
| 1110038B12Rik | processed_transcript | 157,8101552 | 0,753692327 | 0,242575646 | 3,10704038 | 0,001889706 | 0,044939759 |
| Epn1 | protein_coding | 334,7765711 | 0,754727814 | 0,147612464 | 5,112900312 | 3,17E-07 | 4,23E-05 |
| Gimap7 | protein_coding | 4436,659712 | 0,757694117 | 0,148994525 | 5,085382265 | 3,67E-07 | 4,83E-05 |
| Arap2 | protein_coding | 996,4695323 | 0,760257525 | 0,166732477 | 4,559744667 | 5,12E-06 | 4,68E-04 |
| Itih5 | protein_coding | 925,9540053 | 0,761825832 | 0,223207795 | 3,413078969 | 6,42E-04 | 0,021233232 |
| Snord78 | snoRNA | 180,0382606 | 0,762181861 | 0,181910144 | 4,189881028 | 2,79E-05 | 0,001835799 |
| Tmem259 | protein_coding | 208,8958333 | 0,762433828 | 0,247936959 | 3,075111638 | 0,002104238 | 0,048123525 |
| Tbl1x | protein_coding | 1896,325856 | 0,762815527 | 0,104041354 | 7,331849315 | 2,27E-13 | 1,01E-10 |
| Vmn2r97 | protein_coding | 574,8082869 | 0,771161335 | 0,187143908 | 4,120686291 | 3,78E-05 | 0,002286336 |
| Hmgcr | protein_coding | 2029,432219 | 0,773845931 | 0,126659995 | 6,109631764 | 9,99E-10 | 2,18E-07 |
| Srebf2 | protein_coding | 3633,200365 | 0,778625488 | 0,106018651 | 7,344231257 | 2,07E-13 | 9,53E-11 |
| Bach2 | protein_coding | 684,4772934 | 0,779773527 | 0,188180896 | 4,143744371 | 3,42E-05 | 0,002111597 |
| Tmem63b | protein_coding | 248,9742045 | 0,788556017 | 0,239789524 | 3,288534053 | 0,001007106 | 0,029046109 |
| Ifrd2 | protein_coding | 404,7067847 | 0,792447901 | 0,179164047 | 4,423029693 | 9,73E-06 | 7,75E-04 |
| Taf1d | protein_coding | 516,0885177 | 0,795938057 | 0,191969307 | 4,146173508 | 3,38E-05 | 0,002111597 |
| 8030462N17Rik | protein_coding | 433,8158797 | 0,802191346 | 0,202656252 | 3,95838439 | 7,55E-05 | 0,004122078 |
| Dhrs3 | protein_coding | 556,4235581 | 0,805108086 | 0,194006428 | 4,14990418 | 3,33E-05 | 0,002091739 |
| Ankrd55 | protein_coding | 195,6957713 | 0,807032746 | 0,226747497 | 3,559169376 | 3,72E-04 | 0,014403018 |
| Hnrnpul2 | protein_coding | 141,846994 | 0,808554345 | 0,23295358 | 3,470881811 | 5,19E-04 | 0,018272553 |
| Csnk1g2 | protein_coding | 374,9154461 | 0,811456698 | 0,203624107 | 3,985071846 | 6,75E-05 | 0,003797776 |
| Arhgef10 | protein_coding | 375,5937758 | 0,811544365 | 0,246466858 | 3,292711932 | 9,92E-04 | 0,028863942 |
| Sc5d | protein_coding | 2000,502511 | 0,822557443 | 0,145106168 | 5,668659397 | 1,44E-08 | 2,44E-06 |
| 4833407H14Rik | lincRNA | 343,0110506 | 0,824660696 | 0,211153639 | 3,905500746 | 9,40E-05 | 0,005009259 |
| Ogg1 | protein_coding | 132,9139417 | 0,830813913 | 0,266214195 | 3,12084753 | 0,001803314 | 0,043566402 |
| Nrm | protein_coding | 383,5455155 | 0,831578001 | 0,197547029 | 4,209519138 | 2,56E-05 | 0,001727358 |
| Abcg1 | protein_coding | 4907,357257 | 0,832440634 | 0,158811557 | 5,241688011 | 1,59E-07 | 2,28E-05 |
| Snhg1 | processed_transcript | 343,5082523 | 0,837296949 | 0,190143221 | 4,403506715 | 1,07E-05 | 8,37E-04 |
| Gpr34 | protein_coding | 962,9703807 | 0,837523756 | 0,151656498 | 5,522504926 | 3,34E-08 | 5,25E-06 |
| Alg8 | protein_coding | 170,6747875 | 0,861497111 | 0,188826459 | 4,562374965 | 5,06E-06 | 4,66E-04 |
| Plcxd2 | protein_coding | 613,259178 | 0,864139045 | 0,232051584 | 3,723909277 | 1,96E-04 | 0,008690424 |
| Gngt2 | protein_coding | 188,0776834 | 0,870349152 | 0,232811297 | 3,738431786 | 1,85E-04 | 0,008377017 |
| AI504432 | lincRNA | 322,7401054 | 0,871592629 | 0,230698567 | 3,778058263 | 1,58E-04 | 0,007409661 |
| Rsad2 | protein_coding | 301,4144301 | 0,873314126 | 0,220586818 | 3,959049475 | 7,52E-05 | 0,004122078 |
| Larp1 | protein_coding | 207,1005625 | 0,874779187 | 0,245602549 | 3,561767549 | 3,68E-04 | 0,014304158 |
| Tmem256 | protein_coding | 321,771398 | 0,876071282 | 0,209215928 | 4,187402419 | 2,82E-05 | 0,001845612 |
| Mbd6 | protein_coding | 381,9776685 | 0,881257548 | 0,203201209 | 4,33687158 | 1,45E-05 | 0,001077004 |
| Shmt1 | protein_coding | 338,2567923 | 0,885090319 | 0,20037277 | 4,417218558 | 1,00E-05 | 7,91E-04 |
| Trim56 | protein_coding | 545,6493849 | 0,904474201 | 0,188901842 | 4,788064478 | 1,68E-06 | 1,86E-04 |
| Snhg12 | lincRNA | 117,9755196 | 0,912505379 | 0,278194964 | 3,280093087 | 0,001037728 | 0,029598218 |
| Igsf23 | protein_coding | 476,8888686 | 0,923671084 | 0,186100299 | 4,963297144 | 6,93E-07 | 8,59E-05 |
| Sema4a | protein_coding | 1048,492119 | 0,930538823 | 0,200405253 | 4,643285585 | 3,43E-06 | 3,27E-04 |
| Pde4d | protein_coding | 270,4195321 | 0,937845514 | 0,239545259 | 3,915107804 | 9,04E-05 | 0,004854043 |
| Car12 | protein_coding | 199,1092849 | 0,941960976 | 0,220907385 | 4,264053808 | 2,01E-05 | 0,001406567 |
| Cdkn1b | protein_coding | 1172,105578 | 0,943283307 | 0,154354969 | 6,111130156 | 9,89E-10 | 2,18E-07 |
| Hsf2 | protein_coding | 170,8995878 | 0,944425235 | 0,265647083 | 3,555187671 | 3,78E-04 | 0,01453561 |
| Igfbp4 | protein_coding | 3706,253023 | 0,948058899 | 0,213999552 | 4,430191055 | 9,41E-06 | 7,54E-04 |
| Usp18 | protein_coding | 2317,745702 | 0,963710728 | 0,167713696 | 5,746165939 | 9,13E-09 | 1,61E-06 |
| Scarb1 | protein_coding | 245,216603 | 0,984738723 | 0,246805347 | 3,989940799 | 6,61E-05 | 0,003736972 |
| Tmem120b | protein_coding | 162,5663733 | 0,992177585 | 0,275047604 | 3,607294051 | 3,09E-04 | 0,01244489 |
| Satb1 | protein_coding | 2500,346361 | 1,003930202 | 0,163837701 | 6,127589659 | 8,92E-10 | 2,05E-07 |
| Abca1 | protein_coding | 346,0291353 | 1,017072551 | 0,298090423 | 3,411959838 | 6,45E-04 | 0,021266068 |
| Zfas1 | processed_transcript | 391,6007768 | 1,025507175 | 0,149185227 | 6,87405313 | 6,24E-12 | 1,87E-09 |
| Gm48876 | lincRNA | 113,6646203 | 1,030597882 | 0,328285648 | 3,139332738 | 0,001693331 | 0,042353333 |
| Oasl1 | protein_coding | 241,7912516 | 1,033071403 | 0,307413659 | 3,360525379 | 7,78E-04 | 0,024275496 |
| A630081D01Rik | TEC | 63,70293281 | 1,034303709 | 0,33037798 | 3,130667812 | 0,001744093 | 0,043074425 |
| Slc39a1 | protein_coding | 817,8537018 | 1,039494984 | 0,145753706 | 7,131859714 | 9,90E-13 | 3,75E-10 |
| Lss | protein_coding | 347,6224073 | 1,040728621 | 0,218416232 | 4,764886803 | 1,89E-06 | 2,05E-04 |
| Pan3 | protein_coding | 1007,231118 | 1,041959132 | 0,169237839 | 6,156774046 | 7,42E-10 | 1,74E-07 |
| Gm45716 | protein_coding | 151,2077382 | 1,054762177 | 0,296645336 | 3,555633778 | 3,77E-04 | 0,01453561 |
| Dctd | protein_coding | 96,5749062 | 1,055074045 | 0,313133068 | 3,369411132 | 7,53E-04 | 0,02380248 |
| Irak3 | protein_coding | 179,9839375 | 1,062799476 | 0,327002322 | 3,250128219 | 0,00115353 | 0,032258801 |
| Mtln | protein_coding | 194,4613342 | 1,063156518 | 0,221179194 | 4,806765488 | 1,53E-06 | 1,73E-04 |

| | | | | | | | |
|---|---|---|---|---|---|---|---|
| Gm12892 | processed_pseudoge | 62,95819042 | 1,06928789 | 0,337520532 | 3,168067681 | 0,001534558 | 0,039488065 |
| Tm7sf2 | protein_coding | 210,8695289 | 1,077238962 | 0,257619511 | 4,181511548 | 2,90E-05 | 0,001860055 |
| Ckb | protein_coding | 943,2200026 | 1,096304485 | 0,15126216 | 7,247711448 | 4,24E-13 | 1,71E-10 |
| Tcrg-C1 | TR_C_gene | 145,5761784 | 1,103489975 | 0,34240512 | 3,222761316 | 0,001269613 | 0,034386241 |
| Oas2 | protein_coding | 255,2870063 | 1,112773913 | 0,323603433 | 3,438696251 | 5,85E-04 | 0,019935626 |
| Mvd | protein_coding | 901,8098552 | 1,117578613 | 0,168136667 | 6,646846485 | 2,99E-11 | 8,58E-09 |
| Akap1 | protein_coding | 63,16080001 | 1,118802832 | 0,360442484 | 3,103970487 | 0,001909424 | 0,045084782 |
| Ptgir | protein_coding | 299,3926274 | 1,119335046 | 0,270767312 | 4,133937138 | 3,57E-05 | 0,002178816 |
| Pcsk4 | protein_coding | 418,7493132 | 1,125917233 | 0,20684158 | 5,44337958 | 5,23E-08 | 8,02E-06 |
| Gm37169 | TEC | 89,89323941 | 1,126613301 | 0,335840273 | 3,354610489 | 7,95E-04 | 0,024630168 |
| Neb | protein_coding | 620,0738054 | 1,133364424 | 0,265411276 | 4,270219571 | 1,95E-05 | 0,001383273 |
| Sell | protein_coding | 21628,92501 | 1,135752051 | 0,242309835 | 4,687189238 | 2,77E-06 | 2,72E-04 |
| Fdft1 | protein_coding | 1198,615738 | 1,144742517 | 0,119988388 | 9,540444175 | 1,42E-21 | 2,29E-18 |
| C130036L24Rik | processed_transcript | 73,78636976 | 1,146207542 | 0,360417001 | 3,180226072 | 0,001471602 | 0,038404641 |
| Myc | protein_coding | 4741,605711 | 1,149991013 | 0,161850945 | 7,105247421 | 1,20E-12 | 4,42E-10 |
| A930029G22Rik | lincRNA | 60,79251133 | 1,158519645 | 0,35606632 | 3,253662534 | 0,001139275 | 0,031929429 |
| Msmo1 | protein_coding | 1471,480027 | 1,16390529 | 0,109370863 | 10,64182227 | 1,90E-26 | 4,09E-23 |
| Il1rl1 | protein_coding | 448,7726476 | 1,170929299 | 0,332902678 | 3,517332166 | 4,36E-04 | 0,016275323 |
| AA465934 | processed_transcript | 61,88746149 | 1,182676064 | 0,345464223 | 3,423440071 | 6,18E-04 | 0,020705518 |
| Gm14125 | transcribed_processe | 377,2562026 | 1,214811 | 0,33778388 | 3,59641496 | 3,23E-04 | 0,012837607 |
| Rad51b | protein_coding | 153,7109888 | 1,223093481 | 0,257408436 | 4,751567201 | 2,02E-06 | 2,13E-04 |
| Fn3krp | protein_coding | 92,79758416 | 1,231608673 | 0,352290563 | 3,496002457 | 4,72E-04 | 0,017150194 |
| Pik3c2b | protein_coding | 61,40937672 | 1,248466689 | 0,399608795 | 3,124222248 | 0,001782757 | 0,04343521 |
| Mvk | protein_coding | 311,8212075 | 1,262737604 | 0,209328139 | 6,032335684 | 1,62E-09 | 3,31E-07 |
| Mx2 | polymorphic_pseudo | 183,9959249 | 1,263182581 | 0,335740187 | 3,762381236 | 1,68E-04 | 0,007804911 |
| Tmem97 | protein_coding | 213,2738006 | 1,266161604 | 0,246318015 | 5,140353229 | 2,74E-07 | 3,75E-05 |
| Gm42646 | TEC | 265,4281468 | 1,275496073 | 0,169600645 | 7,520585036 | 5,45E-14 | 2,81E-11 |
| Nrip1 | protein_coding | 930,0552526 | 1,289838364 | 0,173343359 | 7,440944773 | 1,00E-13 | 4,96E-11 |
| Gm6781 | processed_pseudoge | 105,6388825 | 1,293821167 | 0,274737115 | 4,709306084 | 2,49E-06 | 2,50E-04 |
| Tent5a | protein_coding | 556,3001342 | 1,296552398 | 0,169375021 | 7,654920936 | 1,93E-14 | 1,13E-11 |
| C230085N15Rik | TEC | 875,9632638 | 1,298562289 | 0,225336593 | 5,762767022 | 8,27E-09 | 1,48E-06 |
| Snord104 | snoRNA | 74,44933199 | 1,299076866 | 0,34156165 | 3,803345212 | 1,43E-04 | 0,006816297 |
| Firre | processed_transcript | 94,6967127 | 1,302278686 | 0,391596194 | 3,325565228 | 8,82E-04 | 0,026455419 |
| Aldoc | protein_coding | 247,8412473 | 1,320249824 | 0,235995873 | 5,594376735 | 2,21E-08 | 3,61E-06 |
| Arglu1 | protein_coding | 753,2586698 | 1,332657023 | 0,176431524 | 7,55339518 | 4,24E-14 | 2,34E-11 |
| Gm35570 | processed_pseudoge | 36,58693876 | 1,362363216 | 0,41203973 | 3,306387996 | 9,45E-04 | 0,027865159 |
| Hmgcs1 | protein_coding | 1281,789717 | 1,369508846 | 0,193344835 | 7,08324506 | 1,41E-12 | 5,04E-10 |
| Acaca | protein_coding | 240,075092 | 1,379142527 | 0,288344138 | 4,78297404 | 1,73E-06 | 1,89E-04 |
| AC121821.1 | processed_transcript | 83,19894988 | 1,394998358 | 0,302342742 | 4,613963435 | 3,95E-06 | 3,72E-04 |
| Bcl6 | protein_coding | 574,7899265 | 1,397988089 | 0,271517767 | 5,148790463 | 2,62E-07 | 3,63E-05 |
| Tbl1xr1 | protein_coding | 1596,922979 | 1,403513972 | 0,166090152 | 8,450314186 | 2,91E-17 | 2,20E-14 |
| 9430081H08Rik | TEC | 40,05866498 | 1,424873825 | 0,412648542 | 3,452996143 | 5,54E-04 | 0,019369337 |
| Gm3571 | processed_pseudoge | 142,9810726 | 1,434258668 | 0,222038223 | 6,459512451 | 1,05E-10 | 2,66E-08 |
| Cd86 | protein_coding | 1123,528334 | 1,450619857 | 0,155771465 | 9,312487743 | 1,25E-20 | 1,46E-17 |
| Insig1 | protein_coding | 1153,804771 | 1,45877234 | 0,15796533 | 9,234762706 | 2,59E-20 | 2,57E-17 |
| Acsl3 | protein_coding | 354,7933204 | 1,472077117 | 0,22498773 | 6,542921782 | 6,03E-11 | 1,59E-08 |
| Fdps | protein_coding | 128,1040338 | 1,480537831 | 0,243027241 | 6,09206533 | 1,11E-09 | 2,39E-07 |
| Fhl3 | protein_coding | 151,2087501 | 1,500861692 | 0,268764755 | 5,584295051 | 2,35E-08 | 3,78E-06 |
| B430306N03Rik | protein_coding | 109,2562536 | 1,522327614 | 0,394297743 | 3,860858051 | 1,13E-04 | 0,005712396 |
| Gm12940 | processed_transcript | 287,0142315 | 1,53424184 | 0,21898748 | 7,006071033 | 2,45E-12 | 8,10E-10 |
| Cyp51 | protein_coding | 1792,742796 | 1,593754367 | 0,167824471 | 9,496555284 | 2,17E-21 | 3,11E-18 |
| Plac8 | protein_coding | 94,17942058 | 1,621428707 | 0,51782648 | 3,131220144 | 0,001740816 | 0,043074425 |
| Nlgn2 | protein_coding | 40,48897085 | 1,63365178 | 0,479143022 | 3,409528479 | 6,51E-04 | 0,0214018 |
| Gm38034 | TEC | 37,08882945 | 1,637520824 | 0,509793695 | 3,212124514 | 0,001317573 | 0,035387803 |
| E430014B02Rik | TEC | 160,1193444 | 1,683065644 | 0,384131889 | 4,381478585 | 1,18E-05 | 9,15E-04 |
| Ldlr | protein_coding | 1833,63119 | 1,717086898 | 0,18540691 | 9,261180702 | 2,02E-20 | 2,17E-17 |
| Gm48236 | lincRNA | 27,79908876 | 1,728868691 | 0,501677993 | 3,446172079 | 5,69E-04 | 0,019714359 |
| Gm20696 | protein_coding | 511,0748106 | 1,738269545 | 0,269961689 | 6,438948988 | 1,20E-10 | 2,93E-08 |
| Gm16086 | lincRNA | 28,49137106 | 1,778665364 | 0,573732826 | 3,100163149 | 0,001934141 | 0,045584904 |
| Gm13546 | antisense | 149,8600705 | 1,790028362 | 0,31347861 | 5,710208949 | 1,13E-08 | 1,97E-06 |
| Zfp608 | protein_coding | 78,79247855 | 1,790505843 | 0,371066086 | 4,825301775 | 1,40E-06 | 1,62E-04 |
| Osgin1 | protein_coding | 604,3913437 | 1,793351511 | 0,192141375 | 9,333499936 | 1,02E-20 | 1,32E-17 |
| Ky | protein_coding | 77,17677314 | 1,828003453 | 0,386368755 | 4,731240377 | 2,23E-06 | 2,30E-04 |
| Gm32633 | lincRNA | 143,4688512 | 1,851905196 | 0,352176937 | 5,258451084 | 1,45E-07 | 2,13E-05 |
| Lztfl1 | protein_coding | 383,1130139 | 1,918007626 | 0,249667503 | 7,682247795 | 1,56E-14 | 9,60E-12 |
| Gm6776 | processed_pseudoge | 33,46587875 | 1,922663505 | 0,438768446 | 4,381954815 | 1,18E-05 | 9,15E-04 |
| Gm37248 | TEC | 182,0228201 | 1,954836447 | 0,326168561 | 5,993331916 | 2,06E-09 | 4,14E-07 |
| Dhcr24 | protein_coding | 87,37391843 | 1,96293428 | 0,316446081 | 6,203060797 | 5,54E-10 | 1,32E-07 |
| Sqle | protein_coding | 1405,869022 | 2,003189648 | 0,128668039 | 15,56866536 | 1,19E-54 | 7,66E-51 |

| | | | | | | | |
|---|---|---|---|---|---|---|---|
| Gm13502 | processed_pseudoge | 198,5034244 | 2,028222405 | 0,242495698 | 8,363952104 | 6,07E-17 | 4,12E-14 |
| Fgd2 | protein_coding | 31,72874786 | 2,054897245 | 0,654750185 | 3,138444696 | 0,00169847 | 0,042353333 |
| Idi1 | protein_coding | 624,2997424 | 2,085314713 | 0,188861089 | 11,04152645 | 2,41E-28 | 6,21E-25 |
| Gm19585 | lincRNA | 770,6443714 | 2,105025261 | 0,157200105 | 13,39073696 | 6,85E-41 | 2,94E-37 |
| Pde3b | protein_coding | 73,68132456 | 2,124076986 | 0,51456894 | 4,127876403 | 3,66E-05 | 0,002226478 |
| Scd2 | protein_coding | 2778,589368 | 2,139465026 | 0,131344716 | 16,28893113 | 1,18E-59 | 1,52E-55 |
| Ncf1 | protein_coding | 450,3116181 | 2,260495748 | 0,268901533 | 8,406407074 | 4,23E-17 | 3,03E-14 |
| Tmem273 | protein_coding | 139,9597381 | 2,306416043 | 0,329202449 | 7,006071941 | 2,45E-12 | 8,10E-10 |
| Gm43672 | lincRNA | 248,0424019 | 2,312833322 | 0,22393918 | 10,327953 | 5,27E-25 | 9,70E-22 |
| A430072P03Rik | processed_transcript | 30,23856443 | 2,351229392 | 0,525307058 | 4,475914336 | 7,61E-06 | 6,45E-04 |
| Scd1 | protein_coding | 37,09793132 | 2,361821406 | 0,522534834 | 4,519931022 | 6,19E-06 | 5,50E-04 |
| Stra6 | protein_coding | 204,3169904 | 2,390015751 | 0,347924542 | 6,869350859 | 6,45E-12 | 1,89E-09 |
| Ccr9 | protein_coding | 413,0109001 | 2,40691358 | 0,208694044 | 11,53321642 | 8,97E-31 | 2,89E-27 |
| Grifin | protein_coding | 59,14771579 | 2,444256885 | 0,6443076 | 3,793617963 | 1,48E-04 | 0,007036949 |
| Sorcs2 | protein_coding | 285,8076518 | 2,639850671 | 0,469392716 | 5,623970249 | 1,87E-08 | 3,08E-06 |
| Bpifa2 | protein_coding | 62,31426255 | 2,894152391 | 0,926289324 | 3,124458327 | 0,001781328 | 0,04343521 |
| Scd4 | protein_coding | 35,99247455 | 2,943182178 | 0,765698474 | 3,843787441 | 1,21E-04 | 0,005984158 |
| A930002I21Rik | lincRNA | 71,48133801 | 3,255285172 | 0,472507031 | 6,889389911 | 5,60E-12 | 1,72E-09 |
| Tuba8 | protein_coding | 34,3638834 | 3,312270145 | 0,79963665 | 4,142219025 | 3,44E-05 | 0,002111597 |
| Cyp11a1 | protein_coding | 25,44328736 | 3,526835899 | 0,789864613 | 4,465114454 | 8,00E-06 | 6,70E-04 |
| Cyp26b1 | protein_coding | 41,19204133 | 3,738299206 | 0,861944301 | 4,337054263 | 1,44E-05 | 0,001077004 |
| Epas1 | protein_coding | 127,447874 | 4,378970013 | 0,501332992 | 8,734653573 | 2,44E-18 | 1,97E-15 |
| 1700061F12Rik | lincRNA | 40,97279913 | 5,439944296 | 0,737676442 | 7,374431372 | 1,65E-13 | 7,88E-11 |
| Xist | lincRNA | 719,0420469 | 9,855136578 | 1,085533388 | 9,078612122 | 1,10E-19 | 1,01E-16 |

**Supplementary Table 3. GSEA hallmark gene sets in naive Treg cells**

| pathway | pval | padj | log2err | ES | NES | size | leadingEdge |
|---|---|---|---|---|---|---|---|
| HALLMARK_G2M_CHECKPOINT | 3,2448E-15 | 1,1965E-13 | 1,00731796 | 0,64733292 | 2,44085637 | 181 | Pbk/Hmmr/Ccnb2/Bub1/Ccna2/Ttk/Kif2c/Aurkb/Cdkn3/Nek2/Tpx2/Cdk1/Kif11/Cenpf/Espl1/Plk1/Exo1/Cdc6/Chek1/Mki67/Cenpe/Cks1b/Rad54l/Marcks/Aurka/Kif4/Kif22/Ccnf/Top2a/Gins2/Racgap1/Mybl2/Cks2/Kif15/Stmn1/Cdc20/Sqle/Kif23/Bard1/Tacc3/Knl1/Kmt5a/Slc7a5/Atf5/Traip/Pole/Ccnd1/Cdc45/Tfdp1/Kif20b/Pura/Prim2/Polq/Orc6/Brca2/Cdkn1b/Kpna2/Myc/Chaf1a/Dbf4/Kpnb1/Cul3/Hnrnpd/Cenpa/Pola2/Odc1/Rps6ka5 |
| HALLMARK_E2F_TARGETS | 4,7859E-15 | 1,1965E-13 | 0,99698622 | 0,63141945 | 2,41454681 | 194 | E2f8/Hmmr/Ccnb2/Depdc1a/Mxd3/Kif2c/Aurkb/Cdkn3/Cdk1/Kif18b/Spc24/Espl1/Plk1/Chek1/Mki67/Cdca3/Cit/Cenpe/Cdca8/Cks1b/Rad51ap1/Aurka/Kif4/Kif22/Spc25/Top2a/Spag5/Cenpm/Racgap1/Mybl2/Tk1/Cks2/Stmn1/Chek2/Cdc20/Dlgap5/Asf1b/Bard1/Tacc3/Cdkn1a/Hells/Dctpp1/Ung/Pole/Diaph3/Gins1/Ube2t/Wdr90/Prim2/Mms22l/Orc6/Brca2/Cdkn1b/Kpna2/Shmt1/Myc |
| HALLMARK_MITOTIC_SPINDLE | 6,4349E-06 | 0,00010725 | 0,61052688 | 0,48885812 | 1,84334145 | 183 | Ccnb2/Bub1/Ttk/Kif2c/Nek2/Tpx2/Cdk1/Kif11/Cenpf/Kntc1/Espl1/Plk1/Cenpe/Anln/Marcks/Aurka/Kif4/Kif22/Top2a/Ect2/Pif1/Fgd6/Racgap1/Kif15/Trio/Dlgap5/Kif23/Vcl/Arhgap29/Dynll2/Rasa1/Bcar1/Kif20b/Mid1ip1/Sass6/Mid1/Brca2/Arhgef11/Arf6/Rictor/Sos1/Tubd1/Cttn/Bcr/Sac3d1 |
| HALLMARK_SPERMATOGENESIS | 0,00010302 | 0,00128777 | 0,5384341 | 0,56192923 | 1,86991494 | 76 | Ccnb2/Bub1/Mlf1/Ttk/Kif2c/Cdkn3/Nek2/Cdk1/Ncaph/Aurka/Gfi1/Ace/Nphp1/Tnp2/Pcsk4 |
| HALLMARK_EPITHELIAL_MESENCHYMAL_TRANSITION | 0,00088436 | 0,00884356 | 0,47727082 | -0,517461 | -1,7210373 | 90 | Fn1/Tgfbi/Col1a1/Mmp14/Nid2/Lrp1/Col1a2/Adam12/Snai2/Vcam1/Areg/Capg/Jun/Ecm1/Comp/Gem/Fgf2/Serpine2 |
| HALLMARK_COMPLEMENT | 0,00113977 | 0,00949805 | 0,45505987 | -0,4648936 | -1,6349024 | 134 | Fn1/Pla2g7/Mmp14/C3/Lrp1/Gzmk/Src/Ang/Serpinc1/Plek/Gca/Pdgfb/Cpq/F8/Gnb4/Pfn1/Anxa5/Irf1/Ctss/Plscr1/Lck/Cxcl2/Calm1/Dpp4/Ctsd/Pdp1/Lyn/C1qa/Pim1/Prcp/Lcp2/Lta4h/Gnai3/Psen1/Fcer1g |
| HALLMARK_IL2_STAT5_SIGNALING | 0,00153588 | 0,01097055 | 0,45505987 | -0,4169075 | -1,5347039 | 185 | Serpinc1/Capg/Gpr83/Itgae/Irf6/Gbp3/Cdkn1c/Myo1e/Muc1/Ecm1/Penk/Il2rb/Ltb/Ikzf2/Cdcp1/Tnfrsf9/Ndrg1/Nrp1/Plpp1/Cst7/Cd81/Ctla4/Socs1/Tnfrsf8/Fah/Cd79b/Swap70/Tnfrsf18/Irf8/Sell/Tiam1/Csf1/Smpdl3a/Drc1/Il2ra/Galm/Gpx4/Slc2a3/Plscr1/Cd83/Rhoh/Capn3/Rgs16/Tnfrsf4/Il4ra/Cdc42se2/Gata1/Ccnd2/Traf1/Cish/Rnh1/Slc39a8/P4ha1/Wls/Il3ra/Pim1/Nt5e/Socs2/Etfbkmt/Tlr7/Sh3bgrl2/Enpp1/Nfil3/Serpinb6a/Ikzf4/Irf4/Xbp1/Gabarapl1/Tnfsf10/Syt11/Prkch/Tnfrsf1b/Eef1akmt1 |
| HALLMARK_MYC_TARGETS_V2 | 0,00529644 | 0,03062424 | 0,40701792 | 0,53358825 | 1,70108298 | 57 | Plk1/Tmem97/Hk2/Slc29a2/Utp20/Dctpp1/Ung/Ndufaf4/Myc/Srm/Slc19a1/Gnl3/Aimp2/Pprc1/Rrp12/Supv3l1/Tfb2m/Pa2g4/Nop16/Pus1/Nolc1/Ipo4/Rcl1/Rrp9/Bysl/Hspd1/Tbrg4/Mybbp1a/Grwd1/Nop56/Cbx3/Hspe1/Nip7/Rabepk/Ppan |
| HALLMARK_COAGULATION | 0,00551236 | 0,03062424 | 0,40701792 | -0,5351791 | -1,6971995 | 67 | Rapgef3/Fn1/Mmp14/C3/Lrp1/Ang/Serpinc1/Plek/Pdgfb/Comp/Cpq/F8/Crip2 |
| HALLMARK_ESTROGEN_RESPONSE_LATE | 0,00677575 | 0,03079888 | 0,40701792 | 0,41158546 | 1,48286916 | 127 | Cyp26b1/Ltf/Slc1a4/Cdc6/Papss2/Ckb/Top2a/Gins2/Id2/Celsr2/Tst/Cdc20/Rapgefl1/Nrip1/Scarb1/Xrcc3/Lsr/Car12/Egr3/Slc7a5/Anxa9/Gla/Abhd2/Cacna2d2/Ccnd1/Ret/Jak2/Fdft1/Clic3/Lamc2/Mest/Sult2b1/Gale/Igfbp4/Kif20a/Fkbp5/Impa2/Farp1/Tmprss3/Nbl1/Il17rb/Rbbp8/Slc16a1/Slc2a8/Wfs1/Chpt1 |
| HALLMARK_APICAL_JUNCTION | 0,00658034 | 0,03079888 | 0,40701792 | -0,4409322 | -1,5318878 | 121 | Nectin2/Tgfbi/Actg2/Lima1/Icam5/Cd209b/Vcam1/Src/Cx3cl1/Sirpa/Acta1/Gamt/Itga9/Actg1/Actb/Pfn1/Cnn2/Rac2/Tmem8b/Pik3r3/Baiap2/Sorbs3/Msn/Cercam/Col16a1 |
| HALLMARK_MTORC1_SIGNALING | 0,00841302 | 0,03505427 | 0,3807304 | 0,37305186 | 1,41491153 | 187 | Bub1/Plk1/Slc1a4/Asns/Scd2/Aurka/Plod2/Ccnf/Bcat1/Bhlhe40/Idi1/Sqle/Tmem97/Hk2/Tm7sf2/Cdkn1a/Scd1/Insig1/Ldlr/Acaca/Slc7a5/Dhcr24/Ung/Hmgcs1/Cyp51/Gla/Acsl3/Fads1/Stard4/Tpi1/Cxcr4/Psme3/Ifi30/Hmgcr/Elovl6/Pno1/Etf1/Sdf2l1/Cfp/Glrx/Fkbp2/Sytl2/Tfrc/Nmt1/Srd5a1/Nup205 |

| | | | | | | | |
|---|---|---|---|---|---|---|---|
| HALLMARK_INTERFERON_GAMMA_RESPONSE | 0,0134435 | 0,05170578 | 0,3807304 | -0,393517 | -1,4268347 | 170 | St3gal5/Vcam1/Gbp3/H2-Q7/H2-D1/Irf5/Cd38/Il2rb/B2m/Txnip/Gm8909/Ly6e/H2-Q10/Socs1/Slamf7/Samhd1/Irf8/Tapbp/Peli1/Csf2rb2/Psme1/Irf1/Gbp10/Psmb8/Cd74/Stat1/Plscr1/Itgb7/Cmpk2/Psmb10/Nfkbia/Gch1/Il4ra/Tap1/Gbp9/Eif4e3/Pim1/Ube2l6/H2-Aa/Lcp2/Nod1/Ifi35/Irf4/Btg1/Tnfsf10/Zbp1/Irf9/Psme2/Lats2/Icam1/Gbp8/Casp8/Bpgm/Psma2/Nmi/Il15ra/Gbp4/Pfkp/Hif1a/Nampt/Il10ra/Psmb2/Mvp/Stat3/Rnf31/Ptpn6 |
| HALLMARK_TNFA_SIGNALING_VIA_NFKB | 0,01974715 | 0,07052554 | 0,35248786 | -0,3970316 | -1,4283754 | 158 | Gfpt2/Dram1/Ccl4/Fut4/Areg/Cd80/Plek/Fosb/Ccl20/Hbegf/Jun/Ccrl2/Gem/Tnfrsf9/Sdc4/Fosl2/Csf1/Irf1/Litaf/Slc2a3/Cd83/Slc2a6/Stat5a/Nfkbia/Gch1/Dusp4/Dusp2/Btg2/Cxcl2/Tap1/Eif1/Btg3/Traf1/Gadd45a/Nr4a2/Pnrc1/Rnf19b/Sqstm1/Relb/Nfil3/Btg1/Sat1/Rela/Dusp1/Tubb2a/Icam1/Ptpre/Plk2/Egr2/Klf10/Ppp1r15a/Phlda1/Clcf1/Il15ra/Pfkfb3/Klf9/Mcl1/Nampt |
| HALLMARK_CHOLESTEROL_HOMEOSTASIS | 0,02224345 | 0,07414485 | 0,35248786 | 0,4720962 | 1,50204392 | 59 | Scd2/Aldoc/Idi1/Lss/Sqle/Tmem97/Mvk/Hsd17b7/Tm7sf2/Scd1/Ldlr/Fdps/Atf5/Hmgcs1/Cyp51/Mvd/Fdft1/Stard4/Nsdhl/Atxn2/Hmgcr/Pmvk/Gstm7 |
| HALLMARK_UV_RESPONSE_UP | 0,02573456 | 0,07568988 | 0,35248786 | -0,3979915 | -1,4055567 | 130 | Mmp14/Nptxr/H2-Q1/Mapk8ip2/H2-Q2/Cdo1/Plcl1/Fosb/Cdkn1c/Cdkn2b/Epcam/Maoa/Irf1/Cnp/Ephx1/Tmbim6/Chka/Nfkbia/Gch1/Btg2/Tap1/Btg3/Lyn/Tuba4a/Sqstm1/Selenow/Dnaja1/Btg1/Bid |
| HALLMARK_APICAL_SURFACE | 0,0251238 | 0,07568988 | 0,35248786 | -0,6329011 | -1,6278494 | 23 | Gstm5/Cx3cl1/Akap7/Cd160/Il2rb/Tmem8b/Sulf2/Crocc/Lyn/Il2rg |
| HALLMARK_KRAS_SIGNALING_UP | 0,0483871 | 0,1344086 | 0,2765006 | -0,3867612 | -1,3474984 | 124 | Aldh1a2/Gfpt2/Clec4a3/Evi5/Il1rl2/Kif5c/Adgra2/Ccl20/Scg5/Hbegf/F13a1/Nrp1/Mtmr10/Irf8/Tnnt2/Map7/Ctss/Psmb8/Dnmbp/Map3k1/Laptm5/Hdac9/Rgs16/Mafb/Tmem158/Itgb2/Ccnd2/Mmp11/Traf1/Ets1/Vwa5a/Mmd/Avl9/Lcp1 |
| HALLMARK_ESTROGEN_RESPONSE_EARLY | 0,09438202 | 0,24837374 | 0,21925035 | 0,34946897 | 1,26119636 | 129 | Cyp26b1/Mybl1/Krt18/Slc1a4/Papss2/Bhlhe40/Celsr2/Hes1/Olfml3/Akap1/Rapgefl1/Nrip1/Scarb1/Deptor/Car12/Egr3/Slc7a5/Anxa9/Gla/Abhd2/Ccnd1/Ret/Jak2/Fdft1/Clic3/Nav2/Ugcg/Sult2b1/Igfbp4/Myc/Fkbp5/Farp1/Slc19a2/Tmprss3/Nbl1/Il17rb/Rbbp8/Tgm2/Celsr1/Slc16a1/Wfs1/Chpt1/Pex11a |
| HALLMARK_INFLAMMATORY_RESPONSE | 0,16370107 | 0,40925267 | 0,14375899 | -0,3364205 | -1,1841383 | 127 | Fzd5/Mmp14/Itgb8/Nmur1/Cx3cl1/Axl/Ccl20/Best1/Hbegf/Il2rb/Ccrl2/Nod2/Tnfrsf9/Ptafr/Ly6e/Tapbp/Sell/Csf1/Irf1/Rgs1/Lck/Bdkrb1/Rgs16/Nfkbia/Gch1/Itga5/Btg2/Acvr2a/Il4ra/Emp3/Lyn/Lta/Lcp2/Calcrl/Cd82/Gnai3/Slamf1/Psen1/Tnfsf10/Rela/Tnfrsf1b/Icam1/Ptpre/Slc28a2 |
| HALLMARK_ALLOGRAFT_REJECTION | 0,17482517 | 0,41625042 | 0,13725078 | -0,317432 | -1,1487144 | 163 | Ccl4/Cd8a/Cd80/Capg/H2-Q7/Igsf6/Il2rb/Ltb/Gbp2/B2m/Ly86/Prf1/Cd96/H2-Q10/H2-T23/Socs1/Irf8/Cd2/Tapbp/Rps19/Csf1/Ctss/H2-Ob/Il2ra/Cd7/Spi1/Cd74/Stat1/Srgn/Hdac9/Lck/Psmb10/Il16/Acvr2a/Il4ra/Rps9/Itgb2/Tap1/H2-Oa/Cd1d1/Ccnd2/Eif5a/Ets1/Cd3g/Lyn/Flna/H2-Aa/Lcp2/Cd3d/Ptprc/Il2rg/Irf4/Hcls1/Tap2/Cd3e/Icam1/Prkcb/Rps3a1/Sit1 |
| HALLMARK_WNT_BETA_CATENIN_SIGNALING | 0,20086393 | 0,42397661 | 0,14290115 | 0,43725582 | 1,21423011 | 29 | Csnk1e/Ppard/Notch4/Myc/Hey1/Skp2/Maml1/Axin2/Hdac5/Axin1/Hdac2/Rbpj/Ncor2 |
| HALLMARK_P53_PATHWAY | 0,20350877 | 0,42397661 | 0,12625399 | -0,3108884 | -1,1294584 | 167 | Dram1/St14/Cdkn2b/Cebpa/Hbegf/Jun/Txnip/Ndrg1/Slc35d1/Cd81/Socs1/Ldhb/Hspa4l/Prmt2/Ephx1/Dgka/Baiap2/Rpl18/Rgs16/Btg2/Krt17/Tap1/Ccnd2/Slc7a11/S100a10/Ctsd/Steap3/Vwa5a/Gadd45a/Ccng1/Trib3/Rnf19b/Cd82/Def6/Rpl36/Btg1/Sat1/Zfp36l1/Ptpre/Perp/Plk2/Gm2a |
| HALLMARK_NOTCH_SIGNALING | 0,19544592 | 0,42397661 | 0,13500203 | -0,4705564 | -1,2102916 | 23 | Fzd5/Tcf7l2 |
| HALLMARK_INTERFERON_ALPHA_RESPONSE | 0,25577265 | 0,5084394 | 0,11146267 | -0,3389651 | -1,1162518 | 87 | Gbp3/H2-Q7/Ccrl2/Gbp2/B2m/Txnip/Ly6e/H2-Q10/Psme1/Sell/Csf1/Irf1/Cnp/Psmb8/Cd74/Mov10/Plscr1/Cmpk2/Il4ra/Tap1 |
| HALLMARK_KRAS_SIGNALING_DN | 0,26438849 | 0,5084394 | 0,11012226 | -0,3508034 | -1,1242357 | 73 | Tex15/Zfp112/Cd80/Brdt/Hc/Stag3/Gamt/Synpo/Tcf7l1/Selenop/Btg2/Cpeb3/Fgf16/Cyp39a1/Arpp21/Pdcd1/Nr4a2 |
| HALLMARK_ANDROGEN_RESPONSE | 0,29748284 | 0,53121935 | 0,11828753 | 0,31960541 | 1,09261838 | 86 | Klk1b27/Scd2/Idi1/Cenpn/Scd1/Gucy1a1/Insig1/Dhcr24/Hmgcs1/Srf/Abhd2/Acsl3/Ccnd1/Fads1/Maf/Akt1/Sec24d/Hmgcr/Pmepa1/Fkbp5 |
| HALLMARK_ANGIOGENESIS | 0,29643527 | 0,53121935 | 0,10552094 | -0,5226342 | -1,1700714 | 12 | Slco2a1/Nrp1/Pglyrp1/Ccnd2 |

| | | | | | | | |
|---|---|---|---|---|---|---|---|
| HALLMARK_MYOGENESIS | 0,39788732 | 0,66314554 | 0,08455574 | -0,3031236 | -1,026084 | 100 | Col1a1/Mef2c/Itga7/Adam12/Pgam2/Dapk2/Hbegf/Fgf2/Acta1/Sorbs1/Cnn3/Dmd/Tnnt2/Erbb3/Lsp1/Sorbs3/Tpm3 |
| HALLMARK_UV_RESPONSE_DN | 0,38908451 | 0,66314554 | 0,08578444 | -0,3041273 | -1,0284459 | 99 | Col1a1/Col1a2/Snai2/Plcb4/Cdc42bpa/Nrp1/Id1/Fhl2/Prkar2b/Pik3r3/Acvr2a/Pparg/Kalrn/Nr1d2 |
| HALLMARK_PEROXISOME | 0,45535714 | 0,68220429 | 0,09026355 | 0,29491777 | 0,99174944 | 79 | Top2a/Idi1/Abcb9/Fdps/Hsd17b11/Dhcr24/Fads1/Dlg4/Sult2b1/Ctps/Pex11a/Acsl1/Retsat/Elovl5/Acsl4/Abcb4/Sod2/Ctbp1/Pex2/Gnpat/Slc25a4/Idh1/Idh2/Abcb1a/Mlycd/Smarcc1/Eci2 |
| HALLMARK_GLYCOLYSIS | 0,45977011 | 0,68220429 | 0,09139243 | 0,26989847 | 0,98566233 | 145 | Hmmr/Depdc1a/Cdk1/Ppfia4/Sdc1/Chst2/Aurka/Plod2/Ppp2cb/Stmn1/Hk2 |
| HALLMARK_IL6_JAK_STAT3_SIGNALING | 0,46389892 | 0,68220429 | 0,07767986 | -0,3189537 | -0,9966755 | 65 | Il13ra1/Jun/Cd38/Ltb/Socs1/Csf2rb2/Il9r/Csf1/Irf1/Il2ra/Stat1/Il4ra/Cxcl2/Il3ra/Pim1/Il17ra/Il2rg/Cd9/Tnfrsf1b/Irf9/Itga4/Il10rb/Il15ra |
| HALLMARK_HYPOXIA | 0,42490842 | 0,68220429 | 0,08312913 | -0,2914478 | -1,0228667 | 133 | Tgfbi/Kif5a/Pdgfb/Cdkn1c/Pgam2/Jun/Csrp2/Ndrg1/Sdc4/Fosl2/Cavin1/Eno1/Map3k1/Slc2a3/Tktl1/Ldha/Ets1/P4ha1/Grhpr/Pim1/Pnrc1/Gapdh/Nfil3/Bnip3l/Akap12/Btg1/Lxn/Hk1/Dusp1/Prkca/Sdc3/Slc2a1/Ilvbl/Ext1/Ppp1r15a/Casp6/Pfkfb3/Pfkp/Rragd/Aldoa/Pfkl/Wsb1/Rora |
| HALLMARK_HEDGEHOG_SIGNALING | 0,53653846 | 0,76648352 | 0,07343814 | -0,3830106 | -0,9694938 | 22 | Nrp2/Adgrg1/Nrp1 |
| HALLMARK_BILE_ACID_METABOLISM | 0,56183746 | 0,7803298 | 0,06689663 | -0,2931626 | -0,9341557 | 68 | Nr3c2/Efhc1/Abca5/Optn |
| HALLMARK_ADIPOGENESIS | 0,59219858 | 0,80026835 | 0,06450312 | -0,2599867 | -0,9450527 | 172 | C3/Itga7/Ptger3/Sowahc/Ubc/Sorbs1/Ucp2/Fah/Atp1b3/Cavin1/Sqor/Gpx4/Pqlc3/Chchd10/Stat5a/Pparg/Gadd45a/Ndufa5/Gpam/Cd151/Crat/Itsn1/Reep5/Ndufb7/Uqcrc1/Scp2/Aldh2/Aplp2/Pfkfb3/Mgst3/Phyh/Aldoa/Pfkl/Atp5o/Apoe/Nabp1/Ghitm/Acads/Angptl4/Ifngr1/Sod1/Samm50/Slc1a5/Esrra/Acox1/Immt/Cox6a1/Ech1/Cox8a/Esyt1 |
| HALLMARK_HEME_METABOLISM | 0,61403509 | 0,8079409 | 0,06224904 | -0,2597127 | -0,9324378 | 157 | C3/Hbb-bt/Xk/Endod1/Igsf3/Pdzk1ip1/Slc30a1/Optn/Slc4a1/Tns1/Ucp2/E2f2/Cast/Abcg2/Sptb/Ypel5/Btg2/Atp6v0a1/Gata1/Alas2/Slc7a11 |
| HALLMARK_APOPTOSIS | 0,69594595 | 0,89223839 | 0,06831109 | 0,24617038 | 0,88818994 | 131 | Emp1/Krt18/Anxa1/Top2a/Lmna/Gna15/Plat/Fasl/Cdkn1a/Gucy2e/Egr3/Ptk2/Bik/Bmf/Ifitm3/Ccnd1/Nefh/Cdkn1b/Casp4 |
| HALLMARK_OXIDATIVE_PHOSPHORYLATION | 0,74099485 | 0,92624357 | 0,05237591 | -0,2426719 | -0,8966625 | 194 | Ldhb/Cox7a2l/Gpx4/Nqo2/Atp5c1/Atp5b/Ldha/Pdp1/Oat/Ndufa5/Atp5a1/Cyb5a/Cox4i1/Uqcrh/Nnt/Ndufb2/Cox6b1/Sdha/Ndufb7/Atp6v0c/Hadha/Uqcrc1/Atp6v1f/Ndufs2/Ndufa7/Slc25a5/Bax/Rhot1/Mgst3/Tomm22/Phyh/Atp5o/Atp1b1/Slc25a3/Pdha1/Atp6v1d/Glud1/Atp6v1g1/Casp7/Hccs/Isca1/Hadhb/Gpi1/Atp6v0e/Immt/Atp5d/Mdh1/Cox6a1/Atp5h/Uqcrc2/Ech1/Cox8a/Cox6c/Atp5j/Uqcrfs1/Ndufs3/Atp5l/Cyc1/Pdhb/Ndufs7/Atp6v1e1/Slc25a11/Cycs/Mpc1/Mdh2/Atp6ap1/Ogdh/Iscu |
| HALLMARK_MYC_TARGETS_V1 | 0,93571429 | 1 | 0,05676724 | 0,21037515 | 0,80648245 | 200 | Ccna2/Cdc20/Tyms/Cdc45/Eif3b/Tfdp1/Rfc4/Kpna2/Myc/Fam120a/Kpnb1/Trim28/Dut/Hnrnpd/Srm/Etf1/Odc1/Gnl3/Ctps/Ap3s1/Aimp2/Gspt1/Hnrnpu/Ddx21/Mad2l1/Hdgf/Ndufab1/Pole3/Pcbp1/Rad23b/Pa2g4/Nop16/Tomm70a/Rps10/Nolc1/Ifrd1/Srsf1/Hdac2/Snrpd1/Nap1l1/Ssbp1/Rrp9/Prdx3/Hspd1/Cad/C1qbp/Apex1/Usp1/Set/Syncrip/Txnl4a/Pabpc4/Tardbp/Mcm2/Smarcc1/Nme1/Eif3j2/Cdk2/Snrpb2/Mrps18b/Nop56/Srpk1/Cbx3/Hspe1/Nhp2/Ncbp2/Pwp1/Dhx15 |
| HALLMARK_UNFOLDED_PROTEIN_RESPONSE | 0,94700461 | 1 | 0,05468085 | 0,21697427 | 0,75425238 | 99 | Slc1a4/Asns/Cks1b/Slc7a5/Dnaja4/Paip1/Khsrp/Tatdn2/Eif2ak3/Wfs1/Bag3/Hyou1/Dkc1/Exosc2 |
| HALLMARK_PROTEIN_SECRETION | 0,9751693 | 1 | 0,05237591 | 0,19775414 | 0,68017681 | 88 | Krt18/Abca1/Gla/Sec24d/Sgms1/Clcn3/Cd63/Vamp7/Ap3s1/Golga4/Atp1a1/Bet1/Rab9/Dop1a/Arfgap3/Ica1/Rab2a/Cln5/Uso1/Rab5a/Stx16 |
| HALLMARK_FATTY_ACID_METABOLISM | 0,99548533 | 1 | 0,05142649 | 0,18058684 | 0,64744803 | 122 | Idi1/Adh1/Hsd17b7/Tdo2/Kmt5a/Uros/Hsd17b11/Nthl1/Dhcr24/Hmgcs1/Nsdhl/Lgals1/Ccdc58/Odc1/Sdhd/Reep6/Acsl1/Retsat/Elovl5/Hsph1/Nbn/Acsl4/Acat3/Cpox/Gcdh/Fasn/Dld/G0s2/Idh1/Car2/Apex1/Mif/Mlycd/Acot2/Eci2/Gstz1/Mcee/Fh1/Eno2 |
| HALLMARK_PANCREAS_BETA_CELLS | 0,94080338 | 1 | 0,05101141 | 0,24711495 | 0,58425668 | 13 | Pcsk1/Elp4/Akt3 |

| | | | | | | | |
|---|---|---|---|---|---|---|---|
| HALLMARK_DNA_REPAIR | 1 | 1 | 0,05153091 | 0,14553722 | 0,53065177 | 142 | Nme4/Rad51/Taf9/Tyms/Prim1/Rfc4/Pola1/Dut/Sac3d1/Pola2/Smad5/Rfc5/Bola2/Stx3/Pom121/Rpa3/Polh/Adcy6/Gtf2h3/Snapc4/Polr2d/Polr2h/Vps37b/Snapc5/Polr2f/Polb/Polr2g/Cstf3/Nme1/Aprt/Rfc3/Rnmt/Tarbp2/Mpg/Gtf2h5/Ncbp2 |
| HALLMARK_REACTIVE_OXYGEN_SPECIES_PATHWAY | 0,98327138 | 1 | 0,04310368 | -0,1951795 | -0,5713382 | 47 | Ftl1/Lsp1/Gpx4/Fes/Ndufs2/Mbp/Stk25/Pfkp/Junb/Sod1/Ipcef1/Lamtor5/Hmox2/Hhex/Sbno2/Pdlim1/Egln2/Prnp |
| HALLMARK_TGF_BETA_SIGNALING | 0,93160813 | 1 | 0,04531524 | -0,2417868 | -0,6980833 | 43 | Cdkn1c/Id3/Id1/Rhoa/Ppp1ca/Ube2d3/Hdac1/Rab31/Klf10/Ppp1r15a/Arid4b/Junb |
| HALLMARK_PI3K_AKT_MTOR_SIGNALING | 0,9177102 | 1 | 0,04450705 | -0,2246556 | -0,7516001 | 94 | Gna14/E2f1/Pfn1/Cfl1/Tiam1/Pik3r3/Lck/Arhgdia/Trib3/Rptor/Sqstm1/Ppp1ca/Il2rg/Nod1/Ube2d3/Rac1/Pak4/Prkcb/Slc2a1/Actr3/Arf1/Them4 |
| HALLMARK_XENOBIOTIC_METABOLISM | 0,8438061 | 1 | 0,04850598 | -0,2300548 | -0,8108936 | 129 | Hgfac/Gsta3/Ddc/Cdo1/Mpp2/Slc35d1/Maoa/Fah/Irf8/Ccl25/Ephx1/Esr1/Tmbim6/Pros1/Psmb10/Gch1/Cndp2/Slc46a3/Slc12a4/Ndrg2/Cyb5a/Ap4b1/Gabarapl1/Ptges3 |

**Supplementary Table 4. GSEA hallmark gene sets in effector Treg cells**

| pathway | pval | padj | log2err | ES | NES | size | leadingEdge |
|---|---|---|---|---|---|---|---|
| HALLMARK_CHOLESTEROL_HOMEOSTASIS | 6,1093E-06 | 0,00030546 | 0,61052688 | 0,64639197 | 2,24109856 | 54 | Scd1/Scd2/Idi1/Sqle/Ldlr/Cyp51/Fdps/Hmgcs1/Aldoc/Tmem97/Mvk/Fdft1/Mvd/Tm7sf2/Lss/Sc5d/Srebf2/Hmgcr/Hsd17b7/Pcyt2/Tnfrsf12a/Cpeb2/Fabp5/Acat3/Nsdhl |
| HALLMARK_MTORC1_SIGNALING | 3,8418E-05 | 0,00096044 | 0,55733224 | 0,44248116 | 1,8275249 | 184 | Scd1/Scd2/Idi1/Sqle/Dhcr24/Ldlr/Cyp51/Acsl3/Insig1/Acaca/Hmgcs1/Slc1a4/Cdkn1a/Tmem97/Fgl2/Tm7sf2/Cxcr4/Sc5d/Ddit4/Hmgcr/Sytl2/Ccnf/Elovl6/Gclc/Slc7a5/Gga2/Ung/Fkbp2/Psmg1/Hspe1/Stard4/Elovl5/Nmt1/Rdh11/Pdap1/Cfp/Nfil3/Nufip1/Dhcr7/Tfrc/Fads1/Eef1e1/Rrp9/Shmt2/Ppa1 |
| HALLMARK_MYC_TARGETS_V1 | 0,00149698 | 0,0249496 | 0,45505987 | 0,37566762 | 1,56340266 | 200 | Myc/Cdc45/Nop16/Rfc4/Nme1/Srm/Ndufab1/Hspe1/Hdgf/Trim28/Pcbp1/Pabpc4/Snrpd1/Apex1/Tardbp/Lsm7/Set/Nolc1/Pa2g4/Tyms/Rrp9/Ssbp1/Erh/Kpnb1/Phb/Ruvbl2/Rack1/C1qbp/Cbx3/Txnl4a/Snrpa1/Mrpl23/Aimp2/Ifrd1/U2af1/Fam120a/Eif2s2/Snrpd2/Tfdp1/Nop56/Impdh2/Nhp2/Eif3b/Snrpa/Pwp1/Hnrnpd/Xrcc6/Lsm2/Eef1b2/Cct5/Rps10/Rnps1/Cdk4/Pold2/Eif1ax/Vdac1/Dut/Bub3/Etf1/Cdk2/Ctps/Srpk1/Eif2s1/Ranbp1/Gnl3/Npm1/Prdx3/Psma7/Hdac2/Hnrnpa1/Snrpd3/Ran/Rpl22 |
| HALLMARK_MYC_TARGETS_V2 | 0,00258362 | 0,0322953 | 0,4317077 | 0,52604538 | 1,83421225 | 57 | Tmem97/Slc29a2/Myc/Dctpp1/Nop16/Ung/Srm/Hspe1/Pus1/Mrto4/Nolc1/Pa2g4/Rrp9/Ndufaf4/Phb/Cbx3/Nip7/Aimp2/Rabepk/Nop56/Nop2/Ppan/Rrp12/Cdk4/Ipo4/Rcl1/Exosc5/Tfb2m/Gnl3/Npm1/Slc19a1/Pprc1/Grwd1/Tbrg4/Farsa/Wdr74 |
| HALLMARK_REACTIVE_OXYGEN_SPECIES_PATHWAY | 0,00340216 | 0,03402158 | 0,4317077 | -0,6866019 | -2,313633 | 46 | Mpo |
| HALLMARK_COMPLEMENT | 0,00450674 | 0,03755613 | 0,40701792 | -0,4030697 | -1,5789857 | 118 | Car2/Mmp14/Plek/Cpq/Gnb4/Serpinc1/Casp3/Spock2/Lipa/Cd40lg/Actn2/Timp2/Cd46/Dgkg/Dgkh/Ang/Lcp2/Pfn1/Irf1/Prcp/Plscr1/Ctss/Ctsd/Casp9/Anxa5/Psen1/Casp7/Sh2b3/Pik3cg/Ctsb/Cd55/Cd55b/Pdgfb/Lck/Dpp4/Prkcd/Gnai3/Lyn/Gmfb/Ctso |
| HALLMARK_ESTROGEN_RESPONSE_LATE | 0,00809369 | 0,05781204 | 0,3807304 | 0,41344415 | 1,60826573 | 109 | Cyp26b1/Cdc6/Slc1a4/Nrip1/Celsr2/Fdft1/Ckb/Scarb1/Chpt1/Igfbp4/Car12/Gins2/Jak2/Rps6ka2/Areg/Fkbp5/Abhd2/Slc7a5/Tob1/Fabp5/Ppif/Slc22a5/Bcl2/Elovl5 |
| HALLMARK_ANDROGEN_RESPONSE | 0,01102057 | 0,06337543 | 0,3807304 | 0,43983445 | 1,64057057 | 82 | Scd1/Scd2/Idi1/Dhcr24/Acsl3/Insig1/Srf/Hmgcs1/Cenpn/Hmgcr/Rab4a/Fkbp5/Abhd2/Pmepa1/Dbi/Elovl5 |
| HALLMARK_ESTROGEN_RESPONSE_EARLY | 0,01140758 | 0,06337543 | 0,3807304 | 0,40491606 | 1,57509212 | 109 | Cyp26b1/Slc1a4/Nrip1/Celsr2/Myc/Fdft1/Akap1/Scarb1/Chpt1/Igfbp4/Car12/Dhrs3/Jak2/Rps6ka2/Areg/Fkbp5/Nav2/Abhd2/Slc7a5/Tob1/Ppif/Slc22a5/Bcl2/Elovl5/Tubb2b/Ugcg/Siah2/Fasn |
| HALLMARK_IL2_STAT5_SIGNALING | 0,01476045 | 0,07380224 | 0,3807304 | -0,3683317 | -1,52836 | 177 | Car2/Alcam/Cst7/Itgae/Plpp1/Sh3bgrl2/Capg/Irf8/Gpr83/Serpinc1/Irf6/Ptch1/Gucy1b1/Myo1e/Casp3/Tnfrsf9/Drc1/Cd79b/Nt5e/Gbp3/Tiam1/Igf1r/Il2rb/Bmpr2/Il18r1/Ltb/Phtf2/Fah/She/Slc2a3/Lclat1/Prnp/Enpp1/Cyfip1/Nrp1/Csf1/Tlr7/Plscr1/Twsg1/Swap70/Penk/Traf1/Cd81 |
| HALLMARK_UV_RESPONSE_UP | 0,02529339 | 0,11315839 | 0,35248786 | -0,363941 | -1,4351046 | 121 | Car2/H2-Q1/Mmp14/H2-Q2/Fosb/Asns/Aqp3/Ntrk3/Casp3/Plcl1/Btg3/Clcn2/Hspa13/Cebpg/Ephx1/Il6st/Irf1/Btg2/Epcam/Bcl2l11/Tap1/Tst/E2f5/H2-M3/Abcb1a/Tmbim6/Hmox1/Aldoa |
| HALLMARK_ANGIOGENESIS | 0,02715801 | 0,11315839 | 0,35248786 | -0,7021303 | -1,5543797 | 9 | S100a4/Nrp1/Pglyrp1/Itgav |
| HALLMARK_MYOGENESIS | 0,03654975 | 0,1218325 | 0,32177592 | 0,39701313 | 1,46627852 | 77 | Scd1/Scd2/Fdps/Cdkn1a/Ckb/Foxo4/Mapk12/Pygm/Gpx3/Pfkm/Nav2/Hdac5/Hbegf/Eno3/Plxnb2 |
| HALLMARK_FATTY_ACID_METABOLISM | 0,03383436 | 0,1218325 | 0,32177592 | 0,35900769 | 1,40688345 | 117 | Idi1/Dhcr24/G0s2/Hmgcs1/Nthl1/Hsd17b7/Cryz/Eno2/Uros/Acat3/Nsdhl/Slc22a5/Kmt5a/Sdhd/Elovl5/Ccdc58/Eno3/Apex1/Mif/Rdh11/Glul/Fasn |

| | | | | | | | |
|---|---|---|---|---|---|---|---|
| HALLMARK_ALLOGRAFT_REJECTION | 0,03476273 | 0,1218325 | 0,32177592 | -0,3894531 | -1,5749377 | 150 | Gcnt1/Cd7/Capg/Il18rap/Irf8/Prf1/Ifng/Cd80/Cd96/H2-Q7/H2-Aa/Il2/Flna/Il2rb/Gbp2/Ltb/Il18/Stat1/Cd40lg/Cd74/B2m/Hif1a/Ly75/Hdac9/Srgn/Tapbp/Lcp2/F2r/Il16/Csf1/Ets1/Ctss/Fgr/Zap70/Ptprc/Tlr2/H2-Q10/Tap1/H2-T23/Bcat1/Itgb2/Cd28/H2-M3/Aars/Cd2/Lck/Cd3e/Sit1/Cd247/Cd3g/Lyn/Cd47/Cd1d1/Ifnar2/Tnf/Ccnd3/Psmb10/Ptpn6 |
| HALLMARK_MITOTIC_SPINDLE | 0,04080329 | 0,12751029 | 0,32177592 | -0,3271451 | -1,3477608 | 169 | Palld/Synpo/Arhgap5/Kif11/Fgd6/Myo1e/Rapgef5/Ccdc88a/Clip1/Flna/Tiam1/Kif1b/Cdk1/Pdlim5/Als2/Nedd9/Incenp/Cntrl/Racgap1/Dock2/Hdac6/Ssh2/Flnb/Rab3gap1/Cdk5rap2/Tbcd/Bcl2l11/Plekhg2/Epb41l2/Smc4/Abr/Cdc42/Tubgcp5/Arhgap10/Dlg1/Smc1a/Arhgef11/Abl1/Clasp1/Myh9/Arhgap27/Arhgef2/Arhgef3/Tubgcp6/Cep57/Cep250/Numa1/Stk38l/Epb41/Csnk1d/Arhgdia/Sun2/Stau1/Pcm1 |
| HALLMARK_G2M_CHECKPOINT | 0,04608208 | 0,12800579 | 0,32177592 | 0,32902245 | 1,34323113 | 165 | Sqle/Cdc6/Chek1/Myc/Rad54l/Cdkn1b/Gins2/Orc6/Mybl2/Sap30/Uck2/Pole/Cdc45/Ccnf/Upf1/Slc7a5/Exo1/Notch2/Tmpo/Chaf1a/Kmt5a/Traip/Cks1b/Kif22/Ube2s/Snrpd1/Prim2/Dkc1/Cenpe/Pura/Cbx1/Stmn1/Bcl3/Nolc1/E2f4/Sfpq/Kpnb1/Hmga1/Ccnb2 |
| HALLMARK_E2F_TARGETS | 0,0451505 | 0,12800579 | 0,2765006 | 0,32017336 | 1,32260869 | 181 | Cdkn1a/Chek1/Myc/Cdkn1b/Shmt1/Rad51ap1/Orc6/Ccne1/Mybl2/Dctpp1/Pole/Rad51c/Trip13/Nme1/Ung/Tmpo/Psmc3ip/Cenpm/Rpa3/Lyar/Cks1b/Dlgap5/Kif22/Tk1/Ube2s/Bub1b/Prim2/Cdca3/Cenpe/Tipin/Stmn1/Nolc1/Tfrc/Pa2g4/Lig1/Hmga1/Cbx5/Ccnb2/Spc24/Mcm3/Gins4/Diaph3/Nop56/Naa38/Pold1/Hnrnpd/Xrcc6/Mlh1/Cdk4/Pop7/Pold2 |
| HALLMARK_INFLAMMATORY_RESPONSE | 0,05811966 | 0,15294118 | 0,24504179 | 0,3370443 | 1,3140109 | 114 | Ldlr/Stab1/Cxcr6/Cdkn1a/Scarf1/Myc/Sell/Ptgir/Abca1/Ahr/Il10/Slc31a1/Tnfsf9/Rgs16/Mxd1/Gna15/Hbegf/Bst2/Itgb3/Ptger2/Chst2/Lif/Ptger4/Ccl5/Gpr183/Ebi3/Kcna3/Ifngr2/Irf7 |
| HALLMARK_EPITHELIAL_MESENCHYMAL_TRANSITION | 0,06117647 | 0,15294118 | 0,28201335 | -0,3813246 | -1,3845569 | 65 | Matn2/Vcam1/Mmp14/Capg/Jun/Gem/Sgcb/Nt5e/Flna/Tgfbr3/Vim/Pfn2 |
| HALLMARK_DNA_REPAIR | 0,0681431 | 0,16224548 | 0,22496609 | 0,31755083 | 1,27812485 | 141 | Nme4/Adcy6/Stx3/Rad51/Rfc4/Nme1/Bola2/Gtf2h3/Polr2h/Rpa3/Nt5c/Prim1/Fen1/Polr2d/Polr2j/Guk1/Sac3d1/Tyms/Nme3/Lig1/Polr1d/Polr2g/Impdh2/Pold1/Polr2i/Gtf2h5/Polr2f/Smad5/Nelfe/Polr2k/Taf12/Brf2/Dut/Polr2e/Gtf2a2 |
| HALLMARK_HEME_METABOLISM | 0,07616708 | 0,17310699 | 0,25720647 | -0,326432 | -1,306321 | 143 | Car2/Hbb-bt/Tspan5/Daam1/Igsf3/Asns/Aqp3/Abcg2/Endod1/Tns1/Ctse/Optn/Kdm7a/Ucp2/Btg2/Ypel5/E2f2/Rnf19a/Bpgm/Tent5c/Xpo7/Btrc/Ctsb/Lrp10/Dcaf11/Clcn3/Myl4/Cast/Slc2a1/Ezh1/Ubac1/Ccnd3/Pdzk1ip1/Cir1/Nek7/Ncoa4/Top1/Bach1/Sdcbp |
| HALLMARK_INTERFERON_GAMMA_RESPONSE | 0,09245742 | 0,18735363 | 0,23112671 | -0,3047533 | -1,2510821 | 166 | Vcam1/St3gal5/Irf8/Slamf7/Bank1/Casp3/H2-Q7/Txnip/H2-Aa/Gbp3/Il2rb/H2-D1/Stat1/Cd74/H2-Eb1/B2m/Hif1a/Eif4e3/Tapbp/Isoc1/Lcp2/Trim26/Ifit2/Gm8909/Irf1/Plscr1/Ly6e/Bpgm/H2-Q10/Itgb7/Casp7/Samhd1/Tap1/Stat3/Ube2l6/Epsti1/Lats2/Peli1/Il15ra/H2-M3/Mthfd2/Pfkp/Slc25a28/Psme1/Ifih1/Gbp9/Psmb8/Oas3/Vamp5/Gbp4/Gpr18/Sppl2a/Ifnar2/Psmb10/Gbp10/Ptpn6/Ifi27/Mvp |
| HALLMARK_KRAS_SIGNALING_UP | 0,09367681 | 0,18735363 | 0,22496609 | -0,3335422 | -1,2816223 | 105 | Car2/Irf8/F2rl1/Kif5c/Scg5/Gucy1a1/Vwa5a/Spon1/Hdac9/Hsd11b1/Dnmbp/Map7/Lcp1/Etv5/Ikzf1/Nrp1/Ly96/Avl9/Ets1/Ctss/Ppp1r15a/Dock2/Glrx/Traf1/Bpgm/Akap12/Rabgap1l/Tmem176b/Tnfrsf1b/Mmp11/Itgb2/Abcb1a/Pecam1/Sdccag8/Map3k1/Psmb8/Tmem158/Laptm5 |
| HALLMARK_COAGULATION | 0,08775982 | 0,18735363 | 0,23112671 | -0,3829126 | -1,3472239 | 56 | Mmp14/Plek/Cpq/Crip2/Serpinc1/Ctse/Gng12/Capn2/Ang/Casp9/Ctsb/Mmp11/Anxa1/Pdgfb/Pecam1 |
| HALLMARK_P53_PATHWAY | 0,1060241 | 0,20389249 | 0,21392786 | -0,2999375 | -1,2230977 | 153 | Stom/Trib3/Fgf13/S100a4/Jun/Hspa4l/Txnip/St14/Vwa5a/Ldhb/Ppm1d/Nudt15/Cd82/Ptpre/Baiap2/Abhd4/Wwp1/Hexim1/Rab40c/Dgka/S100a10/Ephx1/F2r/Ccng1/Btg2/Ctsd/Ppp1r15a/Apaf1/Kif13b/Cd81/Prmt2/Tap1/Tsc22d1/Acvr1b/Coq8a/Gadd45a |
| HALLMARK_SPERMATOGENESIS | 0,13636364 | 0,25252525 | 0,18138313 | -0,3557742 | -1,2671112 | 58 | Art3/Tulp2/Hspa4l/Scg5/Rad17/Cdk1/Ezh2/She/Nf2/Map7/Gmcl1/Agfg1 |

| | | | | | | | |
|---|---|---|---|---|---|---|---|
| HALLMARK_HYPOXIA | 0,14309764 | 0,25553151 | 0,1501698 | 0,30537913 | 1,1996626 | 119 | Plac8/Cdkn1a/Aldoc/Angptl4/Scarb1/Cdkn1b/Car12/Cxcr4/Pygm/Ddit4/Sap30/Eno2/Foxo3/Klf7/Chst2/Bcl2/Tktl1/Eno3/Sdc4/Galk1/Mif/Siah2/Nfil3/Fam162a |
| HALLMARK_APOPTOSIS | 0,15496368 | 0,26717876 | 0,17520405 | -0,3018692 | -1,1879227 | 120 | Mgmt/Pak1/Prf1/Jun/Casp3/Cdc25b/Txnip/Btg3/Tgfbr3/Il18/Pmaip1/Timp2/Madd/F2r/Nedd9/Irf1/Btg2/Casp9/Psen1/Ppp3r1/Casp7/Bcl2l11/Tap1/Hmgb2/Anxa1/Gadd45a/Plcb2/Hmox1/Cd2/Dnm1l/Pea15a/Diablo/Dap |
| HALLMARK_IL6_JAK_STAT3_SIGNALING | 0,16046512 | 0,26744186 | 0,16823817 | -0,3437339 | -1,2286182 | 60 | Il13ra1/Jun/Il17rb/Il18r1/Ltb/Stat1/Il6st/Irf1/Csf1/Tlr2/Tnfrsf1b/Stat3/Acvr1b/Il15ra/Il17ra/Hmox1 |
| HALLMARK_INTERFERON_ALPHA_RESPONSE | 0,19723183 | 0,31811586 | 0,12750532 | 0,31213922 | 1,16873905 | 87 | Tent5a/Mx2/Sell/Oasl1/Usp18/Ifi44/Rsad2/Isg15/Oas1g/Trim25/Oas1a/Bst2/Tdrd7/Ifitm3/Helz2/Pnpt1/Lap3/Irf7/Ncoa7 |
| HALLMARK_APICAL_JUNCTION | 0,21647059 | 0,33823529 | 0,14375899 | -0,2950786 | -1,1315383 | 102 | Vcam1/Actg2/Pik3r3/Syk/Sirpa/Pard6g/Actg1/Exoc4/Baiap2/Acta1/Actn2/Nf2/Actb/Pfn1/Arhgef6/Cnn2/Traf1/Ptprc/Rac2/Msn/Epb41l2/Atp1a3 |
| HALLMARK_WNT_BETA_CATENIN_SIGNALING | 0,23047619 | 0,34920635 | 0,12325723 | 0,40580211 | 1,18771859 | 26 | Ppard/Myc/Hdac5/Axin2/Csnk1e/Kat2a/Rbpj |
| HALLMARK_PEROXISOME | 0,2862069 | 0,42087542 | 0,10244941 | 0,29688294 | 1,09647022 | 77 | Idi1/Dhcr24/Fdps/Dhrs3/Gstk1/Elovl5/Abcb9/Atxn1/Rdh11/Fads1/Slc25a19/Hras/Dlg4/Ctbp1/Idh1/Hsd17b11 |
| HALLMARK_BILE_ACID_METABOLISM | 0,30071174 | 0,42087542 | 0,10135074 | 0,3126016 | 1,08836929 | 58 | Idi1/Dhcr24/Abca1/Pex26/Pfkm/Gstk1/Nedd4/Pecr/Lipe |
| HALLMARK_TNFA_SIGNALING_VIA_NFKB | 0,3030303 | 0,42087542 | 0,09754492 | 0,26381464 | 1,06736282 | 144 | Ldlr/G0s2/Bcl6/Cdkn1a/Myc/Sik1/Abca1/Trip10/Plk2/Kdm6b/Areg/B4galt5/Dusp5/Tubb2a/Tnfsf9/Pmepa1/Mxd1/Lamb3/Ier5/Hbegf/Lif/Ptger4/Ccl5/Gpr183/Map3k8/Sdc4/Nfil3/Ifngr2/Bcl3/Ets2/Zc3h12a/Dnajb4/Icosl |
| HALLMARK_KRAS_SIGNALING_DN | 0,36888112 | 0,47568173 | 0,08835944 | 0,316556 | 1,05521264 | 45 | Mx2/Celsr2/Tg/Rsad2/Slc25a23/Fggy/Ypel1/Gpr19/Skil/Chst2/Rgs11 |
| HALLMARK_PROTEIN_SECRETION | 0,36428571 | 0,47568173 | 0,10755438 | -0,269523 | -1,0204593 | 86 | Kif1b/Anp32e/Pam/Ap3b1/Cltc/Gbf1/Clcn3/Tmx1/Arfip1/Mapk1/Dnm1l/Atp7a/Vps45/Snx2/Stx7/Ocrl/Ctsc/Tom1l1/Lamp2/Stx12/Ap3s1/Sec31a/Tsg101/Ppt1/Stam/Gla/Tpd52/Copb2/Vps4b/Cog2/Rab14/Rab5a/Ap1g1/Arfgap3/Uso1/Arfgef2 |
| HALLMARK_APICAL_SURFACE | 0,37103175 | 0,47568173 | 0,09528798 | -0,4014186 | -1,0659362 | 16 | Cd160/Il2rb/Akap7/Lyn |
| HALLMARK_XENOBIOTIC_METABOLISM | 0,43309002 | 0,54136253 | 0,09821234 | -0,2567916 | -0,9927443 | 113 | Car2/Hgfac/Irf8/Ddc/Xdh/Fah/Pdlim5/Hsd11b1/Abcc3/Ephx1/Cyp2s1/Adh1/Abhd6/Tmem176b/Cndp2/Acp2/Bcat1/Tmbim6/Hacl1/Hmox1/Csad/Pink1 |
| HALLMARK_UV_RESPONSE_DN | 0,46282974 | 0,56442651 | 0,09344492 | -0,256128 | -0,9636925 | 85 | Gcnt1/Prkar2b/Mgmt/Pik3r3/Cdon/Fhl2/Igf1r/Tgfbr3/Nr1d2/Pdlim5/Nrp1/Synj2/Sfmbt1/Scaf8/Prkca/Grk5/Atp2b4/Cacna1a/Dlg1/Sipa1l1/Atp2c1 |
| HALLMARK_HEDGEHOG_SIGNALING | 0,53036437 | 0,63138616 | 0,07667469 | 0,3571205 | 0,92684135 | 17 | Nf1/Tle1/Adgrg1/Ldb1/Rasa1/Ets2/Thy1 |
| HALLMARK_NOTCH_SIGNALING | 0,63653846 | 0,740161 | 0,06494077 | 0,31133548 | 0,85898634 | 20 | Ppard/Sap30/Notch2 |
| HALLMARK_PI3K_AKT_MTOR_SIGNALING | 0,70238095 | 0,79816017 | 0,07061962 | -0,2269805 | -0,8630439 | 91 | Gna14/Trib3/Pik3r3/Tiam1/Cdk1/Pdk1/Pfn1/Cltc/Mapk9/Vav3/Hsp90b1/Pak4/Mapk1/Ppp2r1b/Lck/Slc2a1/Cfl1/Calr |
| HALLMARK_OXIDATIVE_PHOSPHORYLATION | 0,78487395 | 0,87208217 | 0,04879897 | 0,20904952 | 0,86824834 | 193 | Mgst3/Ndufab1/Alas1/Ndufa1/Sdhd/Timm9/Atp5e/Mrps11/Uqcrq/Timm17a/Timm8b/Uqcr11/Cox7a2/Slc25a20/Cox7b/Timm13/Mrpl15/Ndufc2/Ndufb6/Ndufb3/Ndufb4/Nqo2/Mrpl34/Atp5g2/Uqcr10/Cpt1a/Atp6v0b/Idh1/Atp5g1/Cox7c/Atp5k/Polr2f/Timm50/Ndufa4/Vdac1/Ndufa2/Uqcrb/Prdx3/Grpel1/Ndufs6/Mtrf1/Hsd17b10/Fxn/Atp5j2/Atp5g3/Ndufc1/Fh1/Timm10/Ndufa3/Mrps30/Mrpl35/Pdp1/Cs/Ech1/Ndufb2/Cox10/Tcirg1/Mtrr/Cyc1/Etfa/Cox6a1/Ndufs7/Htra2/Mrpl11/Mrps22/Isca1/Ndufv2/Ndufv1/Bax/Sdhc/Sucla2/Ndufb8/Mpc1/Sdhb/Ndufa7/Dld/Cox11/Vdac3/Atp5h/Hspa9/Atp5d/Aldh6a1/Cox5b/Ndufb5/Tomm70a/Atp5l/Eci1/Etfdh/Phb2/Ndufs8/Idh3b |
| HALLMARK_ADIPOGENESIS | 0,82149047 | 0,89292442 | 0,0481184 | 0,20933312 | 0,85064442 | 159 | Adcy6/Bcl6/Angptl4/Abca1/Scarb1/Esrra/Mgst3/Gpx3/Itsn1/Itih5/Elovl6/Vegfb/Tob1/Ndufab1/Lipe/Nmt1/Uqcrq/Uqcr11/Cox7b/Dhcr7/Mrpl15/Pdcd4/Sqor/Pim3/Slc25a1/Aldh2/Ddt/Uqcr10/Idh1/Lpcat3/Ppm1b/Nkiras1 |
| HALLMARK_PANCREAS_BETA_CELLS | 0,92220114 | 0,96062619 | 0,04697587 | 0,26331445 | 0,59907764 | 10 | Akt3/Foxo1/Stxbp1/Srp9/Elp4 |
| HALLMARK_TGF_BETA_SIGNALING | 0,90929204 | 0,96062619 | 0,05468085 | -0,2139788 | -0,6931649 | 38 | Id3/Bmpr2/Tgfbr1/Rab31/Ppp1r15a |
| HALLMARK_UNFOLDED_PROTEIN_RESPONSE | 1 | 1 | 0,03800562 | 0,15864014 | 0,60935826 | 99 | Slc1a4/Ddit4/Paip1/Tubb2a/Slc7a5/Exosc2/Cks1b/Dkc1/Nolc1/Rrp9/Eif4ebp1/Zbtb17/Nop56/Nhp2/Atf4/Tatdn2/Banf1/Dcp2/Exosc5/Exosc1/Eif2s1/Npm1/Nfya/Wfs1/Ifit1bl1 |

| | | | | | | | |
|---|---|---|---|---|---|---|---|
| HALLMARK_GLYCOLYSIS | 0,98292683 | 1 | 0,05559471 | -0,1761655 | -0,7021311 | 136 | Il13ra1/Hmmr/Nt5e/Phka2/Pfkfb1/Cdk1/Pam/Glce/Ang/Homer1/Glrx/Chpf2/Hk2/Eno1/Aldoa/Pfkp/Qsox1/Prps1 |

**Supplementary Table 5. Top 50 Canonical Pathways between NCOR1-cKO and WT naive Treg cells**

| Ingenuity Canonical Pathways | -log(p-value) | zScore | Ratio | Molecules |
|---|---|---|---|---|
| Superpathway of Cholesterol Biosynthesis | 9,26 | 3,605551 | 0,542 | CYP51A1,DHCR24,FDFT1,FDPS,HMGCR,HMGCS1,IDI1,LSS,MSMO1,MVK,NSDHL,SQLE,TM7SF2 |
| Cholesterol Biosynthesis I | 6,85 | 2,828427 | 0,667 | CYP51A1,DHCR24,FDFT1,LSS,MSMO1,NSDHL,SQLE,TM7SF2 |
| Cholesterol Biosynthesis II (via 24,25-dihydrolanosterol) | 6,85 | 2,828427 | 0,667 | CYP51A1,DHCR24,FDFT1,LSS,MSMO1,NSDHL,SQLE,TM7SF2 |
| Cholesterol Biosynthesis III (via Desmosterol) | 6,85 | 2,828427 | 0,667 | CYP51A1,DHCR24,FDFT1,LSS,MSMO1,NSDHL,SQLE,TM7SF2 |
| Crosstalk between Dendritic Cells and Natural Killer Cells | 5,74 | -1,26491 | 0,258 | ACTB,ACTG1,ACTG2,CAMK2D,CD40LG,CD80,CD83,CD86,FASLG,HLA-A,HLA-E,HLA-F,IL2RB,LTA,LTB,NECTIN2 |
| RhoA Signaling | 4,9 | -2,82843 | 0,207 | ACTB,ACTG1,ACTG2,ARHGAP5,ARHGAP9,ARHGEF1,BAIAP2,CDC42EP3,CFL1,MPRIP,MSN,MYL12B,PFN1,RDX,ROCK2,SEPTIN1,SEPTIN8,SEPTIN9 |
| Zymosterol Biosynthesis | 4,04 | 2 | 0,8 | CYP51A1,MSMO1,NSDHL,TM7SF2 |
| Caveolar-mediated Endocytosis Signaling | 3,99 | | 0,22 | ACTB,ACTG1,ACTG2,B2M,FLNA,HLA-A,HLA-E,HLA-F,ITGAE,ITGB2,ITGB3,ITGB7,ITGB8 |
| Dendritic Cell Maturation | 3,86 | -2,82843 | 0,16 | B2M,CD1D,Cd1d2,CD40LG,CD80,CD83,CD86,FCGR3A/FCGR3B,HLA-A,HLA-DOB,HLA-E,HLA-F,IRF8,JAK2,LTA,LTB,NFKBIA,PLCB4,PLCL1,RELB,STAT1 |
| Allograft Rejection Signaling | 3,77 | | 0,256 | B2M,CD40LG,CD80,CD86,FASLG,H2-K2/H2-Q9,HLA-A,HLA-DOB,HLA-E,HLA-F |
| PD-1, PD-L1 cancer immunotherapy pathway | 3,72 | 3 | 0,182 | B2M,CD80,CDKN1B,FOXP3,HLA-A,HLA-DOB,HLA-E,HLA-F,IL2RA,IL2RB,JAK2,LAT,LCK,LCP2,STAT5A,STAT5B |
| Th1 and Th2 Activation Pathway | 3,66 | | 0,152 | CD40LG,CD80,CD86,HLA-A,HLA-DOB,IL18R1,IL1RL1,IL2RA,IL2RB,IL4R,IL6R,IRF1,ITGB2,JAK2,LTA,NFATC3,SOCS1,STAT1,STAT5A,STAT5B,TNFRSF4,TNFSF11 |
| RhoGDI Signaling | 3,6 | 1,290994 | 0,157 | ACTB,ACTG1,ACTG2,ARHGAP5,ARHGAP9,ARHGDIA,ARHGDIB,ARHGEF1,ARHGEF10,Cdc42,CFL1,GNA15,GNAI3,GNB1,MSN,MYL12B,RAC2,RDX,RHOH,ROCK2 |
| Leukocyte Extravasation Signaling | 3,49 | -0,44721 | 0,145 | ACTB,ACTG1,ACTG2,ARHGAP5,ARHGAP9,CLDN10,GNAI3,ITGB2,ITGB3,MMP14,MMP9,MSN,NCF1,RAC2,RAPGEF4,RASSF5,RDX,RHOH,ROCK2,SELPLG,SIPA1,THY1,VCAM1 |
| Antigen Presentation Pathway | 3,47 | | 0,286 | B2M,HLA-A,HLA-DOB,HLA-E,HLA-F,PSMB8,TAP1,TAPBP |
| B Cell Development | 3,28 | | 0,304 | CD79B,CD80,CD86,HLA-A,HLA-DOB,IGHM,PTPRC |
| Type I Diabetes Mellitus Signaling | 3,27 | 0 | 0,167 | CD80,CD86,FASLG,HLA-A,HLA-DOB,HLA-E,HLA-F,IRAK1,IRF1,JAK2,LTA,NFKBIA,SOCS1,SOCS2,SOCS7,STAT1 |
| Autoimmune Thyroid Disease Signaling | 3,24 | | 0,267 | CD40LG,CD80,CD86,FASLG,HLA-A,HLA-DOB,HLA-E,HLA-F |
| iCOS-iCOSL Signaling in T Helper Cells | 3,17 | -1,94145 | 0,163 | Calm1 (includes others),CAMK2D,CD40LG,CD80,CD86,HLA-A,HLA-DOB,IL2RA,IL2RB,LAT,LCK,LCP2,NFATC3,NFKBIA,PPP3CC,PTPRC |
| Altered T Cell and B Cell Signaling in Rheumatoid Arthritis | 3,16 | | 0,183 | CD40LG,CD79B,CD80,CD86,FASLG,HLA-A,HLA-DOB,LTA,LTB,RELB,SLAMF1,TNFRSF13B,TNFSF11 |
| T Helper Cell Differentiation | 2,99 | | 0,185 | BCL6,CD40LG,CD80,CD86,FOXP3,HLA-A,HLA-DOB,IL18R1,IL2RA,IL4R,IL6R,STAT1 |
| Superpathway of Geranylgeranyldiphosphate Biosynthesis I (via Mevalonate) | 2,98 | 2,236068 | 0,385 | FDPS,HMGCR,HMGCS1,IDI1,MVK |
| Role of JAK2 in Hormone-like Cytokine Signaling | 2,82 | | 0,259 | JAK2,SOCS1,SOCS2,SOCS7,STAT1,STAT5A,STAT5B |
| Cell Cycle: G2/M DNA Damage Checkpoint Regulation | 2,69 | -1,66667 | 0,205 | AURKA,CCNB2,CDK1,CHEK1,CHEK2,CKS1B,PLK1,TOP2A,YWHAZ |
| Communication between Innate and Adaptive Immune Cells | 2,64 | | 0,177 | B2M,CD40LG,CD79B,CD80,CD83,CD86,HLA-A,HLA-E,HLA-F,IGHM,TNFRSF13B |
| Systemic Lupus Erythematosus In T Cell Signaling Pathway | 2,64 | -1,63299 | 0,124 | B2M,BCL6,CD40LG,CD80,CD86,Cdc42,FASLG,FOXP3,GNAI3,H2-K2/H2-Q9,HLA-A,HLA-DOB,HLA-E,HLA-F,LAT,MSN,PPP2CB,PPP3CC,RAC2,RDX,RHOH,ROCK2,RPTOR,SELPLG |
| Agranulocyte Adhesion and Diapedesis | 2,63 | | 0,145 | ACTB,ACTG1,ACTG2,CLDN10,FN1,GNAI3,ITGB2,ITGB7,MMP14,MMP9,MSN,RDX,SDC4,SELL,SELPLG,VCAM1 |
| CTLA4 Signaling in Cytotoxic T Lymphocytes | 2,59 | | 0,167 | B2M,CD80,CD86,CTLA4,HLA-A,HLA-E,HLA-F,JAK2,LAT,LCK,LCP2,PPP2CB |
| ILK Signaling | 2,55 | -1,29099 | 0,133 | ACTB,ACTG1,ACTG2,Cdc42,CFL1,FLNA,FN1,GSK3A,ITGB2,ITGB3,ITGB7,ITGB8,MMP9,MYC,PPP1R14B,PPP2CB,RAC2,RHOH,RICTOR |

| | | | | |
|---|---|---|---|---|
| Mevalonate Pathway I | 2,54 | 2 | 0,4 | HMGCR,HMGCS1,IDI1,MVK |
| Interferon Signaling | 2,54 | -1,13389 | 0,233 | IFI35,IRF1,JAK2,PSMB8,SOCS1,STAT1,TAP1 |
| Th1 Pathway | 2,5 | 0,27735 | 0,146 | CD40LG,CD80,CD86,HLA-A,HLA-DOB,IL18R1,IL6R,IRF1,ITGB2,JAK2,LTA,NFATC3,SOCS1,STAT1,TNFSF11 |
| Protein Kinase A Signaling | 2,46 | -0,75593 | 0,11 | AKAP1,Calm1 (includes others),CAMK2D,CDKN3,CNGA1,DUSP10,DUSP2,DUSP4,DUSP5,DUSP6,FLNA,GNAI3,GNB1,GSK3A,H3-3A/H3-3B,MAP3K1,MTMR3,MYL12B,NFATC3,NFKBIA,PDE3B,PDE4A,PLCB4,PLCL1,PPP1CA,PPP1R14B,PPP3CC,PTPN9,PTPRC,ROCK2,SMPDL3A,YWHAZ |
| Signaling by Rho Family GTPases | 2,43 | -2,84019 | 0,123 | ACTB,ACTG1,ACTG2,ARHGEF1,ARHGEF10,BAIAP2,Cdc42,CDC42EP3,CFL1,GNA15,GNAI3,GNB1,MSN,MYL12B,NCF1,RAC2,RDX,RHOH,ROCK2,SEPTIN1,SEPTIN8,SEPTIN9 |
| Epoxysqualene Biosynthesis | 2,36 | | 1 | FDFT1,SQLE |
| Role of NFAT in Regulation of the Immune Response | 2,35 | -1,69775 | 0,128 | Calm1 (includes others),CD79B,CD80,CD86,CSNK1E,FCGR3A/FCGR3B,GNA15,GNAI3,GNB1,GSK3A,HLA-A,HLA-DOB,LAT,LCK,LCP2,NFATC3,NFKBIA,PLCB4,PPP3CC |
| Role of Macrophages, Fibroblasts and Endothelial Cells in Rheumatoid Arthritis | 2,34 | | 0,113 | APC2,Calm1 (includes others),CAMK2D,FCGR3A/FCGR3B,FN1,FZD5,IL16,IL17RA,IL18R1,IL1RL1,IL6R,IRAK1,JAK2,LTA,LTB,MYC,NFATC3,NFKBIA,PLCB4,PLCL1,PPP3CC,ROCK2,SOCS1,TNFSF11,TRAF1,VCAM1,WNT10A |
| Regulation of Actin-based Motility by Rho | 2,32 | -3 | 0,162 | ACTB,ACTG2,ARHGDIA,BAIAP2,Cdc42,CFL1,MPRIP,MYL12B,PFN1,RAC2,RHOH |
| Actin Cytoskeleton Signaling | 2,31 | -2,82843 | 0,124 | ACTB,ACTG1,ACTG2,APC2,ARHGAP24,ARHGEF1,BAIAP2,CFL1,FLNA,FN1,MPRIP,MSN,MYL12B,PFN1,RAC2,RDX,ROCK2,SSH3,TIAM1,TRIO |
| Estrogen-mediated S-phase Entry | 2,3 | 0,447214 | 0,24 | CCNA2,CDK1,CDKN1B,E2F2,MYC,TFDP1 |
| Role of JAK family kinases in IL-6-type Cytokine Signaling | 2,3 | | 0,24 | IL6R,JAK2,SOCS1,STAT1,STAT5A,STAT5B |
| Graft-versus-Host Disease Signaling | 2,29 | | 0,212 | CD80,CD86,FASLG,HLA-A,HLA-DOB,HLA-E,HLA-F |
| Virus Entry via Endocytic Pathways | 2,29 | | 0,148 | ACTB,ACTG1,ACTG2,B2M,FLNA,HLA-A,HLA-E,HLA-F,ITGB2,ITGB3,ITGB7,ITGB8,RAC2 |
| Natural Killer Cell Signaling | 2,28 | -0,22942 | 0,126 | B2M,CFL1,FASLG,FCGR3A/FCGR3B,HLA-A,HLA-E,HLA-F,HSPA8,IL18R1,IL2RB,JAK2,LAT,LCK,LCP2,MAP3K1,NECTIN2,NFATC3,RAC2,RASSF5 |
| Primary Immunodeficiency Signaling | 2,21 | | 0,206 | CD40LG,IGHM,LCK,PTPRC,TAP1,TNFRSF13B,UNG |
| Remodeling of Epithelial Adherens Junctions | 2,12 | | 0,17 | ACTB,ACTG1,ACTG2,ARF6,EXOC2,TUBA1A,TUBA4A,TUBB2B,TUBB4B |
| Integrin Signaling | 2,08 | -3,15296 | 0,118 | ACTB,ACTG1,ACTG2,ARF6,ARHGAP5,CAPN10,CAPN2,CAPN3,Cdc42,ITGAE,ITGB2,ITGB3,ITGB7,ITGB8,MPRIP,MYL12B,PFN1,RAC2,RHOH,TSPAN3 |
| Systemic Lupus Erythematosus In B Cell Signaling Pathway | 2,03 | -0,8165 | 0,111 | Calm1 (includes others),CCND2,CD40LG,CD5,CD79B,FASLG,IGHM,IL6R,IRAK1,IRF5,JAK2,LCK,LILRB3,LTA,LTB,MYC,NFATC3,PAG1,PPP3CC,RAC2,STAT1,STING1,TNFSF11,TRAF1 |
| Kinetochore Metaphase Signaling Pathway | 1,92 | 2,309401 | 0,138 | AURKB,CDCA8,CDK1,CENPE,H2AZ1,MASTL,NEK2,NUF2,PLK1,PPP1CA,PPP1R14B,REC8 |
| Trans, trans-farnesyl Diphosphate Biosynthesis | 1,9 | | 0,667 | FDPS,IDI1 |

**Supplementary Table 6. Top 50 Canonical Pathways between NCOR1-cKO and WT effector Treg cells**

| Ingenuity Canonical Pathways | -log(p-value) | zScore | Ratio | Molecules |
|---|---|---|---|---|
| Superpathway of Cholesterol Biosynthesis | 16,1 | 3,741657 | 0,583 | CYP51A1,DHCR24,FDFT1,FDPS,HMGCR,HMGCS1,IDI1,LSS,MSMO1,MVK,NSDHL,SC5D,SQLE,TM7SF2 |
| Cholesterol Biosynthesis I | 12 | 3 | 0,75 | E,TM7SF2 |
| dihydrolanosterol) | 12 | 3 | 0,75 | E,TM7SF2 |
| Cholesterol Biosynthesis III (via Desmosterol) | 12 | 3 | 0,75 | E,TM7SF2 |
| LXR/RXR Activation | 5,91 | -0,44721 | 0,148 | ABCA1,ABCG1,ACACA,CYP51A1,FDFT1,HMGCR,IL1RL1,IL1RN,LDLR,NCOR1,SCD,SREBF1 |
| Zymosterol Biosynthesis | 5,64 | 2 | 0,8 | CYP51A1,MSMO1,NSDHL,TM7SF2 |
| TR/RXR Activation | 4,93 | | 0,145 | ACACA,HIF1A,LDLR,NCOR1,PDE3B,PIK3C2B,SCARB1,SREBF1,SREBF2,TBL1XR1 |
| Superpathway of Geranylgeranyldiphosphate Biosynthesis I (via Mevalonate) | 4,88 | 2,236068 | 0,385 | FDPS,HMGCR,HMGCS1,IDI1,MVK |
| Mevalonate Pathway I | 4,06 | 2 | 0,4 | HMGCR,HMGCS1,IDI1,MVK |
| Epoxysqualene Biosynthesis | 3,16 | | 1 | FDFT1,SQLE |
| T Cell Exhaustion Signaling Pathway | 3,12 | -0,33333 | 0,0827 | A,JAK2,NFATC1,PDK1,PIK3C2B,PPM1J,RAP2B,STAT1,TGFBR1 |
| Stearate Biosynthesis I (Animals) | 3,07 | 0,447214 | 0,172 | ACSL1,ACSL3,DHCR24,ELOVL6,MBOAT7 |
| LPS/IL-1 Mediated Inhibition of RXR Function | 2,91 | -1,34164 | 0,078 | ABCA1,ABCG1,ACSL1,ACSL3,HMGCS1,IL1RL1,IL1RN,MGMT,SCARB1,SMOX,SREBF1 |
| Trans, trans-farnesyl Diphosphate Biosynthesis | 2,69 | | 0,667 | FDPS,IDI1 |
| Th1 and Th2 Activation Pathway | 2,31 | | 0,069 | A,IL1RL1,JAK2,LTA,NFATC1,NOTCH2,PIK3C2B,STAT1,TG |
| Th1 Pathway | 2,24 | 0,377964 | 0,0777 | CD86,HLA-A,JAK2,LTA,NFATC1,NOTCH2,PIK3C2B,STAT1 |
| Dendritic Cell Maturation | 2,12 | -0,70711 | 0,0687 | Cd1d2,CD86,HLA-A,IL1RN,IRF8,JAK2,LTA,PIK3C2B,STAT1 |
| p38 MAPK Signaling | 2,11 | -0,37796 | 0,0805 | IL1RL1,IL1RN,MKNK2,MYC,PLA2G4F,STAT1,TGFBR1 |
| Oleate Biosynthesis II (Animals) | 2,02 | | 0,333 | SCD,Scd2 |
| IL-7 Signaling Pathway | 2 | -0,44721 | 0,0857 | BCL6,CDKN1B,MYC,NFATC1,PIK3C2B,STAT1 |
| Oncostatin M Signaling | 1,93 | | 0,118 | EPAS1,JAK2,RAP2B,STAT1 |
| plastidic) | 1,9 | | 0,158 | LCLAT1,MBOAT7,PTPMT1 |
| Influenza | 1,88 | | 0,286 | FDPS,RSAD2 |
| Thrombopoietin Signaling | 1,87 | 1,341641 | 0,0926 | JAK2,MYC,PIK3C2B,RAP2B,STAT1 |
| Transition Pathway | 1,84 | | 0,0621 | FBR1,WNT10A |
| Hepatic Fibrosis Signaling Pathway | 1,79 | 0 | 0,05 | CDKN1B,HIF1A,IL1RL1,IL1RN,JAK2,MYC,NCF1,PDK1,PIK3C2B,RAP2B,ROCK2,TGFBR1,VCAM1,WNT10A |
| Fatty Acid Activation | 1,76 | | 0,25 | ACSL1,ACSL3 |
| Pregnenolone Biosynthesis | 1,76 | | 0,25 | CYP11A1,CYP26B1 |
| Caveolar-mediated Endocytosis Signaling | 1,72 | | 0,0847 | ACTG2,FLNA,FLOT1,HLA-A,ITGAE |
| Cardiac Hypertrophy Signaling (Enhanced) | 1,67 | -0,24254 | 0,0451 | YC,NFATC1,PDE3B,PDK1,PIK3C2B,RAP2B,ROCK2,TGFBR1,WNT10A |
| Histidine Degradation VI | 1,66 | | 0,222 | CYP11A1,CYP26B1 |
| Killer Cells | 1,64 | | 0,0806 | ACTG2,CAMK2D,CD86,HLA-A,LTA |
| Role of Macrophages, Fibroblasts and Endothelial Cells in Rheumatoid Arthritis | 1,62 | | 0,0502 | CAMK2D,IL1RL1,IL1RN,JAK2,LTA,MYC,NFATC1,PIK3C2B,RAP2B,ROCK2,VCAM1,WNT10A |
| Lanosterol Biosynthesis | 1,58 | | 1 | LSS |
| Melatonin Degradation III | 1,58 | | 1 | MPO |
| GM-CSF Signaling | 1,58 | 0,447214 | 0,0781 | CAMK2D,JAK2,PIK3C2B,RAP2B,STAT1 |
| γ-linolenate Biosynthesis II (Animals) | 1,57 | | 0,2 | ACSL1,ACSL3 |
| T Helper Cell Differentiation | 1,56 | | 0,0769 | BCL6,CD86,HLA-A,STAT1,TGFBR1 |
| Neuroinflammation Signaling Pathway | 1,55 | 0,632456 | 0,0507 | A,JAK2,NCF1,NFATC1,PIK3C2B,PLA2G4F,STAT1,TGFBR1,VCAM1 |
| Th2 Pathway | 1,52 | 2,236068 | 0,0614 | CD86,HLA-A,IL1RL1,JAK2,NOTCH2,PIK3C2B,TGFBR1 |
| Chronic Myeloid Leukemia Signaling | 1,51 | | 0,0667 | CDKN1B,HDAC5,MYC,PIK3C2B,RAP2B,TGFBR1 |
| Mitochondrial L-carnitine Shuttle Pathway | 1,49 | | 0,182 | ACSL1,ACSL3 |
| Triacylglycerol Biosynthesis | 1,48 | | 0,111 | ELOVL6,LCLAT1,MBOAT7 |
| Superpathway of Melatonin Degradation | 1,44 | | 0,107 | CYP51A1,MPO,SMOX |
| Ubiquinol-10 Biosynthesis (Eukaryotic) | 1,42 | | 0,167 | CYP11A1,CYP26B1 |
| CNTF Signaling | 1,41 | 1 | 0,0816 | JAK2,PIK3C2B,RAP2B,STAT1 |
| CTLA4 Signaling in Cytotoxic T Lymphocytes | 1,39 | | 0,0694 | CD86,HLA-A,JAK2,PIK3C2B,PPM1J |
| IL-4 Signaling | 1,39 | | 0,0694 | HLA-A,JAK2,NFATC1,PIK3C2B,RAP2B |
| ERK/MAPK Signaling | 1,37 | 0 | 0,0537 | TAT1 |
| PDGF Signaling | 1,37 | 1,341641 | 0,0685 | JAK2,MYC,PIK3C2B,RAP2B,STAT1 |

**Supplementary Table 7. Naive and effector Treg cell gene sets**

| Top 100 genes naive Treg cells | | | Top 100 genes effector Treg cells | | |
|---|---|---|---|---|---|
| gene_name | log2FoldChange | padj | gene_name | log2FoldChange | padj |
| Tnnt2 | -6,985729012 | 2,94E-04 | Vax2 | 18,553409 | 9,41E-08 |
| Sele | -5,815397895 | 4,08E-04 | Slc7a10 | 9,590563625 | 5,53E-13 |
| Lhfp | -5,715068291 | 5,46E-04 | Gm6831 | 9,126315862 | 6,98E-12 |
| Tspan9 | -5,475674133 | 6,04E-12 | Gm8798 | 8,729935349 | 1,89E-10 |
| 2610307P16Rik | -5,382908324 | 1,19E-04 | Ccn4 | 8,721220085 | 6,38E-11 |
| Bach2os | -5,283348372 | 0,001960755 | 4930558J18Rik | 8,596431316 | 2,15E-10 |
| Gm20705 | -5,274020721 | 8,63E-04 | Ky | 8,211810424 | 1,35E-09 |
| Myo5c | -5,15774563 | 0,01969748 | Krt83 | 7,967263901 | 4,83E-09 |
| G0s2 | -5,128987463 | 4,25E-08 | Pvrig | 7,913980147 | 3,60E-06 |
| Gm11662 | -4,993642516 | 0,093988898 | Ces2c | 7,689711393 | 9,43E-41 |
| Gm42899 | -4,921288092 | 0,047019117 | Il1rn | 7,6291296 | 5,31E-08 |
| 1700017M07Rik | -4,91731562 | 0,017728472 | Gm49180 | 7,613065702 | 1,02E-07 |
| Gm16327 | -4,809036528 | 0,012645484 | Lpin3 | 7,60142537 | 5,11E-06 |
| Gm17212 | -4,700966963 | 0,075141625 | Gm5159 | 7,599016545 | 0,016233938 |
| Gm9694 | -4,680099746 | 0,022850289 | Klk1b27 | 7,413290324 | 3,57E-06 |
| Gm37283 | -4,651666521 | 0,027531912 | Ces2d-ps | 7,303171155 | 7,94E-14 |
| Sorcs2 | -4,650747416 | 2,30E-21 | Lrrc49 | 7,293781486 | 1,81E-04 |
| Ampd1 | -4,59148535 | 4,09E-09 | B3galt5 | 7,290302496 | 3,80E-04 |
| Gm14125 | -4,4731621 | 7,08E-40 | Gpr35 | 7,282560213 | 3,09E-04 |
| Gm17936 | -4,461856619 | 3,74E-04 | Olfr60 | 7,250261368 | 3,41E-05 |
| Tex26 | -4,416369674 | 0,074458156 | Fam81a | 7,195136409 | 1,16E-05 |
| Spats2 | -4,310836572 | 0,055451774 | Olfr922 | 7,180143457 | 1,80E-06 |
| Fam186b | -4,168410179 | 0,019997072 | D430019H16Rik | 7,167146648 | 4,94E-04 |
| Igkv10-96 | -4,132900056 | 0,048594269 | 1700001O22Rik | 7,162725165 | 1,03E-06 |
| Gm37982 | -4,048063586 | 0,070510397 | Gm37347 | 7,133764373 | 2,79E-06 |
| Tmem108 | -3,908322116 | 2,07E-10 | Serpina9 | 7,130773006 | 6,81E-04 |
| Cd8a | -3,811084631 | 0,063087197 | Klrg1 | 7,117563814 | 9,12E-27 |
| Cnn3 | -3,805654072 | 6,90E-79 | Gm5541 | 7,032332975 | 5,28E-06 |
| Adcy6 | -3,771213139 | 1,83E-09 | Insrr | 7,001603135 | 9,24E-06 |
| Gm47879 | -3,727075072 | 0,089563752 | Ispd | 6,983288211 | 3,90E-05 |
| Sell | -3,725319559 | 1,55E-50 | Selenom | 6,910920511 | 5,11E-06 |
| Nr6a1os | -3,719151185 | 0,06798522 | Tigit | 6,897979839 | 2,46E-11 |
| Tcrg-C1 | -3,686401449 | 7,86E-27 | Frmd5 | 6,894460244 | 4,63E-05 |
| Ly6c1 | -3,6744317 | 2,12E-42 | Pkhd1l1 | 6,894208913 | 0,003263739 |
| Gm19078 | -3,670723365 | 0,05578776 | Npas4 | 6,889554335 | 5,20E-04 |
| Ly6c2 | -3,65296675 | 1,49E-28 | Krt17 | 6,859568479 | 0,001647124 |
| Cdhr3 | -3,642687227 | 1,34E-06 | Tox2 | 6,855138997 | 4,81E-10 |
| 2810410L24Rik | -3,636026898 | 0,044329457 | Nckap5 | 6,831995654 | 3,08E-05 |
| Ptger3 | -3,631235078 | 0,002479428 | Srxn1 | 6,793843051 | 3,01E-06 |
| Gm36931 | -3,618735192 | 6,97E-05 | Cxcl2 | 6,780191123 | 3,96E-04 |
| Sox4 | -3,499475414 | 2,68E-04 | Asb2 | 6,75065671 | 8,80E-80 |
| Gm4815 | -3,493970216 | 5,92E-08 | Gm42495 | 6,736791549 | 1,10E-05 |
| Igfbp4 | -3,491358135 | 2,37E-57 | Gm49751 | 6,71979095 | 0,006189664 |
| 4933417C20Rik | -3,414955465 | 0,080854279 | Msrb3 | 6,712971138 | 6,72E-05 |
| 4930556H04Rik | -3,412456042 | 0,003446727 | Gm42500 | 6,706150375 | 1,52E-05 |
| Col7a1 | -3,411486497 | 0,028408613 | Col16a1 | 6,651403981 | 0,0037116 |
| Grifin | -3,402720257 | 2,85E-06 | Sostdc1 | 6,619671451 | 0,004721844 |
| Aqp11 | -3,388616903 | 0,001151671 | 2210011C24Rik | 6,597645661 | 2,58E-04 |
| Acvr1c | -3,338835589 | 0,04737226 | C030034L19Rik | 6,596860214 | 9,96E-20 |
| Homer3 | -3,30032529 | 1,64E-07 | Ighd4-1 | 6,59528733 | 1,66E-11 |
| Drc1 | -3,291904536 | 0,025451986 | Ces2b | 6,592051788 | 6,60E-05 |
| C230085N15Rik | -3,18237559 | 3,51E-43 | Ifng | 6,582498202 | 1,45E-05 |
| Amigo2 | -3,15132974 | 2,23E-22 | Krt87 | 6,573721541 | 7,65E-05 |
| Trgj1 | -3,14420678 | 0,055163591 | Myh3 | 6,554119854 | 9,22E-04 |
| Selp | -3,129869412 | 0,002787172 | Zfp612 | 6,476097391 | 5,57E-08 |
| BC049715 | -3,127081484 | 0,026837268 | Gm12169 | 6,37237613 | 3,09E-04 |
| Gm37248 | -3,111696984 | 8,93E-20 | Ccl1 | 6,363108869 | 0,002306453 |
| Dock4 | -3,108731161 | 0,033289594 | Moxd2 | 6,341627856 | 1,52E-04 |
| Gm15708 | -3,073635591 | 1,06E-39 | Cfap126 | 6,338569078 | 3,59E-05 |
| Trim47 | -3,060447154 | 0,097106599 | Tshz2 | 6,323635784 | 0,001006436 |
| Tcf7l2 | -3,051451256 | 0,007454222 | Zan | 6,318085943 | 0,023508636 |
| Dapk1 | -3,048384948 | 6,11E-15 | Arnt2 | 6,316274274 | 1,82E-05 |
| Bach2it1 | -3,038580998 | 0,006649316 | Gm3278 | 6,291998767 | 3,21E-04 |
| Tcrg-C4 | -3,028167605 | 3,07E-16 | Gm47418 | 6,290268378 | 0,018279957 |
| Gm48582 | -2,9934522 | 0,016746755 | Gfra1 | 6,266804264 | 6,32E-05 |
| Gm47523 | -2,99315415 | 0,086353234 | Cyb561 | 6,249242093 | 0,001459263 |

| | | | | | |
|---|---|---|---|---|---|
| Stra6 | -2,990624035 | 6,99E-16 | Metrnl | 6,199670309 | 0,001067773 |
| 4833419F23Rik | -2,985352509 | 0,003349574 | Lrrn3 | 6,177708079 | 4,65E-04 |
| Tubb2a-ps2 | -2,935028496 | 2,80E-05 | Speg | 6,135882553 | 8,88E-04 |
| Cybrd1 | -2,921608848 | 0,052898556 | 5730409E04Rik | 6,134354705 | 0,002339923 |
| Kcnh3 | -2,909554127 | 0,001710858 | Atp6v0d2 | 6,120496452 | 2,23E-49 |
| Tcrg-C3 | -2,894220013 | 0,00466347 | Spock2 | 6,120018134 | 1,29E-13 |
| Gm10130 | -2,883446311 | 3,32E-21 | Alox8 | 6,117972836 | 3,60E-17 |
| Gm20696 | -2,877535488 | 1,75E-24 | Emp1 | 6,082274812 | 0,003456015 |
| Prkar1b | -2,874976214 | 0,010539049 | Mageh1 | 6,056245511 | 0,001276826 |
| Gm15915 | -2,791857807 | 0,054812492 | Ccdc80 | 6,05206848 | 9,87E-04 |
| Gm37266 | -2,784798415 | 0,015444318 | Gm49735 | 6,043203949 | 0,001921038 |
| Tdrp | -2,780247009 | 1,93E-21 | Gm37729 | 6,036342547 | 0,004062624 |
| Satb1 | -2,772942072 | 1,26E-61 | Parpbp | 6,011034282 | 0,001485257 |
| Gm20518 | -2,743709205 | 0,064913946 | Noxa1 | 5,974793124 | 2,05E-07 |
| Gm13546 | -2,735647476 | 1,45E-16 | 4930428O21Rik | 5,970914601 | 0,047971958 |
| Gata1 | -2,708372359 | 2,43E-40 | Il10 | 5,956288869 | 1,36E-06 |
| Plaur | -2,67994701 | 0,066421802 | Pltp | 5,930779537 | 0,013812189 |
| B430306N03Rik | -2,677605434 | 3,50E-10 | Esm1 | 5,929109315 | 0,001327358 |
| Gm37176 | -2,675416212 | 0,001988512 | Fam69c | 5,917354027 | 0,096739942 |
| Col11a2 | -2,661698342 | 2,14E-10 | Gm11730 | 5,91322592 | 7,58E-04 |
| 1700025G04Rik | -2,65934046 | 9,66E-08 | Tlr4 | 5,896308039 | 0,006563585 |
| Art4 | -2,649009561 | 2,62E-06 | Ankrd29 | 5,892503704 | 1,88E-05 |
| Treml2 | -2,641551044 | 6,96E-36 | Fgl2 | 5,886327549 | 6,60E-22 |
| Gm21994 | -2,640316532 | 1,26E-23 | Dkkl1 | 5,879279958 | 2,13E-04 |
| Tcrg-C2 | -2,632572557 | 3,94E-07 | Gm37759 | 5,861688336 | 0,001406004 |
| Gm37593 | -2,616132294 | 1,19E-06 | Slc24a3 | 5,829535655 | 0,002878086 |
| Zcchc24 | -2,605294669 | 0,042893263 | Npdc1 | 5,815169809 | 4,08E-04 |
| Tubb2b | -2,602010161 | 3,83E-17 | Vdr | 5,805663342 | 0,015353726 |
| Gm49066 | -2,594480134 | 0,016373515 | Chst2 | 5,803925441 | 6,63E-07 |
| Emid1 | -2,585664057 | 1,41E-05 | Nxnl2 | 5,781235157 | 0,002460744 |
| Pyroxd2 | -2,571318669 | 0,061622166 | Mid2 | 5,766234584 | 0,013339966 |
| Gm45155 | -2,560526509 | 0,002772742 | Gstt3 | 5,763522935 | 0,002108239 |
| Atp1b1 | -2,551907176 | 5,47E-37 | Vegfc | 5,758299123 | 0,009104042 |
| Gm14124 | -2,522007838 | 1,45E-04 | Nsun7 | 5,752899031 | 0,002306453 |
